# Systematic Discovery and Design of Synthetic Immune Receptors in Plants

**DOI:** 10.1101/2025.03.17.642524

**Authors:** Bruno Pok Man Ngou, Michele Wyler, Marc W. Schmid, Takehiro Suzuki, Naoshi Dohmae, Yasuhiro Kadota, Ken Shirasu

**Author notes:** These authors contributed equally to this work (Michele Wyler, Marc W. Schmid).

## Abstract

Plants deploy a diverse array of pattern recognition receptors (PRRs) that perceive microbe-associated molecular patterns (MAMPs) to activate immune responses. Leucine-rich repeat receptor-like kinase subgroup XII (LRR-RLK-XII) represents one of the largest PRR families due to lineage-specific diversification. Bioinformatics, synthetic biology, and biochemical approaches were integrated to characterize PRRs from 285 plant species. We identified a receptor, named “SCORE”, that perceives cold-shock protein (CSP) peptides. SCORE orthologs from multiple angiosperm lineages exhibit remarkable CSP recognition polymorphisms, indicating recurrent selection for pathogen recognition through substitutions at key amino acid residues. Through functional phylogenomics and protein structure predictions, we generated synthetic SCORE variants capable of detecting multiple phytopathogen CSP peptides, thus revealing the diverse PRR recognition landscape in plants. Our strategy hold promises for engineering plant immune receptors, particularly for perennial crops.

## Main Text

Living organisms perceive and respond to environmental stimuli often *via* specialized cell-surface localized receptors (*1*). Receptors that sense microbe-associated or pathogen-associated molecular patterns (MAMPs/PAMPs) are collectively termed “pattern recognition receptors” (PRRs) (*2*). In plants, PRRs are classified as either receptor-like kinases (RLKs) or receptor-like proteins (RLPs), which lack kinase domains (*3*). RLKs and RLPs with leucine-rich repeat ectodomains (LRR-RLKs and LRR-RLPs) represent the two largest PRR families (*4*). LRR-RLKs are further subdivided into 20 subgroups according to their kinase domain sequences (*5, 6*). Of these, subgroup XII (LRR-RLK-XIIs) mediates MAMP perception and immunity. Notable examples include the *Arabidopsis thaliana* flagellin receptor FLS2, elongation factor receptor EFR, and the tyrosine-sulphated protein receptor XA21 from rice (*7–9*). Upon binding their cognate peptide ligands via the extracellular LRR domain, these receptors recruit somatic embryogenesis receptor-like kinases (SERKs) which, in turn, trigger immune signaling cascades (*10, 11*).

Over the course of land plant evolution, LRR-RLK-XIIs have expanded in a lineage-specific manner, yielding considerable receptor diversity. While the *A. thaliana* genome encodes only ten LRR-RLK-XIIs, many other plant genomes harbor hundreds of copies (*12*). These lineage-specific receptors can be transferred between plant species, as exemplified by the introduction of EFR into multiple crop species, thus conferring broad spectrum bacterial resistance (*13, 14*). Yet, despite thirty years of research, fewer than ten LRR-RLK-XII receptor-ligand pairs have been characterized (*15*). Identifying receptor targets in non-model plant species remains particularly challenging due to technical limitations (*16*). Here, we describe a strategy to map the recognition landscape of LRR-RLK-XIIs in plants, paving the way for the discovery of novel PRRs that can be synthetically engineered for improving disease resistance.

### Clustering of LRR-RLK-XIIs

Recently, 13,185 LRR-RLK-XIIs were identified from 350 plant genomes (*12*). To streamline their characterization, we clustered closely related receptors that likely perceive and bind the same ligand. In LRR-RLKs, the inner LRR residues are critical for peptide perception and thus determine ligand specificity; consequently, these residues tend to be conserved among receptors that share ligand specificity (clusters). To identify these clusters, we utilized repeat conservation mapping (RCM) (*17*). Specifically, we aligned the ectodomains of all 13,185 LRR-RLK-XIIs, then extracted the residues in the inner LRR motifs. Consensus residues (i.e. leucines), were removed, and the remaining non-consensus residues were aligned (Fig. 1A). Conservation of non-consensus residues within a cluster implies a higher likelihood of shared ligand specificity. Clusters with low conservation scores are unlikely to perceive similar ligands and thus were subdivided (Fig. 1B). Iterative clustering and RCM analysis produced thousands of receptor clusters. From these, we curated 210 receptor subgroups that represent 4,177 LRR-RLK-XII (31.7% of all identified receptors) across 285 plant species (Fig. 1C-E; Fig. S1, 2). Most receptor subgroups are lineage-specific and less than five percent are conserved across angiosperms (Fig. 1F; Fig. S3, 4), underscoring the lineage-specific diversification of LRR-RLK-XIIs.

**Fig. 1.**
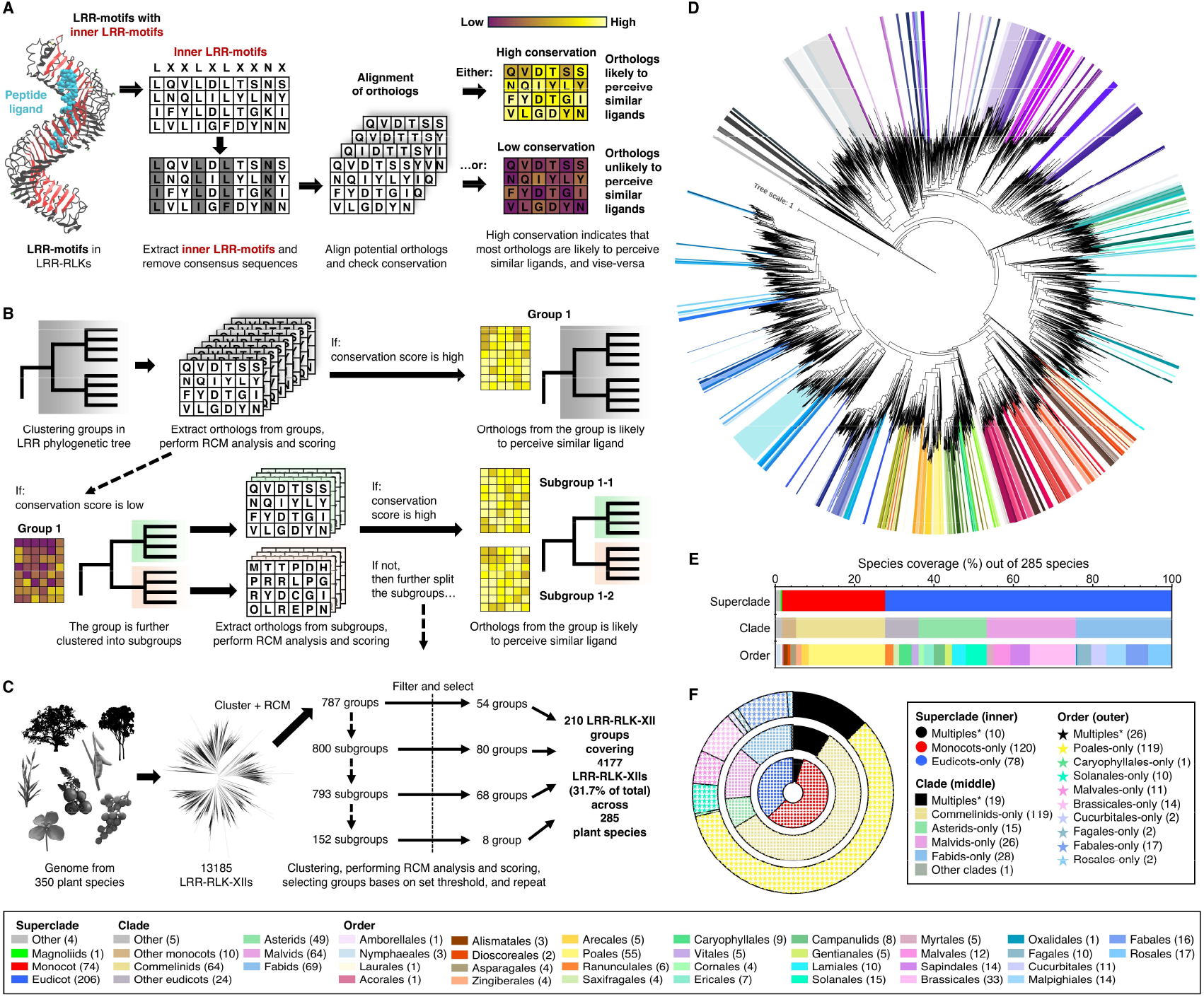
Clustering of LRR-RLK-XIIs through repeat conservation mapping (RCM). **(A)** Schematic of the RCM process. LRR motifs of LRR-RLKs are aligned, masking conserved residues to focus on non-consensus residues. Conservation levels in these alignments infer ligand recognition similarities among orthologs. **(B)** Groups with high conservation scores are retained, and low scoring groups are subdivided and iteratively re-analyzed until stable, high conservation subgroups are identified. **(C)** 4,177 receptors comprising 210 groups from 13,185 LRR-RLK-XII genes across 350 plant genomes were selected via iterative RCM for further screening. **(D)** Phylogenetic tree of LRR-RLK-XII ectodomains, highlighting the 210 groups as colored clusters. **(E)** Species distribution of the 285 analyzed plant species, categorized by superclade, clade, and order. **(F)** Lineage specificity of the 210 selected LRR-RLK-XII groups, with receptors classified as superclade-, clade-, order-specific (colored, inner/middle/outer ring), or distributed across multiple taxa (black). For detailed RCM analysis and methods, refer to methods and supplementary Fig 1-4.

### Exploring the LRR-RLK-XII recognition landscape to identify novel receptors

Our goal was to identify receptors that activate immune responses to bacterial infection among these 210 LRR-RLK-XIIs, through transient expression system in *Nicotiana benthamiana*. Because exogenous LRR-RLK-XIIs could be masked by endogenous PRRs, we generated chimeric receptors by replacing the XII kinase domain with the Xb kinase domain from the Brassinosteroid insensitive 1 (BRI1) receptor. When a ligand is detected, the Xb kinase domain activates responses, such as dephosphorylation of BRI1-EMS-SUPPRESSOR 1 (BES1) (*18, 19*). These developmental outputs are not masked by immune responses, making them ideal indicators of activation from exogenously expressed receptors (*4*) (Fig. 2A). We generated EFR-BRI1, FLS2-BRI1, CORE-BRI1, and MIK2-BRI1 chimeras and confirmed their activation in response to corresponding peptide ligands, full length MAMPs, boiled bacterial suspensions, and bacterial suspensions, as evidenced by BES dephosphorylation (Fig. 2B-E; Fig. S5). Next, we screened all 210 LRR-RLK-XII-BRI1 chimeras with *Agrobacterium tumefaciens* (*Agro*) suspensions and identified seven receptor candidates (Fig. 2F). Receptor candidate 181, an LRR-RLK-XII from *Citrus maxima* (pomelo), was activated by both live and boiled *Agro* suspensions (Fig. 2G), and 181-BRI1 responded to boiled suspensions of multiple bacterial species (Fig. 2H). It is also phosphorylated and interact with the SERK co-receptor, BAK1, in response to boiled *Agro* (Fig. 2I).

**Fig. 2.**
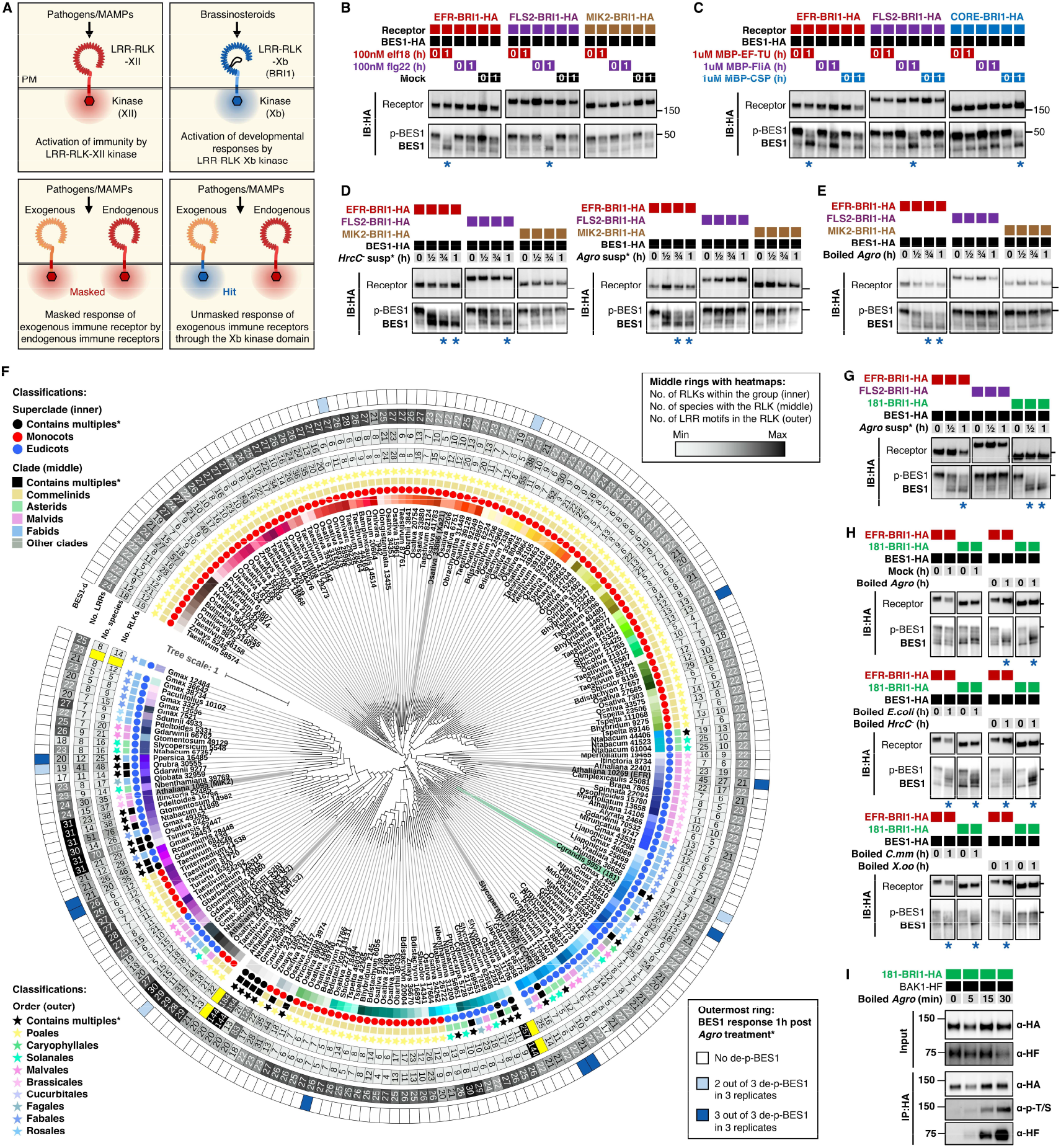
Screening for *Agrobacterium*-responsive LRR-RLK-XII receptors. **(A)** Screening strategy using BRI1 chimeras. LRR ectodomains from 210 LRR-RLK-XII receptors were fused with BRI1 and screened in *N. benthamiana* for BES1 dephosphorylation in response to pathogens or MAMPs. **(B-E)** Validation of BES1 dephosphorylation using known receptor chimeras (EFR-BRI1, FLS2-BRI1, MIK2-BRI1 or CORE-BRI1) treated with the corresponding peptides (**B**), full length MAMP proteins (**C**), live bacterial suspensions (**D**), or boiled bacterial extracts (**E**). Samples were collected at indicated time points and the dephosphorylation of BES1 was assayed by immunoblotting. Asterisks indicate dephosphorylation of BES1. **(F)** Screening results for 210 LRR-RLK-XIIs. Innermost colored blocks correspond to colored clusters in Fig. 1D. Superclade-/Clade-/Order receptor specificity is indicated in the inner ring. The number of LRR-RLK-XIIs within each group, the number of plant species within each group and the number of LRR motifs in the representative receptor in each group are indicated by the middle heatmap rings. The outermost ring represents screening results. Dark blue indicates repeated BES1 dephosphorylation with *A. tumefaciens* (*Agro*). Highlighted are novel receptor 181 from *Citrus grandis* (green) and known receptors (grey). **(G)** Confirmation of 181-BRI1 response to *Agro* suspensions. EFR-BRI1 acts as a positive control and FLS2-BRI1 is a negative control. **(H)** 181-BRI1 respond to boiled *Agro, E*.*coli, P. syringae pv. tomato DC3000 hrcC*^*-*^ (*HrcC*^*-*^), and *C. michiganensis michiganensis* (*Cmm*) suspensions, but not to *X. oryzae pv. oryzicola* (*Xoo*) suspensions. EFR-BRI1 acts as a positive control. (**I**) Immunoprecipitation of 181-BRI1. 181-BRI1 interacts with BAK1 and is phosphorylated upon *Agro* treatment. Coomassie blue staining for immunoblots are shown in supplementary Fig 5.

### SCORE perceives cold-shock proteins

As proof-of-concept for our strategy of characterizing novel LRR-RLK-XIIs, we identified the unknown ligand perceived by 181. The receptor responded to filtrates that passed through spin columns with a cutoff larger than 10 kDa, indicating that the corresponding MAMP is approximately 5-10 kDa (Fig. 3A). In addition, proteinase K treatment abolishes its MAMP activity, suggesting that 181 perceives a protein ligand (Fig. 3A). To identify that ligand, we fractionated the *Agro* suspension by anion exchange chromatography followed by size-exclusion chromatography. The most active fractions, A8 and A9, contain proteins between 5-10 kDa with a relatively high pI value (Fig. 3B, C; Fig. S6). Fractions A6-A9 were analyzed by mass spectrometry for proteins that were more abundant in fractions A8 and A9 compared to fractions A6 and A7. Surprisingly, three of the top five candidates were cold shock proteins (CSPs) (Fig. 3D; Fig. S7). CSPs contain RNA recognition motifs (RNP) and play critical roles in translation and cold tolerance (Fig. 3D) (*20*). These proteins are attractive candidates as PRR ligands because they are conserved and highly expressed in bacteria. Indeed, a Solanaceous-specific LRR-RLK-XII, known as CORE (COld shock protein REceptor) perceives a conserved 22-amino-acid peptide from bacterial CSPs (csp22) (Fig. 3D) (*21, 22*).

**Fig. 3.**
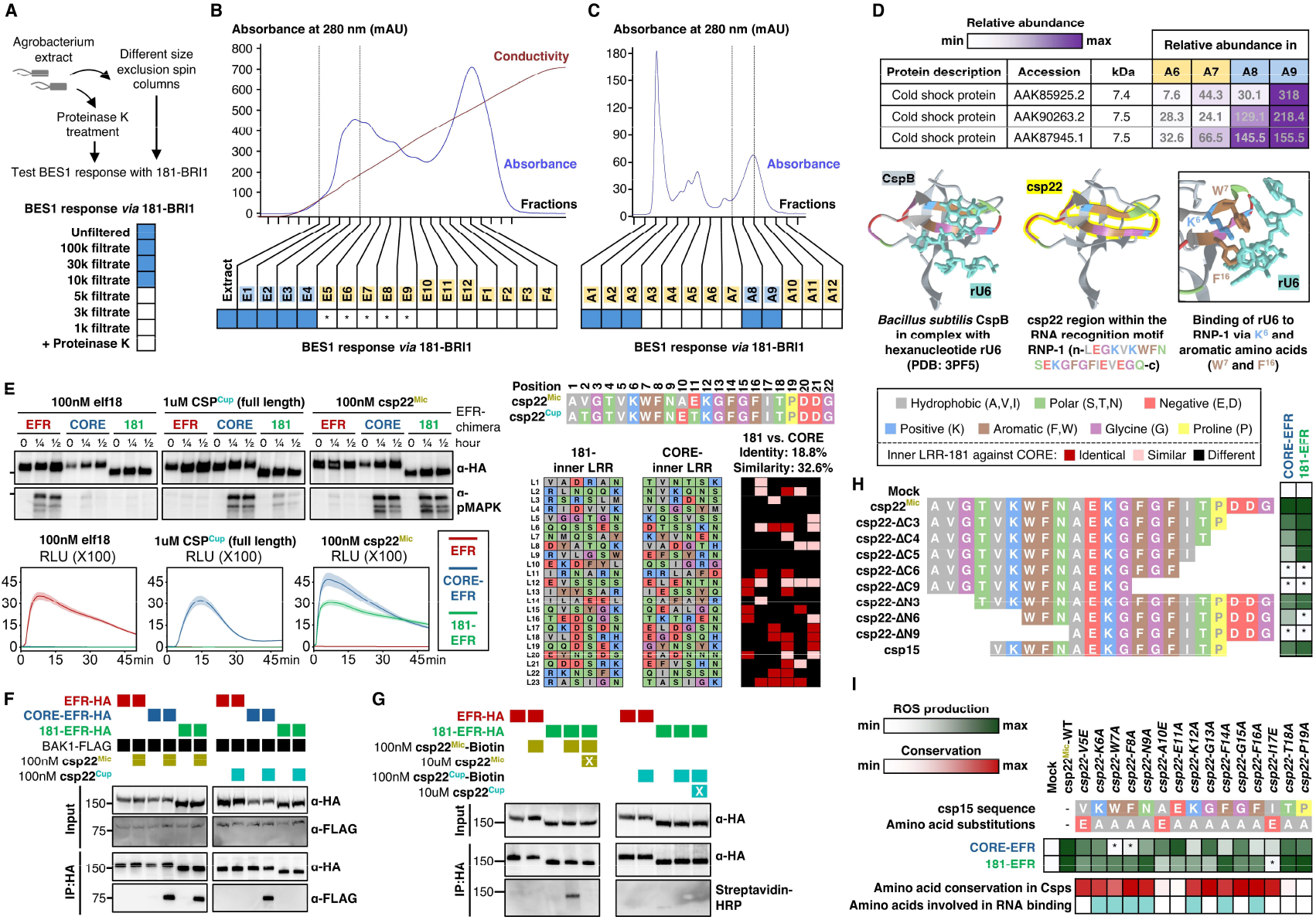
The novel receptor SCORE perceives cold-shock protein (CSP) peptides. **(A)** *Agro* extracts were treated with Proteinase K or fractionated using size exclusion spin columns, and the resulting fractions were tested for activity against 181-BRI1. **(B)** *Agro* extracts were fractioned through anion exchange chromatography, and the activities of each fraction were tested with 181-BRI1. **(C)** Active fractions (E1-E4) from (**B**) were pooled and further separated by size exclusion chromatography, with each fraction tested for 181-BRI1 activity. In (**A-C**), blue-filled squares indicate BES1 dephosphorylation, asterisk (*) indicates week dephosphorylation, and unfilled squares indicate no dephosphorylation. **(D)** Mass spectrometry analysis of fractions A6-9 from (**C**). Cold shock proteins (CSPs) were highly abundant in fractions A8 and A9. The structure of the *Bacillus subtilis* CSP (CspB) complexed with the hexanucleotide rU6 is shown below. Within the RNA recognition motif (RNP), the canonical csp22 peptide is highlighted in yellow. Amino acids are color-coded by properties as indicated in the accompanying box. **(E)** Full-length EFR, CORE-EFR and 181-EFR chimeras were expressed in *N. benthamiana* leaves and treated with 100nM elf18, 1uM full-length CSP from *Cupriavidus* (CSP^Cup^), or 100nM csp22 from *Micrococcus* (csp22^Mic^). MAPK activation and ROS production confirmed activation of the corresponding EFR chimeras. The sequences of csp22^Mic^ and csp22^Cup^ are shown with positions numbered above. Inner LRR residues of 181 and CORE are shown below. Alignment of inner LRR residues between 181 and CORE is also shown with identical residues (dark red), residues with similar properties (light red), and differing residues (black). **(F)** Immunoprecipitation of EFR, CORE-EFR and 181-EFR. 181-BRI1 interacts with BAK1 only in the presence of csp22^Mic^, but not csp22^Cup^, while CORE-BRI1 interacts with BAK1 in the presence of either peptide. **(G)** Immunoprecipitation of EFR and 181-EFR after crosslinking with ethylene glycol bis(succinimidyl succinate) (EGS). 181-BRI1 can be cross-linked to csp22^Mic^-biotin, but not to csp22^Cup^-biotin. EFR is the negative control for both **(F)** and **(G). (H)** CORE-EFR and 181-EFR chimeras expressed in *N. benthamiana* leaves were treated with 100nM of corresponding peptides, listed at left. ROS production is visualized as green heatmaps with an asterisk (*) indicating no significant ROS production compared to mock treatment. **(I)** CORE-EFR and 181-EFR chimeras expressed in *N. benthamiana* leaves were treated with 100nM of corresponding peptides shown on top. ROS production is represented in green heatmaps with an asterisk (*) indicating no significant ROS production compared to the mock treatment. The red heatmap shown below represents the conservation score of each residue in csp15, and cyan blocks indicate residues involved in RNA binding by CSPs. Data related to Fig 3 are shown in supplementary Fig 6-12.

To determine if 181 perceives CSPs, we generated receptor chimeras by fusing CORE and 181 ectodomains to the intracellular domain of EFR. We then expressed full length EFR, CORE-EFR and 181-EFR in 4-week-old *N. benthamiana*. Immune responses were tested using elongation factor peptide (elf18; perceived by EFR), full length CSP from *Cupriavidus* (CSP^Cup^), and canonical csp22 peptide from *Micrococcus* (csp22^Mic^). While CORE-EFR responded to both CSP^Cup^ and csp22^Mic^, 181-EFR was specifically activated by csp22^Mic^, but not by CSP^Cup^. These results were confirmed by BAK1 interaction assays and *in vivo* binding tests with biotinylated csp22^Cup^ and csp22^Mic^ (Fig. 3F, G; Fig. S8, 9). CORE and 181 are not closely related and have evolved independently to perceive csp22 (Figure 2C). Given that these receptors share only 18.8% identity and 32.6% similarity in their inner LRR surfaces, we propose that they differ in their sensitivity to csp22 variants (Fig. 3E; Fig. S10). CORE was more sensitive to C-terminus truncations in csp22, while 181 was more sensitive to N-terminus truncations (Fig. 3H; Fig. S11). Both receptors recognized a core 15-amino acid peptide (csp15) derived from the central region of csp22. CORE was also sensitive to alanine substitutions of conserved aromatic amino acids in csp22, whereas 181 was sensitive to alanine substitutions at positions 10-13 (Fig. 3I; Fig. S12). These results are consistent with the differential recognition of csp22^Cup^ versus csp22^Mic^, which differ at positions 10 and 11 (Fig. 3E). Since positions 10 and 11 are highly variable among CSPs, we named 181 “Selective COld shock protein Receptor” (SCORE).

### Diverse recognition specificity of SCOREs towards CSP variants

CSPs are found in the genomes of viruses, archaea, bacteria, protozoa, plants, fungi and animals, albeit with varying prevalence (Fig. 4A). Specifically, 87-99% of the available bacterial, protozoan, plant, and animal genomes contain at least one CSP, whereas CSP frequency in viruses, archaea and fungi ranges from 1-55% (Fig. 4A). Most of the residues within csp15 are highly conserved, except for those corresponding to positions 10, 11, 18 and 19 in csp22 (or 6, 7, 14 and 15 in csp15) (Fig. 4A), and some residues are enriched in a kingdom-specific manner at positions 6 and 7 in csp15. To capture the breadth of CSP diversity, we synthesized 103 of the most prevalent csp15 peptides, representing 73% (35,881) of all searched species with CSPs (Fig. S13). We also traced the origin and evolutionary trajectory of SCORE in plants, and identified more than 60 potential SCORE orthologs, primarily within the Malvids clade (orders Myrtales and Sapindales), but also in Oxalidales, Ranunculales and Magnoliidae (Fig. 4B, C; Fig. S14-16). To determine if these potential orthologs truly perceive CSPs, we cloned 20 of their ectodomains into EFR chimeras and assayed their immune responses against the 103 synthesized csp15 peptides (Fig. 4C, D). Thirteen of them (including the pomelo SCORE, or “CM”) robustly recognized various csp15 peptides, four responded weakly, and four others gave no response (Fig. 4D; Fig. S17). Notably, the Magnoliidae species *Drimys winteri* encodes a functional SCORE (designated as “DW”), suggesting that SCORE evolved in the last common ancestor of core angiosperms but was subsequently lost from multiple angiosperm lineages. Importantly, SCORE orthologs exhibit extensive polymorphisms in csp15 recognition, likely due to variability at positions 6 and 7 in csp15 (Fig. 4D). Based on these differences, SCOREs fall into three main categories: (1) those favoring csp15 peptides with negatively charged residues at positions 6 and 7 (e.g. *Punica granatum* ortholog, designated as “PG”), (2) those preferring peptides lacking negatively charged residues at these positions (e.g. CM), and (3) those that perceive most csp15 peptides regardless of the residues at these positions (e.g. CORE) (Fig. 4D).

**Fig. 4.**
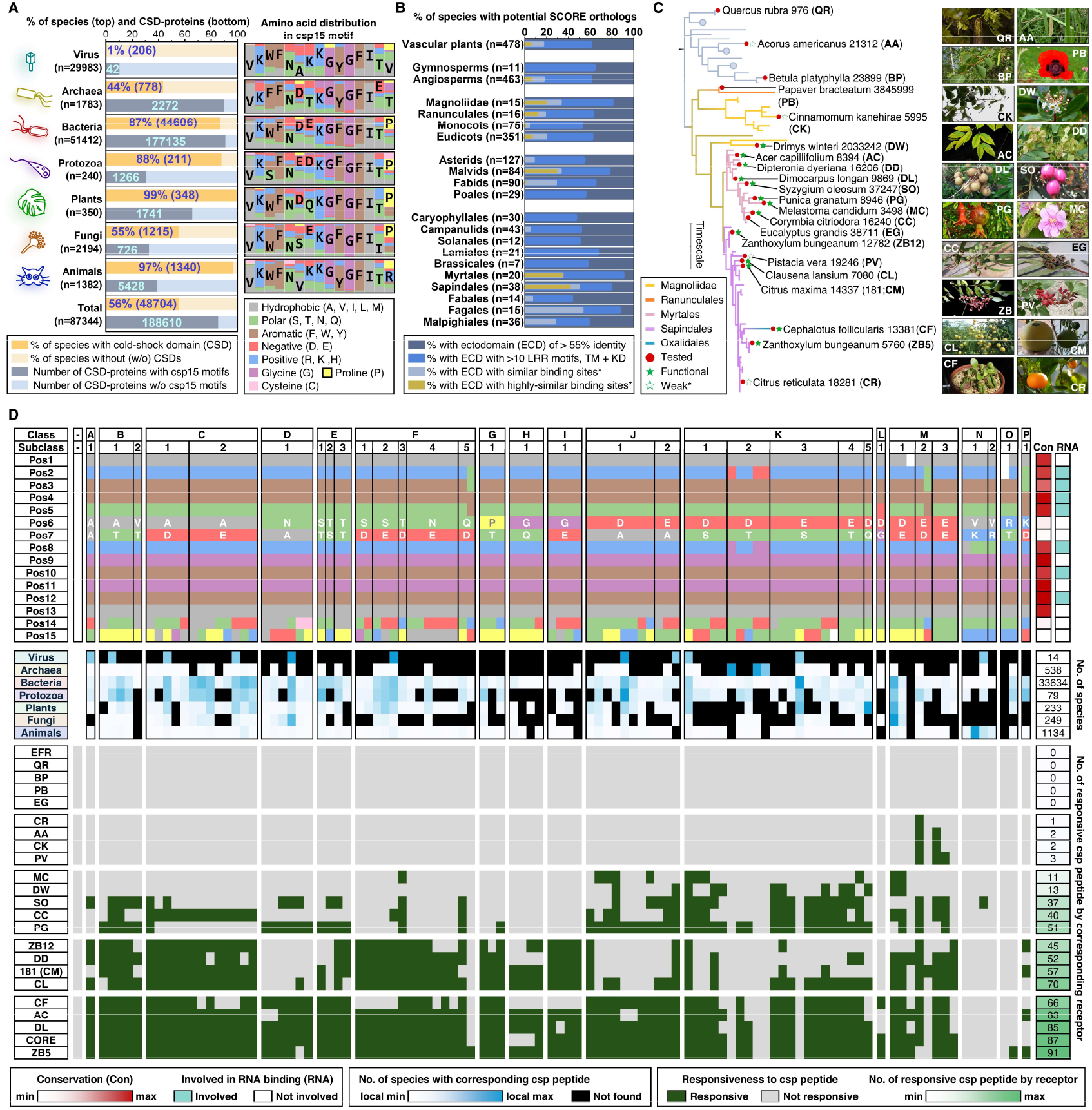
Diverse recognition specificity of SCOREs for CSP variants. **(A)** Prevalence of CSPs across kingdoms. CSPs are defined as proteins containing at least one cold shock domain. (Left) Yellow bars represent the percentage (%) of species in each kingdom encoding at least one CSP. Dark grey bars represent the number of CSPs with csp15 motifs, and light grey bars represent the number of CSPs without csp15 motifs. (Right) All csp15 motifs from CSPs in each kingdom were extracted and aligned. The right bars represent csp15 motif amino acid distributions for each kingdom. Amino acids are categorized by properties, as indicated in the lower right box. **(B)** Distribution of potential SCORE orthologs across land plant lineages. Potential SCORE orthologs were identified through sequential filtering with multiple criteria. Criterion: 1. receptors with more than 55% identity to SCORE (dark blue), 2. receptors from criterion 1 with at least 10 LRRs, a transmembrane domain (TM) and kinase domain (KD) (cerulean), 3. receptors from criterion 2 with binding sites similar to csp15 as predicted by Alphafold2 (*39*) (light blue; *see Fig. 5B and methods), 4. receptors from filter 3 with highly similar csp15 binding sites compared to *C. maxima* SCORE (gold). The overlaying-colored bars represent the percentage of species containing the corresponding receptor type for each filter. *n* represents the total species with receptors identified in filter 1 (set to 100%). For details, refer to supplementary Fig 14-15. **(C)** A simplified phylogenetic tree of SCORE orthologs with csp15 binding sites that are highly similar to CM (gold category in **B**). Branches are labelled according to taxonomic order. Red dots mark the 21 tested SCORE orthologs. Green stars indicate functional CSP receptors, hollow stars denote weak CSP receptors, and unmarked orthologs do not respond to CSPs. **(D)** The top chart represents the 103 csp15 peptides that were synthesized in this study. The amino acid properties at each position (Pos) are indicated with corresponding colors used in Fig. 4A. Peptides are grouped into classes and subclasses based on residues at positions 6 and 7. The red heatmap (right) represents residue conservation scores, and cyan blocks mark RNA binding residues. The blue heatmap (middle) represents the prevalence of the 103 csp15 peptides in each kingdom. Black blocks indicate peptides absent from every species within a kingdom. The number of species searched in each kingdom is shown on the right. Bottom chart represents the CSP recognition capacity of SCOREs in plants. EFR, CORE-EFR and 21 SCORE orthologs (expressed as EFR kinase chimeras) were expressed in *N. benthamiana* leaves. The leaves were then treated with 100nM of corresponding peptides (top). ROS production was then measured for 15 minutes. A peptide was considered active if ROS exceeded the baseline by 2.5x (green). Non-responsive peptides are labelled in grey. EFR served as a negative control and CORE acts as a positive control. Green heatmap (right) indicates the number of responsive peptides (out of 103) for each receptor. Data related to Fig 4 are shown in supplementary Fig 13-17. Acknowledgements and Creative Commons licenses for the plant images in (C) are provided in supplementary text.

### Mechanistic insights into SCORE-csp15 interactions

To understand SCORE ligand specificity, we generated Alphafold2 (AF2) models of SCORE (CM) bound to the csp15^Mic^ peptide (Fig. 5A; Fig. S18) (*23*). AF2 models predict that the 3^rd^-16^th^ LRR motifs of SCORE contribute to csp15 binding. Notably, the N-terminus of csp15 peptide interacts with the 13^th^-16^th^ LRR motifs of SCORE, while the C-terminus interacts with the 3^rd^-5^th^ LRR motifs (Fig. 5A). There are three main interaction sites. First, the “GFGF” motif in csp15 interacts with a highly conserved “QSEMQSY” segment located in the 6^th^-7^th^ LRR motif. Second, the N-terminal valine and lysine residues of csp15 interact mostly with glycine, negatively charged, and aromatic residues in the 13^th^-16^th^ LRR motifs. These negatively charged and aromatic residues are also highly conserved in SCOREs. Third, the two variable amino acids in positions 6 and 7 of csp15 (alanine and glutamate in csp15^Mic^) interact with multiple residues within the 8^th^-11^th^ LRR motifs (Fig. 5A, B; Fig. S19). Because these residues exhibit high variability across SCORE orthologs, they likely are responsible for the diverse recognition specificities of SCOREs toward different CSP peptides (Fig. 5B; Fig. S19).

**Fig. 5.**
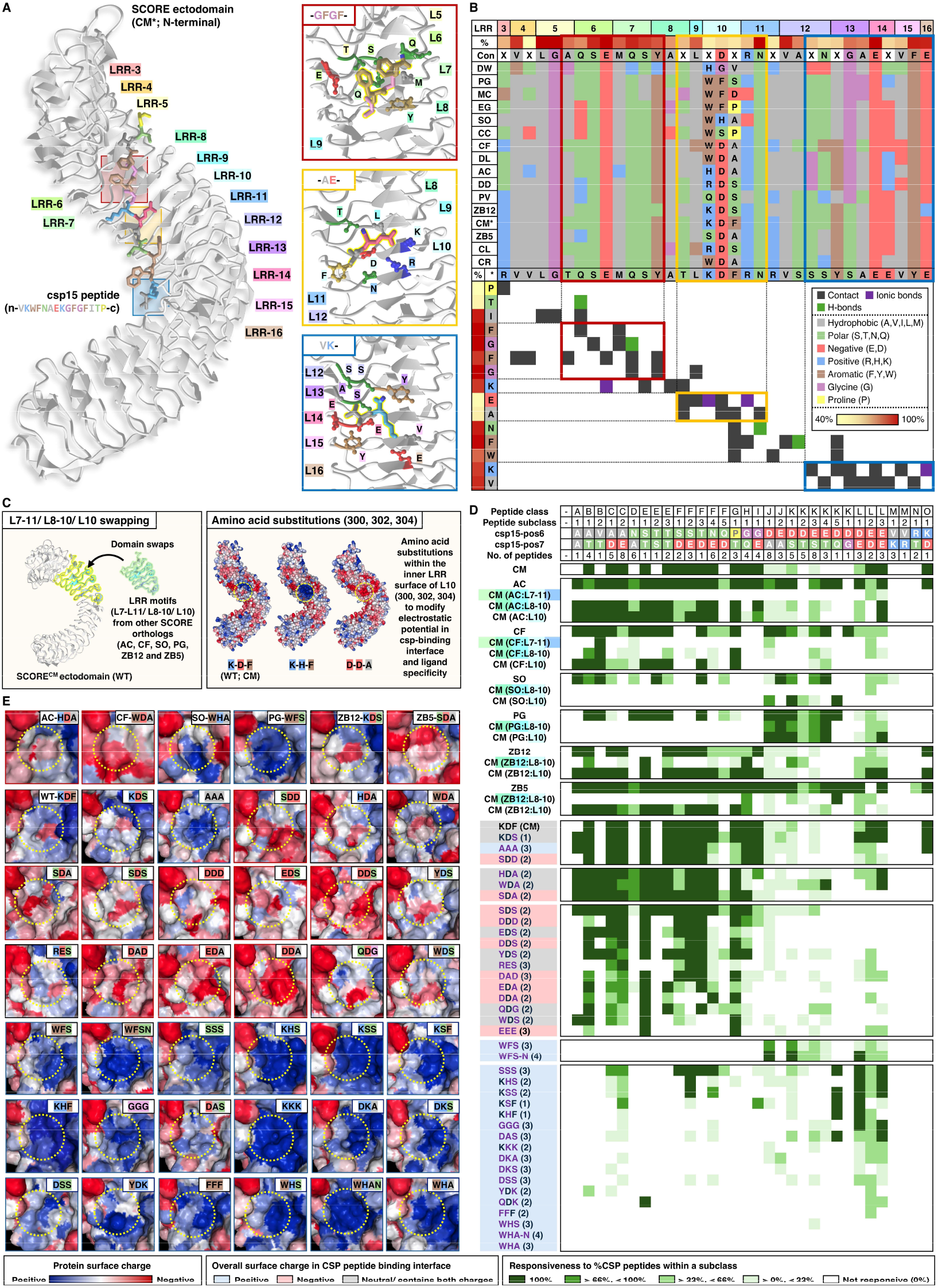
Mechanistic insights into SCORE-csp15 interactions. **(A)** Protein structure prediction of SCORE(CM)-csp15 interaction by Alphafold2 (*23*). LRR motifs involved in csp15 interactions (3^rd^-16^th^) are labelled, with three main interaction sites boxed in red, yellow, and blue. SCORE residues involved in csp15 interactions are labelled in the box on the right. **(B)** Interaction map of SCORE and csp15. Top chart represents the alignment of residues involved in csp15 binding across functional SCORE orthologs. Amino acids are color-coded by properties, as indicated in the box below. Bottom chart represents contact sites between SCORE (*C. maxima*, CM) and csp15, as indicated by colored blocks (right). Contact sites within red, yellow, and blue rectangles correspond to the boxes in (**A**). **(C)** Generation of synthetic SCORE receptors. Left box refers to the swapping of 10^th^, 8^th^-10^th^, 7^th^-11^th^ LRR motifs (L) from different SCORE orthologs into SCORE (CM). Right box refers to the amino acid substitutions at positions 300, 302, or 304 in SCORE (CM). **(D)** CSP recognition specificity of SCORE variants. 103 csp15 peptides were grouped into classes and subclasses according to the amino acids at positions 6 and 7 (top). CM and its variants were expressed in *N. benthamiana* leaves, treated with 100nM of corresponding peptides, and ROS production was measured for 15 minutes. Peptides were deemed active if ROS production exceeded baseline by 2.5x. Peptide subclasses activating a given receptor are indicated as green (darker green indicates more active peptides within a subclass). Non-responsive peptide subclasses are shown in white. Labels of receptor variants: for domain swapping variants, CM, AC, CF, SO, PG, ZB5 and ZB12 refer to the original SCORE orthologs tested in Fig. 4D. CM with different LRR motifs from other orthologs are indicated. For example, CM(AC:L7-11) refers to CM exchanged with the 7^th^-11^th^ LRR motifs from AC. Substitution variants are labelled by amino acid changes at positions 300, 302, or 304 in CM. For example, KDF(WT) refers to the original CM, while AAA refers to CM with residues A, A, A at positions 300, 302, and 304 in CM. WHAN and WFSN have an N residue at position 305 in CM (instead of F). Variants are color-coded by their overall surface charge as indicated in the box below. Amino acid substitutions are indicated in purple and the number of substitutions is shown in parentheses. **(E)** Surface charge of in the inner LRR surface between 8^th^-11^th^ LRR in SCOREs. Protein structures of SCORE orthologs and variants were predicted by Alphafold3 (*39*). Surface charges were calculated by protein-sol (*24*). Labels for receptors are as described in (**C**). Data related to Fig 5 are shown in supplementary Fig 18-24.

To validate these findings, we assessed if csp15 recognition specificity could be transplanted between SCORE orthologs by exchanging their LRR motifs. Specifically, we swapped the 7^th^-11^th^, 8^th^-10^th^ and 10^th^ LRR motifs of multiple orthologs into CM and tested the resulting chimeras (Fig. 5C). Surprisingly, the 7^th^-11^th^ LRR swaps were mostly non-functional, but some 8^th^-10^th^ and 10^th^ LRR swaps altered csp15 recognition profile in wildtype (WT) CM (Fig. 5D; Fig. S22). Moreover, the single 10^th^ LRR swap resulted in a more pronounced change in ligand specificity than the 8^th^-10^th^ LRR swap. Because the three key residues (300^th^, 302^nd^, and 304^th^ in CM) within the 10^th^ LRR are highly variable among SCORE orthologs, we conclude that the 10^th^ LRR motif predominantly governs csp15 recognition specificity in SCOREs (Fig. 5D).

### Synthetic SCORE designs with novel recognition specificities

We engineered synthetic SCORE variants capable of recognizing CSPs from plant pathogens and pests. Residues 6 and 7 of csp15 form ionic bonds with the 8^th^-11^th^ LRR motifs in SCORE, suggesting that the surface charge of the SCORE inner LRR region influences ligand specificity. Using AF3 models of functional SCORE orthologs, we analyzed their surface charges with protein-sol (*23, 24*), and found that, although most charges within the SCORE ectodomain surface are conserved, the inner LRR surface between the 8^th^-10^th^ LRR varies across SCORE orthologs (Fig. 5C, E; Fig. S20, 21, 23). For instance, SCOREs that preferentially recognize csp15 peptides with negatively charged residues at positions 6 and 7, such as PG, generally display a positively charged surface in this region (e.g. PG) (Fig. 5E; Fig. S20). By contrast, SCOREs recognizing csp15 peptides lacking negatively charged residues in these positions typically exhibit neutral or mixed charges (e.g. CM), whereas SCOREs that detect most csp15 variants possess predominantly negative surface charges (e.g. ZB5) (Fig. 5E; Fig. S20). In addition, by substituting residues in positions 300, 302 and 304 in CM, we altered the surface charges of its inner LRR region. For example, changing K-D-F to D-D-S resulted in a negatively charged surface, whereas K-H-F conferred a positive charged surface (Fig. 5C, E; Fig. S23). We concluded that these three residues govern the surface charges of the 8^th^-11^th^ LRR motifs, and thus determine SCORE ligand specificity.

Next, we generated 37 CM substitution variants and assessed their recognition specificities for csp15 (Fig. 5E). Consistent with our predictions, variants sharing similar inner LRR surface charges exhibited similar csp15 recognition patterns (Fig. 5D; Fig. S23, 24). Although we did not produce a CM variant that recognizes most csp15 peptides, we were able to develop variants that perceive CSPs from multiple pathogens and pests (Fig. 5D, 6; Fig. S25). WT CM is unable to respond to CSP peptides from *Ralstonia* sp., *Erwinia* sp., *Xanthomonas* sp., *Candidatus Liberibacter asiaticus*, or root-knot nematodes. However, multiple amino acid substitutions, ranging from single-to quadruple site changes, conferred robust responsiveness to CSPs from these pathogens (Fig. 6; Fig. S26). Multiple bacterial and fungal pathogens, such as *Pseudomonas syringae, Fusarium graminearum*, and *Ustilaginoidea virens*, produce csp15 variants with negatively charged residues at positions 6 and 7 (aspartic acid followed by glutamate; “DE”). WT CM is able to weakly perceive this variant, but a single amino acid substitution from K-D-F to K-S-F greatly enhanced its sensitivity (Fig. 6; Fig. S26). By strategically engineering a few key residues, we improved SCORE’s capacity to recognize pathogen- and pest-derived CSPs.

**Fig. 6.**
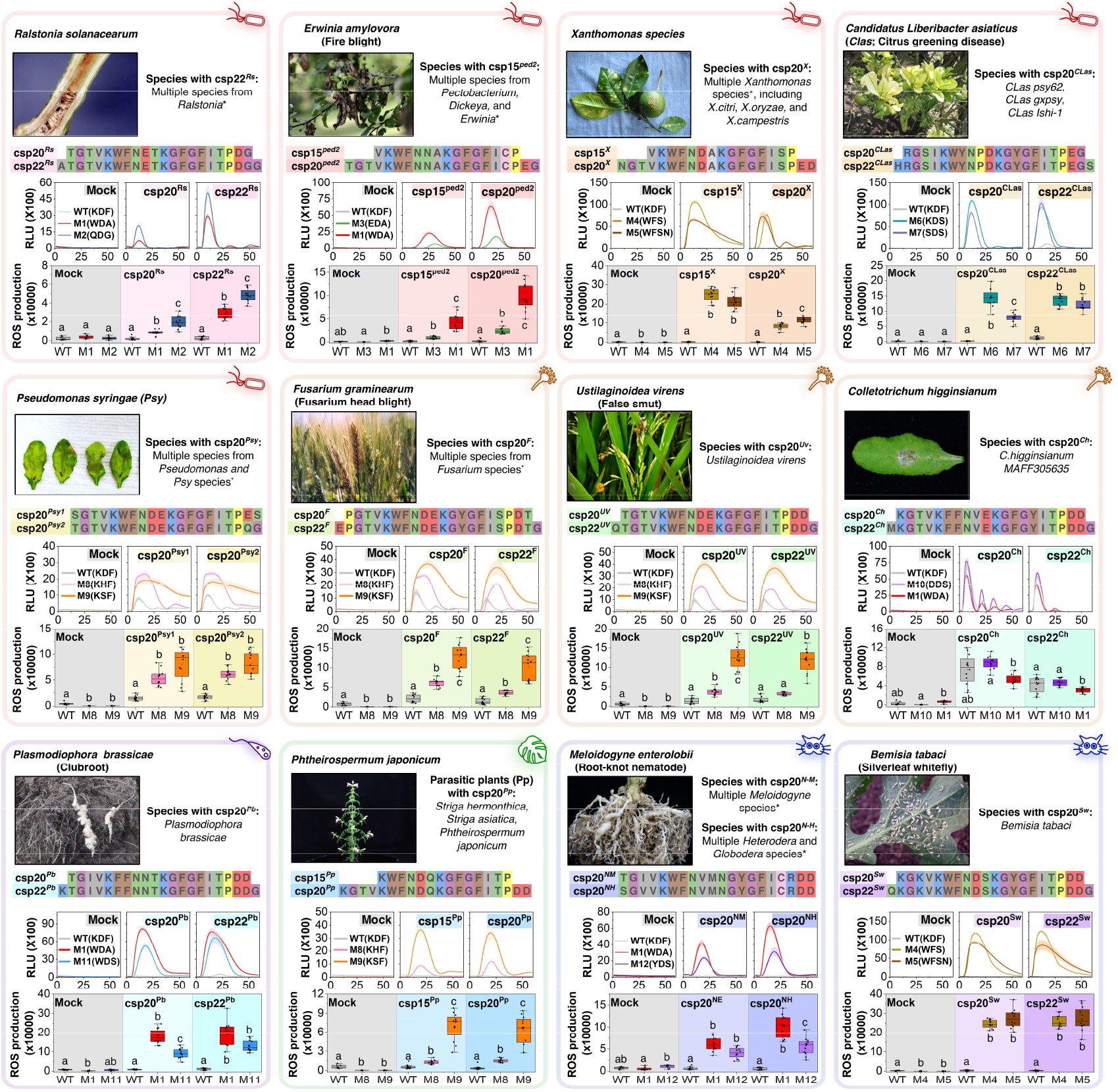
Synthetic SCOREs confer recognition of CSP peptides from multiple plant pathogens and pests. Wildtype (WT) SCORE has residues K, D, F and Y at positions 300, 302, 304 and 305, respectively. Synthetic SCORE variants (M1-M12) were engineered by substituting these residues as indicated. WT SCORE and its variants were expressed in *N. benthamiana* leaves, followed by treatment with 100nM of the indicated CSP peptides. ROS production was measured for one hour. Amino acid substitutions in M1-M12 either boosted CSP recognition by WT, or conferred CSP sensitivity to previously non-responsive variants. Box plots indicate ROS production by WT SCORE or its variants over an hour following mock or peptide treatments. For box plots, center line represents the median, box boundaries are the 25th and 75th percentiles, whiskers indicate the 1.5 × interquartile range (IQR) from the 25th and 75th percentiles. Data points from 8 technical replicates (for mock treatment) and 12 technical replicates (for peptide treatments) were analyzed with a one-sided Kruskal-Wallis test followed by Dunn’s multiple comparisons test. Data points with different letters indicate significant differences of P < 0.05. *For full list of species with the indicated CSP variants, refer to Fig. S25. Acknowledgements and Creative Commons licenses for the plant disease images are provided in supplementary text.

## Discussion

Plants rely on an extensive repertoire of cell-surface receptors to detect and respond to rapidly changing environmental conditions. Pathogen pressure drives the expansion of both intracellular and cell-surface immune receptor repertoires (*12*). But immune receptors can also be lost or pseudogenized due to fitness costs associated with retaining them within the genome (*25, 26*). We propose that this “mosaic” pattern of PRRs in land plants, where LRR-RLK-XIIs from various clusters are interspersed across genomes, reflects a mixture of evolutionarily ancient and conserved receptors (such as FLS2 and CERK1), lineage-specific receptors (such as CORE, MIK2 and EFR), and ancient but lineage-specific receptors (such as SCORE) (Fig. S3, 4) (*16, 27–29*). As a result, each species possesses a unique set of PRRs, shaped by its specific ecological pressures.

Since the discovery of the first plant cell-surface immune receptor in 1994 (*30*), identification of novel PRRs has received considerable attention, but clearly more work in this area is yet to be done. While genetic approaches have been fruitful in model organisms such as *A. thaliana* and *Solanum* species, characterizing PRRs in non-model plants remains challenging, especially in tree and polyploid species, which have much expanded immune receptor repertoires (*12*). By integrating bioinformatics, synthetic biology, and biochemical approaches, we have demonstrated that PRRs from these species can also be characterized. While this system may not capture receptors that require lineage-specific co-receptors and/or co-factors, and are therefore harder to transfer into crop species, it nonetheless paves the way for identifying PRRs from diverse plant lineages, ultimately facilitating the breeding of disease-resistant crops.

To our knowledge, SCORE and CORE are the first identified PRRs that have convergently evolved to perceive the same peptide ligand. Further work is required to resolve the structural mechanisms by which these receptors perceive CSP peptides, particularly given their distinct csp22 recognition patterns. Conceptually, PRRs perceive MAMPs with highly conserved, functionally critical motifs, making it difficult for pathogens to evade detection. Yet increasing evidence suggests that some PRRs and MAMPs are engaged in a dynamic evolutionary arms race, as exemplified by FLS2 and PERU (*31–35*). We speculate that similar processes shape SCORE, whose orthologs exhibit unprecedented polymorphism in ligand specificity. More importantly, the modifiable recognition specificity of SCOREs provides fresh opportunities for engineering disease resistance in tree crops. For example, citrus canker and citrus greening are both economically significant citrus diseases caused by *Xanthomonas* spp. and *Candidatus Liberibacter asiaticus* (*CLas*), respectively (*36, 37*). Several citrus species perceive csp22^Mic^, but not csp22^CLas^, likely through their endogenous SCORE orthologs (*38*). With modern base-editing tools, it may be feasible to generate citrus variants that carry endogenous SCORE receptors altered to perceive csp22^CLas^, thereby providing resistance against these devastating pathogens.

## Acknowledgments

We thank Y. Nagai, N. Maki, N. Watanabe, and M. Yamamoto for providing technical support, Shirasu lab, for discussions and critical reading of the manuscript. We thank M. Albert for sharing the information on chimeric receptors and BES1-HA experiments. A full list of acknowledgments for the images used in Fig. 4 and 6 is available in the supplementary text.

## Author contributions

Conceptualization: BPMN, YK, KS

Methodology: BPMN, MW, MWS, YS, ND, YK, KS

Investigation: BPMN, MW, MWS, TS, YK

Bioinformatic analyses design: BPMN, MW, MWS, YK Bioinformatic analyses: MW, MWS

Visualization: BPMN

Funding acquisition: BPMN, ND, YK, KS

Project administration: BPMN, MWS, ND, YK, KS

Supervision: ND, YK, KS

Writing – original draft: BPMN

Writing – review & editing: BPMN, MW, MWS, TS, ND, YK, KS

## Data and materials availability

All data are available in the main text or the supplementary materials.

## Supplementary Materials

Materials and Methods

Supplementary Text

Figs. S1 to S26

Reference 1-58

Data S1 to S2

