## Supplementary Materials for "Systematic Discovery and Design of Synthetic Immune Receptors in Plants"

**The PDF file includes:**

Materials and Methods
Supplementary Text
Figs. S1 to S26
References 1-58

**Other Supplementary Materials for this manuscript include the following:**

Data S1 to S2

### Materials and Methods

#### Vector construction

The CDS regions of LRR-RLK-XII ectodomains were synthesized by Twist Bioscience or Genscript, or directly amplified by PCR using KOD One polymerase (Toyobo, Japan). The PCR products were cloned into the epiGreenB5 vector (3×HA) between the ClaI and BamHI restriction sites using the In-Fusion HD Cloning Kit (Clontech, USA) to generate p35S::cell-surface receptor$^{Ecto}$-BRI1$^{TM+KD}$-HA or p35S::cell-surface receptor$^{Ecto}$-EFR$^{TM+KD}$-HA (epiGreenB5-Cauliflower mosaic virus (CaMV) p35S: gene of interest-3×HA). The resulting constructs were transformed into *Agrobacterium tumefaciens* strain *AGL1* for transient expression in *Nicotiana benthamiana*. All chimeric cell-surface receptors generated in this study include the EFR signal peptide and either the BRI1 TM+KD or EFR TM+KD, ensuring consistency in construct design and expression levels. For recombinant protein expression, Genes encoding A-type flagellin from *Pseudomonas aeruginosa* (KJJ22056.1), elongation factor Tu from *Escherichia coli* K-12 (8FR3_A), and cold-shock protein from *Cupriavidus* (WP_029045138.1) were synthesized and codon-optimized for expression in *Escherichia coli*. These genes were cloned into the pMAL-c6T expression vector (New England Biolabs) to add an N-terminal maltose-binding protein (MBP) tag for purification.

#### Transient expression in *Nicotiana benthamiana*

*Agrobacterium tumefaciens* strain *AGL1* carrying the binary expression vectors was cultured on LB agar plates supplemented with selection antibiotics. The bacterial cultures were harvested by centrifugation, and the pellets were resuspended in an infiltration buffer containing 10 mM $MgCl_2$, 10 mM MES (pH 5.6), and 100 μM acetosyringone. The concentrations of AGL1 were adjusted to $OD_{600} = 0.5$ for chimeric receptor constructs and $OD_{600} = 0.25$ for the BES1 construct. The prepared suspensions were syringe-infiltrated into *Nicotiana benthamiana* leaves.

#### BES1 assay

Three days after the infiltration of the *AGL1* suspension, *Nicotiana benthamiana* leaves were treated with either peptide, full-length recombinant protein, bacterial suspension, or boiled bacterial suspension. Leaf discs were collected using a 6-mm-diameter cork borer and snap-frozen at the indicated time points. The samples were then processed for immunoblotting as described below.

#### MAPK assay

Three days after the infiltration of *AGL1* suspension, *Nicotiana benthamiana* leaves were infiltrated with either peptide or full-length recombinant protein. Leave discs were collected with a 6-mm-diameter cork borer and snap-frozen at indicated time points, followed by immunoblotting (see below). Phosphorylation of NbSIPK and NbWIPK (pMAPK) were detected with α-phospho-p44/42 MAPK rabbit monoclonal antibody (D13.14.4E, in 1:2000, Cell Signaling Technology, USA).

**Immunoblotting**

Protein extraction was performed as previously described (4). *Nicotiana benthamiana* leaves tissues were lysed in liquid nitrogen and extracted in 1×NuPAGE™ LDS Sample Buffer (Invitrogen™) with 10mM DTT at 70°C for 10 minutes. Total proteins were then separated by SDS-PAGE and blotted onto a PVDF membrane (Trans-Blot Turbo Transfer System, Bio-Rad). The membrane was then blocked in a solution of either 5% skimmed milk (for BES1 and cell-surface receptor detection) or 5% bovine serum albumin (BSA; for MAPK detection) in Tris-buffered saline, 0.1% Tween 20 detergent (TBST) for an hour. Phosphorylated MAPKs were detected using α-phospho-p44/42 MAPK rabbit monoclonal antibody (D13.14.4E, in 1:2000, Cell Signaling Technology, USA) in a solution of 5% BSA in TBST overnight at 4°C. HA-tagged BES1 or cell surface receptors were detected using Anti-HA-Peroxidase, High Affinity, rat $IgG_1$ antibody (Roche) in a solution of 5% skimmed milk in TBST overnight at 4°C. For detection of MAPKs, this is followed by incubation with α-rabbit IgG-HRP-conjugated secondary antibodies (1:10000, Roche, USA) in a solution of 5% BSA in TBST for an hour at room temperature. HRP signal was then detected by Clarity Western ECL Substrate (Bio-Rad) with a LAS 4000 system (GE Healthcare, USA). The PVDF membranes were stained with Coomassie Brilliant Blue (CBB) to ensure equal protein loading.

**Protein extraction and immunoprecipitation**

Protein extraction for immunoprecipitation was performed as previously described (4). Three to four grams of *Nicotiana benthamiana* leaves were used for immunoprecipitations. Three days following transient expression, *Nicotiana benthamiana* leaves were treated with indicated elicitors and snap-frozen. The tissues were then ground in liquid nitrogen and extracted in extraction buffer (50 mM Tris-HCl at pH 7.5, 150 mM NaCl, 10% glycerol, 5 mM DTT, 2.5 mM NaF, 1 mM $Na_2MoO_4$•2H2O, 0.5% polyvinylpyrrolidone (w/v), 1% Protease Inhibitor Cocktail (P9599; Sigma-Aldrich), 100 µM phenylmethylsulphonyl fluoride and 2% IGEPAL CA-630 (v/v; Sigma-Aldrich), and 2 mM EDTA) at a concentration of 3mL/g tissue powder. Samples were then incubated at 4°C for an hour and debris was removed by centrifugations (at 13,000 rpm for 10 min at 4°C). The supernatant was then collected, and the protein concentrations were adjusted to 5mg/mL and incubated with rotation for an hour at 4°C with 50 µL anti-HA magnetic beads (Miltenyi Biotec) for immunoprecipitation. Magnetic beads were then washed twice with extraction buffer and the HA-tagged protein was eluted with sodium dodecyl sulfate (SDS) sample buffer at 95°C.

**Biotinylated peptide cross-linking assay**

The biotinylated peptide cross-linking assay was conducted as previously described (4). *Nicotiana benthamiana* leaves transiently expressing different receptors were vacuum-infiltrated with the specified biotinylated peptides for 5 minutes. Following peptide infiltration, the solutions were replaced with a 2 mM ethylene glycol bis(succinimidyl succinate) (EGS) cross-linker solution (prepared by dissolving 0.09 g of EGS in 1 mL of DMSO and mixing with 100 mL of 50 mM HEPES, pH 7.4). The leaves were vacuum-infiltrated again in the cross-linker solution for 5 minutes and subsequently incubated for 25 minutes. After incubation, the leaves were snap-frozen, and proteins were extracted from the samples. Receptors were immunoprecipitated as previously described. Biotinylated peptides cross-linked to the receptors were visualized in immunoblots using Pierce™ High Sensitivity Streptavidin-HRP (21130).

**ROS assay**

ROS burst assay was performed as described previously (4). *Nicotiana benthamiana* leaf discs were collected with a 4-mm-diameter cork borer and placed in 96-well plates with 120µl deionized water overnight in the dark (abaxial surface of the leaves facing down). *Nicotiana benthamiana* leaf discs were then treated with either mock (water) or peptide solutions in 20 mM luminol (Wako, Japan) and 0.02 mg $ml^{-1}$ horseradish peroxidase (Sigma-Aldrich). Luminescence was then measured over indicated periods of time with a Tristar2 multimode reader (Berthold Technologies, Germany).

**Size exclusion spin column to fractionate Agrobacterium extracts**

Agrobacterium extracts were fractionated using Sartorius Vivaspin® centrifugal concentrators, following the manufacturer's instructions.

**Anion exchange chromatography/Size-exclusion chromatography**

*Agrobacterium tumefaciens AGL1* was dissolved in Extraction Buffer (20 mM Tris, pH 8.0, 0.1% CHAPS (FUJIFILM Wako Pure Chemical Corporation) supplemented with cOmplete Ultra Protease Inhibitor Cocktail (Merck, Rahway, New Jersey), and boiled for 10 min to extract proteins. The extract was centrifuged at 12,000 g for 10 min to remove aggregated proteins and debris. The clarified supernatant was subjected to purification using an ÄKTA FPLC system (Cytiva, Marlborough, MA). The supernatant was first desalted using a HiPrep 26/10 desalting column (Cytiva) and Desalting Buffer (same as Extraction Buffer). The desalted extract was then applied to a HiTrap Q anion exchange chromatography column pre-equilibrated with Anion Exchange Binding Buffer (20 mM Tris, pH 8.0, 0.1% CHAPS). Bound proteins were eluted with increasing salt concentrations using Anion Exchange Elution Buffer (20 mM Tris, pH 8.0, 0.1% CHAPS, 1 M NaCl). Each fraction was desalted using an ultrafiltration column (10 kDa MWCO) with 10 mM MES, pH 5.6, and subsequently tested for immunogenic activity using the BES1 band shift assay. Fractions with immunogenic activity were then pooled and concentrated with ultrafiltration column (10 kDa MWCO) and subjected to Superdex 200 gel filtration chromatography in Gel Filtration Buffer (20 mM Tris, pH 8.0, 150 mM NaCl). The gel filtration fractions were again desalted using ultrafiltration columns and tested for immunogenic activity by BES1 assay. The fractions, A6-A9 were submitted for LC-MS/MS analysis to identify the protein.

**LC-MS/MS analysis**

The A6-A9 fractions were processed using the SP3 method (40). Proteins were reduced and carbamidomethylated with dithiothreitol (DTT) and iodoacetamide, respectively. The samples were digested with trypsin (tosylphenylalanyl chloromethyl ketone-treated, Worthington Biochemical Co.) overnight at 37 °C. The resulting peptides were desalted using GLtip SDB columns (GL Science, Tokyo, Japan). Peptides were analyzed by liquid chromatography-tandem mass spectrometry (LC-MS/MS) using a Q Exactive HF-X mass spectrometer (Thermo Fisher

Scientific, Bartlesville, OK, USA) coupled with an Easy-nLC 1200 system (Thermo Fisher Scientific). The samples were analyzed with a nanoelectrospray ionization (nanoESI) spray column (NTCC-360, 0.075 mm i.d., 150 mm length, 3 µm particle size; Nikkyo Technos, Inc.). Peptide separation was achieved using solution A (0.1% formic acid) and solution B (80% acetonitrile with 0.1% formic acid) at a flow rate of 300 nL/min, with the following gradient: 0–20 min, linear increase from 0% to 80% solution B. The mass spectrometer was operated in positive ion mode using a data-dependent acquisition (Top 10) method. The acquired data were processed using the MASCOT search engine version 2.8 (Matrix Science, London, UK) and Proteome Discoverer 3.0 (Thermo Fisher Scientific). Protein identification was performed with the following parameters: database, Agrobacterium fabrum str. C58 (in-house database); enzyme, trypsin; fixed modification, none; variable modifications, acetyl (protein N-term), oxidation (M), Gln->pyro-Glu (N-term Q), propionamide (C); mass values, monoisotopic; peptide mass tolerance, 15 ppm; fragment mass tolerance, ± 30 mmu; maximum missed cleavages, 3; and instrument type, ESI-TRAP. Label-free quantification was performed using Proteome Discoverer (version 3.0).

### Peptides

The peptides used in this study were synthesized by GenScript. The concentration of each peptide for individual experiments is provided in the figure captions or Supplementary Table 2.

### Recombinant proteins

The plasmids encoding full-length flagellin, EF-Tu, and cold-shock protein, cloned into the pMAL-c6T expression vector, were transformed into *Escherichia coli* strain Rosetta-gami™ 2(DE3) (Merck). Recombinant protein expression was induced by adding 0.3 mM isopropyl-β-D-thiogalactopyranoside (IPTG) to cultures at an optical density at 600 nm (OD600) of 0.6–0.8, followed by incubation at 16 °C overnight. Bacterial cells were harvested by centrifugation, resuspended in lysis buffer (20 mM Tris-HCl, 200 mM NaCl, pH 7.4), and lysed by sonication. Proteins were purified from the soluble fraction using amylose resin (New England Biolabs), following the manufacturer's protocol. Purified proteins were eluted with 10 mM maltose, and the buffer was exchanged to 20 mM MES (pH 5.6) using a 10 kDa molecular weight cutoff ultrafiltration column (Sartorius). The purity of the proteins was confirmed by SDS-PAGE analysis.

### AlphaFold structure predictions

AlphaFold2 and AlphaFold3 were used to predict the protein structure of $SCORE^{ecto}$-csp15, and $SCORE^{ecto}$-csp15-$BAK1^{ecto}$ (23,39).

### Protein surface charge predictions

Protein surface charges were predicted by ProteinSol online server as instructed (24).

### Protein structure visualization

Predicted protein structures were analysed and visualized by iCn3D (41).

**Repeat conservation mapping (RCM) with LRR-RLK-XIIs**

Initially, 19,633 LRR-RLK-XII candidates were identified from 350 plant genomes (12). These candidates were further filtered by selecting LRR-RLK-XIIs containing a kinase domain, at least 10 LRRs, and 1-2 transmembrane helices between them. The kinase (KD), LRR, and transmembrane (TM) domains were identified as described previously (12) using hmmer (version 3.1b2, (42)) predict-phytolrr (obtained in December 2021, (43)), and tmhmm (version 2.0, (44)). This filtering resulted in 13,185 LRR-RLK-XII candidates. To assess conservation within the LRR domain, a method similar to repeat conservation mapping (RCM) (17) was employed, iteratively grouping LRR-RLK-XIIs based on their LRR domain similarity.

The LRR domains were extracted and aligned using FAMSA (45). The alignments were not trimmed (46), and phylogenetic trees were inferred with FastTree (version 2.1.11 SSE3, option -lg, (47)). The trees were rooted using gotree (v0.4.2, (48)), with one sequence from the most basal species serving as the outgroup. The resulting phylogenetic tree was converted into a distance matrix. Distances above a specified threshold were then masked, and a network was created based on the pairwise distances below this threshold. Communities of similar kinase domains were identified using a modularity optimization algorithm (49). Various threshold values were scanned, and the threshold chosen was the smallest distance plus 0.16 times the largest distance in the matrix. This approach resulted in 351 groups, each containing at least 5 members, and encompassing a total of 12,483 RLKs.

For each group, the LRR domains were extracted and aligned using MUSCLE. Gaps were trimmed with TrimAl (version v1.4.rev22, (50)), and conservation scores were calculated using MstatX (1-weighted entropy, github.com/gcollet/MstatX). A consensus sequence was then generated using em_cons (51). To map the conservation scores onto the potential three-dimensional structure, the positions of the LRR repeats were identified again using Phyto-PredictLRR (43). If no repeats were detected in the consensus sequence, the longest sequence was used as the reference for the plot. The resulting conservation plots were manually screened. To evaluate the conservation in the upper left region of the RCM plot, the median sequence conservation in that area was extracted ($50^{th}$ percentile). Groups with a conservation score below 0.6 were further split if they contained more than 10 sequences from at least 3 species. The splitting procedure was applied using the distance matrix derived from LRR domain alignments, with the median distance serving as the threshold for network construction. This process was repeated up to three times, ultimately resulting in 212 curated LRR-RLK-XII groups, encompassing 4,177 RLKs from 285 species.

**Identification of cold-shock proteins across taxa**

CSP domains were screened across all available proteomes from NCBI GenBank. Proteomes were downloaded using genome_updater (v0.6.3) with the following options: -A "species:1" -d "genbank" -f "protein.faa.gz" -t 20 -m -L curl. Additionally, amino acid collections from InsectBase 2.0, WormBase, and 350 plant genomes were included in the screening (52–54). CSP domains were identified using HMMER (v3.1b2, (42)) with an E-value threshold of 1000, employing the CSP HMM profile available from PFAM (PF00313.23, (55)). All resulting sequences were aligned using FAMSA and trimmed to match the original csp15Mic sequence

(VKWFNAEKGFGFITP). The frequency of each csp15 sequence was counted, and 103 sequences were selected from the 350 most frequent csp15 sequences.

**In-depth search of SCORE homologs in plants**

To identify additional SCORE homologs, homologs were first searched in multiple plant genomes using DIAMOND (version 0.9.26.127 with options -e 1e-10 -k 100000, (56)), using full-length and ectodomain sequences from the *Citrus maxima* SCORE receptor (181). The ectodomains of all matches and the original candidates were aligned and manually filtered. Simultaneously, a manual search in online resources identified the *Drimys winteri* homolog. To further expand the search, all plant genomes used for CSP identification and available plant transcriptomes from the 1KP database (57) are included. These databases were then searched for homologs using all citrus and *Drimys winteri* candidate sequences. Ectodomains of all original and novel candidates were aligned. The following criteria were used to narrow down potential SCORE orthologs: (1) Receptors with more than 55% identity to SCORE. (2) Receptors with at least 10 LRRs, a transmembrane domain (TM), and a kinase domain (KD). (3) Receptors with binding sites for csp15 predicted by Alphafold2 (23) (34 sites; supplementary figure 19). (4) Receptors with highly similar csp15 binding sites to *C. maxima* SCORE, identified from the SCORE phylogenetic tree (supplementary figure 16).

### Supplementary Text

#### Acknowledgments and Creative Commons licenses for the plant images used in Fig 4C

Due to space limitation in the figure legends, we provide the acknowledgments and the Creative Common licenses for the plant images used in Fig 4C here.

Image QR: This image is licensed under the Creative Commons Attribution-Share Alike 3.0 Unported license (https://creativecommons.org/licenses/by-sa/3.0/deed.en). We acknowledge and thank the author Ввласенко for the use of this image. For further information, refer to https://commons.wikimedia.org/wiki/File:Red_oak_(Quercus_rubra)_blooming.jpg

Image AA: This image is licensed under the Creative Commons Attribution-Share Alike 4.0 International license. (https://creativecommons.org/licenses/by-sa/4.0/deed.en). We acknowledge and thank the source Bibliothèque de l'Université Lavalfor the use of this image. For further information, refer to https://commons.wikimedia.org/wiki/File:Acorus_americanus_15-p.bot-acorus.calam-19.jpg

Image BP: This image is licensed under the Creative Commons Attribution-Share Alike 4.0 International license. (https://creativecommons.org/licenses/by-sa/4.0/deed.en). We acknowledge and thank the author Krzysztof Ziarnek, Kenraiz for the use of this image. For further information, refer to https://commons.wikimedia.org/wiki/File:Betula_platyphylla_subsp._mandshurica_kz03.jpg

Image PB: This image is licensed under the Creative Commons Attribution-Share Alike 3.0 Unported license (https://creativecommons.org/licenses/by-sa/3.0/deed.en). We acknowledge and thank the author Johnny S. for the use of this image. For further information, refer to https://commons.wikimedia.org/wiki/File:Papaver_bracteatum-flower05.jpg

Image CK: This image is licensed under the Creative Commons Attribution-Share Alike 4.0 International license. (https://creativecommons.org/licenses/by-sa/4.0/deed.en). We acknowledge and thank the author Ping an Chang for the use of this image. For further information, refer to https://commons.wikimedia.org/wiki/File:%E7%89%9B%E6%A8%9FCinnamomum_kanehirae_20210503094855_10.jpg

Image DD: This image is licensed under the Creative Commons Attribution-Share Alike 4.0 International license. (https://creativecommons.org/licenses/by-sa/4.0/deed.en). We acknowledge and thank the author Agnieszka Kwiecień, Novafor for the use of this image. For further information, refer to https://commons.wikimedia.org/wiki/File:Dipteronia_sinensis_Dwuskrzydlak_chi%C5%84ski_2023-07-21_02.jpg

Image DL: This image is made available under the Creative Commons CC0 1.0 Universal Public Domain Dedication. (https://creativecommons.org/publicdomain/zero/1.0/deed.en). We acknowledge and thank the author Dinkum for the use of this image. For further information, refer to https://commons.wikimedia.org/wiki/File:Longan_fruits.jpg

Image SO: This image is licensed under the Creative Commons Attribution-Share Alike 3.0 Unported license (https://creativecommons.org/licenses/by-sa/3.0/deed.en). We acknowledge and thank author Hans B. for the use of this image. For further information, refer to https://commons.wikimedia.org/wiki/File:Syzygium_oleosum.jpg

Image PG: This image is licensed under the Creative Commons Attribution-Share Alike 3.0 Unported license (https://creativecommons.org/licenses/by-sa/3.0/deed.en). We acknowledge and thank author H. Zellfor the use of this image. For further information, refer to https://commons.wikimedia.org/wiki/File:Punica_granatum_004.JPG

Image MC: This image is licensed under the Creative Commons Attribution-Share Alike 4.0 International license. (https://creativecommons.org/licenses/by-sa/4.0/deed.en). We acknowledge and thank the author M108t for the use of this image. For further information, refer to https://commons.wikimedia.org/wiki/File:Seven_petal_flower_of_Melastoma_candidum_in_Ishigaki_Island.jpg

Image CC: This image is licensed under the Creative Commons Attribution-Share Alike 4.0 International license. (https://creativecommons.org/licenses/by-sa/4.0/deed.en). We acknowledge and thank the author Geekstreet for the use of this image. For further information, refer to https://commons.wikimedia.org/wiki/File:Corymbia_citriodora_-_shedding_bark_5.jpg

Image EG: This image is licensed under the Creative Commons Attribution 2.0 Generic license. (https://creativecommons.org/licenses/by/2.0/deed.en). We acknowledge and thank the authors Forest and Kim Starr for the use of this image. For further information, refer to https://commons.wikimedia.org/wiki/File:Starr-201226-8886-Eucalyptus_grandis-seed_capsules-Hawea_Pl_Olinda-Maui_(50859579713).jpg

Image ZB: This image is licensed under the Creative Commons Attribution-Share Alike 4.0 International license. (https://creativecommons.org/licenses/by-sa/4.0/deed.en). We acknowledge and thank the author Didier Descouensfor for the use of this image. For further information, refer to https://commons.wikimedia.org/wiki/File:Zanthoxylum_piperitum.jpg

Image PV: This image is licensed under the Creative Commons Attribution-Share Alike 4.0 International license. (https://creativecommons.org/licenses/by-sa/4.0/deed.en). We acknowledge and thank the author Krzysztof Ziarnek, Kenraiz for the use of this image. For further information, refer to https://commons.wikimedia.org/wiki/File:Pistacia_vera.jpg

Image CL: This image is made available under the Creative Commons CC0 1.0 Universal Public Domain Dedication. (https://creativecommons.org/publicdomain/zero/1.0/deed.en). We acknowledge and thank the author Anna Frodesiak for the use of this image. For further information, refer to https://commons.wikimedia.org/wiki/File:Clausena_lansium_in_Hainan_-_02.JPG

Image CM: This image is provided by Ryo Sakai. We acknowledge and thank Ryo for the use of this image.

Image CF: This image is provided by Yuki Tanaka. We acknowledge and thank Yuki for the use of this image.

Image CR: This image is licensed under the Creative Commons Attribution-Share Alike 3.0 Unported license (https://creativecommons.org/licenses/by-sa/3.0/deed.en). We acknowledge and thank author 4028mdk09 the use of this image. For further information, refer to https://commons.wikimedia.org/wiki/File:Citrus_reticulata_April_2013_Nordbaden.JPG

The images metioned aboved were resized and cropped. Otherwise, no other modifications were made.

**Acknowledgments and Creative Commons licenses for the plant disease images used in Fig 6**

Due to space limitation in the figure legends, we provide the acknowledgments and the Creative Common licenses for the plant disease images used in Fig 6 here.

Image *Ralstonia solanacearum*: This image is licensed under the Creative Commons Attribution-Share Alike 3.0 Unported license (https://creativecommons.org/licenses/by-sa/3.0/deed.en). We acknowledge and thank author Clemson University for the use of this image. For further information, refer to https://commons.wikimedia.org/wiki/File:Ralstonia_solanacearum_symptoms.jpg

Image *Erwinia amylovora*: This image is licensed under the Creative Commons Attribution-Share Alike 3.0 Unported license (https://creativecommons.org/licenses/by-sa/3.0/deed.en). We acknowledge and thank author Sebastian Stabinger for the use of this image. For further information, refer to https://commons.wikimedia.org/wiki/File:Apple_tree_with_fire_blight.jpg

Image *Xanthomonas species*: This image is made available under the Creative Commons CC0 1.0 Universal Public Domain Dedication (https://creativecommons.org/publicdomain/zero/1.0/deed.en). We acknowledge and thank author Scot Nelson for the use of this image. For further information, refer to https://commons.wikimedia.org/wiki/File:Xanthomonas_axonopodis_P8210062_(8232130407).jpg

Image *Candidatus Liberibacter asiaticus*: This image is licensed under the Creative Commons Attribution-Share Alike 3.0 Unported license (https://creativecommons.org/licenses/by-sa/3.0/deed.en). We acknowledge and thank author Florida Division of Plant Industry Archive, Florida Department of Agriculture and Consumer Services for the use of this image. For further information, refer to https://commons.wikimedia.org/wiki/File:Citrus_greening_UGA5201066.jpg

Image *Pseudomonas syringae*: This image is taken by the author B.P.M.N.

Image *Fusarium graminearum*: This image is licensed under the Creative Commons Attribution-Share Alike 3.0 Unported license (https://creativecommons.org/licenses/by-sa/3.0/deed.en). We acknowledge and thank author division, CSIRO for the use of this image. For further information, refer to https://commons.wikimedia.org/wiki/File:CSIRO_ScienceImage_11243_Fusarium_head_blight_of_barley.jpg

Image *Ustilaginoidea virens*: This image is is licensed under the Creative Commons Attribution-Share Alike 3.0 United States (https://creativecommons.org/licenses/by-sa/3.0/us/deed.en). We acknowledge and thank author Earlycj5 for the use of this image. For further information, refer to https://commons.wikimedia.org/wiki/File:U.Virens.jpg

Image *Colletotrichum higginsianum*: This image is provided by Katsuma Yonehara. We acknowledge and thank Katsuma for the use of this image.

Image *Plasmodiophora brassicae*: This image is licensed under the Creative Commons Attribution-Share Alike 3.0 Unported license (https://creativecommons.org/licenses/by-sa/3.0/deed.en). We acknowledge and thank author Rasbak for the use of this image. For further information, refer to https://commons.wikimedia.org/wiki/File:Knolvoet_bij_bloemkool_(Plasmodiophora_brassicae_on_cauliflower).jpg

Image *Phtheirospermum japonicum*: This image is provided by Yuki Tanaka. We acknowledge and thank Yuki for the use of this image.

Image *Meloidogyne enterolobii*: This image is made available under the Creative Commons CC0 1.0 Universal Public Domain Dedication (https://creativecommons.org/publicdomain/zero/1.0/deed.en). We acknowledge and thank author Plant pests and diseases for the use of this image. For further information, refer to https://commons.wikimedia.org/wiki/File:Tomato_(Solanum_lycopersicum)-_Root-knot_nematodes_-_38850656495.jpg

Image *Bemisia tabaci*: This image is licensed under the Creative Commons Attribution-Share Alike 3.0 Unported license (https://creativecommons.org/licenses/by-sa/3.0/deed.en). We acknowledge and thank author CSIRO for the use of this image. For further information, refer to https://commons.wikimedia.org/wiki/File:CSIRO_ScienceImage_7704_Silverleaf_whitefly_Bemisia_tabaci_biotype_B.jpg

The images metioned aboved were resized and cropped. Otherwise, no other modifications were made.

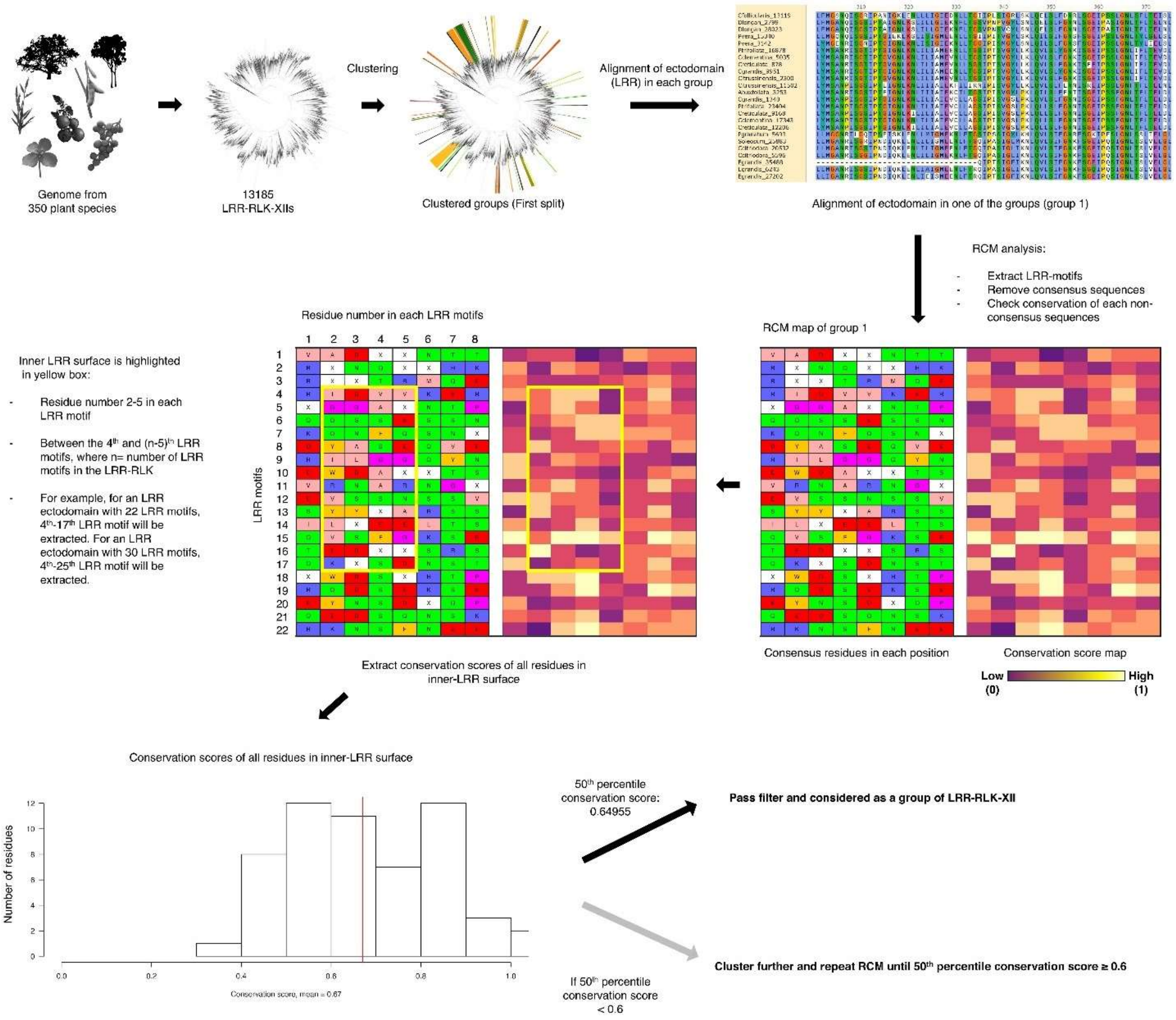


**Fig. S1. Pipeline of LRR-RLK-XII clustering with repeat conservation mapping (RCM).**

A total of 13,185 LRR-RLK-XIIs were previously identified across 350 plant genomes (54). The LRR ectodomains of these receptors were aligned, and the receptors were grouped based on phylogenetic distance. For each group, the LRR alignment was extracted. Consensus sequences were removed, and non-consensus sequences were scored based on their conservation within the group. Subsequently, the inner-LRR surface motifs (indicated by yellow box) were extracted, and their conservation scores were plotted. Receptor groups with a 50th percentile conservation score higher than 0.6 passed the filter and were considered valid. If the conservation score was below 0.6, the receptors were further clustered, and the process was repeated until groups with conservation scores exceeding 0.6 were obtained.

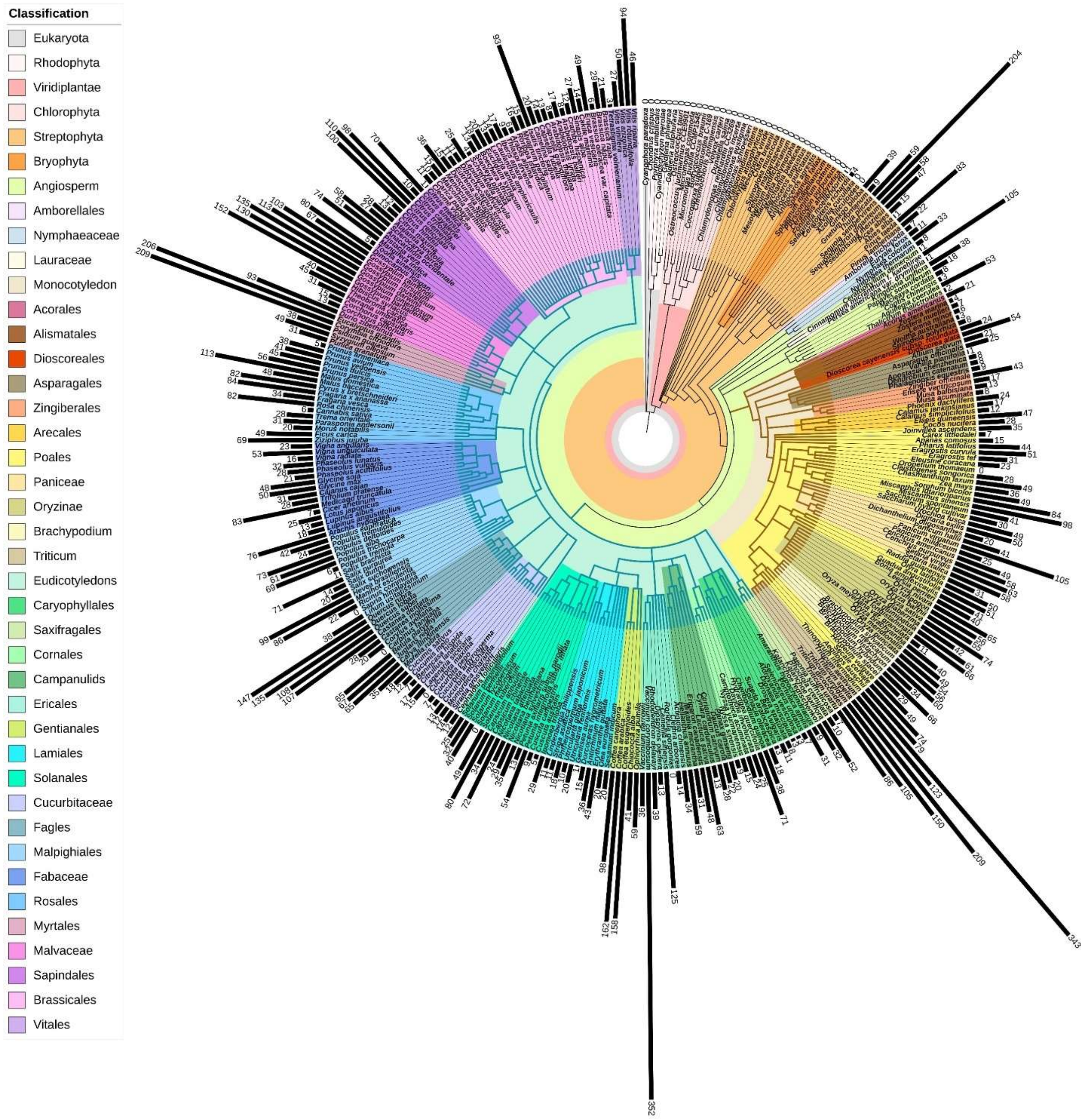


**Fig. S2. The 13,185 LRR-RLK-XIIs across 350 plant genomes.**

Phylogenetic tree of 350 plant species (300 angiosperms, 79 monocots and 208 eudicots). Bar charts represent the number LRR-RLK-XIIs in the corresponding species. Brown branches represent monocots and teal branches represent eudicots.

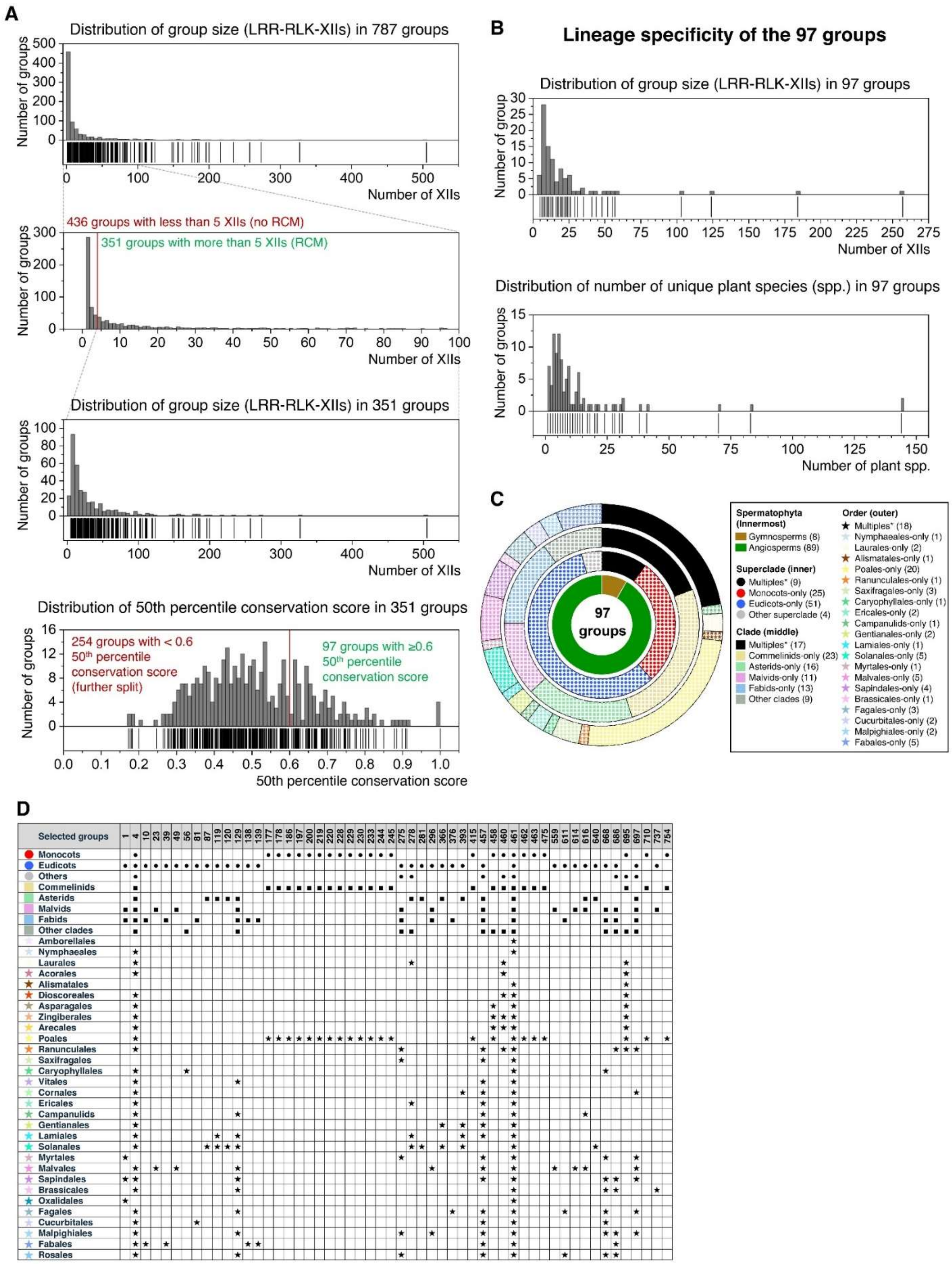
A
Distribution of group size (LRR-RLK-XIIs) in 787 groups
Number of groups
Number of XIIs
436 groups with less than 5 XIIs (no RCM)
351 groups with more than 5 XIIs (RCM)
Distribution of group size (LRR-RLK-XIIs) in 351 groups
Distribution of 50th percentile conservation score in 351 groups
254 groups with < 0.6 50th percentile conservation score (further split)
97 groups with ≥0.6 50th percentile conservation score
50th percentile conservation score
B
Lineage specificity of the 97 groups
Distribution of group size (LRR-RLK-XIIs) in 97 groups
Number of group
Distribution of number of unique plant species (spp.) in 97 groups
Number of plant spp.
C
97 groups
Spermatophyta (Innermost)
Gymnosperms (8)
Angiosperms (89)
Superclade (inner)
Multiples* (9)
Monocots-only (25)
Eudicots-only (51)
Other superclade (4)
Clade (middle)
Multiples* (17)
Commelinids-only (23)
Asterids-only (16)
Malvids-only (11)
Fabids-only (13)
Other clades (9)
Order (outer)
Multiples* (18)
Nymphaeales-only (1)
Laurales-only (2)
Alismatales-only (1)
Poales-only (20)
Ranunculales-only (1)
Saxifragales-only (3)
Caryophyllales-only (1)
Ericales-only (2)
Campanulids-only (1)
Gentianales-only (2)
Lamiales-only (1)
Solanales-only (5)
Myrtales-only (1)
Malvales-only (5)
Sapindales-only (4)
Brassicales-only (1)
Fagales-only (3)
Cucurbitales-only (2)
Malpighiales-only (2)
Fabales-only (5)
D
Selected groups

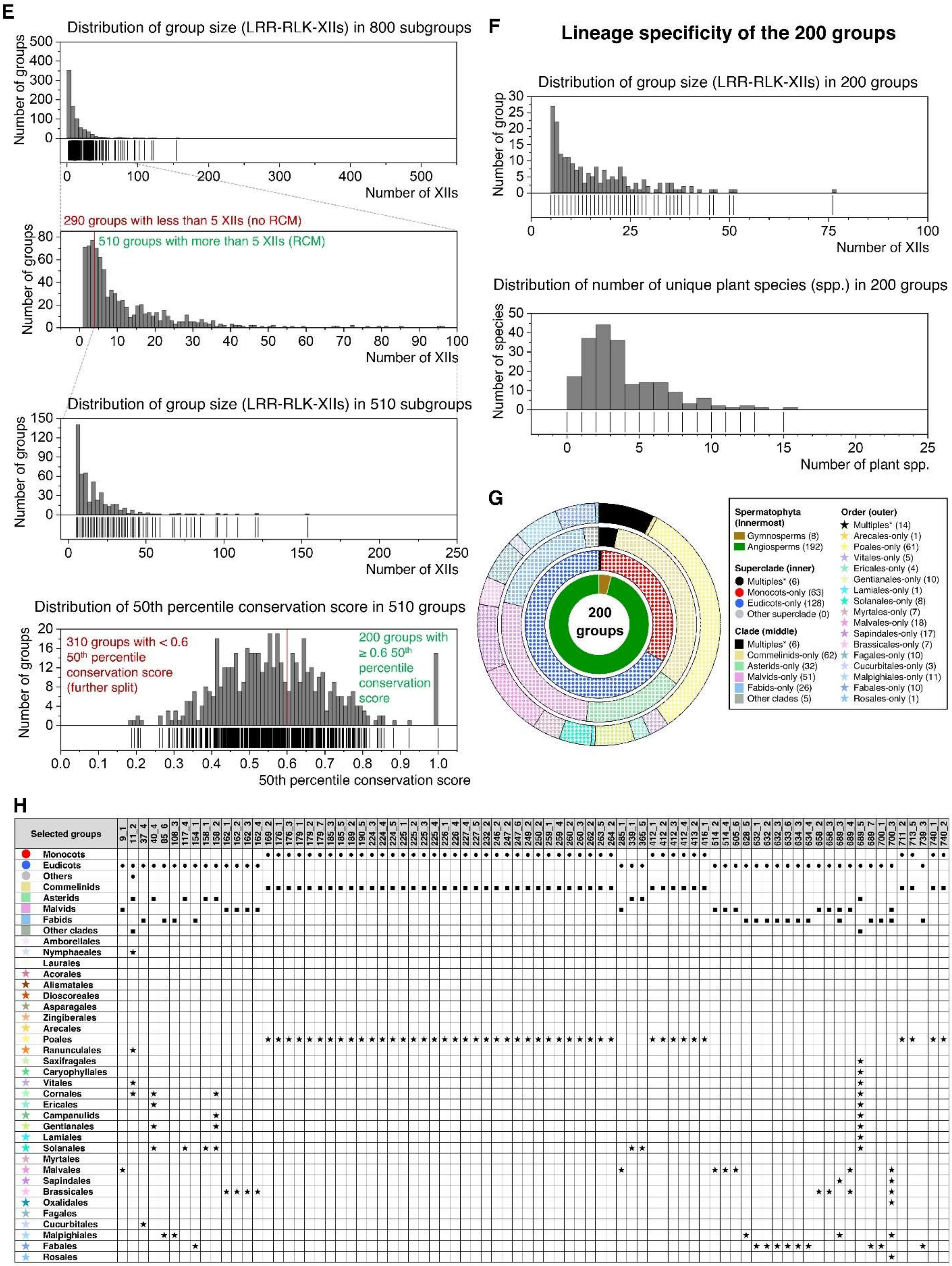
E
Distribution of group size (LRR-RLK-XIIs) in 800 subgroups
Number of groups
Number of XIIs
290 groups with less than 5 XIIs (no RCM)
510 groups with more than 5 XIIs (RCM)
Number of groups
Number of XIIs
Distribution of group size (LRR-RLK-XIIs) in 510 subgroups
Number of groups
Number of XIIs
Distribution of 50th percentile conservation score in 510 groups
310 groups with < 0.6 50th percentile conservation score (further split)
200 groups with ≥ 0.6 50th percentile conservation score
Number of groups
50th percentile conservation score
F
Lineage specificity of the 200 groups
Distribution of group size (LRR-RLK-XIIs) in 200 groups
Number of group
Number of XIIs
Distribution of number of unique plant species (spp.) in 200 groups
Number of species
Number of plant spp.
G
200 groups
Spermatophyta (Innermost)
Gymnosperms (8)
Angiosperms (192)
Superclade (inner)
Multiples* (6)
Monocots-only (63)
Eudicots-only (128)
Other superclade (0)
Clade (middle)
Multiples* (6)
Commelinids-only (62)
Asterids-only (32)
Malvids-only (51)
Fabids-only (26)
Other clades (5)
Order (outer)
Multiples* (14)
Arecales-only (1)
Poales-only (61)
Vitales-only (5)
Ericales-only (4)
Gentianales-only (10)
Lamiales-only (1)
Solanales-only (8)
Myrtales-only (7)
Malvales-only (18)
Sapindales-only (17)
Brassicales-only (7)
Fagales-only (10)
Cucurbitales-only (3)
Malpighiales-only (11)
Fabales-only (10)
Rosales-only (1)
H
Selected groups
Monocots
Eudicots
Others
Commelinids
Asterids
Malvids
Fabids
Other clades
Amborellales
Nymphaeales
Laurales
Acorales
Alismatales
Dioscoreales
Asparagales
Zingiberales
Arecales
Poales
Ranunculales
Saxifragales
Caryophyllales
Vitales
Cornales
Ericales
Campanulids
Gentianales
Lamiales
Solanales
Myrtales
Malvales
Sapindales
Brassicales
Oxalidales
Fagales
Cucurbitales
Malpighiales
Fabales
Rosales

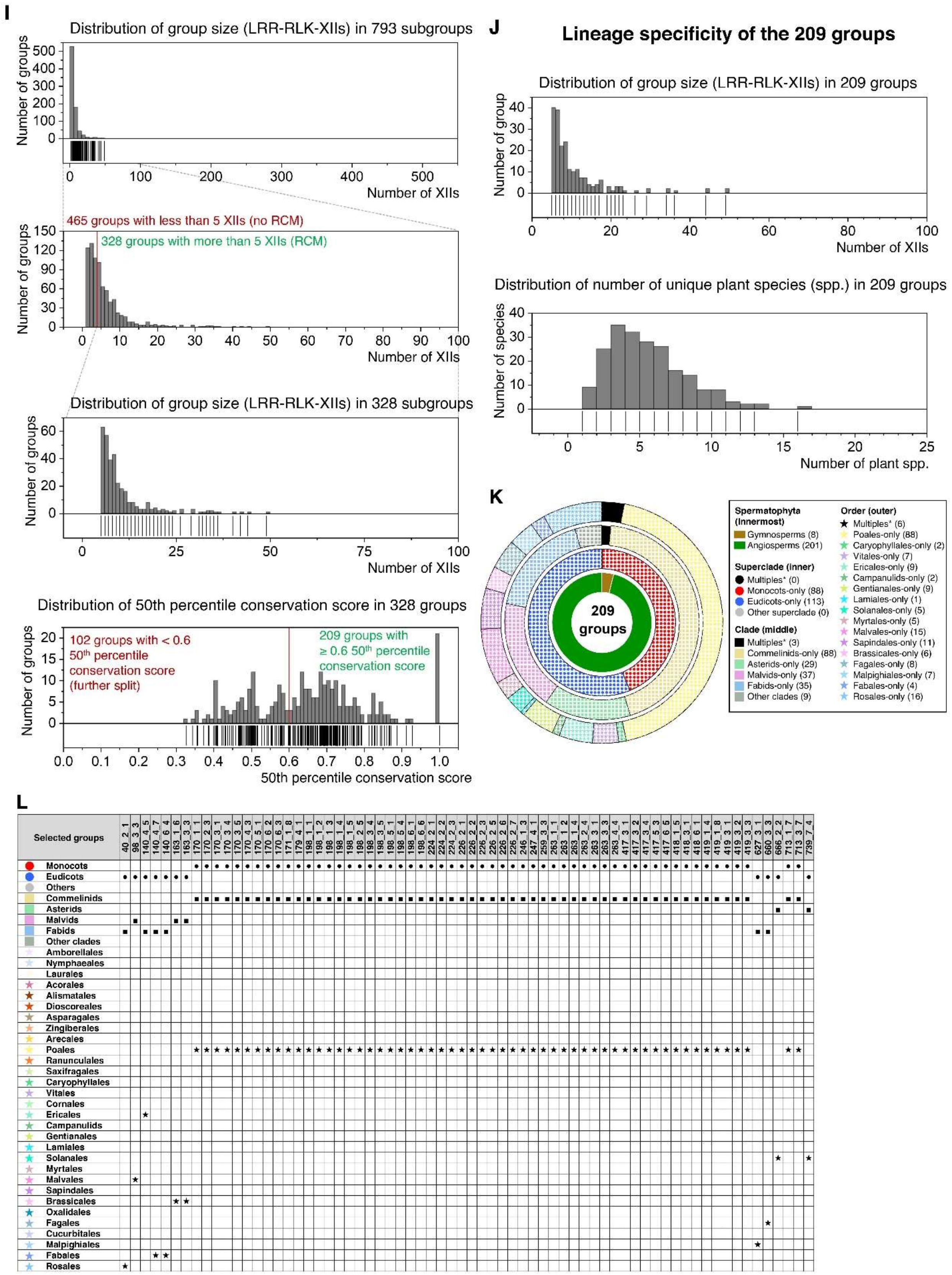
I
Distribution of group size (LRR-RLK-XIIs) in 793 subgroups
Number of groups
Number of XIIs
465 groups with less than 5 XIIs (no RCM)
328 groups with more than 5 XIIs (RCM)
Distribution of group size (LRR-RLK-XIIs) in 328 subgroups
Distribution of 50th percentile conservation score in 328 groups
102 groups with < 0.6 50th percentile conservation score (further split)
209 groups with ≥ 0.6 50th percentile conservation score
50th percentile conservation score
J
Lineage specificity of the 209 groups
Distribution of group size (LRR-RLK-XIIs) in 209 groups
Number of group
Distribution of number of unique plant species (spp.) in 209 groups
Number of species
Number of plant spp.
K
209 groups
Spermatophyta (innermost)
Gymnosperms (8)
Angiosperms (201)
Superclade (inner)
Multiples* (0)
Monocots-only (88)
Eudicots-only (113)
Other superclade (0)
Clade (middle)
Multiples* (3)
Commelinids-only (88)
Asterids-only (29)
Malvids-only (37)
Fabids-only (35)
Other clades (9)
Order (outer)
Multiples* (6)
Poales-only (88)
Caryophyllales-only (2)
Vitales-only (7)
Ericales-only (9)
Campanulids-only (2)
Gentianales-only (9)
Lamiales-only (1)
Solanales-only (5)
Myrtales-only (5)
Malvales-only (15)
Sapindales-only (11)
Brassicales-only (6)
Fagales-only (8)
Malpighiales-only (7)
Fabales-only (4)
Rosales-only (16)
L
Selected groups
Monocots
Eudicots
Others
Commelinids
Asterids
Malvids
Fabids
Other clades
Amborellales
Nymphaeales
Laurales
Acorales
Alismatales
Dioscoreales
Asparagales
Zingiberales
Arecales
Poales
Ranunculales
Saxifragales
Caryophyllales
Vitales
Cornales
Ericales
Campanulids
Gentianales
Lamiales
Solanales
Myrtales
Malvales
Sapindales
Brassicales
Oxalidales
Fagales
Cucurbitales
Malpighiales
Fabales
Rosales

M

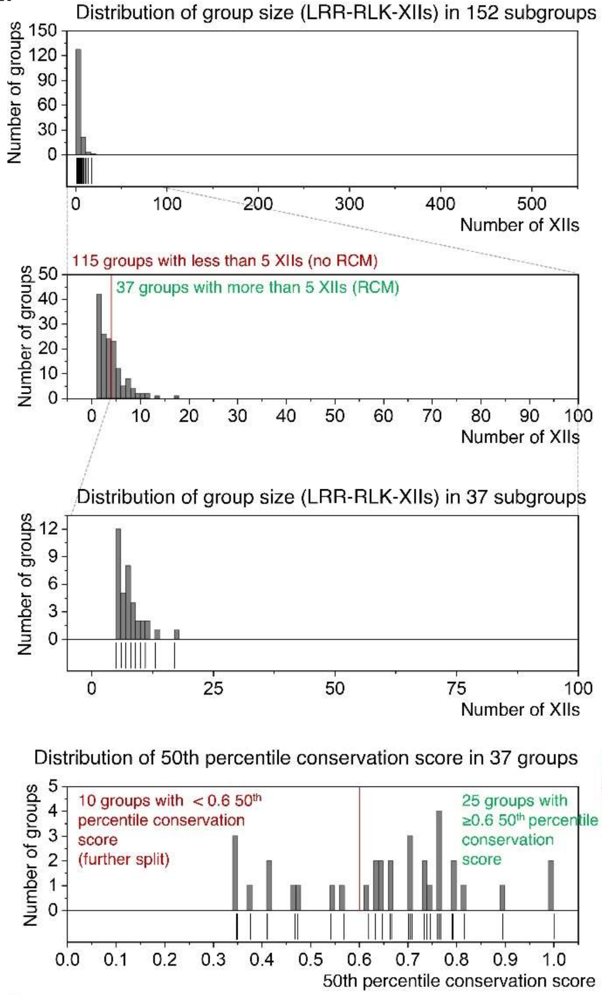


N

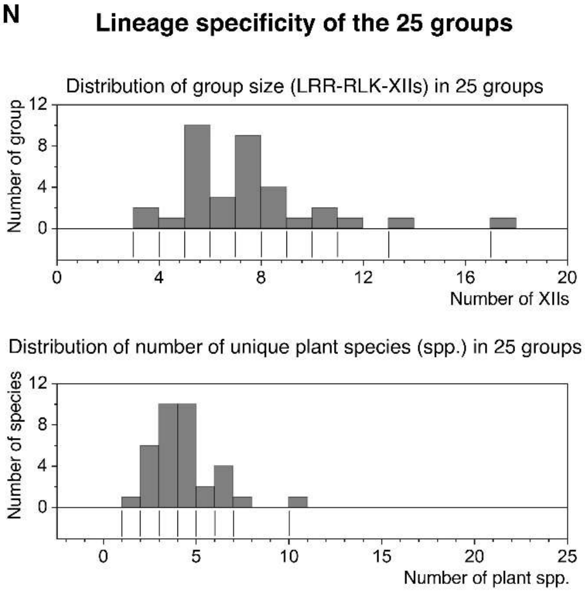


O

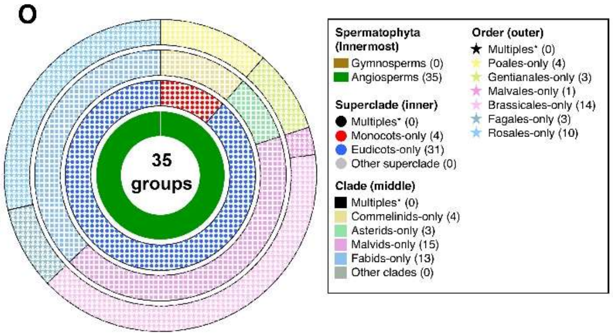


P

| Selected groups | 40_2_3_5 | 98_3_4_2 | 163_2_3_5 | 163_2_3_6 | 163_2_4_2 | 163_3_5_6 | 163_4_4_3 | 198_3_6_2 |
|---|---|---|---|---|---|---|---|---|
| Monocots | | | | | | | | • |
| Eudicots | • | • | • | • | • | • | • | |
| Others | | | | | | | | |
| Commelinids | | | | | | | | ■ |
| Asterids | | | | | | | | |
| Malvids | | ■ | ■ | ■ | ■ | ■ | ■ | |
| Fabids | ■ | | | | | | | |
| Other clades | | | | | | | | |
| Amborellales | | | | | | | | |
| Nymphaeales | | | | | | | | |
| Laurales | | | | | | | | |
| Acorales | | | | | | | | |
| Alismatales | | | | | | | | |
| Dioscoreales | | | | | | | | |
| Asparagales | | | | | | | | |
| Zingiberales | | | | | | | | |
| Arecales | | | | | | | | |
| Poales | | | | | | | | ★ |
| Ranunculales | | | | | | | | |
| Saxifragales | | | | | | | | |
| Caryophyllales | | | | | | | | |
| Vitales | | | | | | | | |
| Cornales | | | | | | | | |
| Ericales | | | | | | | | |
| Campanulids | | | | | | | | |
| Gentianales | | | | | | | | |
| Lamiales | | | | | | | | |
| Solanales | | | | | | | | |
| Myrtales | | | | | | | | |
| Malvales | | ★ | | | | | | |
| Sapindales | | | | | | | | |
| Brassicales | | | ★ | ★ | ★ | ★ | ★ | |
| Oxalidales | | | | | | | | |
| Fagales | | | | | | | | |
| Cucurbitales | | | | | | | | |
| Malpighiales | | | | | | | | |
| Fabales | | | | | | | | |
| Rosales | ★ | | | | | | | |

**Fig. S3. Information on LRR-RLK-XII groups from (A-D) first clustering, (E-H) second clustering, (I-L) third clustering, and (M-P) last clustering. (A, E, I, M)** The distribution of group sizes (number of LRR-RLK-XIIs) from the first to the last clustering is presented as follows: Top Two Plots: These illustrate the distribution of group sizes across the groups obtained at each clustering step. Only groups containing at least five LRR-RLK-XIIs were included in the RCM analysis, as highlighted in green text. Third Plot: This represents the distribution of groups with at least five LRR-RLK-XIIs. Bottom Plots: These show the distribution of the 50$^{th}$ percentile conservation scores for each group containing at least five LRR-RLK-XIIs. Groups with conservation scores of at least 0.6 are highlighted in the green text. **(B, F, J, N)** The top plot represents the distribution of group sizes across the groups that passed the conservation score filter. The bottom plot represents the distribution of species size (number of unique plant species) across these filtered groups. **(C, G, K, O)** The lineage specificity of the LRR-RLK-XII groups that passed the conservation score filter is represented. Receptors are classified as superclade-, clade-, or order-specific, indicated by colored sections (inner, middle, or outer rings, as shown on the right). Groups distributed across multiple taxa are shown in black. **(D, H, L, P)** Distribution of selected LRR-RLK-XII groups across the angiosperm lineage. 54, 80, 66 and 8 groups were manually selected from the first, second, third and last clustering, respectively. Within each subgroup, the presence of receptors within superclades, clades and orders are denoted by circles (●), squares (■), and stars (★), respectively.

| Group number | 1st split | 2nd split | 3rd split | 4th split | Colour code (hex) | Representative gene | Monocots | Eudicots |
|---|---|---|---|---|---|---|---|---|
| 1 | 1 | | | | [illegible] | Cgrandis_9951 | | * |
| 4 | 4 | | | | #8EB8F2 | Osativa_22937 | * | * |
| 9_1 | 9 | 1 | | | [illegible] | Cdarwinii_66775 | | * |
| 10 | 10 | | | | [illegible] | Lpaonicas_16858 | | * |
| 11_2 | 11 | 2 | | | #40C4FF | Vrotundifolia_3166 | | * |
| 28 | 28 | | | | [illegible] | Cdarwinii_70682 | | * |
| 37_4 | 37 | 4 | | | #418AFF | Clanatus_10689 | | * |
| 39 | 39 | | | | [illegible] | Gmax_1466 | | * |
| 39 | 39 | | | | [illegible] | Gmax_26252 | | * |
| 40_2_1 | 40 | 2 | 1 | | [illegible] | Pdulcis_22333 | | * |
| 40_2_5_3 | 40 | 2 | 5 | 3 | [illegible] | Mcoriacea_22335 | | * |
| 40_4 | 40 | 4 | | | #82C3FF | Mabacum_35913 | | * |
| 40 | 40 | | | | #F1F6FF | Cdarwinii_78142 | | * |
| 56 | 56 | | | | [illegible] | Sclerassu_3743 | | * |
| 81 | 81 | | | | #81D4FA | Carpycaparma_7775 | | * |
| 85_6 | 85 | 6 | | | #4FC3F7 | Pinthocarpa_23919 | | * |
| 87 | 87 | | | | #B3E5FC | Mabacum_4534 | | * |
| 96_3_3 | 96 | 3 | 3 | | [illegible] | Glumerosum_77493 | | * |
| 96_3_4_2 | 96 | 3 | 4 | 2 | #80D8FF | Cdarwinii_21507 | | * |
| 106_3 | 106 | 3 | | | [illegible] | Peuphratica_3307 | | * |
| 117_4 | 117 | 4 | | | [illegible] | Slycopersicum_3025 | | * |
| 119 | 119 | | | | [illegible] | Mabacum_17989 | | * |
| 120 | 120 | | | | [illegible] | Mabacum_16890 | | * |
| 129 | 129 | | | | [illegible] | Mabacum_58587 | | * |
| 138 | 138 | | | | #E3F2FD | Lpaonicas_27236 | | * |
| 139 | 139 | | | | #BBDEFB | Gmax_46068 | | * |
| 140_4_5 | 140 | 4 | 5 | | #90CAF9 | Lpaonicas_20009 | | * |
| 140_4_7 | 140 | 4 | 7 | | #64B5F6 | Vvinifera_3445 | | * |
| 140_6_4 | 140 | 6 | 4 | | #42A5F5 | Hannuus_36666 | | * |
| 154_1 | 154 | 1 | | | [illegible] | Mtruncatula_9747 | | * |
| 154_1 | 154 | 1 | | | [illegible] | Gmax_49831 | | * |
| 158_1 | 158 | 1 | | | #00BCD4 | Mabacum_41523 | | * |
| 158_1 | 158 | 1 | | | #00BCD4 | Mabacum_61034 | | * |
| 158_2 | 158 | 2 | | | [illegible] | Mabacum_44408 | | * |
| 162_1 | 162 | 1 | | | #C5CAE9 | Mcrefidatum_19466 | | * |
| 162_2 | 162 | 2 | | | #9FA8DA | Athaliana_22401 | | * |
| 162_3 | 162 | 3 | | | #7986CB | Athaliana_14308 | | * |
| 162_4 | 162 | 4 | | | #5C6BC0 | Camplexicaulis_29081 | | * |
| 163_1_6 | 163 | 1 | 6 | | [illegible] | Mcrefidatum_13666 | | * |
| 163_2_3_5 | 163 | 2 | 3 | 5 | #C5CAE9 | Itinctoria_8794 | | * |
| 163_2_3_6 | 163 | 2 | 3 | 6 | #536DFE | Alyrata_2198 | | * |
| 163_2_4_2 | 163 | 2 | 4 | 2 | #8C9EFF | Athaliana_14106 | | * |
| 163_3_3 | 163 | 3 | 3 | | #303F9F | Spinnata_22094 | | * |
| 163_3_5_6 | 163 | 3 | 5 | 6 | #283593 | Dsocnoides_15782 | | * |
| 163_4_4_3 | 163 | 4 | 4 | 3 | #3949AB | Brapa_7835 | | * |
| 169_2 | 169 | 2 | | | #F5A623 | Taestivum_53848 | * | |
| 170_1_1 | 170 | 1 | 1 | | #F57F17 | Osativa_37432 | * | |
| 170_2_3 | 170 | 2 | 3 | | #FF5722 | Tspelta_38892 | * | |
| 170_3_1 | 170 | 3 | 1 | | #C0CA33 | Bhybridum_20347 | * | |
| 170_3_4 | 170 | 3 | 4 | | #E6F4C2 | Tspelta_22808 | * | |
| 170_3_5 | 170 | 3 | 5 | | #AFB42B | Taestivum_6598 | * | |
| 170_4_3 | 170 | 4 | 3 | | [illegible] | Zmays_14704 | * | |
| 170_5_1 | 170 | 5 | 1 | | #D4E157 | Sbicolor_33104 | * | |
| 170_6_2 | 170 | 6 | 2 | | #DCE775 | Osativa_2592 | * | |
| 170_6_3 | 170 | 6 | 3 | | #E6EE9C | Osativa_15365 | * | |
| 171_1_6 | 171 | 1 | 6 | | #F0F4C3 | Zmays_25660 | * | |
| 176_1 | 176 | 1 | | | #FFEE58 | Osativa_35401 | * | |
| 176_3 | 176 | 3 | | | #FFF176 | Bdistachyon_7038 | * | |
| 177 | 177 | | | | #FFF59D | Osativa_34060 | * | |
| 178 | 178 | | | | #FDD835 | Osativa_9884 | * | |
| 179_1 | 179 | 1 | | | #FFB74D | Taestivum_29115 | * | |
| 179_2 | 179 | 2 | | | #FFCC80 | Taestivum_48483 | * | |
| 179_4_1 | 179 | 4 | 1 | | #FFC030 | Osativa_15105 | * | |
| 179_7 | 179 | 7 | | | #FFE0B2 | Tspelta_60485 | * | |
| 185_3 | 185 | 3 | | | #76FF03 | Osativa_15024 | * | |
| 185_5 | 185 | 5 | | | #B2FF59 | Taestivum_24154 | * | |
| 186 | 186 | | | | #CCFF90 | Tspelta_30877 | * | |
| 189_2 | 189 | 2 | | | #9E9D24 | Osativa_40480 | * | |
| 190_5 | 190 | 5 | | | [illegible] | Bhybridum_44057 | * | |
| 197 | 197 | | | | #64DD17 | Sbicolor_25426 | * | |
| 198_1_1 | 198 | 1 | 1 | | #DCEDC8 | Osativa_33575 | * | |
| 198_1_2 | 198 | 1 | 2 | | #C5E1A5 | Tspelta_23506 | * | |
| 198_1_3 | 198 | 1 | 3 | | #9CCC65 | Bhybridum_9275 | * | |
| 198_1_4 | 198 | 1 | 4 | | #7CB342 | Tspelta_85146 | * | |
| 198_1_5 | 198 | 1 | 5 | | #AED581 | Tspelta_111088 | * | |
| 198_2_5 | 198 | 2 | 5 | | [illegible] | Sbicolor_8156 | * | |
| 198_3_4 | 198 | 3 | 4 | | #689F38 | Taestivum_69172 | * | |
| 198_3_5 | 198 | 3 | 5 | | [illegible] | Bdistachyon_27657 | * | |
| 198_3_6_2 | 198 | 3 | 6 | 2 | #F1F8E9 | Osativa_11284 | * | |
| 198_5_1 | 198 | 5 | 1 | | #b8de9b | Osativa_7303 | * | |
| 198_5_4 | 198 | 5 | 4 | | #1b460c | Osativa_27885 | * | |
| 198_6_1 | 198 | 6 | 1 | | #00C853 | Taestivum_15567 | * | |
| 198_6_6 | 198 | 6 | 6 | | #00E676 | Osativa_21312 | * | |
| 200 | 200 | | | | #69F0AE | Sbicolor_21300 | * | |
| 219 | 219 | | | | #D7CCC8 | Zmays_5246 | * | |
| 220 | 220 | | | | #EFEBE9 | Taestivum_58574 | * | |
| 224_2_1 | 224 | 2 | 1 | | #FF8A80 | Taestivum_44169 | * | |
| 224_2_2 | 224 | 2 | 2 | | [illegible] | Taestivum_28284 | * | |
| 224_2_3 | 224 | 2 | 3 | | [illegible] | Omeyera_29235 | * | |
| 224_3 | 224 | 3 | | | [illegible] | Omeyera_28882 | * | |
| 224_4 | 224 | 4 | | | #EF5350 | Osativa_31553 | * | |
| 224_5 | 224 | 5 | | | [illegible] | Osativa_39379 | * | |
| 225_1 | 225 | 1 | | | #FFEBEE | Taestivum_21448 | * | |
| 225_2 | 225 | 2 | | | #FFCDD2 | Osativa_39019 | * | |
| 225_3 | 225 | 3 | | | #E57373 | Taestivum_41671 | * | |
| 225_4 | 225 | 4 | | | #EF9A9A | Obrachyantha_29273 | * | |
| 226_1 | 226 | 1 | | | #AD1457 | Taestivum_44435 | * | |
| 226_2_1 | 226 | 2 | 1 | | #FF4081 | Bdistachyon_13888 | * | |
| 226_2_2 | 226 | 2 | 2 | | #880E4F | Tspelta_89420 | * | |
| 226_2_3 | 226 | 2 | 3 | | #EC407A | Osativa_4456 | * | |
| 226_2_5 | 226 | 2 | 5 | | #F48FB1 | Zmays_27587 | * | |
| 226_2_6 | 226 | 2 | 6 | | [illegible] | Osativa_8390 | * | |
| 226_2_7 | 226 | 2 | 7 | | [illegible] | Osativa_27583 | * | |
| 226_4 | 226 | 4 | | | #F06292 | Osativa_1813 | * | |
| 227_4 | 227 | 4 | | | [illegible] | Taestivum_44475 | * | |
| 227_5 | 227 | 5 | | | [illegible] | Osativa_41853 | * | |
| 228 | 228 | | | | #F8BBD0 | Pvirgatum_31937 | * | |
| 229 | 229 | | | | #F48FB1 | Ufusca_834 | * | |
| 230 | 230 | | | | [illegible] | Osativa_21331 | * | |
| 232_2 | 232 | 2 | | | #FCE4EC | Bhybridum_43014 | * | |
| 233 | 233 | | | | [illegible] | Hvulgare_37792 | * | |
| 244 | 244 | | | | #FF5252 | Bamplexicaulis_14614 | * | |
| 245 | 245 | | | | [illegible] | Claurum_20891 | * | |
| 246_1_3 | 246 | 1 | 3 | | #FF8A65 | Taestivum_87761 | * | |
| 246_2 | 246 | 2 | | | #FFCCBC | Osativa_31351 | * | |
| 247_2 | 247 | 2 | | | #FFAB91 | Osativa_13361 | * | |
| 247_4_1 | 247 | 4 | 1 | | #FF7043 | Osativa_3641 | * | |
| 247_6 | 247 | 6 | | | #F4511E | Osativa_29734 | * | |
| 249_2 | 249 | 2 | | | #FBE9E7 | Olongistaminata_13456 | * | |
| 250_2 | 250 | 2 | | | [illegible] | Omeyera_7548 | * | |
| 259_1 | 259 | 1 | | | #6D4C41 | Bdistachyon_27355 | * | |
| 259_3_3 | 259 | 3 | 3 | | [illegible] | Phallacum_51932 | * | |
| 259_4 | 259 | 4 | | | #5D4037 | Osativa_33062 | * | |
| 260_2 | 260 | 2 | | | #A1887F | Osativa_9873 | * | |
| 260_3 | 260 | 3 | | | #BCAAA4 | Taestivum_46158 | * | |
| 262_2 | 262 | 2 | | | [illegible] | Taestivum_82124 | * | |
| 263_1_1 | 263 | 1 | 1 | | [illegible] | Taestivum_52848 | * | |
| 263_1_2 | 263 | 1 | 2 | | #FF6E40 | Osativa_3751 | * | |
| 263_1_2 | 263 | 1 | 2 | | #FF8A4C | Osativa_33128 | * | |
| 263_1_4 | 263 | 1 | 4 | | [illegible] | Obrachyantha_31449 | * | |
| 263_2_4 | 263 | 2 | 4 | | #FFFDE7 | Osativa_28507 | * | |
| 263_3_1 | 263 | 3 | 1 | | #FFF9C4 | Bdistachyon_6221 | * | |
| 263_3_3 | 263 | 3 | 3 | | #FFF3E0 | Taestivum_1008 | * | |
| 263_4_4 | 263 | 4 | 4 | | [illegible] | Osativa_41783 | * | |
| 263_5 | 263 | 5 | | | #FF9E80 | Osativa_33866 | * | |
| 263_5 | 263 | 5 | | | #FF9E80 | Osativa_2206 | * | |
| 264_2 | 264 | 2 | | | [illegible] | Osativa_33883 | * | |
| 275 | 275 | | | | #E0F7FA | Pintrocarpa_3974 | | * |
| 278 | 278 | | | | #4DD0E1 | Mabacum_52998 | | * |
| 281 | 281 | | | | #26C6DA | Slycopersicum_7338 | | * |
| 286_1 | 286 | 1 | | | [illegible] | Gmustelinum_88032 | | * |
| 296 | 296 | | | | #00838F | Pintrocarpa_5728 | | * |
| 339_1 | 339 | 1 | | | #18FFFF | Mabacum_28722 | | * |
| 365_5 | 365 | 5 | | | #006064 | Mabacum_59351 | | * |
| 366 | 366 | | | | #84FFFF | Nbenthamiana_31200 | | * |
| 376 | 376 | | | | #0097A7 | Cdentata_21751 | | * |
| 380 | 380 | | | | #80DEEA | Slycopersicum_2297 | | * |
| 412_1 | 412 | 1 | | | #B2DFDB | Bdistachyon_16887 | * | |
| 412_2 | 412 | 2 | | | #4DB6AC | Bdistachyon_20821 | * | |
| 412_3 | 412 | 3 | | | #26A69A | Dsorghi_50483 | * | |
| 412_4 | 412 | 4 | | | #80CBC4 | Zmays_06670 | * | |
| 413_2 | 413 | 2 | | | #E0F2F1 | Sbicolor_1484 | * | |
| 415 | 415 | | | | #00B8D4 | Osativa_17387 | * | |
| 416_1 | 416 | 1 | | | #1DE9B6 | Sbicolor_18494 | * | |
| 417_3_1 | 417 | 3 | 1 | | #81C784 | Osativa_19992 | * | |
| 417_3_2 | 417 | 3 | 2 | | #A5D6A7 | Osativa_39700 | * | |
| 417_3_4 | 417 | 3 | 4 | | #66BB6A | Osativa_6969 | * | |
| 417_5_3 | 417 | 5 | 3 | | #C8E6C9 | Bdistachyon_21788 | * | |
| 417_6_5 | 417 | 6 | 5 | | #E8F5E9 | Bdistachyon_13131 | * | |
| 418_1_5 | 418 | 1 | 5 | | [illegible] | Osativa_22723 | * | |
| 418_5_1 | 418 | 5 | 1 | | #00E5FF | Osativa_24262 | * | |
| 418_6_1 | 418 | 6 | 1 | | [illegible] | Osativa_19822 | * | |
| 418_6_1 | 418 | 6 | 1 | | [illegible] | Osativa_22083 | * | |
| 419_1_4 | 419 | 1 | 4 | | #64FFDA | Tspelta_48632 | * | |
| 419_1_6 | 419 | 1 | 6 | | #A7FFEB | Tspelta_42565 | * | |
| 419_3_1 | 419 | 3 | 1 | | [illegible] | Osativa_9108 | * | |
| 419_3_2 | 419 | 3 | 2 | | #004D40 | Bhybridum_2145 | * | |
| 419_3_3 | 419 | 3 | 3 | | [illegible] | Bdistachyon_6055 | * | |
| 437 | 437 | | | | #F5F5F5 | Gmax_2481 | | * |
| 458 | 458 | | | | #B0BEC5 | Osativa_9147 | * | |
| 460 | 460 | | | | #EEEEEE | Cnuciferа_19391 | * | |
| 461 | 461 | | | | #E0E0E0 | Gmax_38068 | * | * |
| 461 | 461 | | | | #E0E0E0 | Mabacum_47195 | * | * |
| 462 | 462 | | | | #CFD8DC | Osativa_13157 | * | |
| 463 | 463 | | | | #FAFAFA | Zmays_18587 | * | |
| 475 | 475 | | | | [illegible] | Tintermedium_125318 | * | |
| 514_2 | 514 | 2 | | | #CE93D8 | Glomerosum_11890 | | * |
| 514_4 | 514 | 4 | | | #BA68C8 | Gharsedesa_75615 | | * |
| 559 | 559 | | | | #E1BEE7 | Cdarwinii_16752 | | * |
| 606_6 | 606 | 6 | | | [illegible] | Cdarwinii_66762 | | * |
| 611 | 611 | | | | #6200EA | Ppersica_16486 | | * |
| 614 | 614 | | | | [illegible] | Glomeratum_40120 | | * |
| 616 | 616 | | | | [illegible] | Cdarwinii_9277 | | * |
| 627_3_1 | 627 | 3 | 1 | | [illegible] | Sdunnii_4655 | | * |
| 628_3 | 628 | 3 | | | [illegible] | Pdeltoides_3331 | | * |
| 632_1 | 632 | 1 | | | #F8E7F7 | Pasutifolius_13132 | | * |
| 632_2 | 632 | 2 | | | #F3D3F5 | Gmax_3327 | | * |
| 632_3 | 632 | 3 | | | #D3BAE8 | Gmax_12656 | | * |
| 633_6 | 633 | 6 | | | [illegible] | Gmax_7821 | | * |
| 634_3 | 634 | 3 | | | #A8A8DD | Gmax_19484 | | * |
| 634_4 | 634 | 4 | | | #CCEDE4 | Gmax_36734 | | * |
| 640 | 640 | | | | #445C94 | Mabacum_67357 | | * |
| 640 | 640 | | | | #445C94 | Slycopersicum_5548 | | * |
| 658_2 | 658 | 2 | | | [illegible] | Itinctoria_19485 | | * |
| 658_3 | 658 | 3 | | | #4527A0 | Athaliana_1095 | | * |
| 660_3_3 | 660 | 3 | 3 | | #311B92 | Crubella_39859 | | * |
| 666_2_2 | 666 | 2 | 2 | | #7C4DFF | Nbenthamiana_33780 | | * |
| 666 | 666 | | | | [illegible] | Cnubra_33555 | | * |
| 686 | 686 | | | | #EDE7F6 | Talseanea_22447 | | * |
| 689_3 | 689 | 3 | | | [illegible] | Pdeltoides_16788 | | * |
| 689_4 | 689 | 4 | | | [illegible] | Glomeratum_14062 | | * |
| 689_5 | 689 | 5 | | | #9575CD | Mabacum_11998 | | * |
| 689_7 | 689 | 7 | | | #B39DDB | Gmax_46162 | | * |
| 696 | 696 | | | | #D1C4E9 | Osativa_5225 | * | |
| 697 | 697 | | | | #E040FB | Cdarwinii_68169 | | * |
| 700_1 | 700 | 1 | | | [illegible] | Gmax_26453 | | * |
| 700_3 | 700 | 3 | | | [illegible] | Rcommunis_28446 | | * |
| 710 | 710 | | | | #EA80FC | Turartu_16320 | * | |
| 710 | 710 | | | | #EA80FC | Taestivum_61720 | * | |
| 711_2 | 711 | 2 | | | #4A148C | Taestivum_34267 | * | |
| 713_1_7 | 713 | 1 | 7 | | #7B1FA2 | Tintermedium_21038 | * | |
| 713_3_7 | 713 | 3 | 7 | | #6A1B9A | Taestivum_67548 | * | |
| 713_5 | 713 | 5 | | | #8E24AA | Taestivum_21527 | * | |
| 737 | 737 | | | | #424242 | Athaliana_24101 | | * |
| 739_3 | 739 | 3 | | | #37474F | Gmax_30865 | | * |
| 739_5 | 739 | 5 | | | #37474F | Gmax_28544 | | * |
| 739_7_4 | 739 | 7 | 4 | | #212121 | Mabacum_67221 | | * |
| 740_1 | 740 | 1 | | | [illegible] | Taestivum_65488 | * | |
| 740_2 | 740 | 2 | | | [illegible] | Osativa_1914 | * | |
| 754 | 754 | | | | #BDBDBD | Taestivum_48164 | * | |

**Fig. S4. Information on selected 210 LRR-RLK-XII groups.** The table represents information on the 210 selected LRR-RLK-XII groups for the BES1 screening. From left to right: group number (name of each group), group number within each split or cluster ($1^{st}$ to $4^{th}$), colour code for indication in Fig 1D and 2F, name of representative receptors cloned for the screening, and the distribution of selected LRR-RLK-XIIs subgroups across the angiosperm lineage. Within each subgroup, the presence of receptors within superclades, clades and orders are labelled with circles (●), squares (■), and stars (★), respectively.

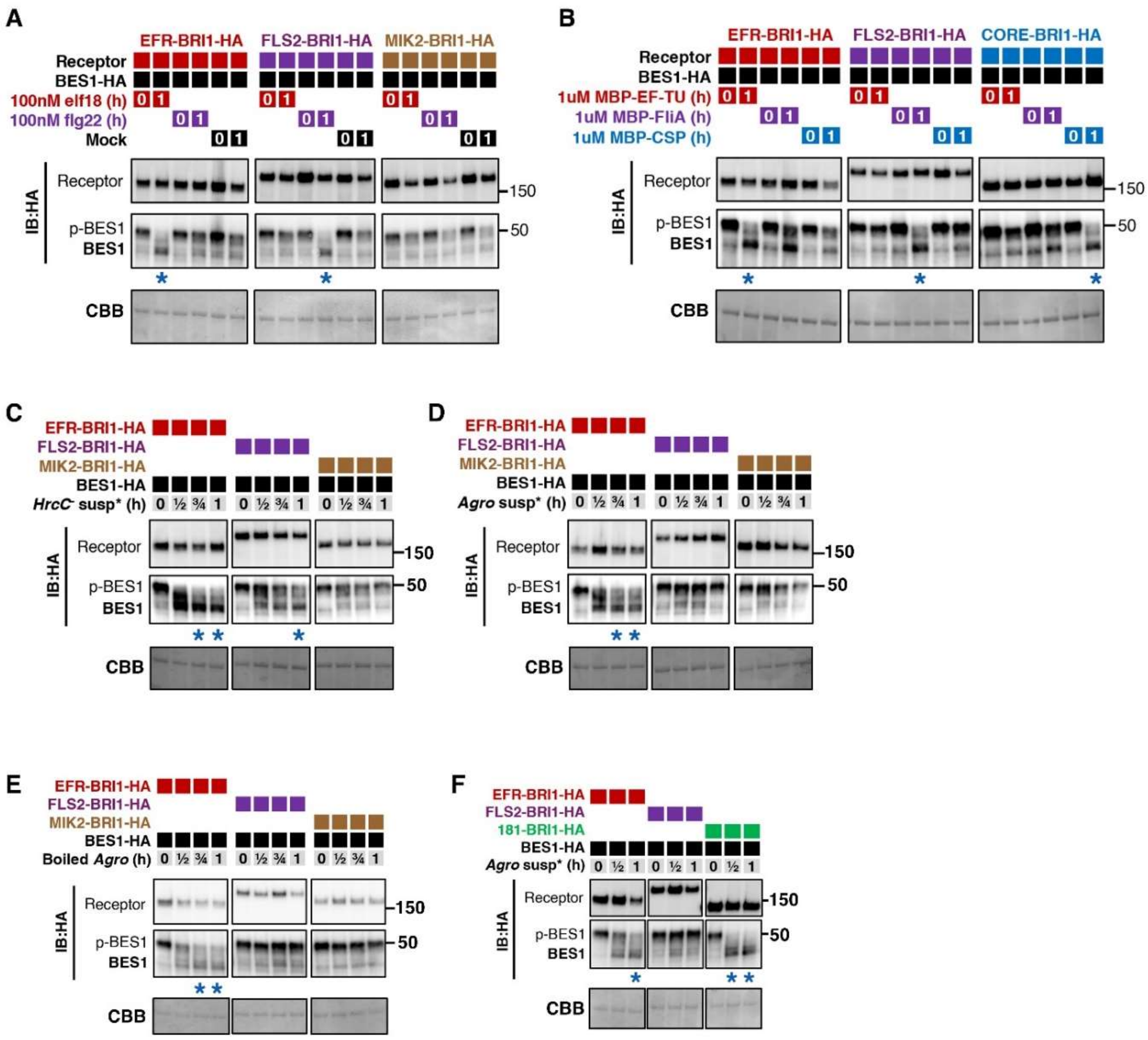


**Fig. S5. Coomassie blue staining for immunoblots in Fig 2.**

Due to space limitations, Coomassie blue staining for immunoblots in Fig 2 are shown here. **(A)** corresponds to Fig 2B, **(B)** corresponds to Fig 2C, **(C-D)** corresponds to Fig 2D, **(E)** corresponds to Fig 2E, and **(F)** corresponds to Fig 2G.

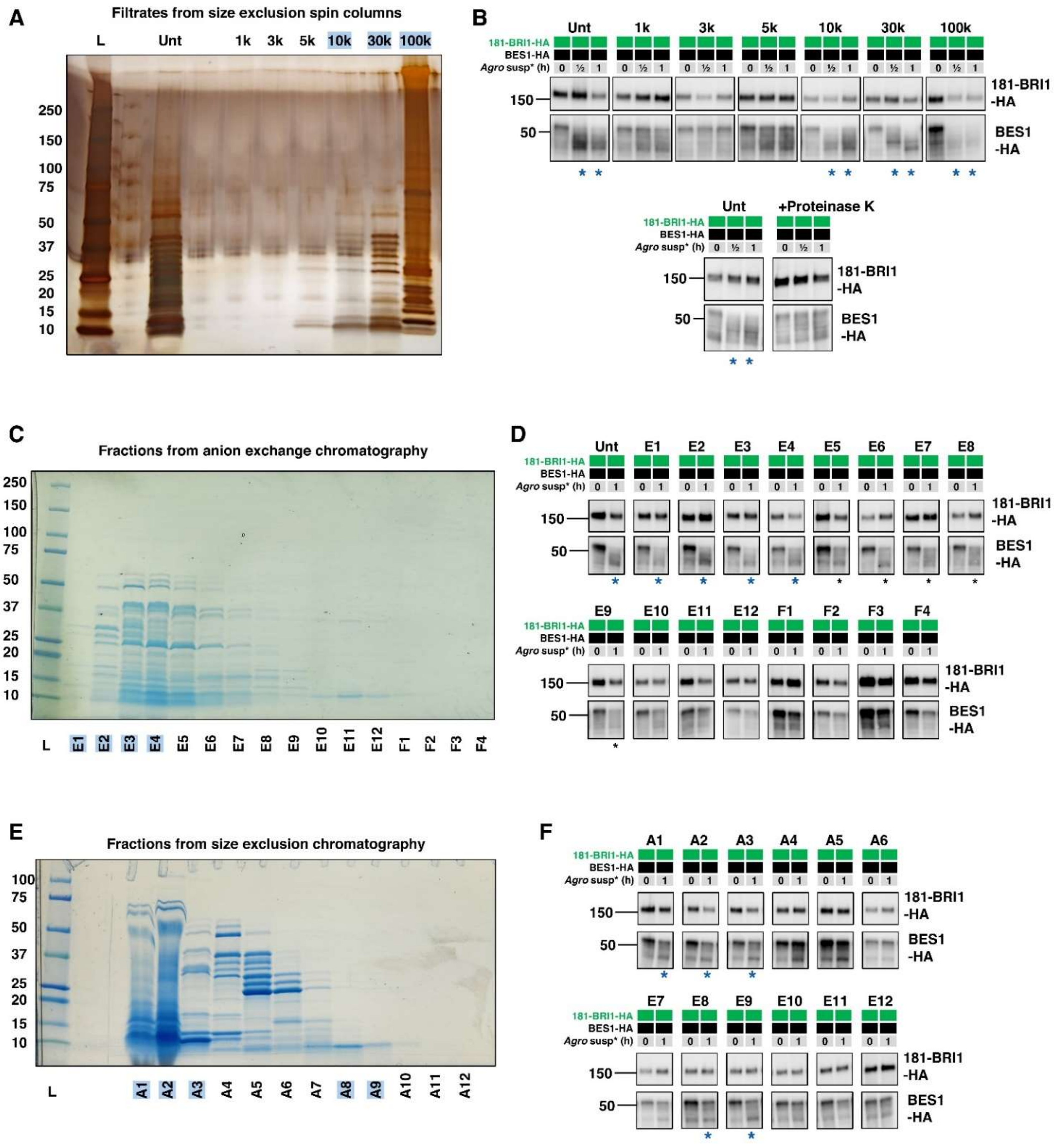


**Fig. S6. Activity of *Agrobacterium* fractions through 181-BRI1.**

**(A-B)** correspond to Fig 3A. **(A)** Silver staining of proteins in *Agrobacterium* extract filtrates from size exclusion spin columns. **(B)** The activity of fractions in (A) were tested with 181-BRI1. To confirm that 181-BRI1 perceives a protein ligand, Agrobacterium extract was treated with Proteinase K. Samples were collected at indicated time points, and BES1 dephosphorylation was checked with immunoblotting. Asterisks denote dephosphorylation of BES1. **(C-D)** correspond to Fig 3B. **(A)** CBB staining of proteins in *Agrobacterium* extract fractions obtained from anion exchange chromatography. **(D)** The activity of fractions shown in (C) were tested with 181-BRI1. Samples were collected at indicated time points, and BES1 dephosphorylation was checked with immunoblotting. Asterisks denote dephosphorylation of BES1. **(E-F)** correspond to Fig 3C. **(E)** CBB staining of proteins in *Agrobacterium* extract fractions obtained from size exclusion chromatography. **(F)** The activity of fractions in (E) were tested with 181-BRI1. Samples were collected at indicated time points, and BES1 dephosphorylation was checked with immunoblotting. Asterisks denote dephosphorylation of BES1.

| Protein FDR Confidence (Combined) | Accession | Description | Protein length | MW (kDa) | pI | Abundance Ratio: (8) / (6) | Abundance Ratio: (9) / (6) | Abundance Ratio: (8) / (7) | Abundance Ratio: (9) / (7) | Abundance (Grouped): 6 | Abundance (Grouped): 7 | Abundance (Grouped): 8 | Abundance (Grouped): 9 |
|---|---|---|---|---|---|---|---|---|---|---|---|---|---|
| High | AAK85925.2 | cold shock protein | 69 | 7.4 | 5.95 | 3.96 | 41.84 | 0.68 | 7.18 | 7.6 | 44.3 | 30.1 | 318 |
| High | AAK85847.2 | thioredoxin | 106 | 11.1 | 4.67 | 256.08 | 9.23 | 6.19 | 0.22 | 1.3 | 53.8 | 332.9 | 12 |
| High | AAK90263.2 | cold shock protein | 69 | 7.5 | 8.62 | 4.56 | 7.72 | 5.36 | 9.06 | 28.3 | 24.1 | 129.1 | 218.4 |
| High | AAK88726.2 | conserved hypothetical protein | 67 | 8.2 | 6.77 | 3.07 | 6.94 | 2.86 | 6.47 | 33.1 | 35.5 | 101.7 | 229.6 |
| High | AAK87945.1 | cold shock protein | 71 | 7.5 | 7.25 | 4.46 | 4.77 | 2.19 | 2.34 | 32.6 | 66.5 | 145.5 | 155.5 |
| High | AAK89888.2 | conserved hypothetical protein | 259 | 30.1 | 9.55 | 3.13 | 3.88 | 0.50 | 0.62 | 28.1 | 175 | 87.9 | 109.1 |
| High | AAK89289.1 | hypothetical protein Atu4143 | 73 | 8.8 | 4.82 | 15.52 | 1.37 | 2.89 | 0.25 | 17.2 | 92.3 | 267 | 23.5 |
| Medium | AAK86076.2 | aldo-keto reductase | 333 | 35.7 | 5.95 | 0.67 | 1.22 | 0.12 | 0.21 | 46.3 | 266.3 | 31 | 56.4 |
| High | AAK87618.2 | BolA/YrbA family protein | 77 | 8.2 | 6.28 | 3.73 | 0.91 | 2.05 | 0.50 | 53.6 | 97.6 | 199.8 | 48.9 |
| High | AAL44894.1 | hypothetical protein Atu4093 | 62 | 7.1 | 4.7 | 2.28 | 0.85 | 3.49 | 1.29 | 83.7 | 54.7 | 190.7 | 70.8 |
| High | AAK87119.2 | conserved hypothetical protein | 145 | 16.1 | 7.14 | 43.21 | 0.83 | 1.36 | 0.03 | 5.2 | 165.8 | 224.7 | 4.3 |
| High | AAK87479.1 | SEC-independent protein translocase protein | 70 | 7.6 | 7.42 | 3.57 | 0.75 | 1.61 | 0.34 | 53 | 117.8 | 189.4 | 39.8 |
| High | AAK85941.1 | molecular chaperone, DnaJ family | 377 | 40.9 | 7.59 | 3.67 | 0.74 | 1.29 | 0.26 | 48.5 | 137.5 | 178 | 36 |
| High | AAK88440.1 | porin | 220 | 22.6 | 9.2 | 0.73 | 0.57 | 0.56 | 0.44 | 110.8 | 144.6 | 81.2 | 63.4 |
| High | AAK85904.1 | 30S ribosomal protein S15 | 89 | 10 | 10.13 | 0.70 | 0.57 | 1.58 | 1.28 | 147.4 | 65.4 | 103.5 | 83.7 |
| High | AAK86356.2 | flaB | 320 | 33 | 4.88 | 0.54 | 0.54 | 0.42 | 0.42 | 118.5 | 152.9 | 64.1 | 64.5 |
| High | AAK89225.1 | cold shock protein | 76 | 8.1 | 6.52 | 0.07 | 0.51 | 0.37 | 2.62 | 224.7 | 43.9 | 16.3 | 115.2 |
| High | AAK87834.2 | conserved hypothetical protein | 288 | 32.1 | 8.87 | 0.97 | 0.50 | 2.40 | 1.24 | 139 | 56.3 | 135.1 | 69.6 |
| High | AAK89098.2 | conserved hypothetical protein | 169 | 19.1 | 5.11 | 0.50 | 0.49 | 1.26 | 1.22 | 167.7 | 66.7 | 84.2 | 81.4 |
| High | AAK88419.1 | 30S ribosomal protein S16 | 126 | 13.7 | 10.04 | 0.50 | 0.46 | 0.87 | 0.79 | 158.3 | 90.8 | 78.9 | 72.1 |
| High | AAK89009.1 | ABC transporter, substrate binding protein (oligopeptide) | 545 | 59.4 | 5.95 | 0.44 | 0.44 | 1.13 | 1.13 | 176 | 68.8 | 77.8 | 77.4 |
| High | AAK89994.1 | ABC transporter, substrate binding protein (iron) | 343 | 36.9 | 5.26 | 0.33 | 0.43 | 0.69 | 0.89 | 177.9 | 86.1 | 59.3 | 76.8 |
| High | AAK90908.2 | ABC transporter, substrate binding protein (plasmid) | 416 | 45.3 | 6.7 | 0.56 | 0.43 | 1.01 | 0.78 | 157.3 | 87.1 | 87.7 | 67.9 |
| High | AAK89102.2 | conserved hypothetical protein | 169 | 19 | 6.24 | 1.17 | 0.41 | 0.83 | 0.29 | 100.4 | 141 | 117.1 | 41.5 |
| Medium | AAK86357.1 | flagella associated protein | 306 | 31.6 | 4.97 | 0.36 | 0.40 | 0.77 | 0.85 | 179.5 | 84.2 | 64.6 | 71.7 |
| High | AAL42103.1 | 30S ribosomal protein S18 | 82 | 9.4 | 11.4 | 0.14 | 0.36 | 0.42 | 1.05 | 217 | 74 | 31.2 | 77.9 |
| High | AAK87947.2 | conserved hypothetical protein | 199 | 21.6 | 4.64 | 3.52 | 0.35 | 1.93 | 0.19 | 59.8 | 108.9 | 210.2 | 21.1 |
| High | AAK87683.1 | 50S ribosomal protein L17 | 141 | 15.4 | 10.45 | 0.73 | 0.35 | 1.14 | 0.54 | 146.7 | 94.4 | 107.6 | 51.3 |
| High | AAK86194.2 | conserved hypothetical protein | 198 | 21.7 | 8.81 | 0.67 | 0.35 | 0.90 | 0.47 | 145.4 | 107.6 | 96.7 | 50.3 |
| High | AAK88464.2 | conserved hypothetical protein | 163 | 17.5 | 9.48 | 0.43 | 0.32 | 1.28 | 0.94 | 191.5 | 64.7 | 82.8 | 61 |
| High | AAL42104.1 | 30S ribosomal protein S6 | 153 | 17.8 | 9 | 0.48 | 0.32 | 0.28 | 0.18 | 114 | 195.5 | 54.5 | 36.1 |
| Medium | AAK86139.1 | 30S ribosomal protein S20 | 88 | 9.5 | 12.07 | 0.57 | 0.31 | 2.10 | 1.13 | 186.2 | 50.5 | 106.1 | 57.3 |
| High | AAK89692.1 | omp16 protein | 177 | 18.8 | 9.47 | 0.25 | 0.30 | 0.55 | 0.67 | 199.6 | 90 | 49.6 | 60.7 |
| High | AAK88363.1 | conserved hypothetical protein | 180 | 18.4 | 8.66 | 0.47 | 0.29 | 0.92 | 0.56 | 176.7 | 90.1 | 82.8 | 50.5 |
| High | AAL41919.2 | dnaK suppressor protein | 106 | 12.1 | 4.96 | 0.34 | 0.28 | 0.93 | 0.75 | 201.4 | 74.3 | 68.9 | 55.4 |
| High | AAK87643.2 | OmpA family protein | 746 | 82.4 | 5.48 | 0.24 | 0.24 | 0.66 | 0.66 | 216.1 | 79.2 | 52.5 | 52.3 |
| High | AAK86703.1 | conserved hypothetical protein | 133 | 15.2 | 8.37 | 4.49 | 0.24 | 0.80 | 0.04 | 35.3 | 197.6 | 158.5 | 8.5 |
| High | AAK87869.1 | lytic murein transglycosylase | 408 | 43.7 | 6.19 | 0.71 | 0.24 | - | - | 205.8 |  | 145.3 | 48.9 |
| High | AAK86496.2 | conserved hypothetical protein | 209 | 22.9 | 9.14 | 0.22 | 0.23 | 1.17 | 1.19 | 243.7 | 46.6 | 54.5 | 55.3 |
| Medium | AAK88682.1 | conserved hypothetical protein | 145 | 15 | 8.53 | 0.09 | 0.20 | 0.36 | 0.79 | 256.9 | 66.7 | 23.8 | 52.6 |
| High | AAK89646.1 | conserved hypothetical protein | 179 | 20.1 | 6.86 | 5.05 | 0.21 | 0.17 | 0.01 | 11.2 | 330 | 56.6 | 2.3 |
| High | AAK86146.1 | GRPE protein | 211 | 22.8 | 4.77 | 1.60 | 0.20 | 0.58 | 0.07 | 71.8 | 199.2 | 114.7 | 14.3 |
| High | AAK87542.2 | conserved hypothetical protein | 218 | 22.8 | 9.06 | 0.97 | 0.19 | 1.46 | 0.28 | 141.5 | 94.5 | 137.7 | 26.3 |
| High | AAK87602.1 | ATP-dependent RNA helicase | 503 | 55.8 | 8.56 | 0.81 | 0.15 | - | - | 203.9 |  | 164.5 | 31.5 |
| High | AAL44562.1 | conserved hypothetical protein | 58 | 6.6 | 8.56 | 0.56 | 0.16 | 0.86 | 0.24 | 169.3 | 109.8 | 94.6 | 26.3 |
| High | AAK89663.1 | 50S ribosomal protein L31 | 73 | 8.1 | 8.56 | 0.64 | 0.13 | 0.39 | 0.08 | 117.7 | 191.6 | 75 | 15.8 |
| High | AAK89695.2 | metalloprotease | 648 | 70.5 | 5.87 | 1.98 | 0.12 | 0.55 | 0.03 | 59.7 | 214.7 | 118.5 | 7.2 |
| High | AAK87877.2 | conserved hypothetical protein | 113 | 11.9 | 5.14 | 0.10 | 0.10 | 0.38 | 0.39 | 274.7 | 70.8 | 26.7 | 27.8 |
| High | AAK89318.1 | ABC transporter, substrate binding protein (dipeptide) | 534 | 59.1 | 5.39 | 0.13 | 0.09 | 0.50 | 0.36 | 270.4 | 69.4 | 34.8 | 25.3 |
| High | AAK87730.2 | conserved hypothetical protein | 149 | 15.9 | 9.11 | 0.43 | 0.09 | 0.98 | 0.21 | 204.8 | 89.2 | 87.5 | 18.6 |
| High | AAK87360.2 | ABC transporter, substrate binding protein (amino acid) | 312 | 33.9 | 5.22 | 0.12 | 0.09 | 0.56 | 0.43 | 282.3 | 59.2 | 33.1 | 25.4 |
| High | AAK89884.1 | glutaredoxin | 100 | 10.9 | 5.36 | 3.77 | 0.09 | 0.34 | 0.01 | 24.9 | 279.1 | 93.8 | 2.2 |
| High | AAK87057.1 | histone-like protein | 91 | 9.3 | 9.19 | 0.18 | 0.08 | 0.73 | 0.34 | 265.3 | 65.1 | 47.6 | 22 |
| High | AAK88804.2 | hypothetical protein Atu4643 | 58 | 6.3 | 7.25 | 0.12 | 0.08 | 0.07 | 0.05 | 132.3 | 240.5 | 16.4 | 10.9 |
| High | AAK88158.2 | ABC transporter, substrate binding protein (amino acid) | 372 | 38.8 | 5.36 | 0.06 | 0.07 | 0.48 | 0.63 | 320.3 | 37.8 | 18 | 23.9 |
| High | AAK89877.2 | peptidyl-prolyl cis-trans isomerase | 288 | 32.1 | 5.35 | 0.13 | 0.07 | 0.49 | 0.27 | 272.1 | 72.8 | 35.6 | 19.5 |
| High | AAK89180.2 | ABC transporter, substrate binding protein (oligopeptide) | 531 | 58.8 | 5.22 | 0.09 | 0.07 | 0.33 | 0.25 | 276 | 78.3 | 26 | 19.7 |
| High | AAK88918.1 | ABC transporter, substrate binding protein (oligopeptide) | 511 | 55.9 | 5.67 | 0.21 | 0.06 | 0.26 | 0.08 | 189.4 | 157.6 | 40.7 | 12.3 |
| High | AAK88731.1 | ABC transporter, substrate binding protein (dipeptide) | 502 | 55.1 | 5.55 | 0.08 | 0.06 | 0.36 | 0.29 | 297.1 | 62.2 | 22.6 | 18.1 |
| High | AAK88025.2 | pseudoazurin | 150 | 15.7 | 7.96 | 0.18 | 0.06 | 0.51 | 0.16 | 252.2 | 88.4 | 45.3 | 14.1 |
| High | AAK89365.2 | O-linked GlcNAc transferase | 298 | 32.7 | 8.51 | 0.22 | 0.05 | 1.02 | 0.26 | 270.3 | 56.8 | 58.2 | 14.7 |
| High | AAK87437.2 | trigger factor | 492 | 54.1 | 4.84 | 0.19 | 0.05 | 0.49 | 0.13 | 243.8 | 96 | 47.5 | 12.7 |
| High | AAK86683.1 | superoxide dismutase | 200 | 22.6 | 6.1 | 0.02 | 0.05 | 0.21 | 0.46 | 340.3 | 35.7 | 7.5 | 16.5 |
| High | AAK87991.1 | conserved hypothetical protein | 61 | 7 | 6.28 | 0.88 | 0.04 | 0.16 | 0.01 | 54.2 | 296.1 | 47.6 | 2.1 |
| High | AAK86504.1 | conserved hypothetical protein | 79 | 9.1 | 6.52 | 0.05 | 0.04 | 0.14 | 0.10 | 274.8 | 101.2 | 13.9 | 10.1 |
| High | AAK87292.2 | hypothetical protein Atu1501 | 114 | 12.6 | 10.45 | 0.11 | 0.04 | 0.51 | 0.17 | 295.9 | 62.1 | 31.4 | 10.6 |
| High | AAK86234.1 | ABC transporter, substrate binding protein (phosphate) | 344 | 36.2 | 5.52 | 0.07 | 0.04 | 0.91 | 0.44 | 336.9 | 26.9 | 24.4 | 11.8 |
| High | AAK89607.2 | ABC transporter, substrate binding protein (proline/glycine betaine) | 345 | 37 | 5.1 | 0.02 | 0.03 | 0.51 | 0.63 | 361.8 | 17.8 | 9 | 11.3 |
| High | AAK87894.1 | ABC transporter, substrate binding protein | 368 | 39 | 5.45 | 0.03 | 0.03 | 0.57 | 0.61 | 359.9 | 18.4 | 10.5 | 11.2 |
| High | AAK89999.1 | ABC transporter, substrate binding protein (iron) | 377 | 41.7 | 6.09 | 0.04 | 0.03 | 0.71 | 0.52 | 356.3 | 19.6 | 14 | 10.1 |
| High | AAK87559.2 | lipoprotein | 357 | 38.8 | 5.99 | 0.05 | 0.03 | 0.63 | 0.32 | 342.5 | 29.5 | 18.6 | 9.4 |
| High | AAK87928.1 | transcription elongation facto | 158 | 17.3 | 4.92 | 0.10 | 0.03 | 0.28 | 0.08 | 271.7 | 94.6 | 26.4 | 7.3 |
| High | AAK89373.2 | endo-1,3-1,4-beta-glycanase | 263 | 29.4 | 6.23 | 0.05 | 0.02 | 0.15 | 0.08 | 290.1 | 89.5 | 13.2 | 7.2 |
| High | AAK86905.1 | conserved hypothetical protein | 406 | 44.5 | 6.47 | 2.21 | 0.03 | 0.33 | 0.00 | 39.9 | 271 | 88.1 | 1 |
| High | AAK86420.1 | ABC transporter, substrate binding protein (putrescine) | 365 | 40.2 | 5.3 | 0.03 | 0.02 | 0.50 | 0.36 | 356.4 | 23.4 | 11.7 | 8.5 |

Molecular weight (kDa): min – max; pI: 14 – 7 – 1; Abundance ratio: min – max; Abundance (relative): min – max

**Fig. S7. MS analysis from fractions A6-A9 in Fig 3C.**

The table provides detailed information on the proteins identified from fractions A6 to A9 in Fig. 3C. The columns, from left to right, include: Protein false discovery rate (FDR) confidence, Accession number of *Agrobacterium tumefaciens Agl1* proteins identified, protein description, protein length (number of amino acids), molecular weight (MW; in kDa), pI value of the full-length protein, protein abundance ratio for A8/A6, A9/A6, A8/A7, A9/A7, and relative protein abundance in fraction A6, A7, A8 and A9. Since 181-BRI1 responds specifically to fractions A8 and A9, the table is ranked by the protein abundance ratios A8/A7 and A9/A7. Cold-shock proteins are highlighted in yellow. For full list of identified peptides, refer to supplementary table 1.

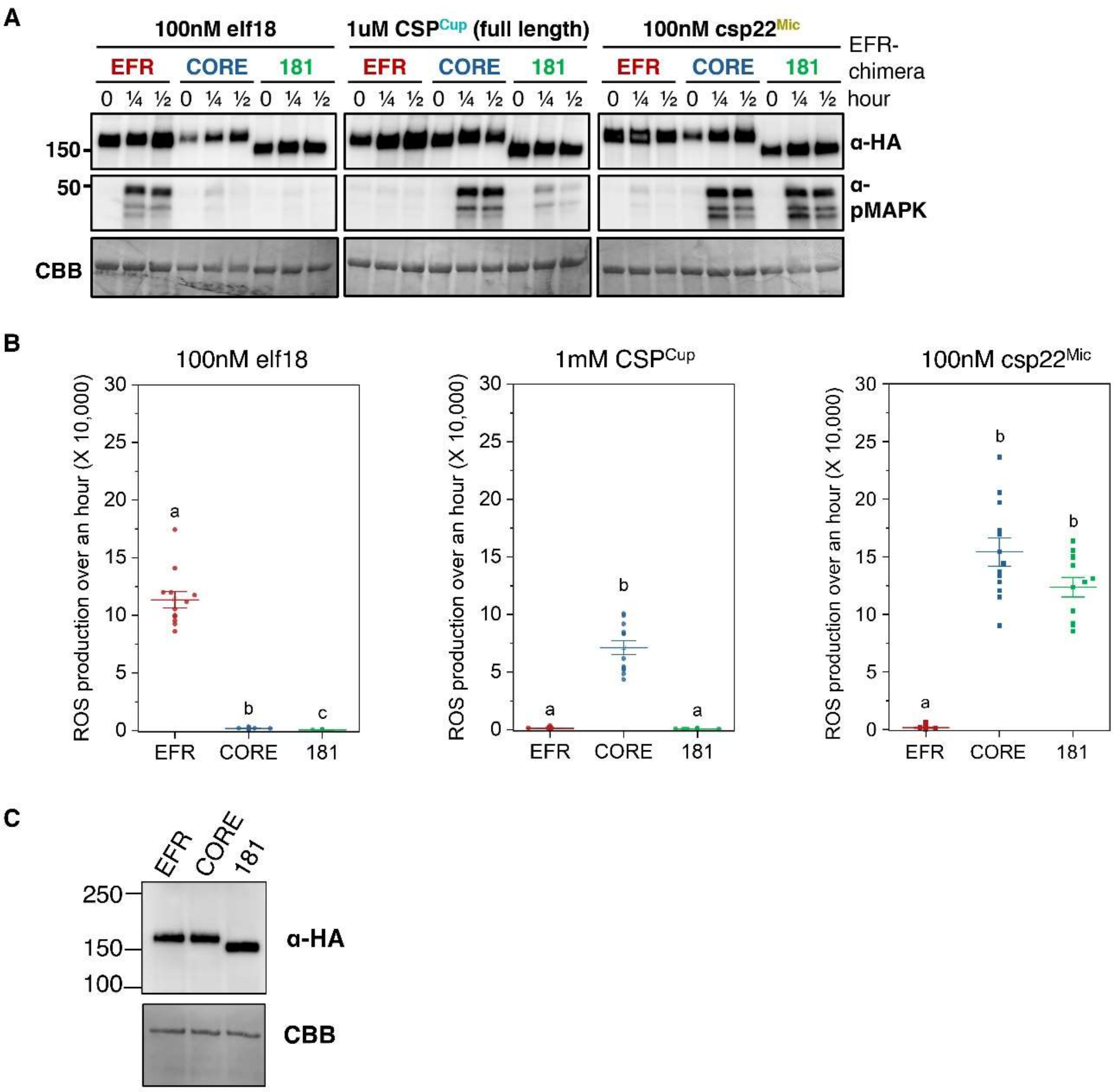


**Fig. S8. Data related to Fig 3E.**

**(A)** Due to space constraints, the CBB-stained blots corresponding to the immunoblots shown in Fig. 3E are displayed here. **(B)** ROS production in Fig 3E. Box plots represent ROS production by EFR, CORE-EFR or 181-EFR over an hour following mock, full-length CSP or csp22$^{Mic}$ peptide treatments. In the plots, the center line represents the median, while the upper and lower lines indicate the 25th and 75th percentiles, respectively. Data points from 12 technical replicates ($n = 12$) were analyzed using a one-sided Kruskal-Wallis test followed by Dunn's multiple comparisons test. Different letters above the data points denote significant differences ($P < 0.05$). **(C)** Samples used in the ROS assays in (B) were analysed with immunoblotting to confirm receptor expression. CBB staining is included as loading controls.

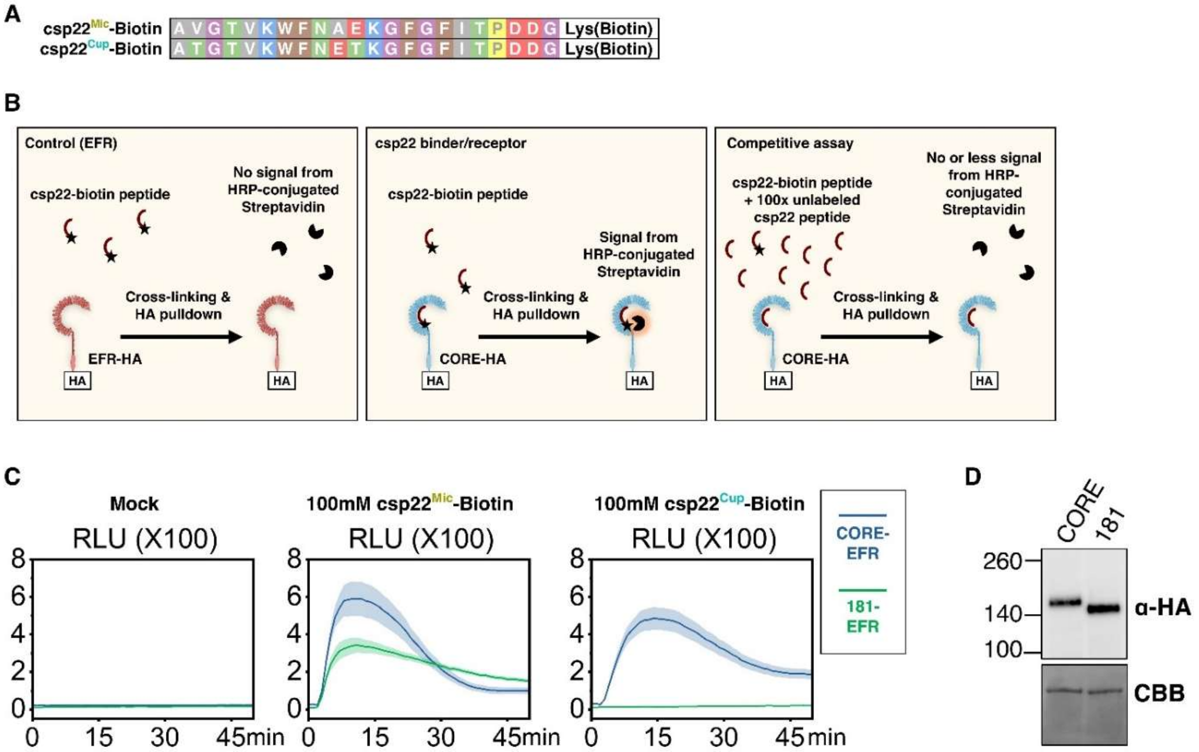


**Fig. S9. Data related to Fig 3G.**

**(A)** Amino acid sequence of csp22$^{Mic}$-Biotin and csp22$^{Cup}$-Biotin peptides. **(B)** Schematic representation of biotinylated peptide cross-linking assay. Biotinylated peptide were cross-linked to their corresponding receptor *via* ethylene glycol bis(succinimidyl succinate) (EGS). After cross-linking, the receptors are pulled down, and biotinylated peptides are detected using HRP-conjugated streptavidin. A competitive assay is included as a control, where an excessive unlabelled peptide (100-fold) is used to outcompete the binding of biotinylated peptide. **(C)** ROS production by CORE-EFR and 181-EFR over an hour following mock, csp22$^{Mic}$-Biotin, and csp22$^{Cup}$-Biotin peptide. **(D)** Samples used in the ROS assays in (B) were analysed with immunoblotting to confirm receptor expression. CBB staining is included as loading controls.

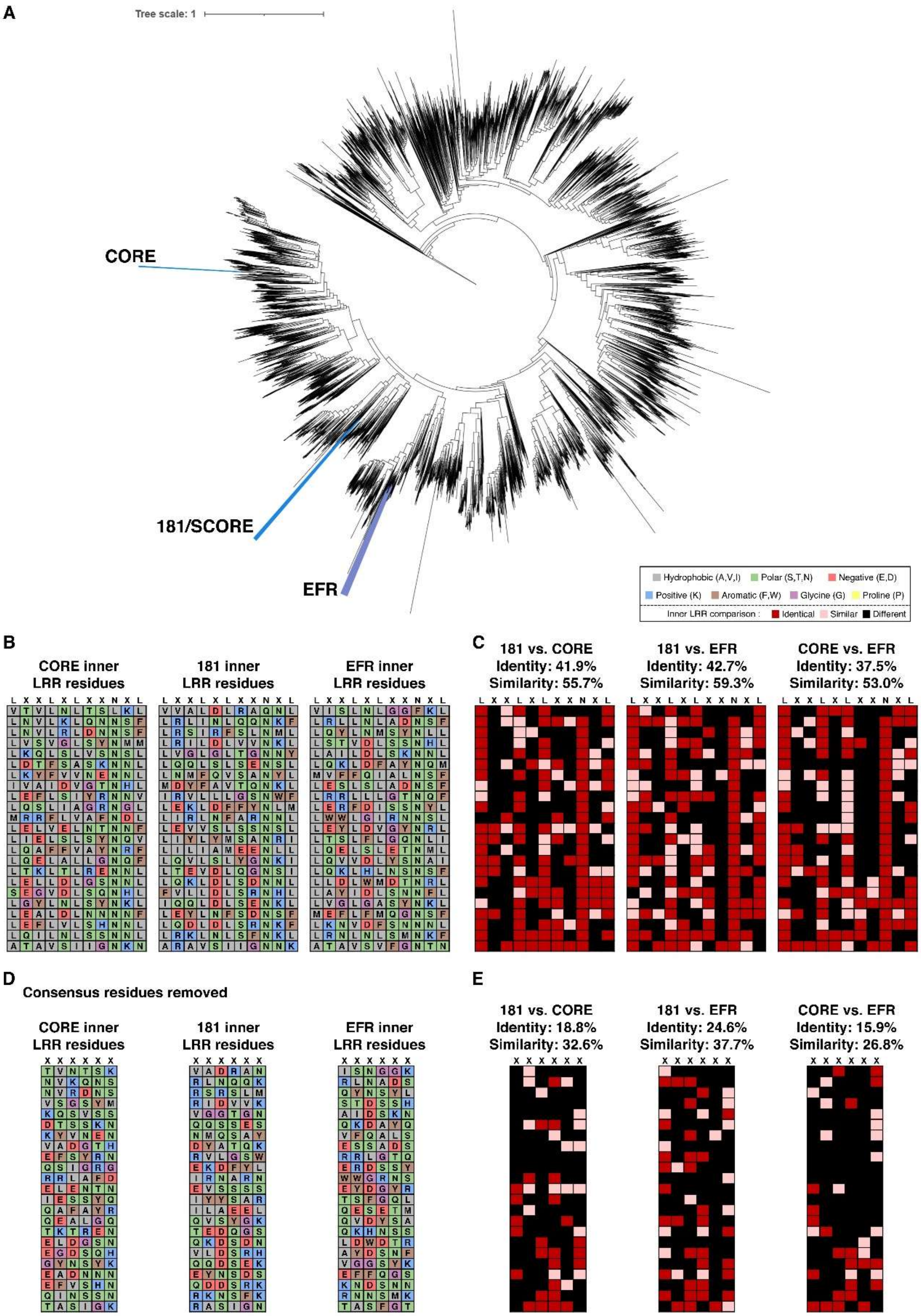
A
Tree scale: 1
CORE
181/SCORE
EFR
Hydrophobic (A,V,I)
Polar (S,T,N)
Negative (E,D)
Positive (K)
Aromatic (F,W)
Glycine (G)
Proline (P)
Inner LRR comparison :
Identical
Similar
Different
B
CORE inner LRR residues
181 inner LRR residues
EFR inner LRR residues
C
181 vs. CORE
Identity: 41.9%
Similarity: 55.7%
181 vs. EFR
Identity: 42.7%
Similarity: 59.3%
CORE vs. EFR
Identity: 37.5%
Similarity: 53.0%
D
Consensus residues removed
CORE inner LRR residues
181 inner LRR residues
EFR inner LRR residues
E
181 vs. CORE
Identity: 18.8%
Similarity: 32.6%
181 vs. EFR
Identity: 24.6%
Similarity: 37.7%
CORE vs. EFR
Identity: 15.9%
Similarity: 26.8%

**Fig. S10. Sequence similarity between EFR$^{Ecto}$, CORE$^{Ecto}$, and 181$^{Ecto}$.**

**(A)** Phylogenetic tree of LRR-RLK-XII ectodomains, with EFR-, 181-. And CORE-clades are highlighted and labelled. **(B)** Inner LRR residues of CORE, 181, and EFR are displayed. The consensus sequence 'LxxLxLxxNxL' is labelled above for reference. **(C)** Comparisons between the inner LRR surface of 181 and CORE, 181 and EFR, and CORE and EFR are shown, highlighing identical residues (dark red), residues with similar properties (light red), and differing residues (black). Identity is calculated as the number of identical residues/number of total residues, while similarity is calculated as the number of identical and similar residues/number of total residues. **(D-E)** The same analyses as in (B-C), but with consensus residues in the LRR motif removed.

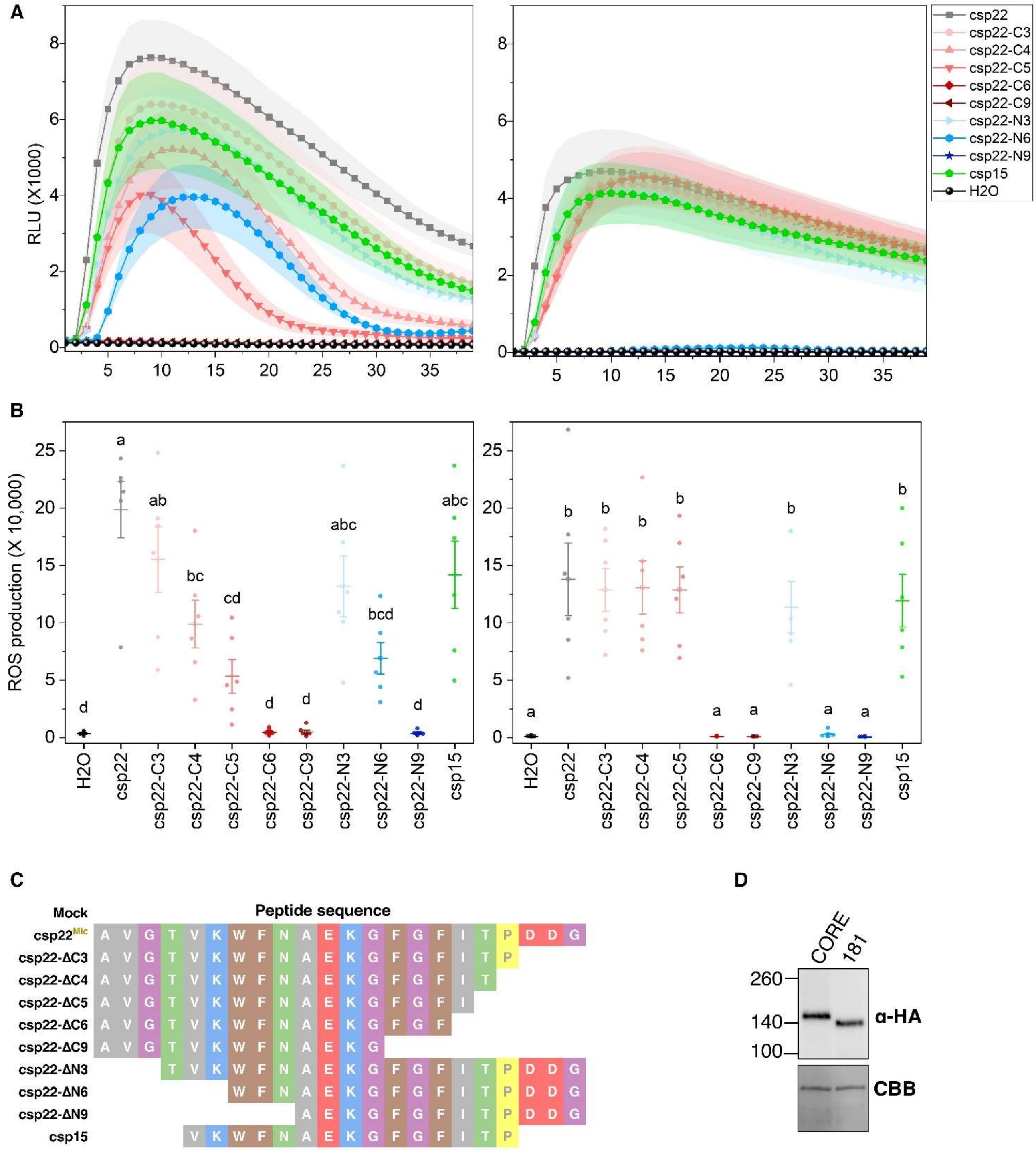


**Fig. S11. Data related to Fig 3H.**

**(A)** ROS production by CORE-EFR (left) and 181-EFR (right) over 50 minutes following treatment with 100 nM of the indicated CSP peptides (listed in the right box). The shaded band represents standard error of mean (S.E.M.). **(B)** Total ROS production in (A). Box plots represent ROS production by CORE-EFR (left) and 181-EFR (right) over 50 minutes, followed by treatment with 100 nM of the indicated CSP peptides. In the plots, centre line, median; upper and lower lines, 25th and 75th percentiles. Data points from 6 technical replicates (n=6) were analyzed with a one-way analysis of variance (ANOVA), followed by post hoc Tukey's honestly significant difference (HSD) test. Data points with different letters indicate significant differences of $P < 0.05$. **(C)** Sequence of peptides used in the ROS assay. **(D)** Samples used in the ROS assays in were analysed with immunoblotting to confirm receptor expression. CBB staining is included as loading controls.

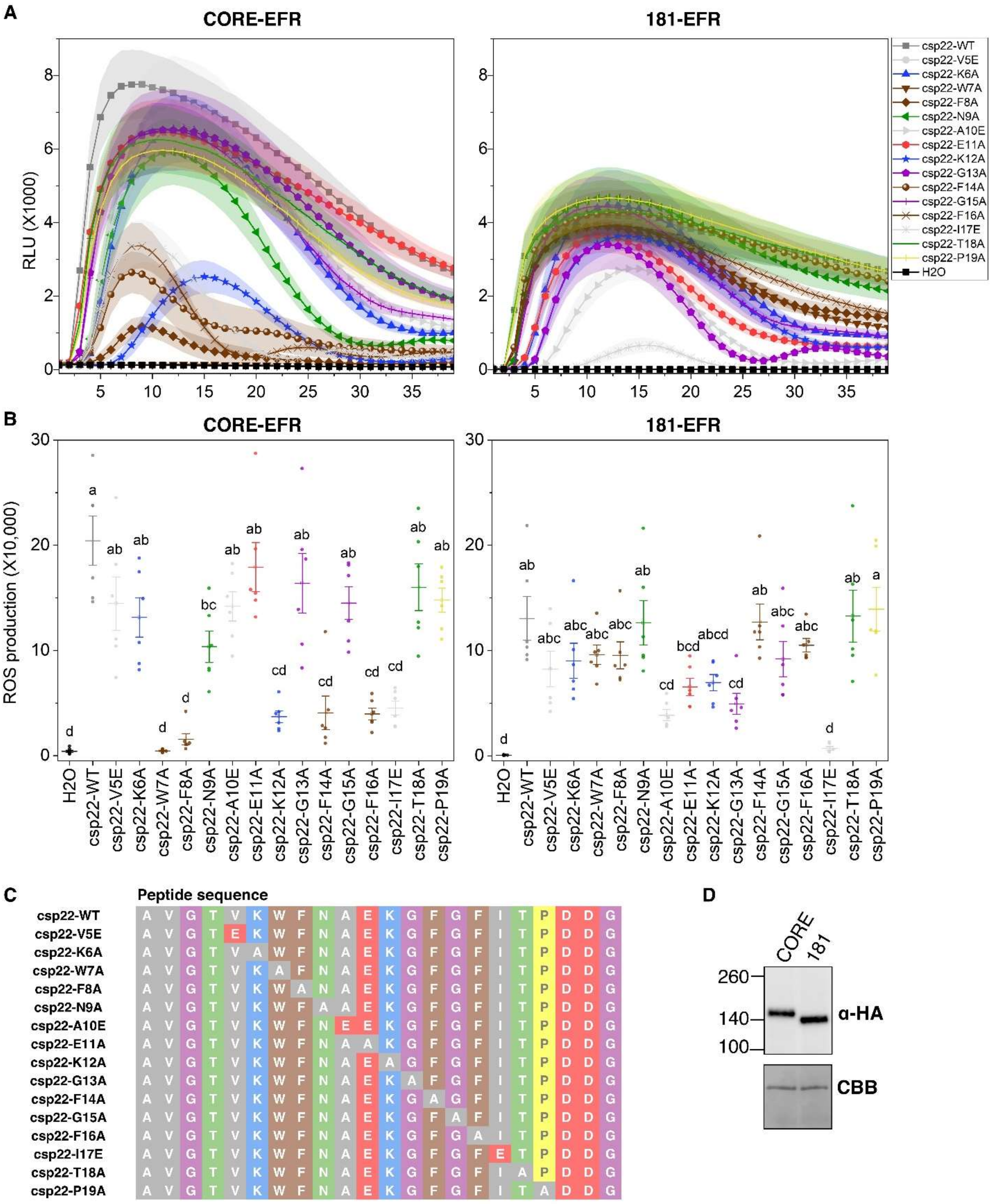


**Fig. S12. Data related to Fig 3I.**

**(A)** ROS production by CORE-EFR (left) and 181-EFR (right) over 50 minutes following treatment with 100 nM of the indicated CSP peptides (listed in the right box). The shaded band represents standard error of mean (S.E.M.). **(B)** Total ROS production in (A). Box plots represent

ROS production by CORE-EFR (left) and 181-EFR (right) over 50 minutes, followed by treatment with 100 nM of the indicated CSP peptides. In the plots, centre line, median; upper and lower lines, 25th and 75th percentiles. Data points from 6 technical replicates (n=6) were analyzed with a one-way analysis of variance (ANOVA), followed by post hoc Tukey's honestly significant difference (HSD) test. Data points with different letters indicate significant differences of $P < 0.05$. **(C)** Sequence of peptides used in the ROS assay. **(D)** Samples used in the ROS assays in (B) were analysed with immunoblotting to confirm receptor expression. CBB staining is included as loading controls.

**A**

| Class | Subclass | Pos1 | Pos2 | Pos3 | Pos4 | Pos5 | Pos6 | Pos7 | Pos8 | Pos9 | Pos10 | Pos11 | Pos12 | Pos13 | Pos14 | Pos15 | Virus | Archaea | Bacteria | Protozoa | Plants | Fungi | Animals | Total |
|---|---|---|---|---|---|---|---|---|---|---|---|---|---|---|---|---|---|---|---|---|---|---|---|---|
| A | 1 | V | K | W | F | N | A | A | K | G | F | G | F | I | E | Q | 1 | 1 | 415 | 2 | 1 | 0 | 17 | 437 |
| B | 1 | V | K | W | F | N | A | T | K | G | F | G | F | I | A | P | 0 | 1 | 309 | 1 | 0 | 1 | 6 | 318 |
| B | 1 | V | K | F | F | N | A | T | K | G | F | G | F | I | T | P | 1 | 1 | 633 | 5 | 1 | 1 | 17 | 659 |
| B | 1 | V | K | W | F | N | A | T | K | G | F | G | F | I | Q | P | 0 | 1 | 1701 | 9 | 6 | 1 | 19 | 1737 |
| B | 1 | V | K | W | F | N | A | T | K | G | Y | G | F | I | Q | P | 0 | 3 | 819 | 5 | 3 | 1 | 12 | 843 |
| B | 2 | V | K | F | F | N | V | T | K | G | F | G | F | I | I | P | 0 | 1 | 65 | 0 | 0 | 0 | 0 | 66 |
| C | 1 | V | K | F | F | N | A | D | K | G | Y | G | F | I | A | P | 0 | 1 | 392 | 2 | 2 | 1 | 3 | 401 |
| C | 1 | V | K | W | F | N | A | D | K | G | F | G | F | I | T | V | 0 | 2 | 380 | 0 | 2 | 2 | 2 | 388 |
| C | 1 | V | K | W | F | N | A | D | K | G | F | G | F | I | T | P | 1 | 2 | 2546 | 7 | 14 | 6 | 18 | 2594 |
| C | 1 | V | K | W | F | N | A | D | K | G | F | G | F | I | T | G | 0 | 2 | 258 | 1 | 0 | 0 | 3 | 264 |
| C | 1 | V | K | W | F | N | A | D | K | G | F | G | F | I | E | V | 0 | 1 | 628 | 2 | 2 | 2 | 6 | 641 |
| C | 2 | V | K | W | F | N | A | E | K | G | Y | G | F | I | A | V | 0 | 0 | 2497 | 6 | 3 | 5 | 4 | 2515 |
| C | 2 | V | K | W | F | N | A | E | K | G | F | G | F | I | A | Q | 0 | 13 | 3152 | 6 | 3 | 10 | 14 | 3198 |
| C | 2 | V | K | W | F | N | A | E | K | G | F | G | F | I | A | P | 0 | 3 | 2449 | 7 | 14 | 8 | 20 | 2501 |
| C | 2 | V | K | W | F | N | A | E | K | G | F | G | F | I | S | R | 0 | 0 | 277 | 0 | 2 | 0 | 4 | 283 |
| C | 2 | V | K | W | F | N | A | E | K | G | F | G | F | I | T | P | 0 | 2 | 788 | 0 | 3 | 0 | 8 | 801 |
| C | 2 | V | K | W | F | N | A | E | K | G | F | G | F | I | E | V | 1 | 3 | 2232 | 5 | 18 | 6 | 15 | 2280 |
| C | 2 | V | K | W | F | N | A | E | K | G | F | G | F | I | E | Q | 0 | 11 | 3177 | 6 | 7 | 3 | 80 | 3284 |
| C | 2 | V | K | W | F | N | A | E | K | G | F | G | F | I | E | R | 0 | 3 | 2626 | 4 | 16 | 3 | 11 | 2663 |
| D | 1 | V | K | W | F | N | N | A | K | G | F | G | F | I | V | E | 0 | 0 | 336 | 1 | 1 | 1 | 14 | 353 |
| D | 1 | V | K | W | F | N | N | A | K | G | Y | G | F | I | N | E | 0 | 3 | 518 | 2 | 1 | 5 | 18 | 547 |
| D | 1 | V | K | W | F | N | N | A | K | G | F | G | F | I | C | S | 0 | 0 | 274 | 0 | 1 | 12 | 9 | 296 |
| D | 1 | V | K | W | F | N | N | A | K | G | F | G | F | I | C | P | 2 | 5 | 2883 | 8 | 8 | 49 | 19 | 2974 |
| D | 1 | V | K | W | F | N | N | A | K | G | Y | G | F | I | I | E | 0 | 0 | 197 | 1 | 1 | 2 | 9 | 210 |
| D | 1 | V | K | W | F | N | N | A | K | G | Y | G | F | I | L | A | 0 | 0 | 339 | 1 | 1 | 1 | 10 | 352 |
| E | 1 | V | K | W | F | N | S | T | K | G | F | G | F | I | Q | P | 0 | 3 | 1858 | 9 | 4 | 3 | 21 | 1898 |
| E | 2 | V | K | W | F | N | T | S | K | G | F | G | F | I | S | R | 0 | 0 | 1755 | 0 | 1 | 0 | 6 | 1762 |
| E | 3 | V | K | W | F | N | T | T | K | G | Y | G | F | I | A | P | 0 | 2 | 1260 | 5 | 6 | 2 | 22 | 1297 |
| E | 3 | V | K | W | F | N | T | T | K | G | F | G | F | I | A | P | 0 | 0 | 485 | 1 | 0 | 0 | 15 | 501 |
| F | 1 | V | K | W | F | N | S | D | K | G | F | G | F | I | T | G | 0 | 0 | 137 | 1 | 1 | 0 | 1 | 140 |
| F | 1 | V | K | W | F | N | S | D | K | G | F | G | F | I | E | Q | 0 | 0 | 242 | 1 | 5 | 0 | 3 | 251 |
| F | 2 | V | K | W | F | N | S | E | K | G | F | G | F | L | S | R | 0 | 2 | 1451 | 5 | 11 | 6 | 12 | 1487 |
| F | 2 | V | K | W | F | N | S | E | K | G | F | G | F | I | E | V | 0 | 3 | 1999 | 6 | 19 | 7 | 23 | 2057 |
| F | 2 | V | K | W | F | N | S | E | K | G | F | G | F | I | E | Q | 2 | 7 | 2502 | 7 | 12 | 2 | 61 | 2593 |
| F | 3 | V | K | F | F | N | T | D | K | G | F | G | F | I | K | P | 0 | 0 | 615 | 2 | 1 | 0 | 5 | 623 |
| F | 4 | V | K | W | F | N | N | E | K | G | F | G | F | I | S | V | 0 | 0 | 155 | 1 | 1 | 0 | 1 | 158 |
| F | 4 | V | K | W | F | N | N | E | K | G | F | G | F | I | T | V | 0 | 0 | 202 | 0 | 1 | 0 | 2 | 205 |
| F | 4 | V | K | W | F | N | N | E | K | G | F | G | F | I | E | V | 0 | 3 | 1159 | 4 | 8 | 1 | 5 | 1180 |
| F | 4 | V | K | W | F | N | N | E | K | G | Y | G | F | I | E | V | 0 | 0 | 217 | 0 | 1 | 0 | 3 | 221 |
| F | 4 | V | K | W | F | N | N | E | K | G | F | G | F | I | E | I | 0 | 0 | 181 | 0 | 1 | 0 | 1 | 183 |
| F | 4 | V | K | W | F | N | N | E | K | G | F | G | F | I | E | M | 0 | 0 | 281 | 0 | 1 | 0 | 4 | 286 |
| F | 5 | V | K | F | F | N | Q | D | K | G | F | G | F | I | T | P | 0 | 0 | 592 | 3 | 5 | 0 | 27 | 627 |
| F | 5 | I | T | T | W | F | Q | D | K | G | F | G | F | I | K | D | 0 | 0 | 92 | 1 | 0 | 0 | 1 | 94 |
| G | 1 | V | K | W | F | N | P | T | K | G | Y | G | F | I | Q | P | 0 | 0 | 667 | 6 | 2 | 5 | 14 | 694 |
| G | 1 | V | K | W | F | N | P | T | K | G | F | G | F | I | Q | P | 0 | 0 | 121 | 0 | 0 | 0 | 3 | 124 |
| G | 1 | V | K | W | F | N | P | T | K | G | Y | G | F | I | K | P | 0 | 0 | 406 | 3 | 2 | 0 | 1 | 412 |
| H | 1 | V | K | W | F | N | G | Q | K | G | F | G | F | I | Q | P | 0 | 0 | 470 | 0 | 1 | 1 | 20 | 492 |
| H | 1 | V | K | W | F | N | G | Q | K | G | Y | G | F | I | Q | P | 0 | 0 | 107 | 0 | 0 | 0 | 1 | 108 |
| H | 1 | V | K | F | Y | N | G | Q | K | G | F | G | F | I | Q | P | 0 | 0 | 101 | 0 | 1 | 0 | 0 | 102 |
| H | 1 | V | K | W | F | N | G | Q | K | G | F | G | F | I | E | P | 0 | 0 | 57 | 0 | 0 | 0 | 4 | 61 |
| I | 1 | V | K | W | F | N | G | E | K | G | F | G | F | I | A | Q | 0 | 0 | 209 | 0 | 0 | 1 | 3 | 213 |
| I | 1 | V | K | W | F | N | G | E | K | G | F | G | F | I | E | V | 0 | 0 | 247 | 1 | 1 | 0 | 1 | 250 |
| I | 1 | V | K | W | F | N | G | E | K | G | F | G | F | I | E | Q | 0 | 0 | 274 | 0 | 3 | 0 | 2 | 279 |
| I | 1 | V | K | W | F | N | G | E | K | G | F | G | F | I | E | R | 0 | 1 | 186 | 0 | 0 | 0 | 0 | 187 |
| J | 1 | V | K | W | F | N | D | A | K | G | F | G | F | I | S | P | 0 | 0 | 518 | 0 | 9 | 2 | 4 | 533 |
| J | 1 | V | K | W | F | N | D | A | K | G | F | G | F | I | S | R | 0 | 2 | 756 | 4 | 7 | 9 | 24 | 802 |
| J | 1 | V | K | W | F | N | D | A | K | G | F | G | F | I | T | S | 0 | 3 | 345 | 0 | 1 | 1 | 2 | 352 |
| J | 1 | V | K | W | F | N | D | A | K | G | F | G | F | I | T | Q | 0 | 1 | 336 | 2 | 0 | 0 | 2 | 341 |
| J | 1 | V | K | W | F | N | D | A | K | G | F | G | F | I | T | P | 2 | 3 | 3072 | 14 | 17 | 6 | 38 | 3152 |
| J | 1 | V | K | W | F | N | D | A | K | G | Y | G | F | I | Q | R | 0 | 0 | 1427 | 2 | 6 | 0 | 9 | 1444 |
| J | 1 | V | K | W | F | N | D | A | K | G | F | G | F | I | E | Q | 0 | 0 | 156 | 1 | 1 | 1 | 4 | 163 |
| J | 1 | V | K | W | F | N | D | A | K | G | F | G | F | I | K | P | 0 | 0 | 259 | 1 | 0 | 0 | 5 | 265 |
| J | 2 | V | K | W | F | N | E | A | K | G | F | G | F | I | A | Q | 0 | 0 | 148 | 1 | 2 | 0 | 1 | 152 |
| J | 2 | V | K | F | F | N | E | A | K | G | F | G | F | I | E | Q | 0 | 1 | 536 | 2 | 2 | 0 | 2 | 543 |
| J | 2 | V | K | W | F | N | E | A | K | G | F | G | F | I | E | Q | 1 | 0 | 493 | 0 | 2 | 2 | 0 | 498 |
| K | 1 | V | K | W | F | N | D | S | K | G | F | G | F | I | T | P | 1 | 3 | 1869 | 5 | 18 | 3 | 18 | 1917 |
| K | 1 | V | K | F | F | N | D | S | K | G | F | G | F | I | T | E | 0 | 1 | 207 | 2 | 0 | 0 | 2 | 212 |
| K | 1 | V | K | F | F | N | D | S | K | G | Y | G | F | I | T | E | 0 | 0 | 128 | 2 | 0 | 0 | 1 | 131 |
| K | 1 | V | K | W | F | N | D | S | K | G | F | G | F | I | E | Q | 0 | 0 | 217 | 1 | 0 | 0 | 15 | 233 |
| K | 1 | V | K | W | F | N | D | S | K | G | F | G | F | L | E | Q | 0 | 0 | 80 | 1 | 0 | 0 | 5 | 86 |
| K | 2 | V | D | F | F | N | D | T | G | G | Y | G | F | I | S | T | 0 | 355 | 9 | 0 | 0 | 0 | 0 | 364 |
| K | 2 | V | K | F | F | N | D | T | K | G | F | G | F | I | T | E | 0 | 0 | 207 | 4 | 0 | 1 | 2 | 214 |
| K | 2 | V | K | W | F | N | D | T | K | G | F | G | F | I | Q | R | 0 | 0 | 437 | 3 | 3 | 1 | 8 | 452 |
| K | 2 | V | D | F | F | N | D | T | G | G | Y | G | F | I | E | T | 0 | 345 | 16 | 0 | 0 | 1 | 1 | 363 |
| K | 2 | V | D | F | F | N | D | T | G | G | Y | G | F | I | D | T | 0 | 193 | 3 | 0 | 0 | 0 | 0 | 196 |
| K | 3 | V | K | W | F | N | E | S | K | G | F | G | F | I | A | Q | 0 | 0 | 223 | 1 | 0 | 0 | 3 | 227 |
| K | 3 | V | K | W | F | N | E | S | K | G | F | G | F | I | S | Q | 0 | 0 | 84 | 0 | 1 | 0 | 2 | 87 |
| K | 3 | V | K | F | F | N | E | S | K | G | Y | G | F | I | T | N | 0 | 0 | 265 | 1 | 0 | 0 | 2 | 268 |
| K | 3 | V | K | W | F | N | E | S | K | G | F | G | F | I | T | P | 1 | 9 | 4795 | 7 | 29 | 11 | 42 | 4894 |
| K | 3 | V | K | F | F | N | E | S | K | G | F | G | F | I | T | E | 0 | 1 | 189 | 0 | 0 | 1 | 1 | 192 |
| K | 3 | V | K | F | F | N | E | S | K | G | Y | G | F | I | T | D | 0 | 3 | 332 | 2 | 0 | 3 | 2 | 342 |
| K | 3 | V | K | W | F | N | E | S | K | G | F | G | F | I | E | Q | 0 | 1 | 751 | 2 | 3 | 3 | 2 | 762 |
| K | 3 | V | K | W | F | N | E | S | K | G | F | G | F | I | E |  | 0 | 0 | 75 | 1 | 0 | 0 | 2 | 78 |
| K | 4 | V | K | W | F | N | E | T | K | G | F | G | F | I | S | Q | 0 | 1 | 322 | 2 | 3 | 0 | 6 | 334 |
| K | 4 | V | K | F | F | N | E | T | K | G | F | G | F | I | T | P | 0 | 0 | 316 | 2 | 2 | 0 | 1 | 321 |
| K | 4 | V | K | W | F | N | E | T | K | G | F | G | F | I | Q | Q | 0 | 2 | 928 | 1 | 5 | 2 | 8 | 946 |
| K | 4 | V | K | W | F | N | D | Q | K | G | F | G | F | I | T | P | 0 | 0 | 147 | 2 | 46 | 0 | 2 | 197 |
| L | 1 | V | K | W | F | D | D | G | K | G | F | G | F | I | T | P | 0 | 1 | 87 | 0 | 0 | 1 | 0 | 89 |
| M | 1 | V | K | W | F | N | D | E | K | G | F | G | F | I | T | P | 0 | 3 | 3641 | 7 | 34 | 72 | 43 | 3800 |
| M | 1 | V | K | W | F | N | D | E | K | G | Y | G | F | I | T | P | 0 | 0 | 570 | 1 | 5 | 12 | 15 | 603 |
| M | 1 |  | K | W | F | N | D | E | K | G | Y | G | F | I | T | P | 0 | 0 | 0 | 0 | 0 | 0 | 0 | 0 |
| M | 2 | V | K | W | F | N | E | D | K | G | F | G | F | I | S | P | 0 | 1 | 267 | 5 | 2 | 1 | 12 | 288 |
| M | 2 | I | T | T | W | F | E | D | K | G | F | G | F | I | K | D | 0 | 0 | 221 | 2 | 0 | 0 | 2 | 225 |
| M | 3 | V | K | W | F | N | E | E | K | G | F | G | F | I | S | Q | 0 | 0 | 126 | 0 | 0 | 0 | 2 | 128 |
| M | 3 | V | K | W | F | N | E | E | K | G | F | G | F | I | T | Q | 0 | 0 | 339 | 1 | 3 | 0 | 5 | 348 |
| M | 3 | V | K | W | F | N | E | E | K | G | F | G | F | L | T | Q | 0 | 1 | 120 | 1 | 1 | 0 | 0 | 123 |
| N | 1 | V | K | W | F | N | V | K | A | G | Y | G | F | I | T | R | 0 | 0 | 2 | 1 | 0 | 0 | 52 | 55 |
| N | 1 | V | K | W | F | N | V | K | S | G | Y | G | F | I | N | R | 0 | 0 | 13 | 0 | 0 | 0 | 684 | 697 |
| N | 1 | V | K | W | F | N | V | K | N | G | Y | G | F | I | N | R | 1 | 1 | 77 | 2 | 0 | 0 | 165 | 246 |
| N | 2 | V | K | W | F | N | V | R | N | G | Y | G | F | I | N | R | 0 | 0 | 1 | 0 | 0 | 0 | 91 | 92 |
| O | 1 |  |  | W | F | N | R | T | K | G | F | G | F | L | T | R | 0 | 0 | 0 | 0 | 0 | 0 | 0 | 0 |
| O | 1 | V | K | W | F | N | R | T | K | G | F | G | F | L | T | R | 0 | 2 | 1080 | 5 | 5 | 0 | 39 | 1131 |
| P | 1 | I | T | T | W | F | K | D | K | G | F | G | F | I | K | D | 0 | 0 | 1242 | 8 | 0 | 0 | 6 | 1256 |

**B**

| Class | Subclass | Pos1 | Pos2 | Pos3 | Pos4 | Pos5 | Pos6 | Pos7 | Pos8 | Pos9 | Pos10 | Pos11 | Pos12 | Pos13 | Pos14 | Pos15 | Virus | Archaea | Bacteria | Protozoa | Plants | Fungi | Animals | Total |
|---|---|---|---|---|---|---|---|---|---|---|---|---|---|---|---|---|---|---|---|---|---|---|---|---|
| A | 1 | V | K | W | F | N | A | A | K | G | F | G | F | I | E | Q | 1 | 1 | 424 | 2 | 1 | 0 | 17 | 446 |
| B | 1 | V | K | W | F | N | A | T | K | G | F | G | F | I | A | P | 0 | 1 | 312 | 1 | 0 | 1 | 6 | 321 |
| B | 1 | V | K | F | F | N | A | T | K | G | F | G | F | I | T | P | 1 | 1 | 644 | 5 | 2 | 1 | 17 | 671 |
| B | 1 | V | K | W | F | N | A | T | K | G | F | G | F | I | Q | P | 0 | 1 | 1879 | 9 | 6 | 1 | 25 | 1921 |
| B | 1 | V | K | W | F | N | A | T | K | G | Y | G | F | I | Q | P | 0 | 3 | 850 | 5 | 3 | 1 | 12 | 874 |
| B | 2 | V | K | F | F | N | V | T | K | G | F | G | F | I | I | P | 0 | 1 | 65 | 0 | 0 | 0 | 0 | 66 |
| C | 1 | V | K | F | F | N | A | D | K | G | Y | G | F | I | A | P | 0 | 1 | 397 | 2 | 2 | 1 | 3 | 406 |
| C | 1 | V | K | W | F | N | A | D | K | G | F | G | F | I | T | V | 0 | 2 | 382 | 0 | 2 | 2 | 3 | 391 |
| C | 1 | V | K | W | F | N | A | D | K | G | F | G | F | I | T | P | 1 | 2 | 2714 | 7 | 16 | 6 | 19 | 2765 |
| C | 1 | V | K | W | F | N | A | D | K | G | F | G | F | I | T | G | 0 | 2 | 261 | 1 | 0 | 0 | 3 | 267 |
| C | 1 | V | K | W | F | N | A | D | K | G | F | G | F | I | E | V | 0 | 1 | 642 | 2 | 2 | 2 | 6 | 655 |
| C | 2 | V | K | W | F | N | A | E | K | G | Y | G | F | I | A | V | 0 | 0 | 2524 | 6 | 4 | 5 | 4 | 2543 |
| C | 2 | V | K | W | F | N | A | E | K | G | F | G | F | I | A | Q | 0 | 13 | 3641 | 6 | 3 | 10 | 14 | 3687 |
| C | 2 | V | K | W | F | N | A | E | K | G | F | G | F | I | A | P | 0 | 3 | 2547 | 7 | 14 | 8 | 20 | 2599 |
| C | 2 | V | K | W | F | N | A | E | K | G | F | G | F | I | S | R | 0 | 0 | 277 | 0 | 2 | 0 | 4 | 283 |
| C | 2 | V | K | W | F | N | A | E | K | G | F | G | F | I | T | P | 0 | 2 | 796 | 0 | 3 | 0 | 8 | 809 |
| C | 2 | V | K | W | F | N | A | E | K | G | F | G | F | I | E | V | 1 | 3 | 2416 | 5 | 19 | 6 | 16 | 2466 |
| C | 2 | V | K | W | F | N | A | E | K | G | F | G | F | I | E | Q | 0 | 11 | 3491 | 9 | 7 | 3 | 82 | 3603 |
| C | 2 | V | K | W | F | N | A | E | K | G | F | G | F | I | E | R | 0 | 3 | 2881 | 4 | 17 | 3 | 12 | 2920 |
| D | 1 | V | K | W | F | N | N | A | K | G | F | G | F | I | V | E | 0 | 0 | 338 | 1 | 1 | 1 | 15 | 356 |
| D | 1 | V | K | W | F | N | N | A | K | G | Y | G | F | I | N | E | 0 | 3 | 521 | 2 | 1 | 5 | 18 | 550 |
| D | 1 | V | K | W | F | N | N | A | K | G | F | G | F | I | C | S | 0 | 0 | 275 | 0 | 1 | 13 | 9 | 298 |
| D | 1 | V | K | W | F | N | N | A | K | G | F | G | F | I | C | P | 2 | 5 | 2984 | 9 | 9 | 50 | 22 | 3081 |
| D | 1 | V | K | W | F | N | N | A | K | G | Y | G | F | I | I | E | 0 | 0 | 199 | 1 | 1 | 2 | 9 | 212 |
| D | 1 | V | K | W | F | N | N | A | K | G | Y | G | F | I | L | A | 0 | 0 | 345 | 1 | 1 | 1 | 10 | 358 |
| E | 1 | V | K | W | F | N | S | T | K | G | F | G | F | I | Q | P | 0 | 3 | 2128 | 12 | 5 | 3 | 26 | 2177 |
| E | 2 | V | K | W | F | N | T | S | K | G | F | G | F | I | S | R | 0 | 0 | 1766 | 0 | 1 | 0 | 6 | 1773 |
| E | 3 | V | K | W | F | N | T | T | K | G | Y | G | F | I | A | P | 0 | 2 | 1298 | 6 | 6 | 2 | 23 | 1337 |
| E | 3 | V | K | W | F | N | T | T | K | G | F | G | F | I | A | P | 0 | 0 | 495 | 2 | 0 | 0 | 15 | 512 |
| F | 1 | V | K | W | F | N | S | D | K | G | F | G | F | I | T | G | 0 | 0 | 142 | 1 | 1 | 0 | 1 | 145 |
| F | 1 | V | K | W | F | N | S | D | K | G | F | G | F | I | E | Q | 0 | 0 | 242 | 1 | 5 | 0 | 3 | 251 |
| F | 2 | V | K | W | F | N | S | E | K | G | F | G | F | L | S | R | 0 | 2 | 1479 | 5 | 12 | 6 | 12 | 1516 |
| F | 2 | V | K | W | F | N | S | E | K | G | F | G | F | I | E | V | 0 | 3 | 2210 | 6 | 20 | 8 | 26 | 2273 |
| F | 2 | V | K | W | F | N | S | E | K | G | F | G | F | I | E | Q | 2 | 7 | 2652 | 8 | 12 | 2 | 63 | 2746 |
| F | 3 | V | K | F | F | N | T | D | K | G | F | G | F | I | K | P | 0 | 0 | 623 | 2 | 1 | 0 | 5 | 631 |
| F | 4 | V | K | W | F | N | N | E | K | G | F | G | F | I | S | V | 0 | 0 | 164 | 1 | 1 | 0 | 1 | 167 |
| F | 4 | V | K | W | F | N | N | E | K | G | F | G | F | I | T | V | 0 | 0 | 204 | 0 | 1 | 0 | 2 | 207 |
| F | 4 | V | K | W | F | N | N | E | K | G | F | G | F | I | E | V | 0 | 3 | 1176 | 4 | 9 | 1 | 5 | 1198 |
| F | 4 | V | K | W | F | N | N | E | K | G | Y | G | F | I | E | V | 0 | 0 | 217 | 0 | 1 | 0 | 3 | 221 |
| F | 4 | V | K | W | F | N | N | E | K | G | F | G | F | I | E | I | 0 | 0 | 184 | 0 | 1 | 0 | 1 | 186 |
| F | 4 | V | K | W | F | N | N | E | K | G | F | G | F | I | E | M | 0 | 0 | 281 | 0 | 1 | 0 | 4 | 286 |
| F | 5 | V | K | F | F | N | Q | D | K | G | F | G | F | I | T | P | 0 | 0 | 603 | 3 | 5 | 0 | 27 | 638 |
| F | 5 | I | T | T | W | F | Q | D | K | G | F | G | F | I | K | D | 0 | 0 | 92 | 1 | 0 | 0 | 1 | 94 |
| G | 1 | V | K | W | F | N | P | T | K | G | Y | G | F | I | Q | P | 0 | 0 | 682 | 9 | 3 | 5 | 14 | 713 |
| G | 1 | V | K | W | F | N | P | T | K | G | F | G | F | I | Q | P | 0 | 0 | 122 | 0 | 0 | 0 | 3 | 125 |
| G | 1 | V | K | W | F | N | P | T | K | G | Y | G | F | I | K | P | 0 | 0 | 410 | 4 | 2 | 0 | 1 | 417 |
| H | 1 | V | K | W | F | N | G | Q | K | G | F | G | F | I | Q | P | 0 | 0 | 487 | 0 | 1 | 1 | 20 | 509 |
| H | 1 | V | K | W | F | N | G | Q | K | G | Y | G | F | I | Q | P | 0 | 0 | 108 | 0 | 0 | 0 | 1 | 109 |
| H | 1 | V | K | F | Y | N | G | Q | K | G | F | G | F | I | Q | P | 0 | 0 | 101 | 0 | 1 | 0 | 0 | 102 |
| H | 1 | V | K | W | F | N | G | Q | K | G | F | G | F | I | E | P | 0 | 0 | 58 | 0 | 0 | 0 | 4 | 62 |
| I | 1 | V | K | W | F | N | G | E | K | G | F | G | F | I | A | Q | 0 | 0 | 215 | 0 | 0 | 1 | 3 | 219 |
| I | 1 | V | K | W | F | N | G | E | K | G | F | G | F | I | E | V | 0 | 0 | 248 | 1 | 1 | 0 | 1 | 251 |
| I | 1 | V | K | W | F | N | G | E | K | G | F | G | F | I | E | Q | 0 | 0 | 275 | 0 | 3 | 0 | 2 | 280 |
| I | 1 | V | K | W | F | N | G | E | K | G | F | G | F | I | E | R | 0 | 1 | 194 | 0 | 0 | 0 | 0 | 195 |
| J | 1 | V | K | W | F | N | D | A | K | G | F | G | F | I | S | P | 0 | 0 | 524 | 0 | 9 | 2 | 4 | 539 |
| J | 1 | V | K | W | F | N | D | A | K | G | F | G | F | I | S | R | 0 | 2 | 774 | 5 | 7 | 9 | 25 | 822 |
| J | 1 | V | K | W | F | N | D | A | K | G | F | G | F | I | T | S | 0 | 3 | 347 | 0 | 1 | 1 | 2 | 354 |
| J | 1 | V | K | W | F | N | D | A | K | G | F | G | F | I | T | Q | 0 | 1 | 384 | 2 | 0 | 0 | 2 | 389 |
| J | 1 | V | K | W | F | N | D | A | K | G | F | G | F | I | T | P | 2 | 3 | 3310 | 14 | 20 | 6 | 40 | 3395 |
| J | 1 | V | K | W | F | N | D | A | K | G | Y | G | F | I | Q | R | 0 | 0 | 1447 | 2 | 7 | 0 | 10 | 1466 |
| J | 1 | V | K | W | F | N | D | A | K | G | F | G | F | I | E | Q | 0 | 0 | 162 | 1 | 1 | 1 | 4 | 169 |
| J | 1 | V | K | W | F | N | D | A | K | G | F | G | F | I | K | P | 0 | 0 | 260 | 1 | 0 | 0 | 5 | 266 |
| J | 2 | V | K | W | F | N | E | A | K | G | F | G | F | I | A | Q | 0 | 0 | 149 | 1 | 2 | 0 | 1 | 153 |
| J | 2 | V | K | F | F | N | E | A | K | G | F | G | F | I | E | Q | 0 | 1 | 548 | 2 | 2 | 0 | 2 | 555 |
| J | 2 | V | K | W | F | N | E | A | K | G | F | G | F | I | E | Q | 1 | 0 | 512 | 0 | 2 | 2 | 0 | 517 |
| K | 1 | V | K | W | F | N | D | S | K | G | F | G | F | I | T | P | 1 | 3 | 2002 | 5 | 19 | 3 | 22 | 2055 |
| K | 1 | V | K | F | F | N | D | S | K | G | F | G | F | I | T | E | 0 | 1 | 211 | 2 | 0 | 0 | 2 | 216 |
| K | 1 | V | K | F | F | N | D | S | K | G | Y | G | F | I | T | E | 0 | 0 | 131 | 2 | 0 | 0 | 2 | 135 |
| K | 1 | V | K | W | F | N | D | S | K | G | F | G | F | I | E | Q | 0 | 0 | 223 | 1 | 0 | 0 | 15 | 239 |
| K | 1 | V | K | W | F | N | D | S | K | G | F | G | F | L | E | Q | 0 | 0 | 90 | 1 | 0 | 0 | 5 | 96 |
| K | 2 | V | D | F | F | N | D | T | G | G | Y | G | F | I | S | T | 0 | 623 | 9 | 0 | 0 | 0 | 0 | 632 |
| K | 2 | V | K | F | F | N | D | T | K | G | F | G | F | I | T | E | 0 | 0 | 215 | 4 | 0 | 1 | 2 | 222 |
| K | 2 | V | K | W | F | N | D | T | K | G | F | G | F | I | Q | R | 0 | 0 | 440 | 4 | 3 | 1 | 8 | 456 |
| K | 2 | V | D | F | F | N | D | T | G | G | Y | G | F | I | E | T | 0 | 615 | 16 | 0 | 0 | 1 | 1 | 633 |
| K | 2 | V | D | F | F | N | D | T | G | G | Y | G | F | I | D | T | 0 | 258 | 3 | 0 | 0 | 0 | 0 | 261 |
| K | 3 | V | K | W | F | N | E | S | K | G | F | G | F | I | A | Q | 0 | 0 | 223 | 1 | 0 | 0 | 3 | 227 |
| K | 3 | V | K | W | F | N | E | S | K | G | F | G | F | I | S | Q | 0 | 0 | 84 | 0 | 1 | 0 | 2 | 87 |
| K | 3 | V | K | F | F | N | E | S | K | G | Y | G | F | I | T | N | 0 | 0 | 265 | 1 | 0 | 0 | 2 | 268 |
| K | 3 | V | K | W | F | N | E | S | K | G | F | G | F | I | T | P | 1 | 9 | 5603 | 8 | 36 | 12 | 45 | 5714 |
| K | 3 | V | K | F | F | N | E | S | K | G | F | G | F | I | T | E | 0 | 1 | 196 | 0 | 0 | 1 | 1 | 199 |
| K | 3 | V | K | F | F | N | E | S | K | G | Y | G | F | I | T | D | 0 | 3 | 337 | 3 | 0 | 3 | 2 | 348 |
| K | 3 | V | K | W | F | N | E | S | K | G | F | G | F | I | E | Q | 0 | 1 | 758 | 2 | 3 | 3 | 2 | 769 |
| K | 3 | V | K | W | F | N | E | S | K | G | F | G | F | I | E | | 0 | 0 | 75 | 1 | 0 | 0 | 2 | 78 |
| K | 4 | V | K | W | F | N | E | T | K | G | F | G | F | I | S | Q | 0 | 1 | 323 | 2 | 3 | 0 | 6 | 335 |
| K | 4 | V | K | F | F | N | E | T | K | G | F | G | F | I | T | P | 0 | 0 | 321 | 2 | 3 | 0 | 1 | 327 |
| K | 4 | V | K | W | F | N | E | T | K | G | F | G | F | I | Q | Q | 0 | 2 | 953 | 1 | 5 | 2 | 8 | 971 |
| K | 4 | V | K | W | F | N | D | Q | K | G | F | G | F | I | T | P | 0 | 0 | 148 | 2 | 51 | 0 | 2 | 203 |
| L | 1 | V | K | W | F | D | D | G | K | G | F | G | F | I | T | P | 0 | 1 | 87 | 0 | 0 | 1 | 0 | 89 |
| M | 1 | V | K | W | F | N | D | E | K | G | F | G | F | I | T | P | 0 | 3 | 4010 | 7 | 36 | 75 | 50 | 4181 |
| M | 1 | V | K | W | F | N | D | E | K | G | Y | G | F | I | T | P | 0 | 0 | 578 | 1 | 5 | 12 | 15 | 611 |
| M | 1 | | K | W | F | N | D | E | K | G | Y | G | F | I | T | P | 0 | 0 | 0 | 0 | 0 | 0 | 0 | 0 |
| M | 2 | V | K | W | F | N | E | D | K | G | F | G | F | I | S | P | 0 | 1 | 269 | 6 | 2 | 1 | 12 | 291 |
| M | 2 | I | T | T | W | F | E | D | K | G | F | G | F | I | K | D | 0 | 0 | 221 | 2 | 0 | 0 | 2 | 225 |
| M | 3 | V | K | W | F | N | E | E | K | G | F | G | F | I | S | Q | 0 | 0 | 126 | 0 | 0 | 0 | 2 | 128 |
| M | 3 | V | K | W | F | N | E | E | K | G | F | G | F | I | T | Q | 0 | 0 | 384 | 1 | 3 | 0 | 5 | 393 |
| M | 3 | V | K | W | F | N | E | E | K | G | F | G | F | L | T | Q | 0 | 1 | 121 | 1 | 1 | 0 | 0 | 124 |
| N | 1 | V | K | W | F | N | V | K | A | G | Y | G | F | I | T | R | 0 | 0 | 2 | 1 | 0 | 0 | 83 | 86 |
| N | 1 | V | K | W | F | N | V | K | S | G | Y | G | F | I | N | R | 0 | 0 | 13 | 0 | 0 | 0 | 853 | 866 |
| N | 1 | V | K | W | F | N | V | K | N | G | Y | G | F | I | N | R | 1 | 1 | 77 | 2 | 0 | 0 | 195 | 276 |
| N | 2 | V | K | W | F | N | V | R | N | G | Y | G | F | I | N | R | 0 | 0 | 1 | 0 | 0 | 0 | 100 | 101 |
| O | 1 | | | W | F | N | R | T | K | G | F | G | F | L | T | R | 0 | 0 | 0 | 0 | 0 | 0 | 0 | 0 |
| O | 1 | V | K | W | F | N | R | T | K | G | F | G | F | L | T | R | 0 | 2 | 1107 | 5 | 5 | 0 | 40 | 1159 |
| P | 1 | I | T | T | W | F | K | D | K | G | F | G | F | I | K | D | 0 | 0 | 1245 | 12 | 0 | 0 | 8 | 1265 |
| | | | | | | | | | | | | | | | | | 14 | 1616 | 79945 | 259 | 437 | 288 | 2220 | 84779 |

**Fig. S13. Data related to Fig 4A.**

The table provides information on the most abundant csp15 peptides identified across multiple kingdoms. These 103 peptides are also used for the ROS experiments in Fig 3D and 4D. The columns, from left to right, include class and subclass of csp15 peptides, ranked based on the amino acids residues in position 6 and 7 in csp15, amino acids residues from position 1-15 (pos) in csp15, **(A)** the number of unique species in each kingdom containing the corresponding peptide. For **(B)**, number of proteins in each kingdom containing the corresponding peptide.

A

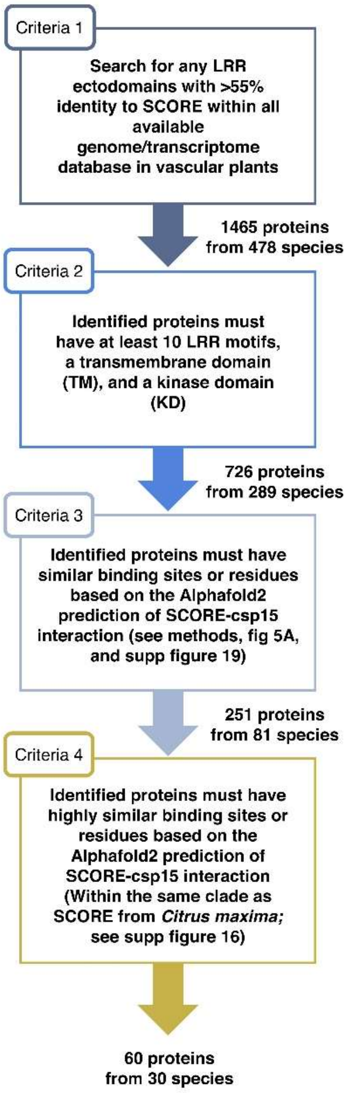


B

| Classification | Criteria 1 | Criteria 2 | Criteria 3 | Criteria 4 |
|---|---|---|---|---|
| Vascular plants | 478 | 289 | 82 | 30 |
| Gymnosperms | 11 | 7 | 0 | 0 |
| Angiosperms | 463 | 281 | 82 | 32 |
| Magnoliidae | 15 | 12 | 5 | 3 |
| Ranunculales | 16 | 10 | 3 | 2 |
| Monocots | 75 | 39 | 3 | 1 |
| Eudicots | 351 | 216 | 70 | 26 |
| Asterids | 127 | 69 | 11 | 0 |
| Malvids | 84 | 65 | 28 | 25 |
| Fabids | 90 | 58 | 26 | 1 |
| Poales | 29 | 16 | 0 | 0 |
| Caryophyllales | 30 | 14 | 0 | 0 |
| Campanulids | 43 | 22 | 2 | 0 |
| Solanales | 12 | 6 | 0 | 0 |
| Lamiales | 21 | 14 | 0 | 0 |
| Brassicales | 7 | 4 | 0 | 0 |
| Myrtales | 20 | 18 | 7 | 7 |
| Sapindales | 38 | 30 | 19 | 16 |
| Fabales | 14 | 6 | 1 | 0 |
| Fagales | 15 | 13 | 8 | 0 |
| Malpighiales | 36 | 21 | 10 | 0 |

C

| Classification | Criteria 1 | Criteria 2 | Criteria 3 | Criteria 4 |
|---|---|---|---|---|
| Vascular plants | 1465 | 726 | 251 | 60 |
| Gymnosperms | 12 | 8 | 0 | 0 |
| Angiosperms | 1446 | 717 | 251 | 60 |
| Magnoliidae | 84 | 44 | 28 | 9 |
| Ranunculales | 102 | 27 | 3 | 3 |
| Monocots | 123 | 55 | 8 | 1 |
| Eudicots | 1121 | 586 | 211 | 47 |
| Asterids | 301 | 129 | 30 | 0 |
| Malvids | 341 | 202 | 65 | 46 |
| Fabids | 368 | 216 | 106 | 1 |
| Poales | 35 | 17 | 0 | 0 |
| Caryophyllales | 68 | 20 | 0 | 0 |
| Campanulids | 104 | 30 | 2 | 0 |
| Solanales | 18 | 7 | 0 | 0 |
| Lamiales | 38 | 18 | 0 | 0 |
| Brassicales | 11 | 6 | 0 | 0 |
| Myrtales | 72 | 40 | 10 | 9 |
| Sapindales | 218 | 130 | 52 | 37 |
| Fabales | 48 | 14 | 3 | 0 |
| Fagales | 112 | 75 | 54 | 0 |
| Malpighiales | 127 | 72 | 42 | 0 |

**Fig. S14. Search for potential SCORE orthologs (data related to Fig 4B).**

**(A)** A detailed outline of the identification of potential SCORE orthologs. Each colored criterion correspond to the color-coded categories in Fig 4B. For additional details, refer to the Methods section. **(B)** The number of unique species containing potential SCORE orthologs meeting each criterion. Criteria 1-4 correspond to those outlined in (A). **(C)** The number of potential SCORE orthologs identified under each criterion.

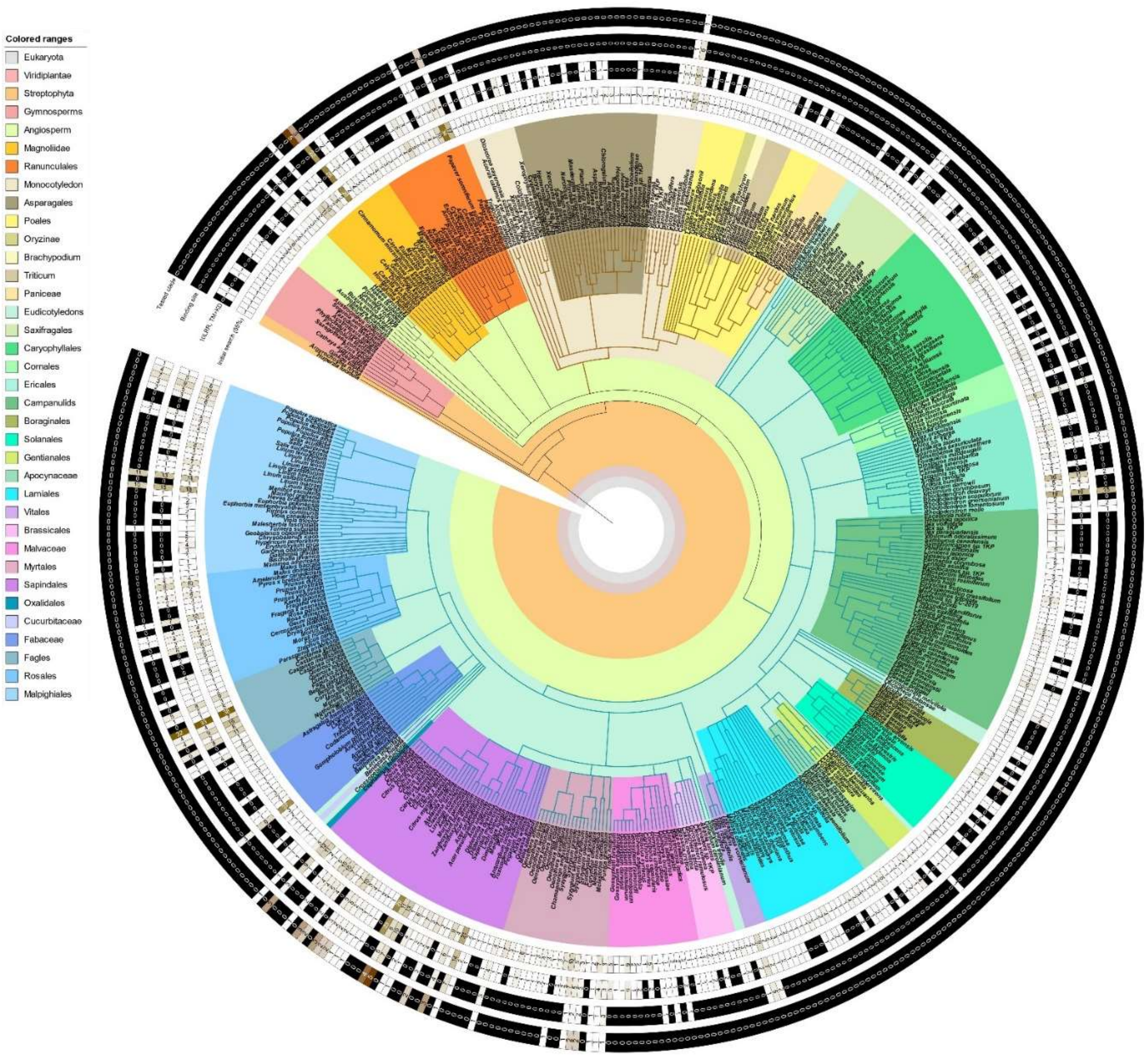


**Fig. S15. Presence of potential SCORE orthologs across angiosperm lineage.**

Phylogenetic tree of 478 plant species with potential SCORE orthologs. Each heatmap indicates the number potential SCORE orthologs identified based on each criterion (1-4; see supplementary Fig 14). Innermost ring: criterion 1; second inner ring: criterion 2; third inner ring: criterion 3; outermost ring: criterion 4. Color coding for the heatmaps: a black box indicates the absence of potential SCORE orthologs, a dark brown box represents a higher number of identified orthologs, and a white or light brown box represents a lower number of identified orthologs.

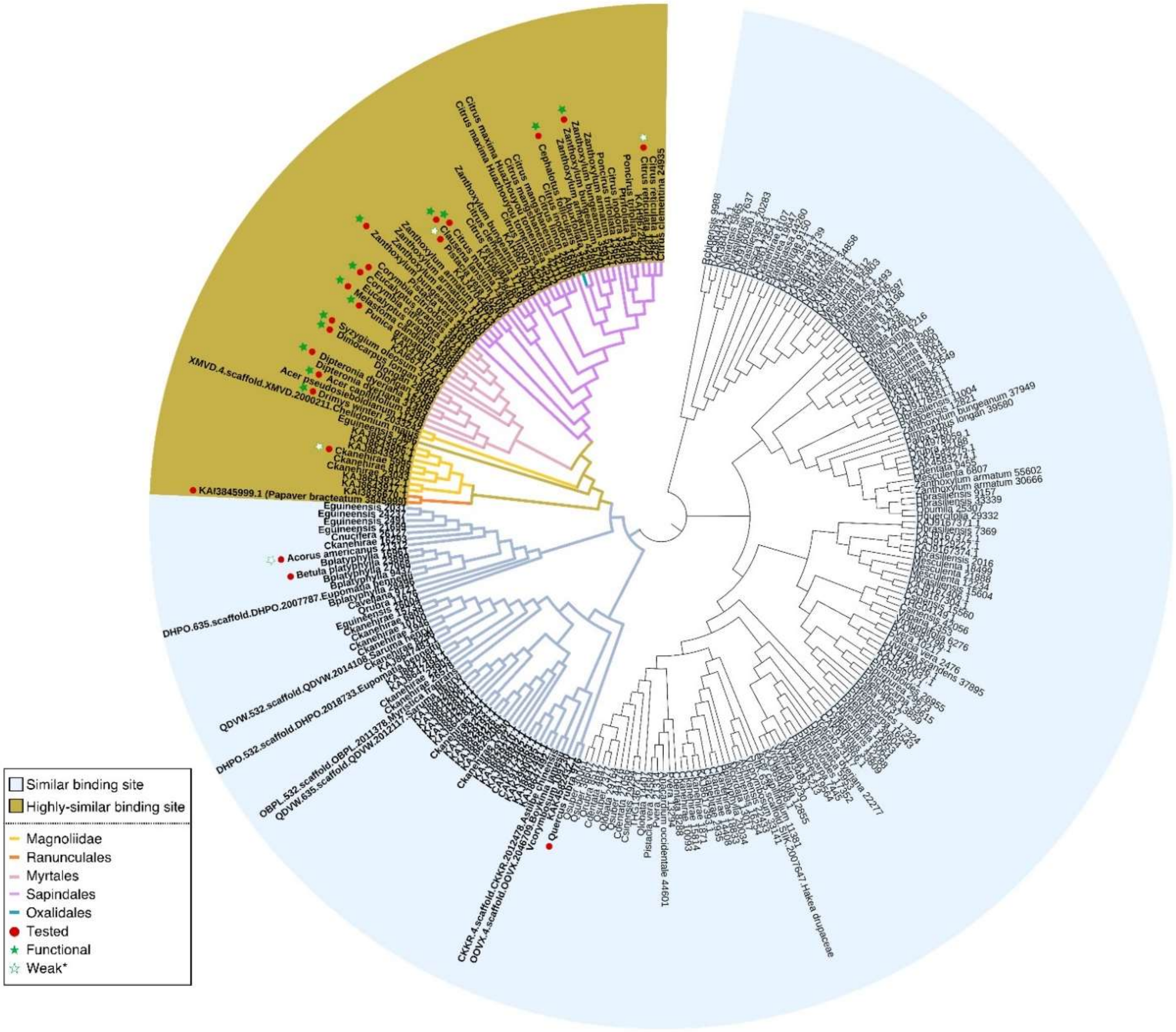


**Fig. S16. Phylogenetic tree of potential SCORE orthologs, related to Fig 4C.**

Phylogenetic tree of all potential SCORE orthologs from vascular plants, identified from criteria 3 (see supplementary Fig 14). SCORE orthologs closely related to Citrus maxima 14337 (the originally identified 181) are highlighted in gold (criterion 4). Branches are labelled according to species order. Red dots mark the 21 tested SCORE orthologs. Green stars indicate functional CSP receptors, while hollow stars represent weak CSP receptors, and unmarked orthologs do not respond to CSPs. The simplified phylogenetic tree in Fig 3C is extracted directly from the clade represented by the light blue branch in this tree.

**A**

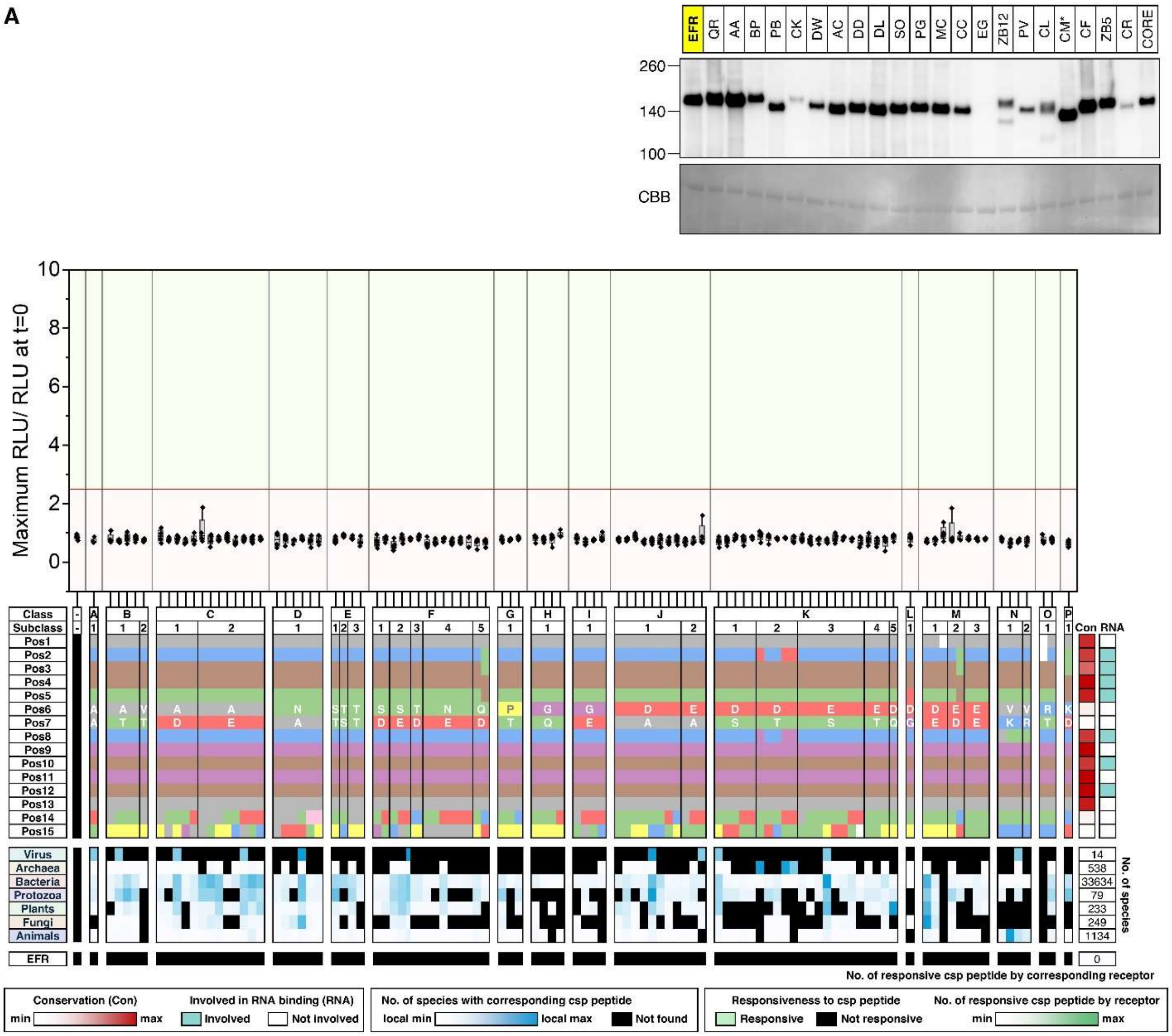
EFR QR AA BP PB CK DW AC DD DL SO PG MC CC EG ZB12 PV CL CM* CF ZB5 CR CORE
260
140
100
CBB
Maximum RLU/ RLU at t=0
10
8
6
4
2
0
Class
Subclass
Pos1
Pos2
Pos3
Pos4
Pos5
Pos6
Pos7
Pos8
Pos9
Pos10
Pos11
Pos12
Pos13
Pos14
Pos15
Con
RNA
Virus
Archaea
Bacteria
Protozoa
Plants
Fungi
Animals
14
538
33634
79
233
249
1134
No. of species
EFR
0
No. of responsive csp peptide by corresponding receptor
Conservation (Con)
min
max
Involved in RNA binding (RNA)
Involved
Not involved
No. of species with corresponding csp peptide
local min
local max
Not found
Responsiveness to csp peptide
Responsive
Not responsive
No. of responsive csp peptide by receptor
min
max

**B**

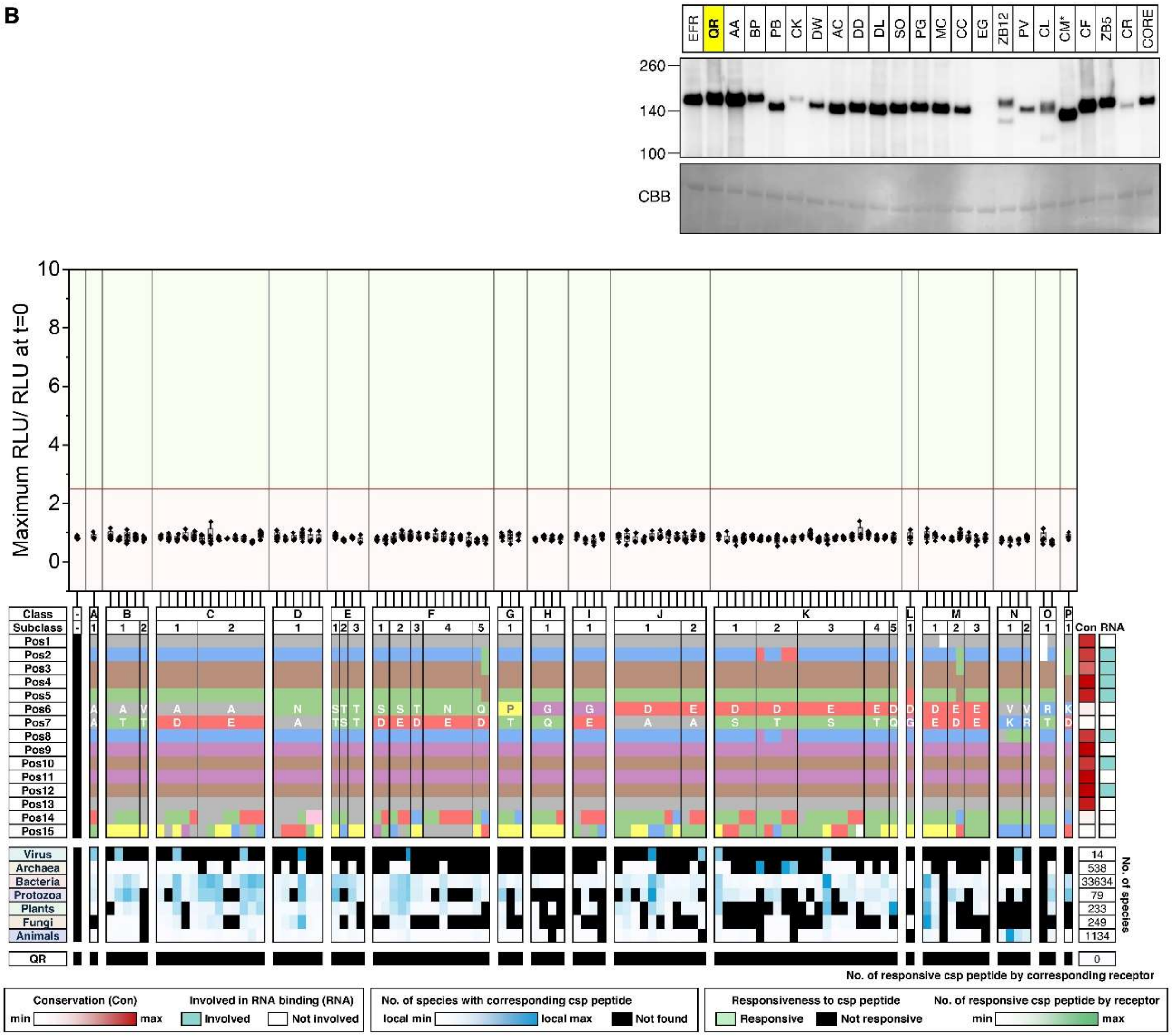

EFR
QR
AA
BP
PB
CK
DW
AC
DD
DL
SO
PG
MC
CC
EG
ZB12
PV
CL
CM*
CF
ZB5
CR
CORE
260
140
100
CBB
Maximum RLU/ RLU at t=0
10
8
6
4
2
0
Class
Subclass
Pos1
Pos2
Pos3
Pos4
Pos5
Pos6
Pos7
Pos8
Pos9
Pos10
Pos11
Pos12
Pos13
Pos14
Pos15
Con
RNA
Virus
Archaea
Bacteria
Protozoa
Plants
Fungi
Animals
14
538
33634
79
233
249
1134
No. of species
QR
0
No. of responsive csp peptide by corresponding receptor
Conservation (Con)
min
max
Involved in RNA binding (RNA)
Involved
Not involved
No. of species with corresponding csp peptide
local min
local max
Not found
Responsiveness to csp peptide
Responsive
Not responsive
No. of responsive csp peptide by receptor
min
max

**C**

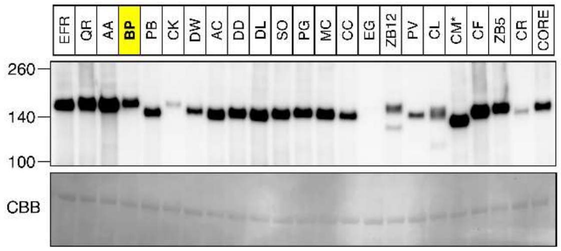
EFR
QR
AA
BP
PB
CK
DW
AC
DD
DL
SO
PG
MC
CC
EG
ZB12
PV
CL
CM*
CF
ZB5
CR
CORE
260
140
100
CBB

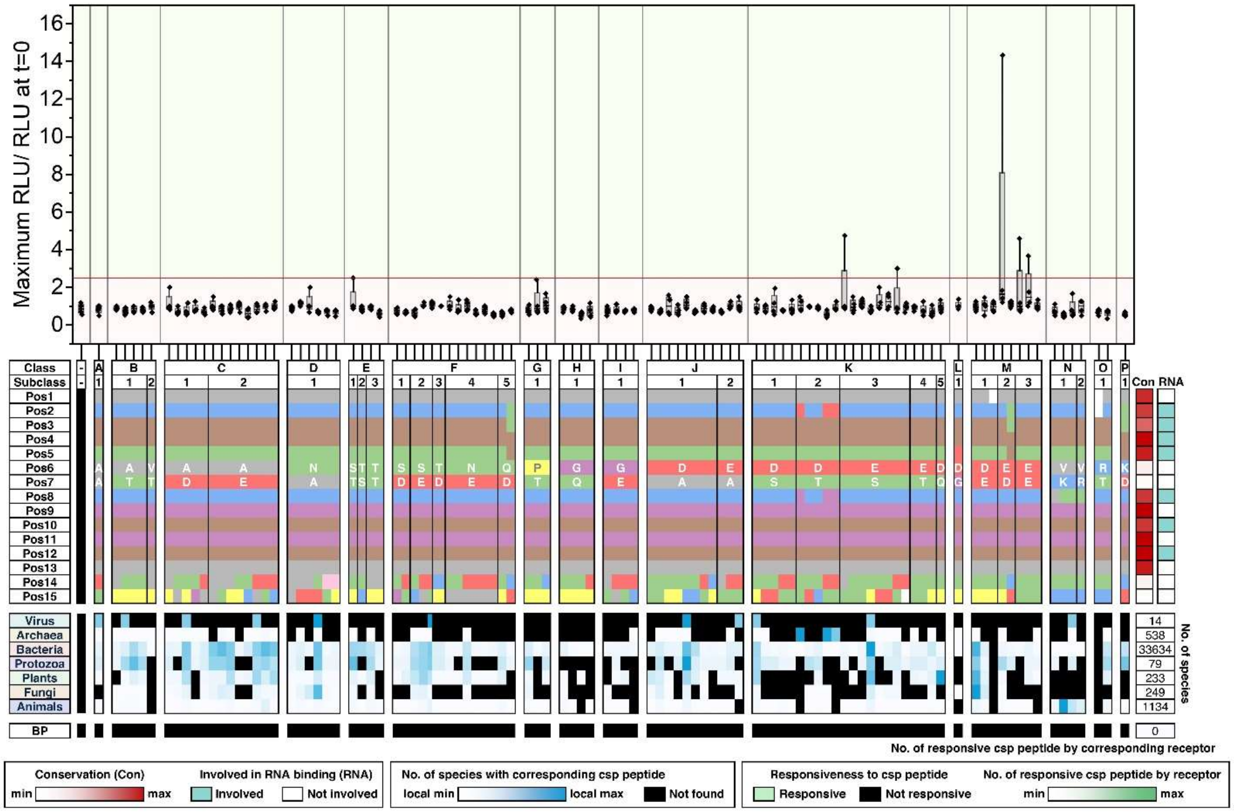
Maximum RLU/ RLU at t=0
16
14
12
10
8
6
4
2
0
Class
Subclass
Pos1
Pos2
Pos3
Pos4
Pos5
Pos6
Pos7
Pos8
Pos9
Pos10
Pos11
Pos12
Pos13
Pos14
Pos15
Con
RNA
Virus
Archaea
Bacteria
Protozoa
Plants
Fungi
Animals
14
538
33634
79
233
249
1134
No. of species
BP
0
No. of responsive csp peptide by corresponding receptor
Conservation (Con)
min
max
Involved in RNA binding (RNA)
Involved
Not involved
No. of species with corresponding csp peptide
local min
local max
Not found
Responsiveness to csp peptide
Responsive
Not responsive
No. of responsive csp peptide by receptor
min
max

D

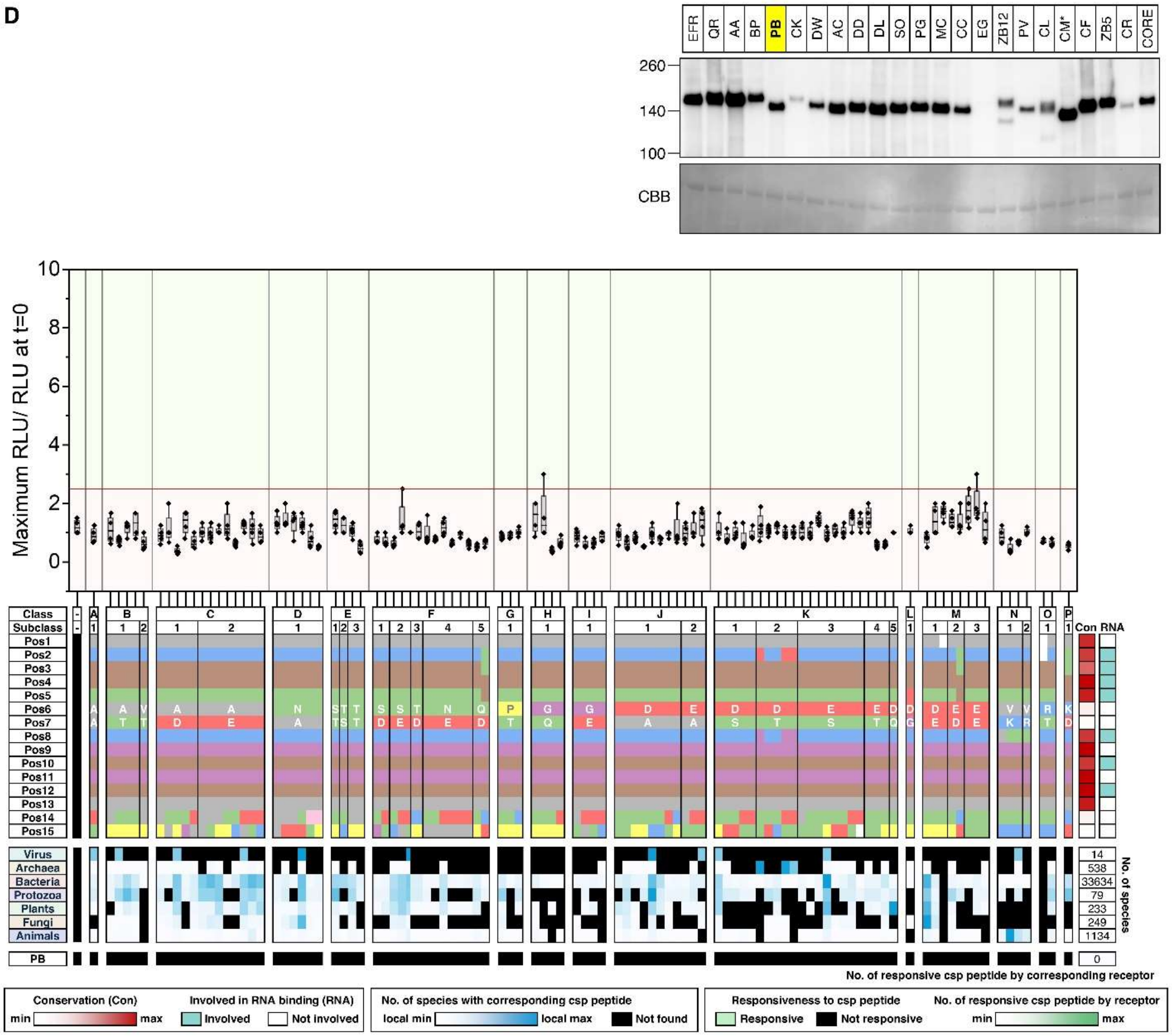

EFR QR AA BP PB CK DW AC DD DL SO PG MC CC EG ZB12 PV CL CM* CF ZB5 CR CORE
260
140
100
CBB
Maximum RLU/ RLU at t=0
Class
Subclass
Pos1
Pos2
Pos3
Pos4
Pos5
Pos6
Pos7
Pos8
Pos9
Pos10
Pos11
Pos12
Pos13
Pos14
Pos15
Con RNA
Virus
Archaea
Bacteria
Protozoa
Plants
Fungi
Animals
14
538
33634
79
233
249
1134
No. of species
PB
0
No. of responsive csp peptide by corresponding receptor
Conservation (Con)
min
max
Involved in RNA binding (RNA)
Involved
Not involved
No. of species with corresponding csp peptide
local min
local max
Not found
Responsiveness to csp peptide
Responsive
Not responsive
No. of responsive csp peptide by receptor
min
max

E

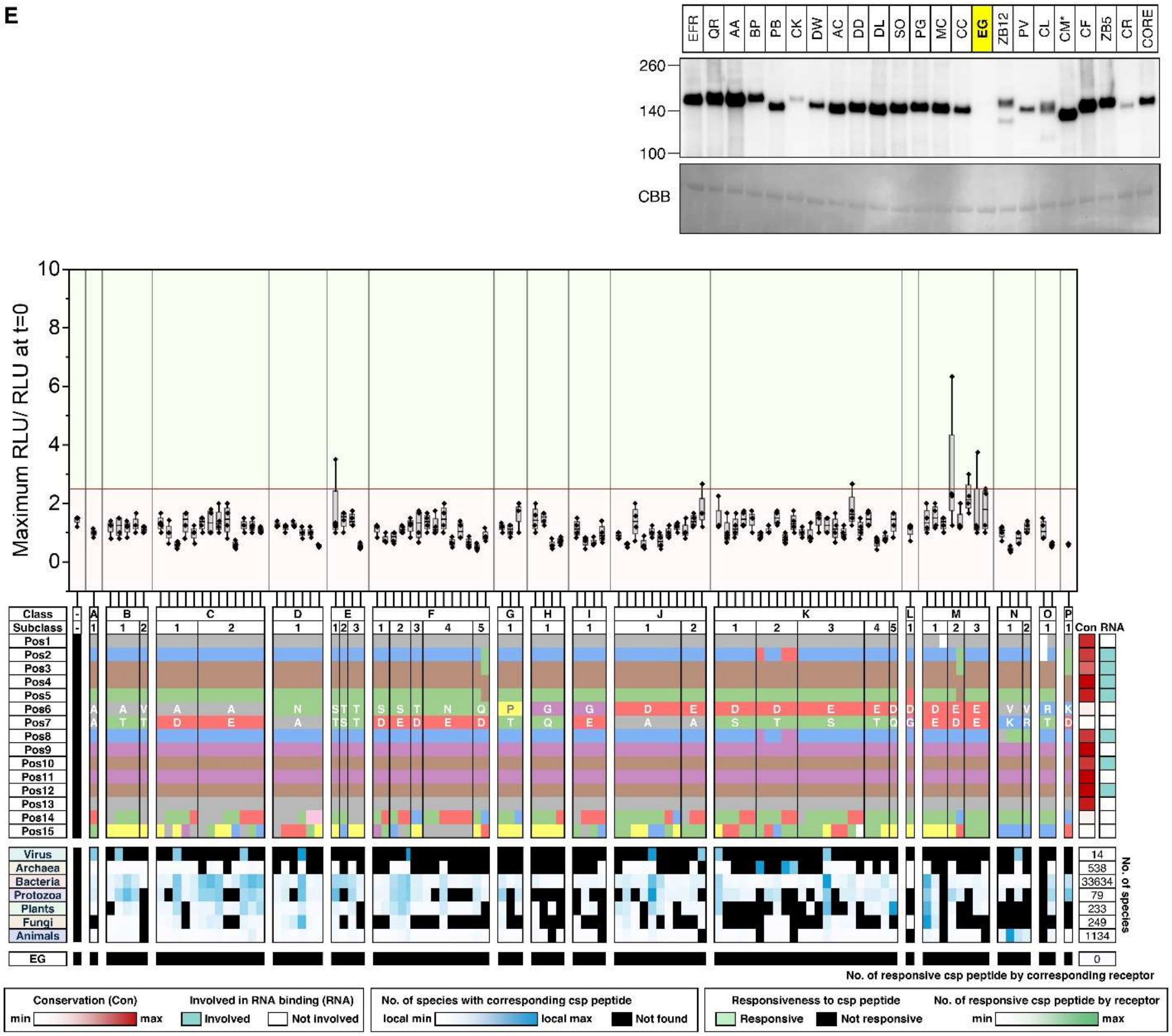
EFR QR AA BP PB CK DW AC DD DL SO PG MC CC EG ZB12 PV CL CM* CF ZB5 CR CORE
260
140
100
CBB
Maximum RLU/ RLU at t=0
Class
Subclass
Pos1
Pos2
Pos3
Pos4
Pos5
Pos6
Pos7
Pos8
Pos9
Pos10
Pos11
Pos12
Pos13
Pos14
Pos15
Con RNA
Virus
Archaea
Bacteria
Protozoa
Plants
Fungi
Animals
14
538
33634
79
233
249
1134
No. of species
EG
0
No. of responsive csp peptide by corresponding receptor
Conservation (Con)
min max
Involved in RNA binding (RNA)
Involved
Not involved
No. of species with corresponding csp peptide
local min local max
Not found
Responsiveness to csp peptide
Responsive
Not responsive
No. of responsive csp peptide by receptor
min max

**F**

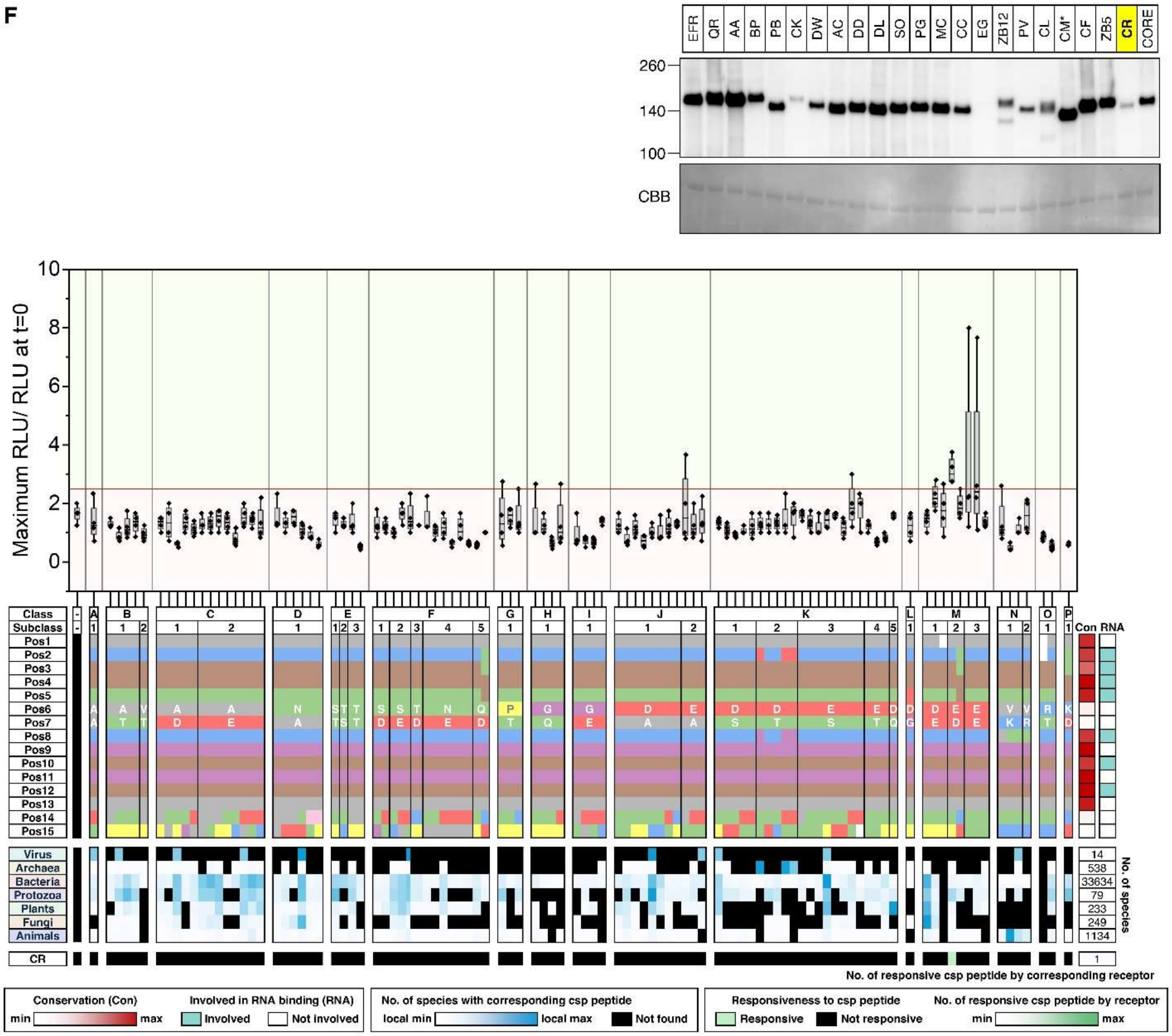
EFR
QR
AA
BP
PB
CK
DW
AC
DD
DL
SO
PG
MC
CC
EG
ZB12
PV
CL
CM*
CF
ZB5
CR
CORE
260
140
100
CBB
Maximum RLU/ RLU at t=0
10
8
6
4
2
0
Class
Subclass
Pos1
Pos2
Pos3
Pos4
Pos5
Pos6
Pos7
Pos8
Pos9
Pos10
Pos11
Pos12
Pos13
Pos14
Pos15
Con
RNA
Virus
Archaea
Bacteria
Protozoa
Plants
Fungi
Animals
CR
14
538
33634
79
233
249
1134
No. of species
1
No. of responsive csp peptide by corresponding receptor
Conservation (Con)
min
max
Involved in RNA binding (RNA)
Involved
Not involved
No. of species with corresponding csp peptide
local min
local max
Not found
Responsiveness to csp peptide
Responsive
Not responsive
No. of responsive csp peptide by receptor
min
max

G

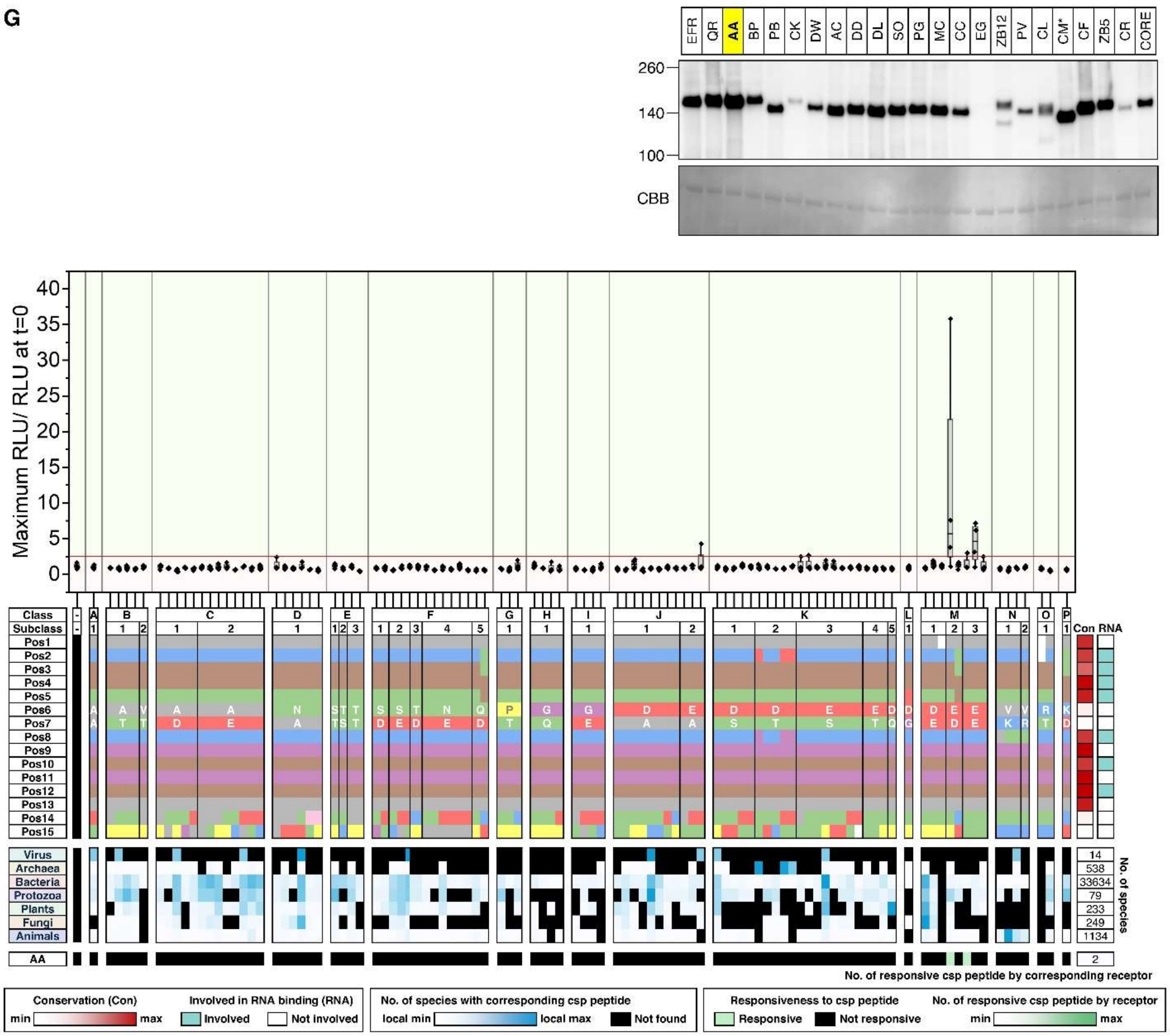

EFR QR AA BP PB CK DW AC DD DL SO PG MC CC EG ZB12 PV CL CM* CF ZB5 CR CORE
260
140
100
CBB
Maximum RLU/ RLU at t=0
Class
Subclass
Virus
Archaea
Bacteria
Protozoa
Plants
Fungi
Animals
AA
No. of species
14
538
33634
79
233
249
1134
2
Con RNA
No. of responsive csp peptide by corresponding receptor
Conservation (Con)
min max
Involved in RNA binding (RNA)
Involved
Not involved
No. of species with corresponding csp peptide
local min local max Not found
Responsiveness to csp peptide
Responsive Not responsive
No. of responsive csp peptide by receptor
min max

H

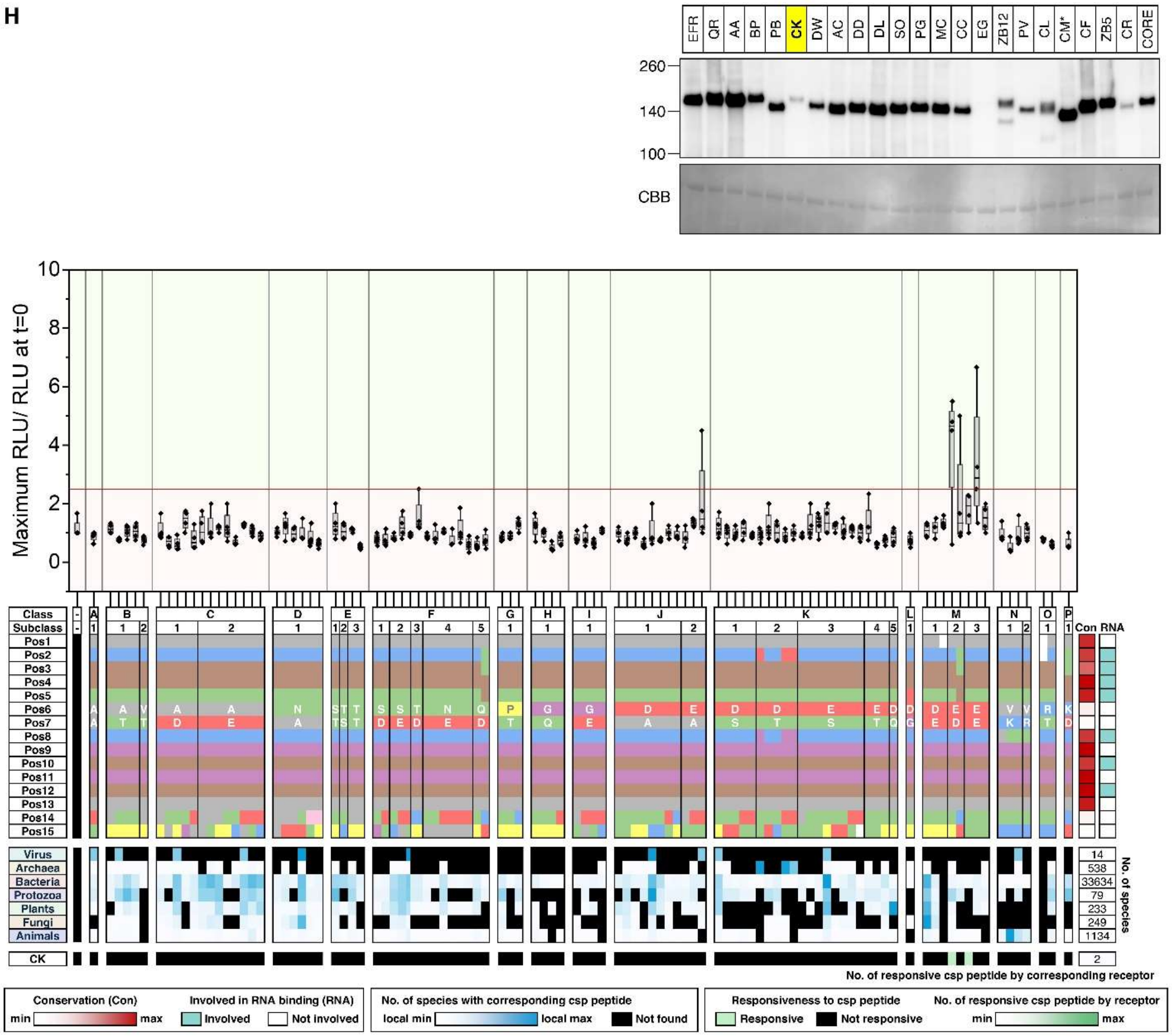

EFR QR AA BP PB CK DW AC DD DL SO PG MC CC EG ZB12 PV CL CM* CF ZB5 CR CORE
260
140
100
CBB
Maximum RLU/ RLU at t=0
Class
Subclass
Pos1
Pos2
Pos3
Pos4
Pos5
Pos6
Pos7
Pos8
Pos9
Pos10
Pos11
Pos12
Pos13
Pos14
Pos15
Con RNA
Virus
Archaea
Bacteria
Protozoa
Plants
Fungi
Animals
14
538
33634
79
233
249
1134
No. of species
CK
2
No. of responsive csp peptide by corresponding receptor
Conservation (Con)
min
max
Involved in RNA binding (RNA)
Involved
Not involved
No. of species with corresponding csp peptide
local min
local max
Not found
Responsiveness to csp peptide
Responsive
Not responsive
No. of responsive csp peptide by receptor
min
max

I

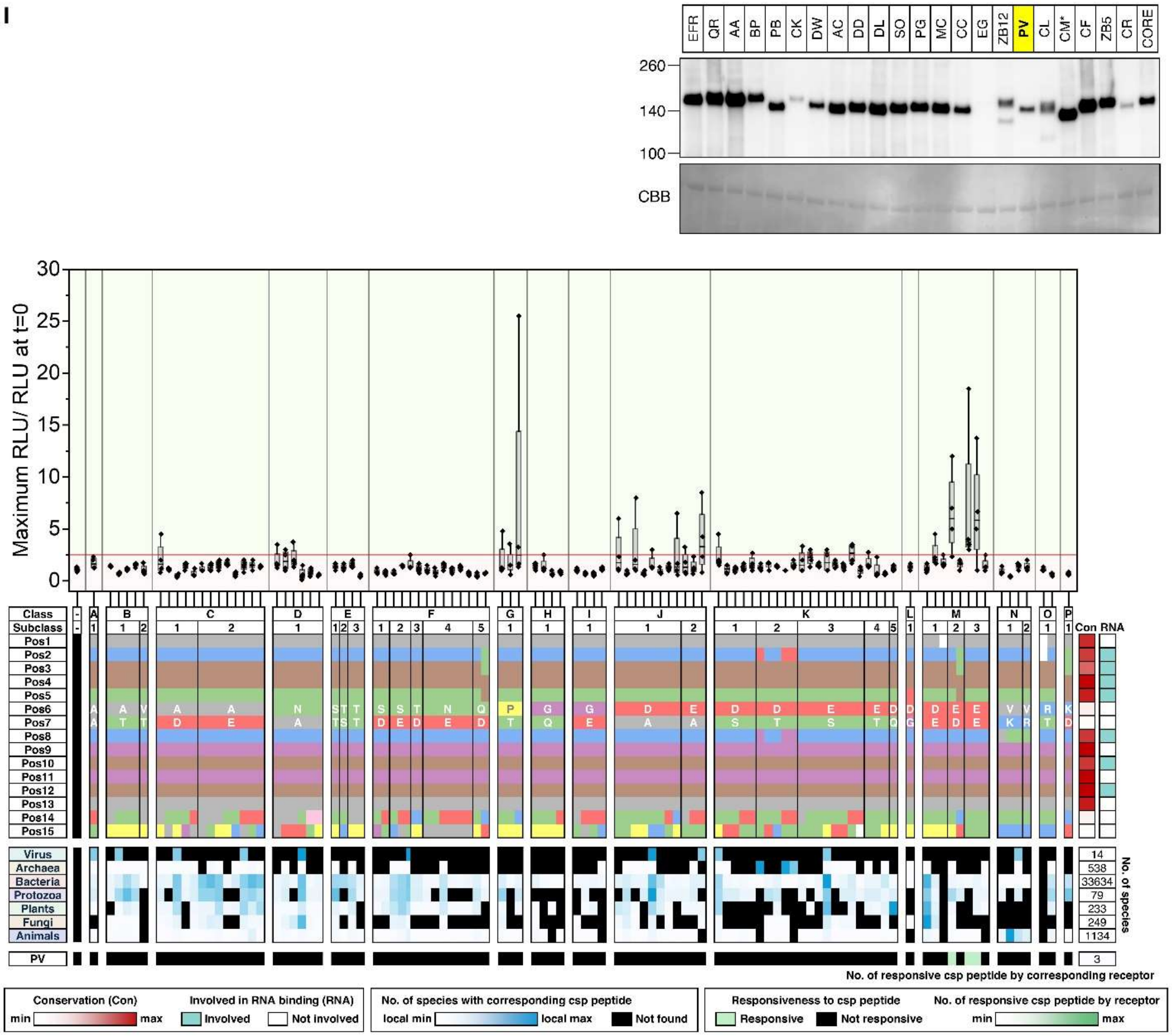

EFR QR AA BP PB CK DW AC DD DL SO PG MC CC EG ZB12 PV CL CM* CF ZB5 CR CORE
260
140
100
CBB
Maximum RLU/ RLU at t=0
Class
Subclass
Pos1
Pos2
Pos3
Pos4
Pos5
Pos6
Pos7
Pos8
Pos9
Pos10
Pos11
Pos12
Pos13
Pos14
Pos15
Con RNA
Virus
Archaea
Bacteria
Protozoa
Plants
Fungi
Animals
14
538
33634
79
233
249
1134
No. of species
PV
3
No. of responsive csp peptide by corresponding receptor
Conservation (Con)
min
max
Involved in RNA binding (RNA)
Involved
Not involved
No. of species with corresponding csp peptide
local min
local max
Not found
Responsiveness to csp peptide
Responsive
Not responsive
No. of responsive csp peptide by receptor
min
max

J

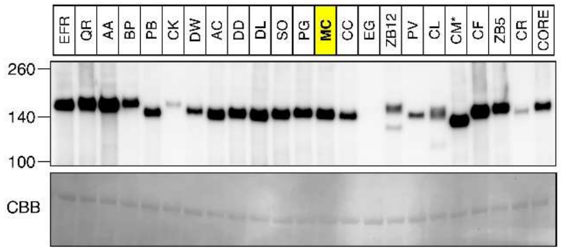
EFR
QR
AA
BP
PB
CK
DW
AC
DD
DL
SO
PG
MC
CC
EG
ZB12
PV
CL
CM*
CF
ZB5
CR
CORE
260
140
100
CBB

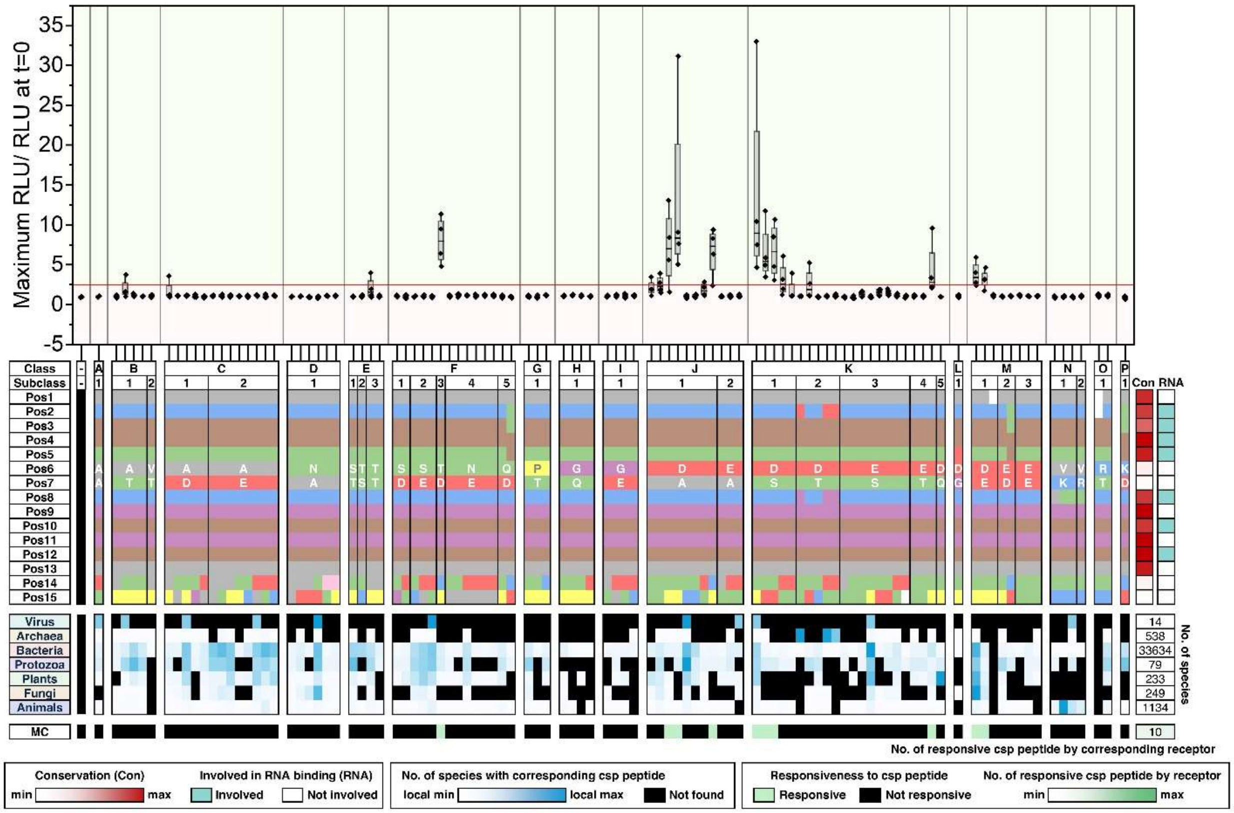
Maximum RLU/ RLU at t=0
Class
Subclass
Con
RNA
Virus
Archaea
Bacteria
Protozoa
Plants
Fungi
Animals
14
538
33634
79
233
249
1134
No. of species
MC
10
No. of responsive csp peptide by corresponding receptor
Conservation (Con)
min
max
Involved in RNA binding (RNA)
Involved
Not involved
No. of species with corresponding csp peptide
local min
local max
Not found
Responsiveness to csp peptide
Responsive
Not responsive
No. of responsive csp peptide by receptor
min
max

K

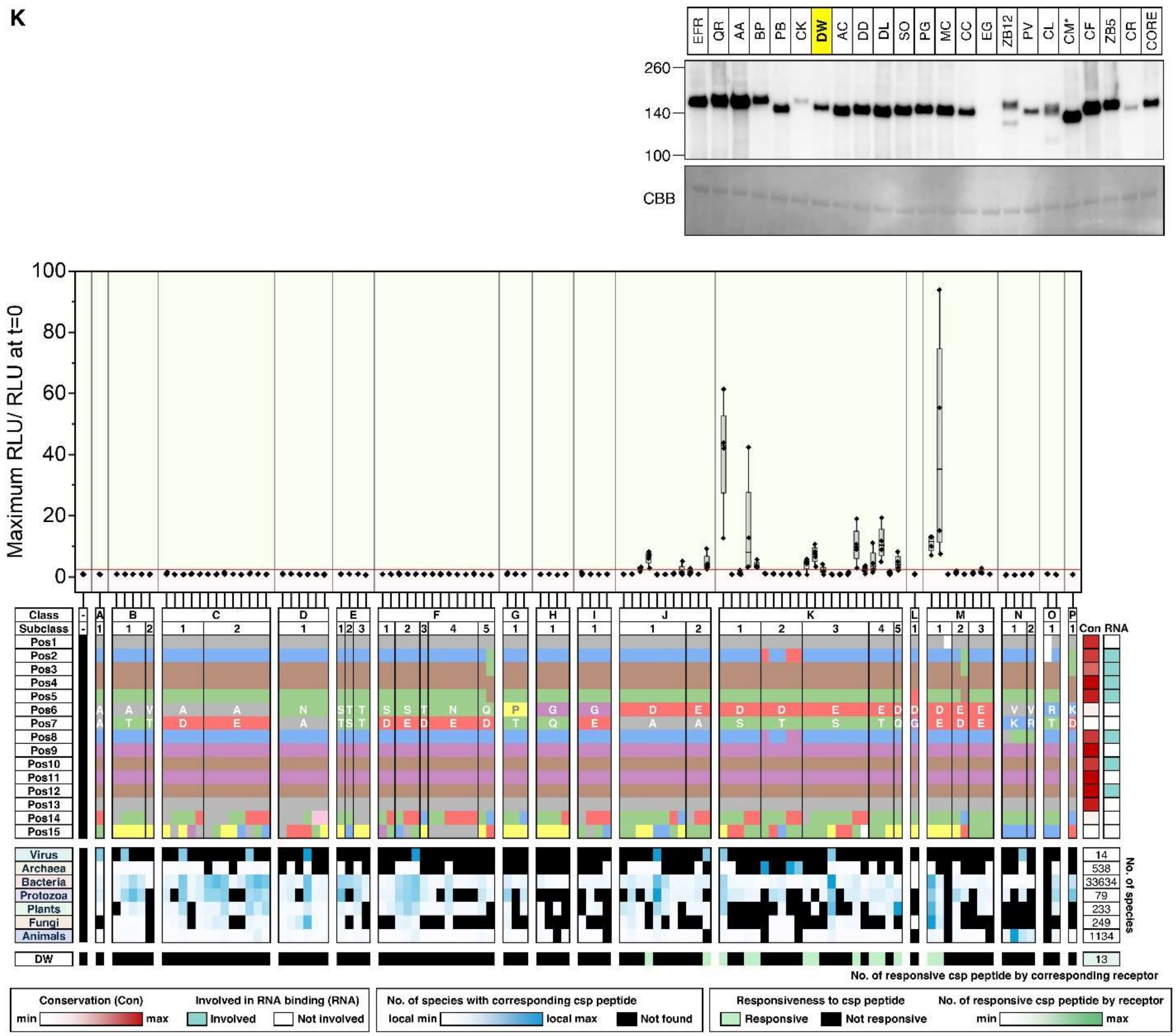

EFR QR AA BP PB CK DW AC DD DL SO PG MC CC EG ZB12 PV CL CM* CF ZB5 CR CORE
260
140
100
CBB
Maximum RLU/ RLU at t=0
Class
Subclass
Virus
Archaea
Bacteria
Protozoa
Plants
Fungi
Animals
DW
No. of species
No. of responsive csp peptide by corresponding receptor
Conservation (Con)
Involved in RNA binding (RNA)
Involved
Not involved
No. of species with corresponding csp peptide
local min
local max
Not found
Responsiveness to csp peptide
Responsive
Not responsive
No. of responsive csp peptide by receptor

L

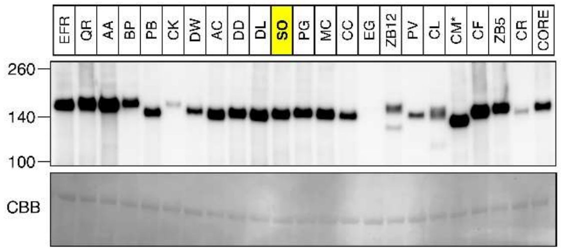
EFR
QR
AA
BP
PB
CK
DW
AC
DD
DL
SO
PG
MC
CC
EG
ZB12
PV
CL
CM*
CF
ZB5
CR
CORE
260
140
100
CBB

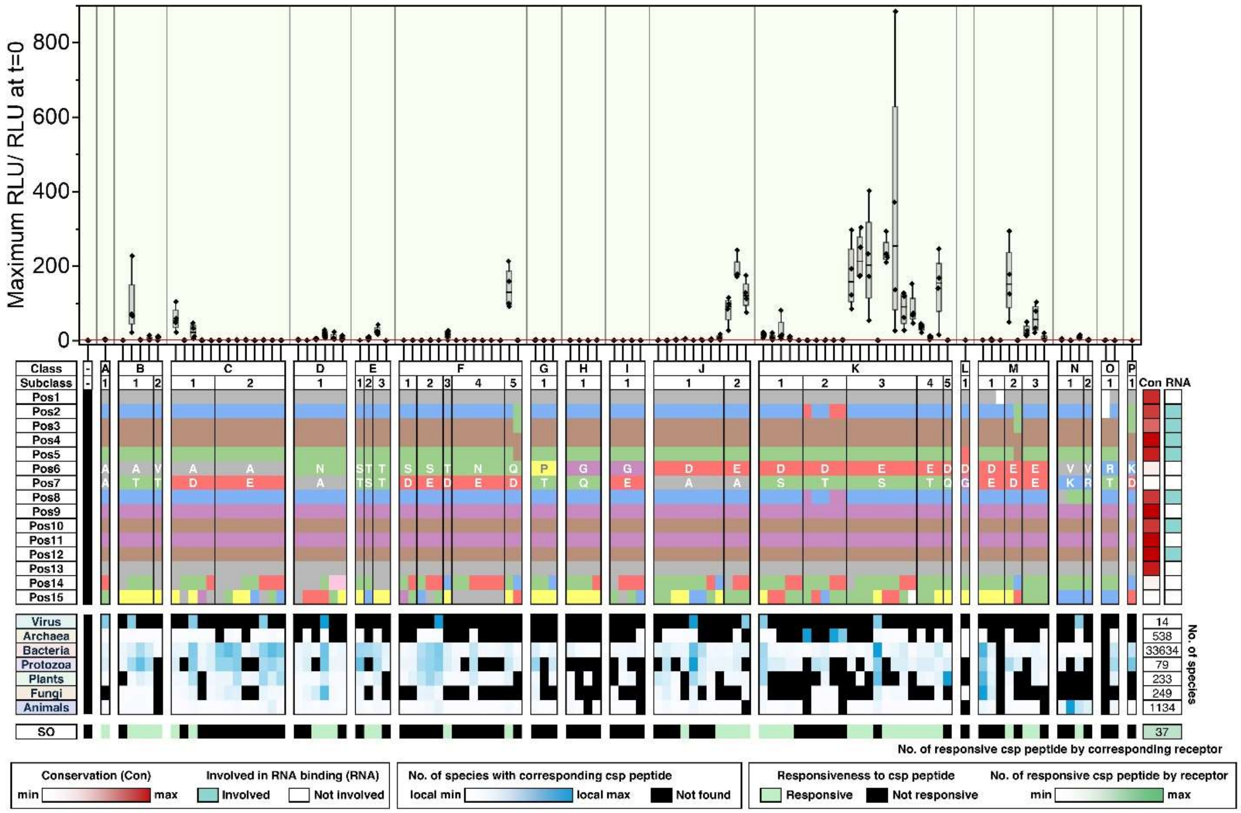
Maximum RLU/ RLU at t=0
800
600
400
200
0
Class
Subclass
Pos1
Pos2
Pos3
Pos4
Pos5
Pos6
Pos7
Pos8
Pos9
Pos10
Pos11
Pos12
Pos13
Pos14
Pos15
Con
RNA
Virus
Archaea
Bacteria
Protozoa
Plants
Fungi
Animals
No. of species
14
538
33634
79
233
249
1134
SO
37
No. of responsive csp peptide by corresponding receptor
Conservation (Con)
min
max
Involved in RNA binding (RNA)
Involved
Not involved
No. of species with corresponding csp peptide
local min
local max
Not found
Responsiveness to csp peptide
Responsive
Not responsive
No. of responsive csp peptide by receptor
min
max

**M**

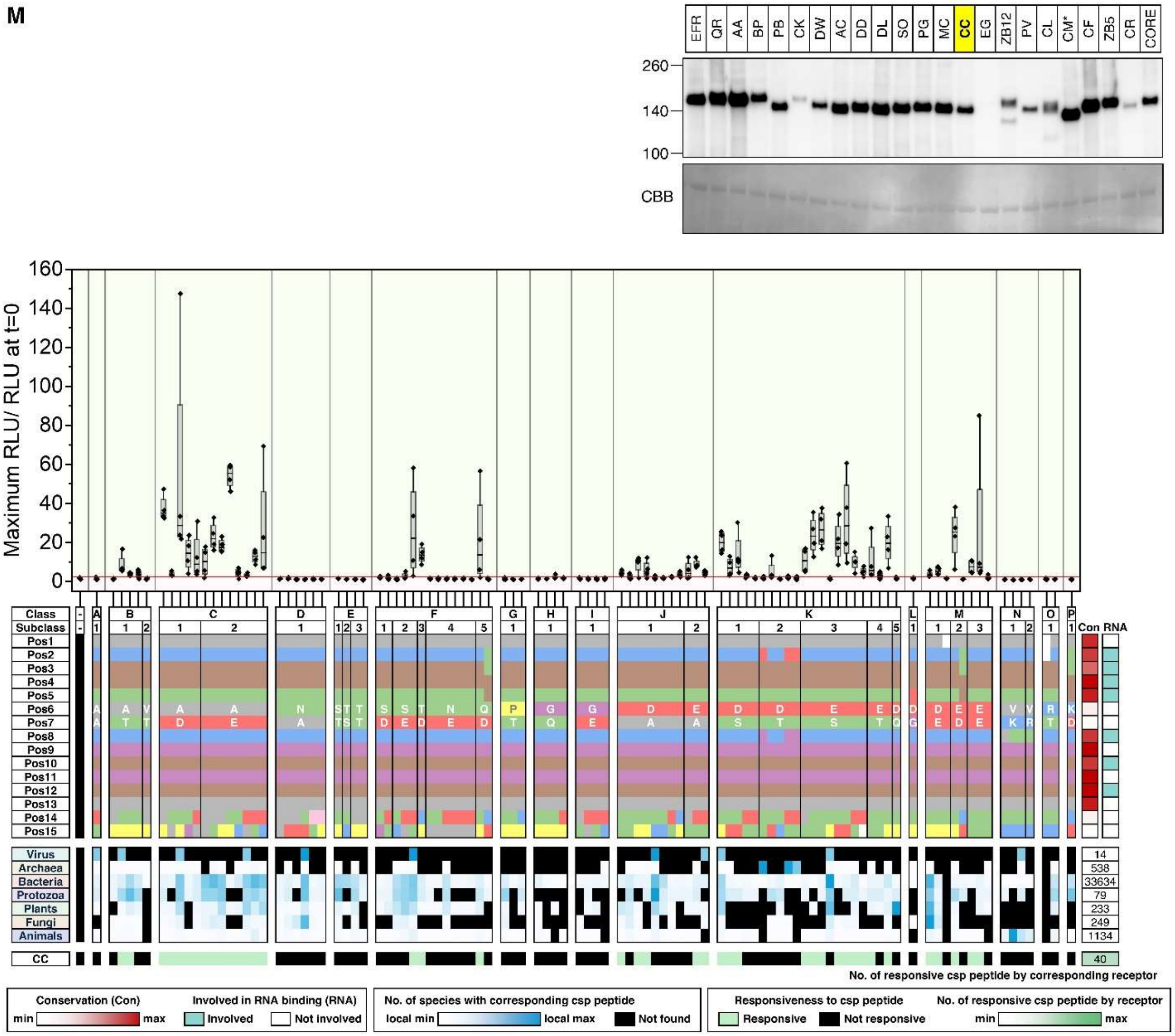

EFR QR AA BP PB CK DW AC DD DL SO PG MC CC EG ZB12 PV CL CM* CF ZB5 CR CORE
260
140
100
CBB
Maximum RLU/ RLU at t=0
Class
Subclass
Virus
Archaea
Bacteria
Protozoa
Plants
Fungi
Animals
CC
Con RNA
No. of responsive csp peptide by corresponding receptor
Conservation (Con)
min
max
Involved in RNA binding (RNA)
Involved
Not involved
No. of species with corresponding csp peptide
local min
local max
Not found
Responsiveness to csp peptide
Responsive
Not responsive
No. of responsive csp peptide by receptor

N

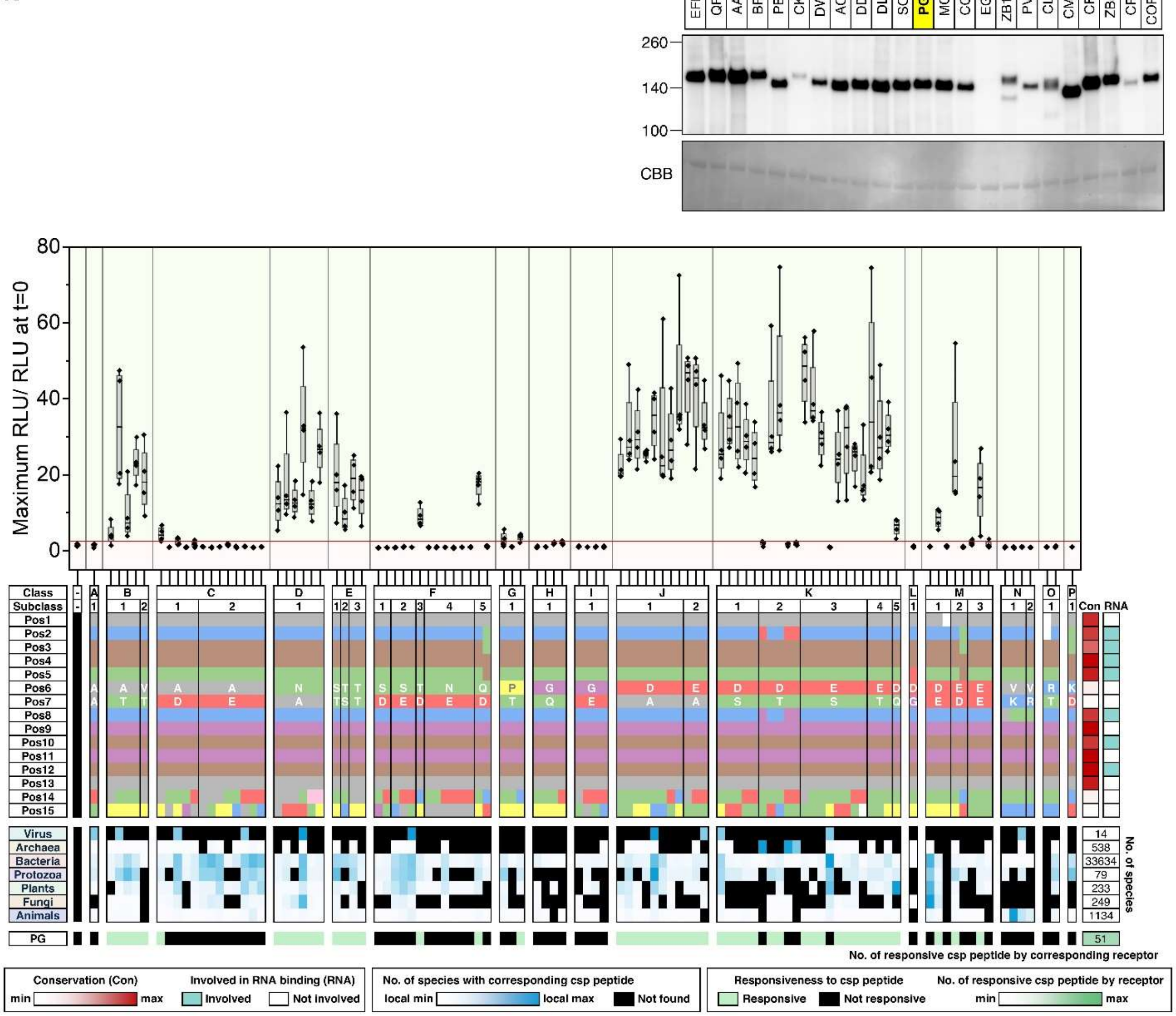
EFR QR AA BP PB CK DW AC DD DL SO PG MC CC EG ZB12 PV CL CM* CF ZB5 CR CORE
260
140
100
CBB
Maximum RLU/ RLU at t=0
Class
Subclass
Con RNA
No. of species
No. of responsive csp peptide by corresponding receptor
Conservation (Con)
min
max
Involved in RNA binding (RNA)
Involved
Not involved
No. of species with corresponding csp peptide
local min
local max
Not found
Responsiveness to csp peptide
Responsive
Not responsive
No. of responsive csp peptide by receptor
min
max

o

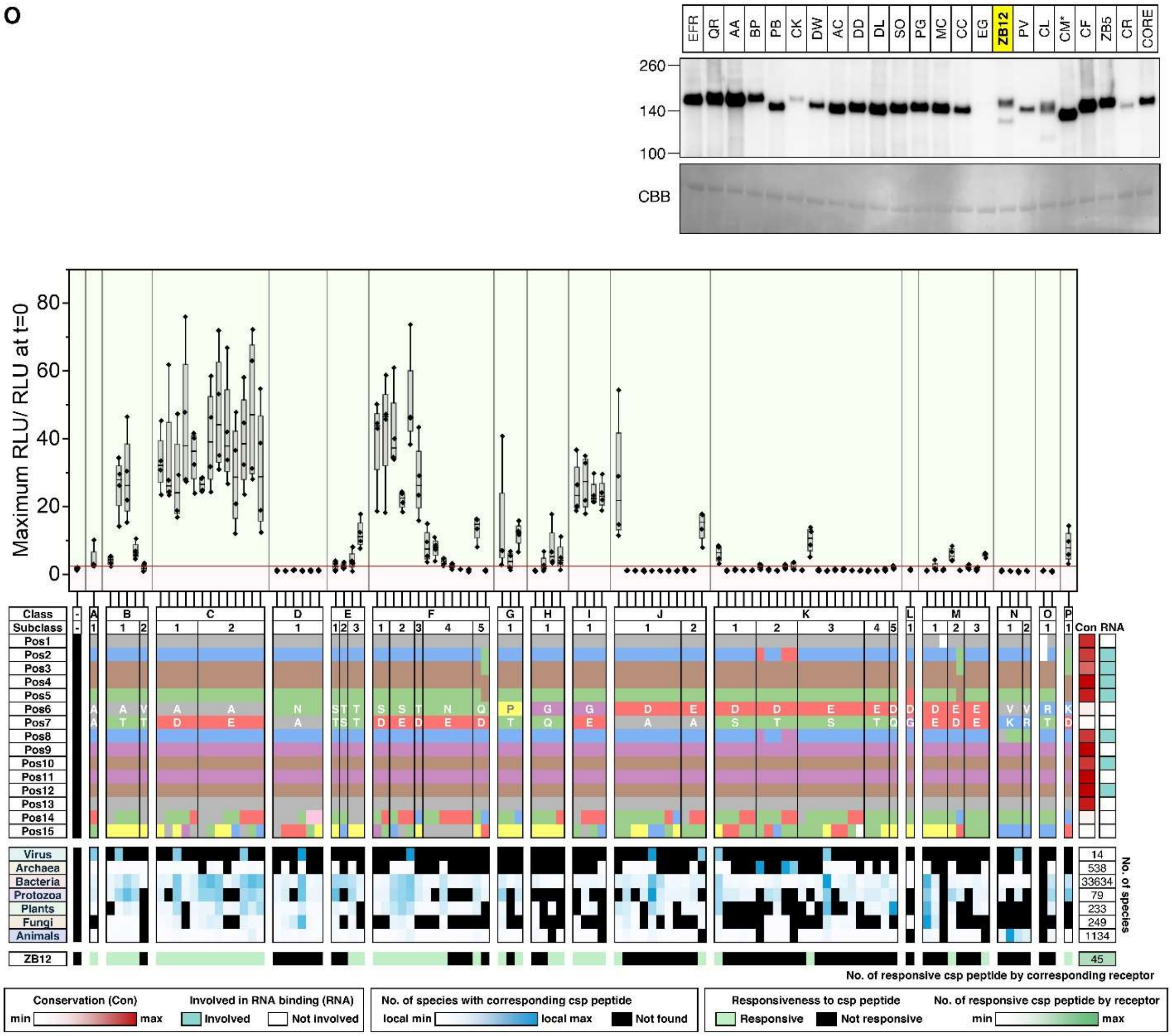
EFR
QR
AA
BP
PB
CK
DW
AC
DD
DL
SO
PG
MC
CC
EG
ZB12
PV
CL
CM*
CF
ZB5
CR
CORE
260
140
100
CBB
Maximum RLU/ RLU at t=0
80
60
40
20
0
Class
Subclass
Pos1
Pos2
Pos3
Pos4
Pos5
Pos6
Pos7
Pos8
Pos9
Pos10
Pos11
Pos12
Pos13
Pos14
Pos15
Con
RNA
Virus
Archaea
Bacteria
Protozoa
Plants
Fungi
Animals
14
538
33634
79
233
249
1134
No. of species
ZB12
45
No. of responsive csp peptide by corresponding receptor
Conservation (Con)
min
max
Involved in RNA binding (RNA)
Involved
Not involved
No. of species with corresponding csp peptide
local min
local max
Not found
Responsiveness to csp peptide
Responsive
Not responsive
No. of responsive csp peptide by receptor
min
max

**P**

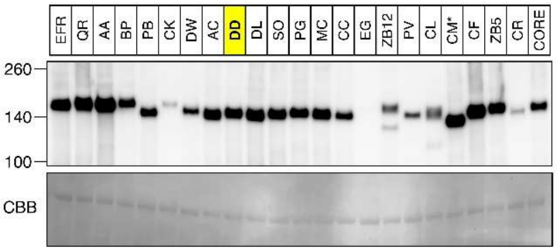
EFR
QR
AA
BP
PB
CK
DW
AC
DD
DL
SO
PG
MC
CC
EG
ZB12
PV
CL
CM*
CF
ZB5
CR
CORE
260
140
100
CBB

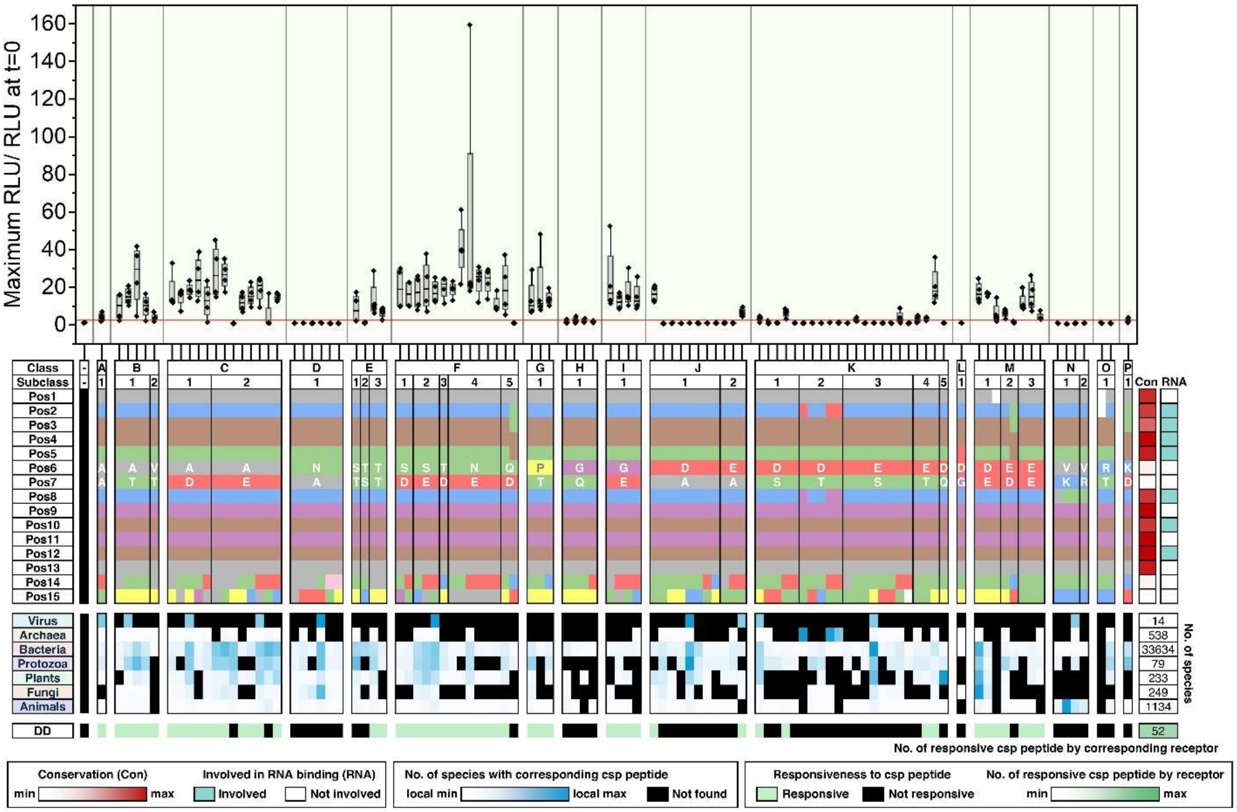
Maximum RLU/ RLU at t=0
160
140
120
100
80
60
40
20
0
Class
Subclass
Pos1
Pos2
Pos3
Pos4
Pos5
Pos6
Pos7
Pos8
Pos9
Pos10
Pos11
Pos12
Pos13
Pos14
Pos15
Con
RNA
Virus
Archaea
Bacteria
Protozoa
Plants
Fungi
Animals
14
538
33634
79
233
249
1134
No. of species
DD
52
No. of responsive csp peptide by corresponding receptor
Conservation (Con)
min
max
Involved in RNA binding (RNA)
Involved
Not involved
No. of species with corresponding csp peptide
local min
local max
Not found
Responsiveness to csp peptide
Responsive
Not responsive
No. of responsive csp peptide by receptor
min
max

Q

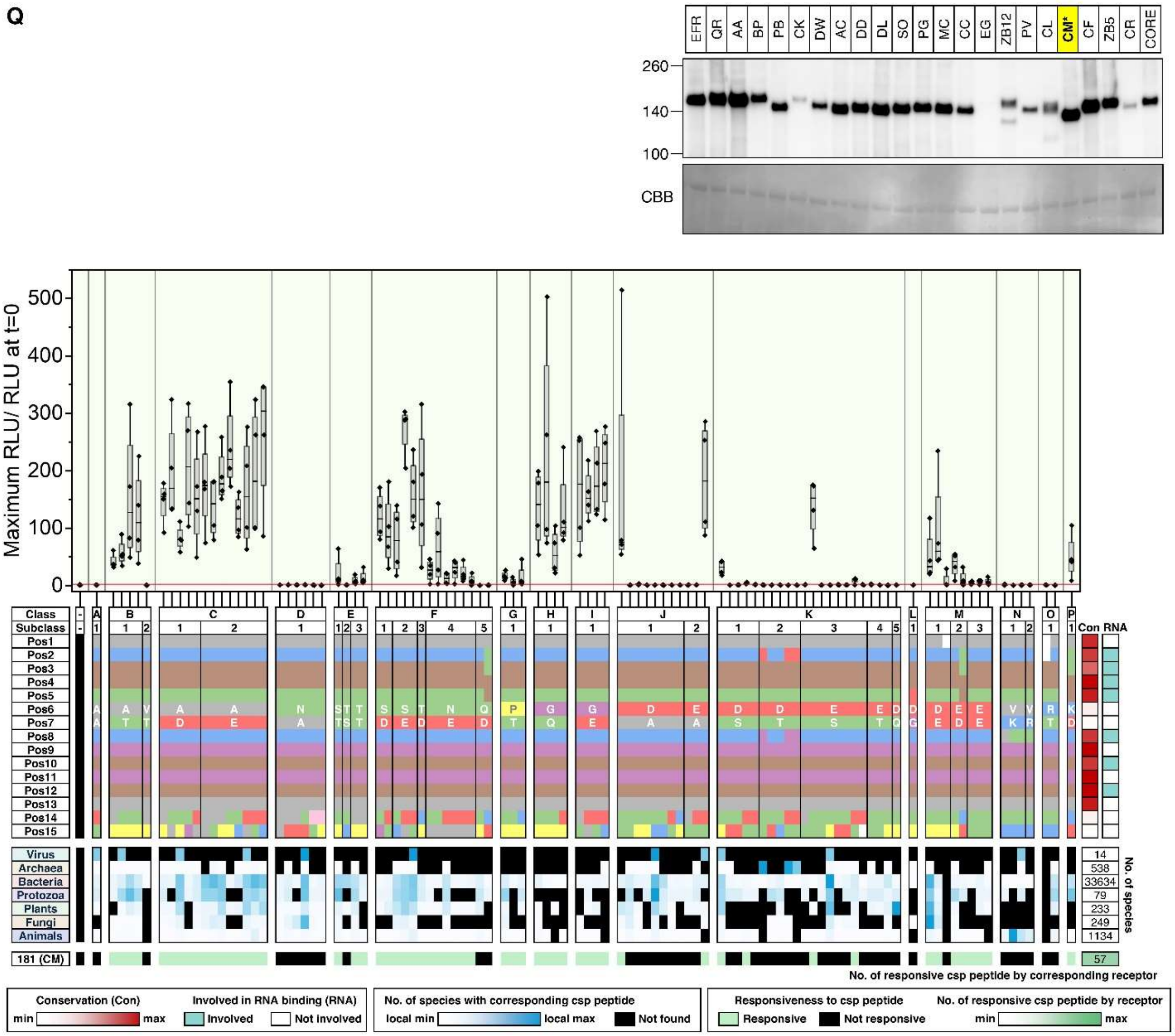
EFR QR AA BP PB CK DW AC DD DL SO PG MC CC EG ZB12 PV CL CM* CF ZB5 CR CORE
260
140
100
CBB
Maximum RLU/ RLU at t=0
Class
Subclass
Con RNA
Virus
Archaea
Bacteria
Protozoa
Plants
Fungi
Animals
No. of species
14
538
33634
79
233
249
1134
181 (CM)
57
No. of responsive csp peptide by corresponding receptor
Conservation (Con)
min
max
Involved in RNA binding (RNA)
Involved
Not involved
No. of species with corresponding csp peptide
local min
local max
Not found
Responsiveness to csp peptide
Responsive
Not responsive
No. of responsive csp peptide by receptor
min
max

**R**

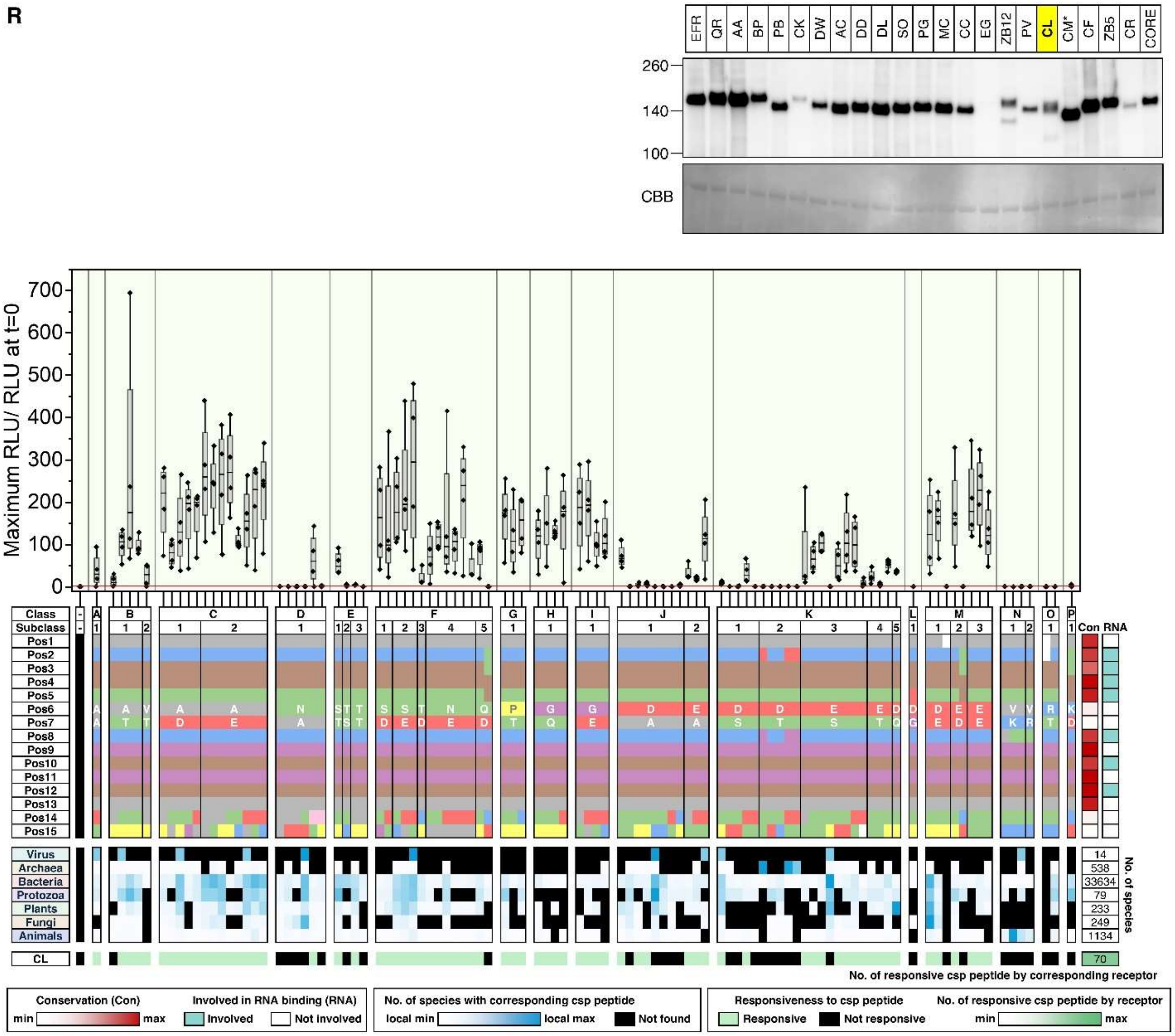

EFR
QR
AA
BP
PB
CK
DW
AC
DD
DL
SO
PG
MC
CC
EG
ZB12
PV
CL
CM*
CF
ZB5
CR
CORE
260
140
100
CBB
Maximum RLU/ RLU at t=0
Class
Subclass
Con
RNA
Virus
Archaea
Bacteria
Protozoa
Plants
Fungi
Animals
CL
No. of species
No. of responsive csp peptide by corresponding receptor
Conservation (Con)
Involved in RNA binding (RNA)
No. of species with corresponding csp peptide
Responsiveness to csp peptide
No. of responsive csp peptide by receptor

**S**

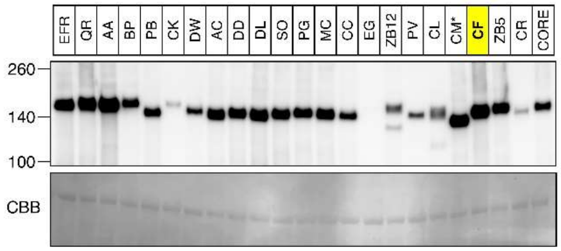
EFR QR AA BP PB CK DW AC DD DL SO PG MC CC EG ZB12 PV CL CM* CF ZB5 CR CORE
260
140
100
CBB

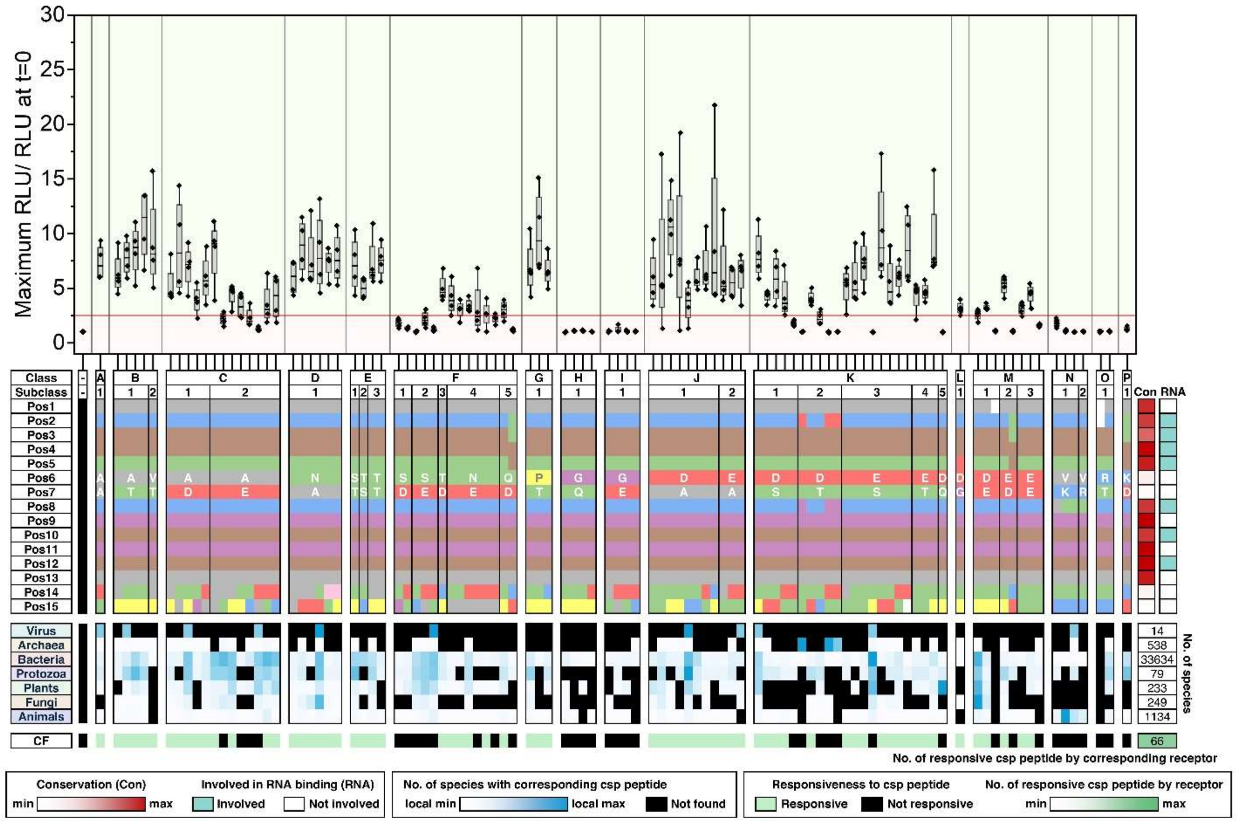
Maximum RLU/ RLU at t=0
30
25
20
15
10
5
0
Class
Subclass
Pos1
Pos2
Pos3
Pos4
Pos5
Pos6
Pos7
Pos8
Pos9
Pos10
Pos11
Pos12
Pos13
Pos14
Pos15
Con RNA
Virus
Archaea
Bacteria
Protozoa
Plants
Fungi
Animals
14
538
33634
79
233
249
1134
No. of species
CF
66
No. of responsive csp peptide by corresponding receptor
Conservation (Con)
min
max
Involved in RNA binding (RNA)
Involved
Not involved
No. of species with corresponding csp peptide
local min
local max
Not found
Responsiveness to csp peptide
Responsive
Not responsive
No. of responsive csp peptide by receptor
min
max

T

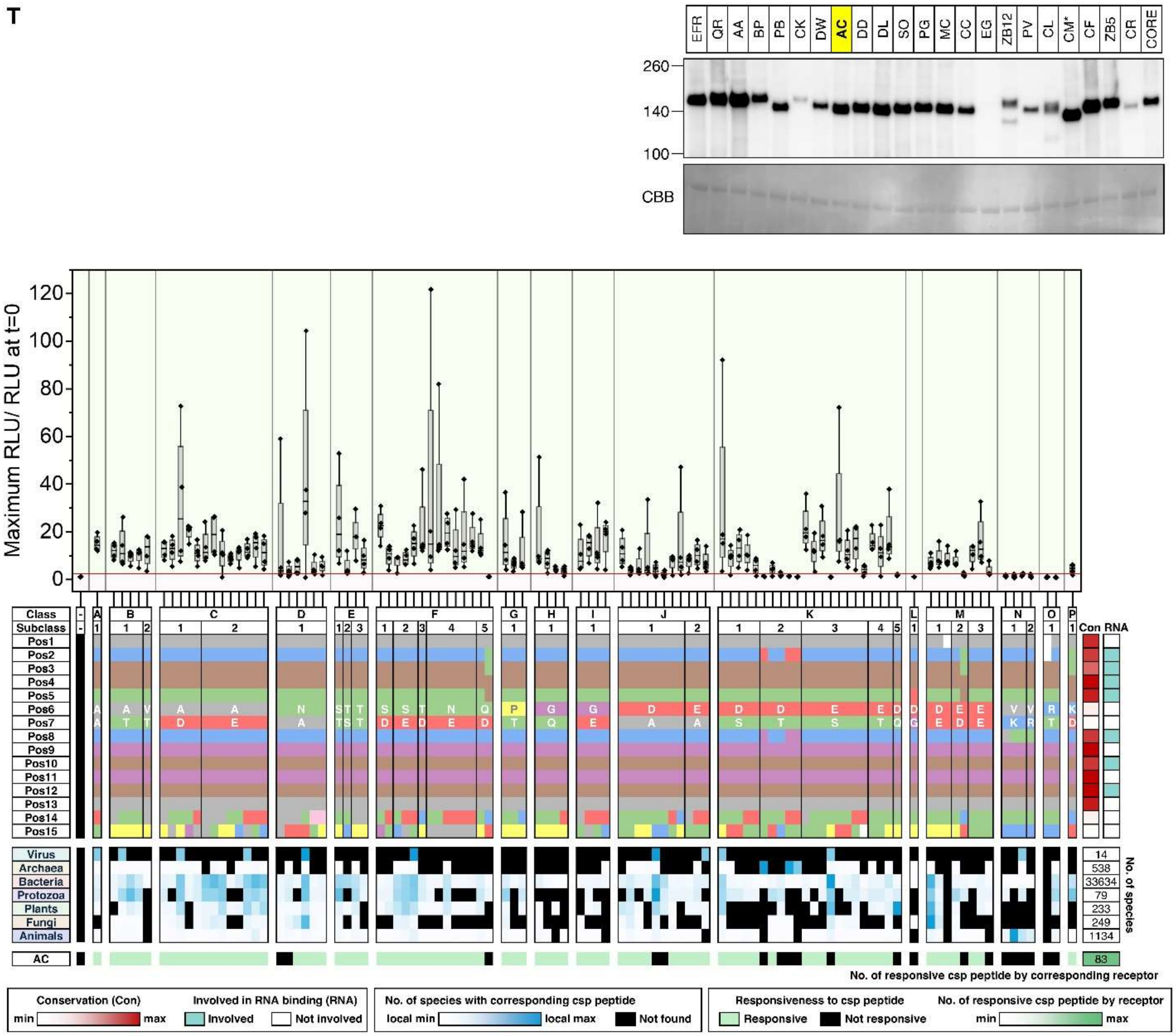
EFR QR AA BP PB CK DW AC DD DL SO PG MC CC EG ZB12 PV CL CM* CF ZB5 CR CORE
260
140
100
CBB
Maximum RLU/ RLU at t=0
Class
Subclass
Virus
Archaea
Bacteria
Protozoa
Plants
Fungi
Animals
AC
Con RNA
No. of species
14
538
33634
79
233
249
1134
83
No. of responsive csp peptide by corresponding receptor
Conservation (Con)
min
max
Involved in RNA binding (RNA)
Involved
Not involved
No. of species with corresponding csp peptide
local min
local max
Not found
Responsiveness to csp peptide
Responsive
Not responsive
No. of responsive csp peptide by receptor
min
max

U

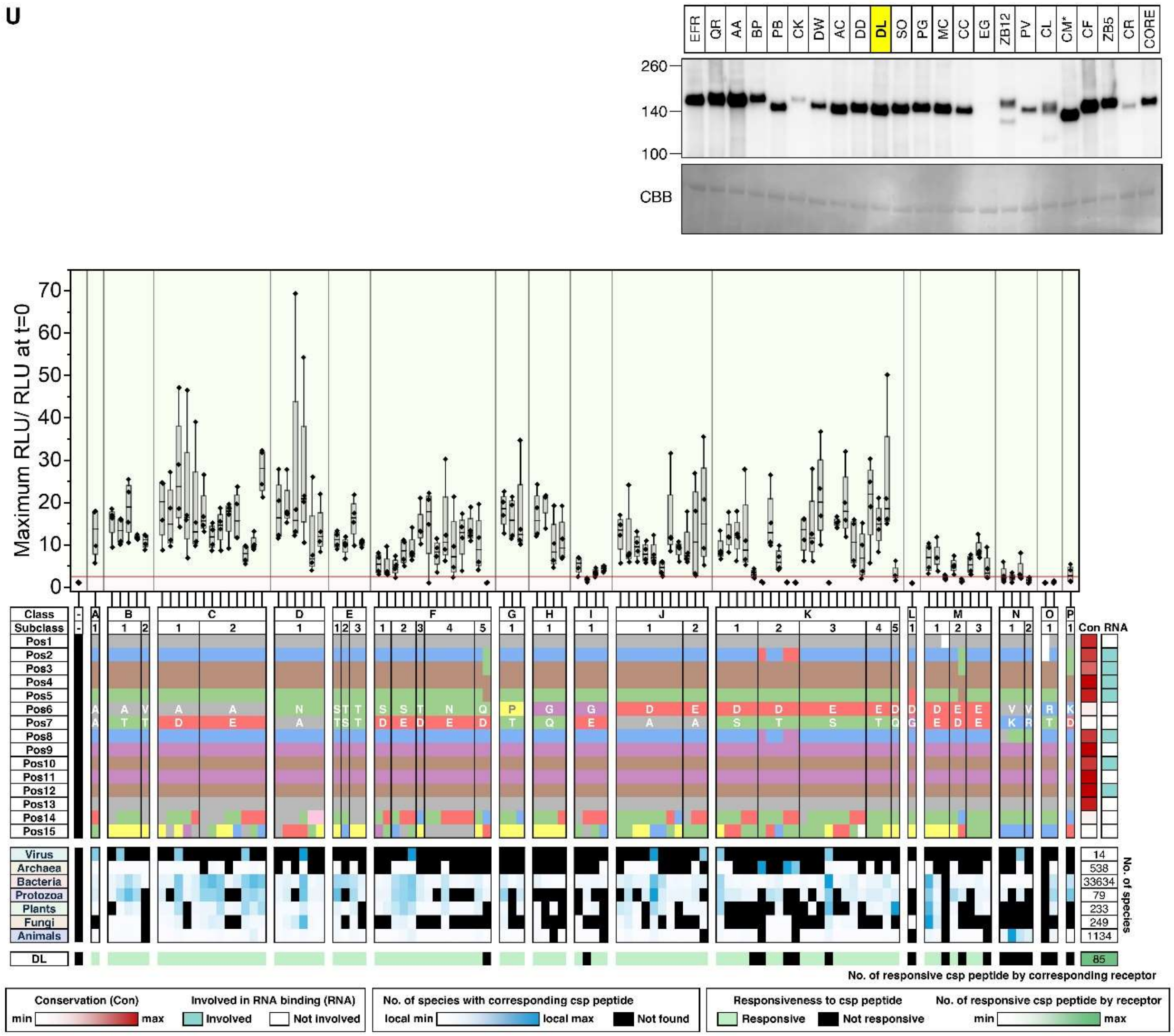
EFR QR AA BP PB CK DW AC DD DL SO PG MC CC EG ZB12 PV CL CM* CF ZB5 CR CORE
260
140
100
CBB
Maximum RLU/ RLU at t=0
Class
Subclass
Pos1
Pos2
Pos3
Pos4
Pos5
Pos6
Pos7
Pos8
Pos9
Pos10
Pos11
Pos12
Pos13
Pos14
Pos15
Con
RNA
Virus
Archaea
Bacteria
Protozoa
Plants
Fungi
Animals
14
538
33634
79
233
249
1134
No. of species
DL
85
No. of responsive csp peptide by corresponding receptor
Conservation (Con)
min
max
Involved in RNA binding (RNA)
Involved
Not involved
No. of species with corresponding csp peptide
local min
local max
Not found
Responsiveness to csp peptide
Responsive
Not responsive
No. of responsive csp peptide by receptor
min
max

**V**

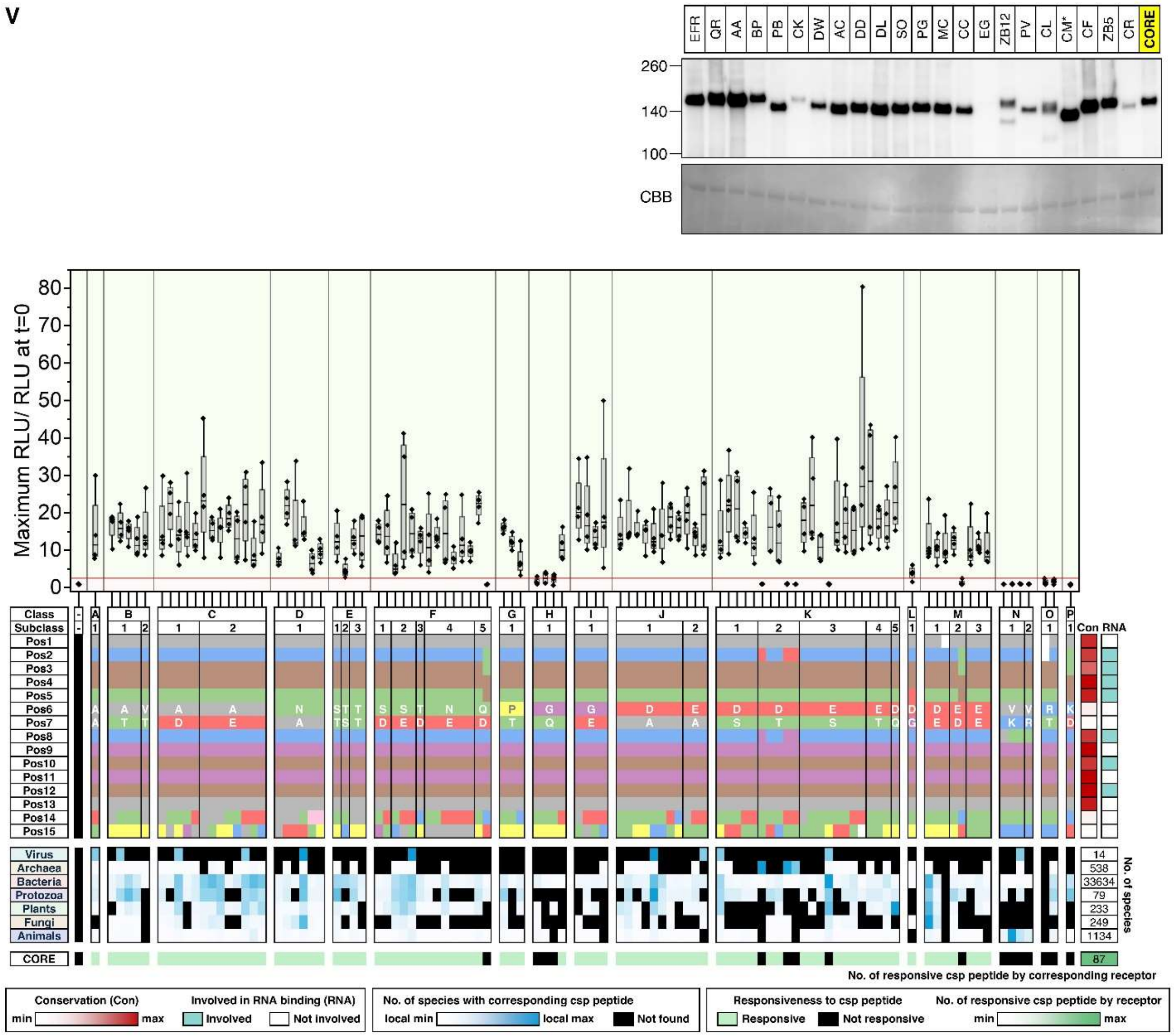

EFR QR AA BP PB CK DW AC DD DL SO PG MC CC EG ZB12 PV CL CM* CF ZB5 CR CORE
260
140
100
CBB
Maximum RLU/ RLU at t=0
Class
Subclass
Pos1
Pos2
Pos3
Pos4
Pos5
Pos6
Pos7
Pos8
Pos9
Pos10
Pos11
Pos12
Pos13
Pos14
Pos15
Con RNA
Virus
Archaea
Bacteria
Protozoa
Plants
Fungi
Animals
No. of species
14
538
33634
79
233
249
1134
CORE
87
No. of responsive csp peptide by corresponding receptor
Conservation (Con)
min max
Involved in RNA binding (RNA)
Involved
Not involved
No. of species with corresponding csp peptide
local min
local max
Not found
Responsiveness to csp peptide
Responsive
Not responsive
No. of responsive csp peptide by receptor
min max

W

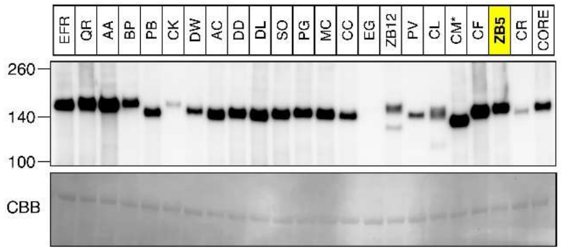
EFR QR AA BP PB CK DW AC DD DL SO PG MC CC EG ZB12 PV CL CM* CF ZB5 CR CORE
260
140
100
CBB

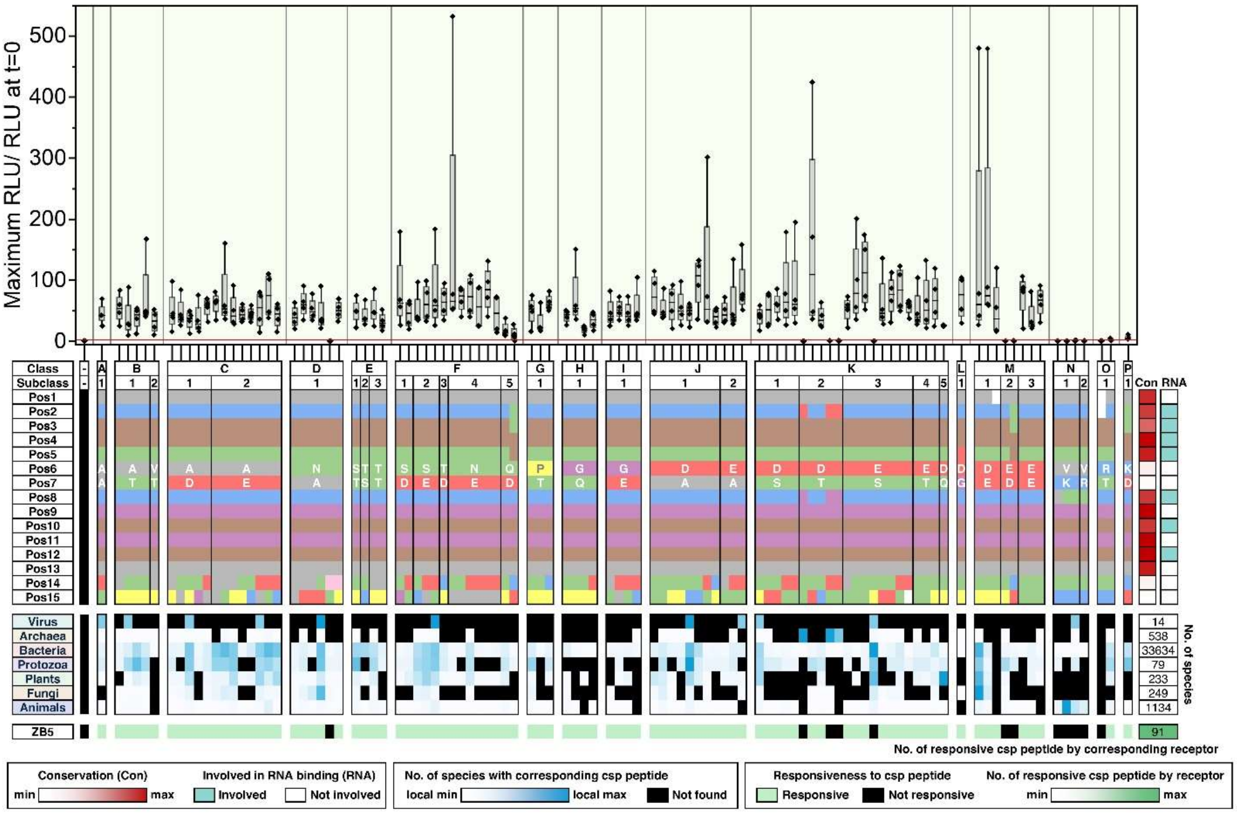
Maximum RLU/ RLU at t=0
500
400
300
200
100
0
Class
Subclass
Pos1
Pos2
Pos3
Pos4
Pos5
Pos6
Pos7
Pos8
Pos9
Pos10
Pos11
Pos12
Pos13
Pos14
Pos15
Con RNA
Virus
Archaea
Bacteria
Protozoa
Plants
Fungi
Animals
14
538
33634
79
233
249
1134
No. of species
ZB5
91
No. of responsive csp peptide by corresponding receptor
Conservation (Con)
min
max
Involved in RNA binding (RNA)
Involved
Not involved
No. of species with corresponding csp peptide
local min
local max
Not found
Responsiveness to csp peptide
Responsive
Not responsive
No. of responsive csp peptide by receptor
min
max

**Fig. S17. ROS data related to Fig 4D.**

Top immunoblot: Samples used in the following ROS assays were analysed with immunoblotting to confirm receptor expression. CBB staining is included as loading controls. Middle bar chart represents the ROS responses to mock treatment or each of the 103 csp15 peptide by the corresponding receptors from (**A-W**). Leaves were then treated with 100 nM of indicated peptides, and ROS production was recorded for 15 minutes. For the box plots, The center line represents the median; the box bounds represent the 25th and 75th percentiles; and the whiskers extend to 1.5 × the interquartile range (IQR) from the 25th and 75th percentiles. Data points from 4 technical replicates were analyzed. A peptide was considered active if at least 3 out of 4 technical replicates showed ROS levels exceeding the baseline by 2.5 times (marked by the red line). Bottom charts: The top chart displays the 103 csp15 peptides synthesized in this study. Amino acid properties at each position (Pos) are color-coded, as in in Fig. 4A. Peptides are grouped into classes and subclasses based on residues at the positions 6 and 7. A red heatmap on the right indicates residue conservation scores, while cyan blocks mark RNA-binding residues in CSPs. The middle blue heatmap shows the prevalence of each csp15 peptide across kingdom. Black blocks represent peptides absent from all species within a kingdom. The number of species analyzed in each kingdom is shown on the right. Bottom chart represents the CSP recognition capacity of the corresponding receptors in plants. As noted above, peptides were considered active if at least 3 out of 4 technical replicates showed ROS levels exceeding the baseline by 2.5 times (see above). Non-responsive peptides are labeled in black. EFR served as a negative control, while CORE was used as a position control. The green heatmap on the right indicates the number of responsive peptides (out of 103) for each receptor.

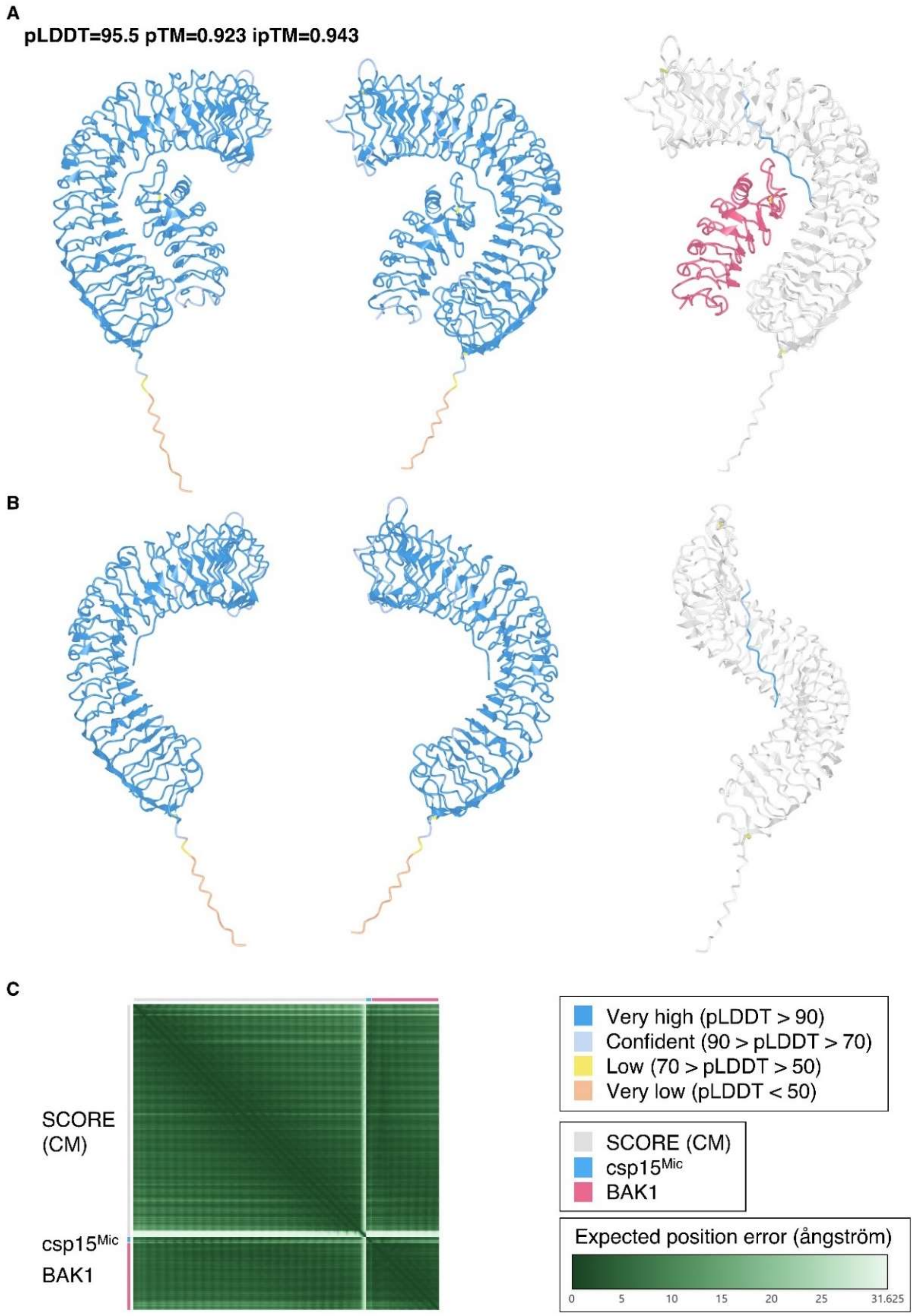


**Fig. S18. Alphafold2 structure prediction of SCORE$^{Ecto}$-csp15-BAK1$^{Ecto}$ interactions.**

**(A)** Alphafold2 structure prediction of SCORE$^{Ecto}$(CM)-csp15-BAK1$^{Ecto}$ complex. The two structures on the left are color-coded by pLDDT scores. The structure on the right is color-coded by chains (SCORE$^{Ecto}$(CM), csp15, and BAK1$^{Ecto}$). Predicted local distance difference test (pLDDT), predicted template modelling (pTM), and interface predicted template modelling (ipTM) scores are indicated. **(B)** The same structure shown in (A), without $BAK1^{Ecto}$. **(C)** The Predicted Aligned Error (PAE) matrix corresponding to the structure prediction. Protein structures were visualized by iCn3D and PAE matrix was visualized by PAE Viewer (41,58).

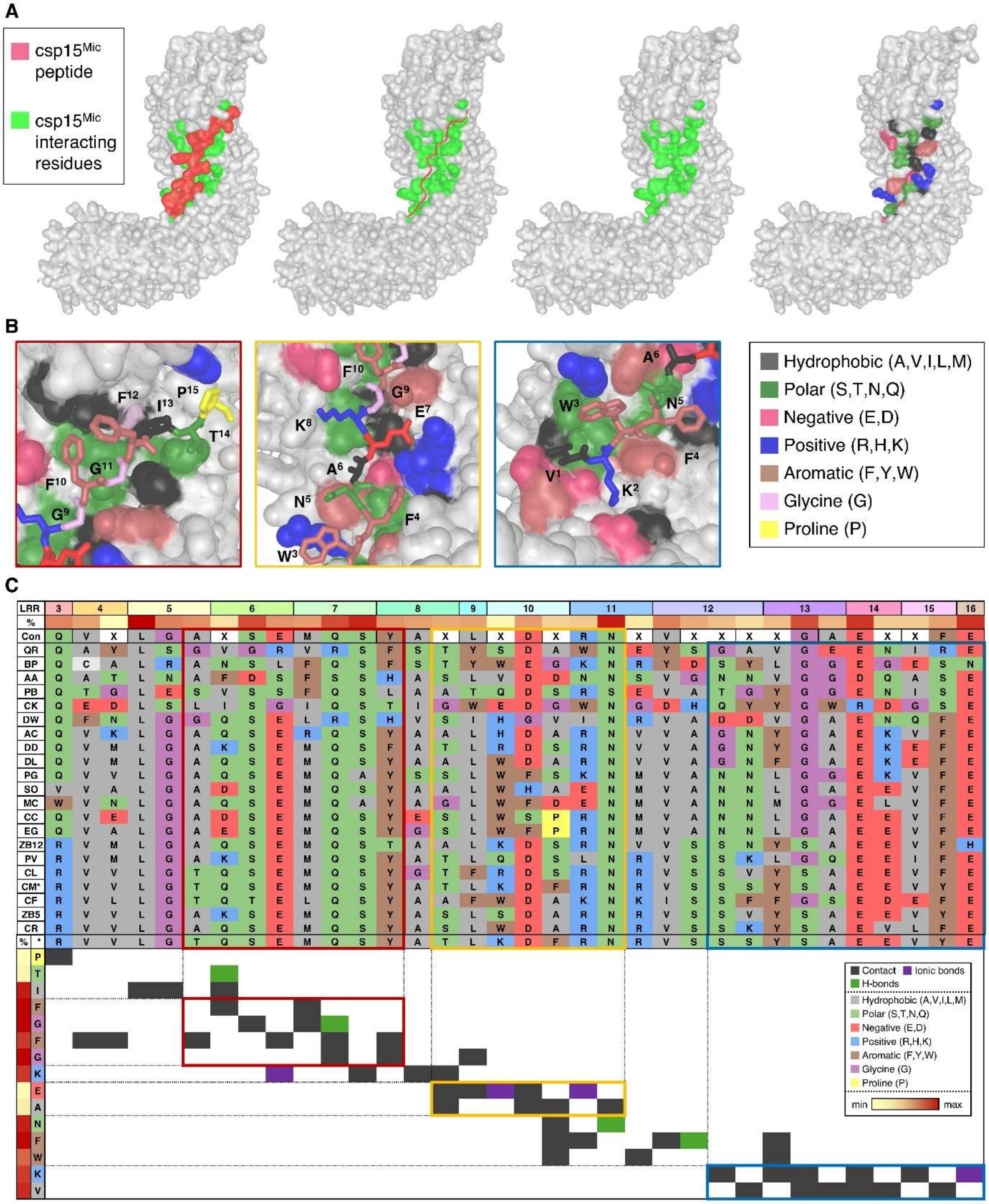


**Fig. S19. Interaction between SCORE$^{Ecto}$ (CM) and csp15.**

**(A)** Alphafold2 structure prediction of SCORE$^{Ecto}$(CM)-csp15 interactions. Left structure: Csp15 peptide is highlighted in red and csp15-interacting residues in SCORE are shown in green. Second left structure: csp15 peptide is displayed as a strand. Second right structure: csp15 peptide is removed to reveal the csp15-interacting residues. Right structure: csp15-interacting residues are color-coded by amino acid property as indicated in the box below. **(B)** The three main interaction sites in red, yellow, and blue boxes in Fig 5A. Residues in SCORE(CM) involved in the interactions with csp15 are color-coded by amino acid property as indicated in the box in the right. **(C)** Interaction map between SCORE and csp15. Top chart represents the alignment of residues involved in csp15 binding across SCORE orthologs. Amino acids are color-coded by properties, as indicated in the box below. Bottom chart represents contact sites between SCORE(CM) and csp15, indicated by colored blocks indicated on the right. Contact sites within red, yellow, and blue rectangles are corresponding to colored boxes in (A). Protein structures are visualized with iCn3D (41).

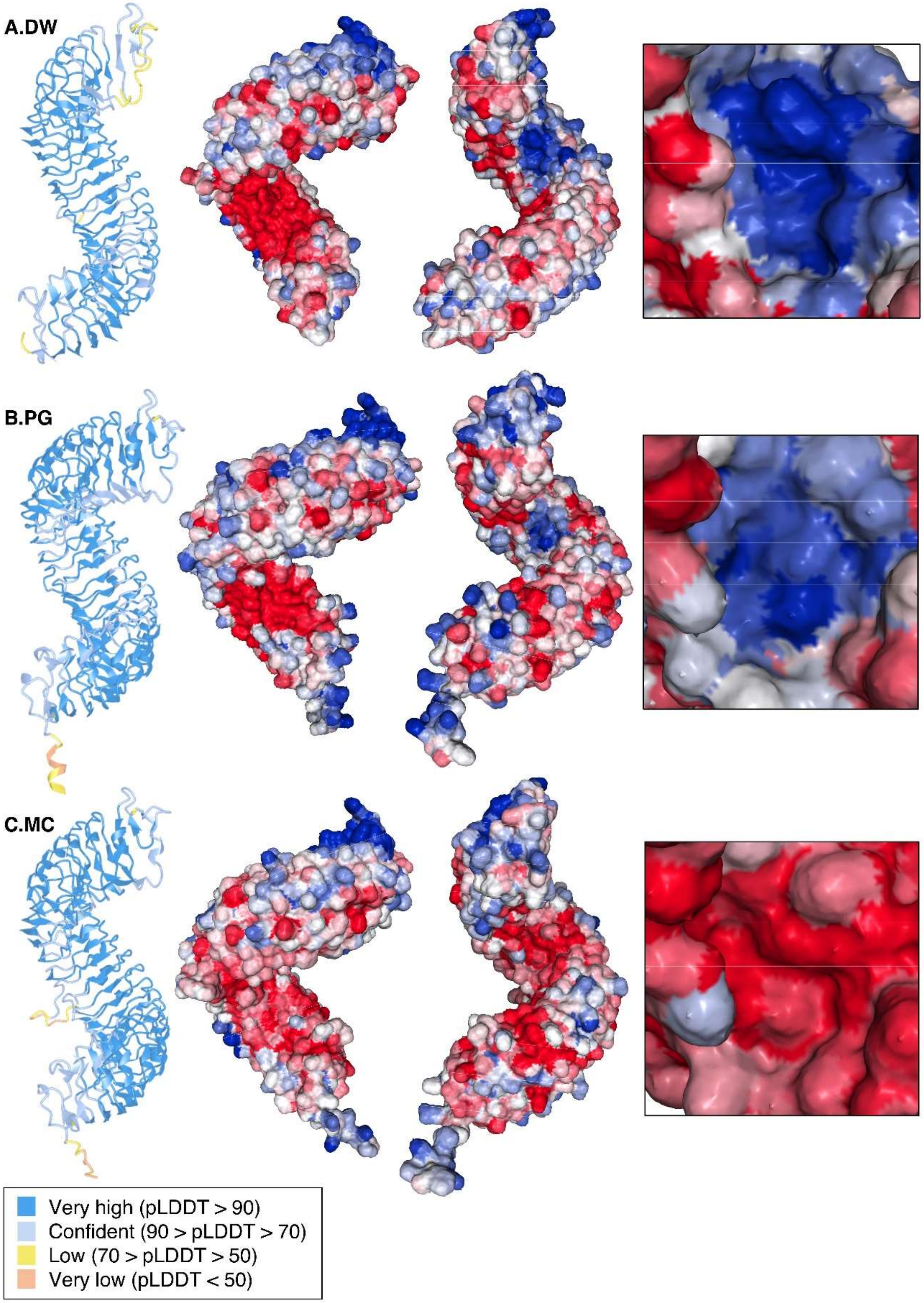
A.DW
B.PG
C.MC
Very high (pLDDT > 90)
Confident (90 > pLDDT > 70)
Low (70 > pLDDT > 50)
Very low (pLDDT < 50)

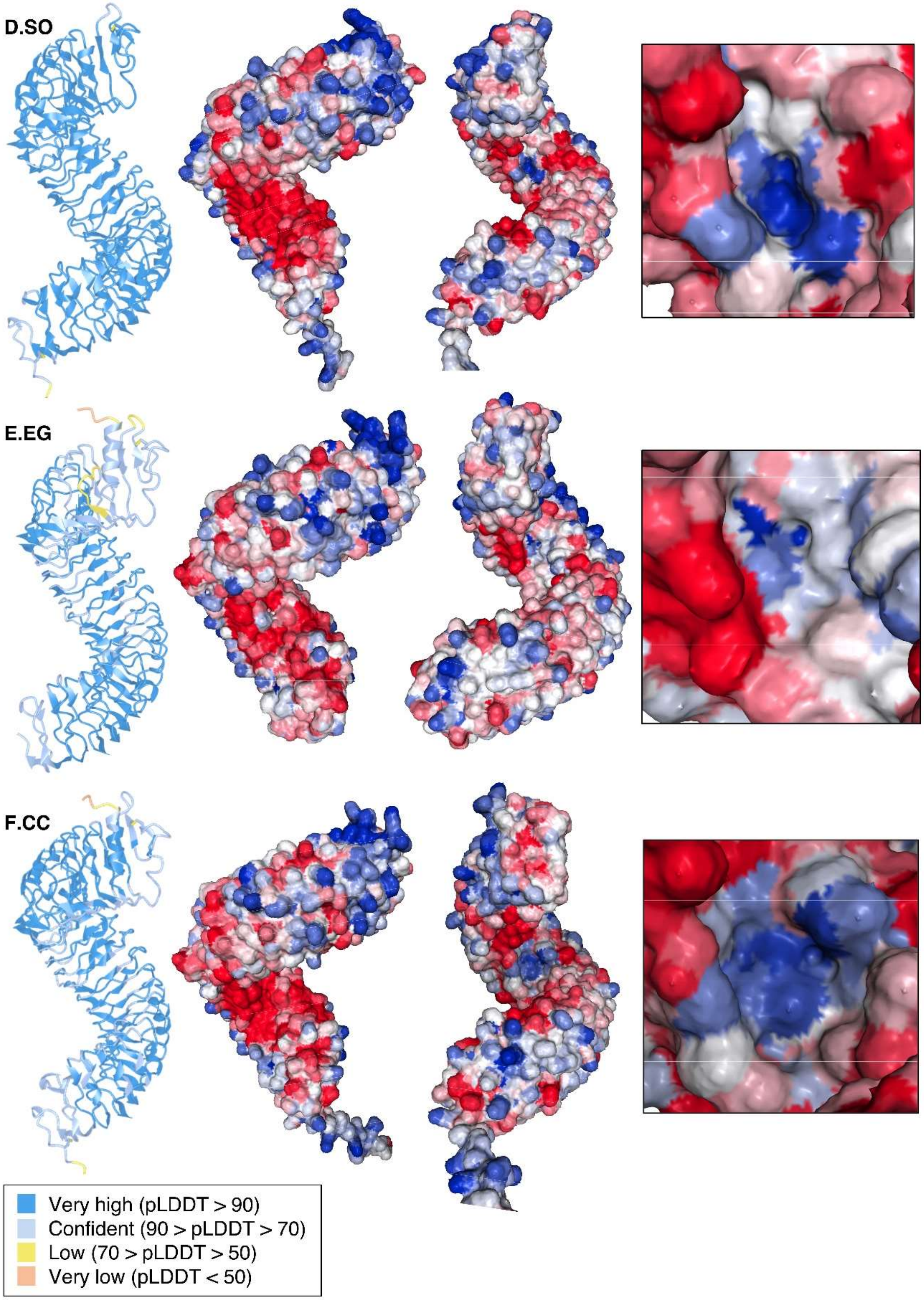
D.SO
E.EG
F.CC
Very high (pLDDT > 90)
Confident (90 > pLDDT > 70)
Low (70 > pLDDT > 50)
Very low (pLDDT < 50)

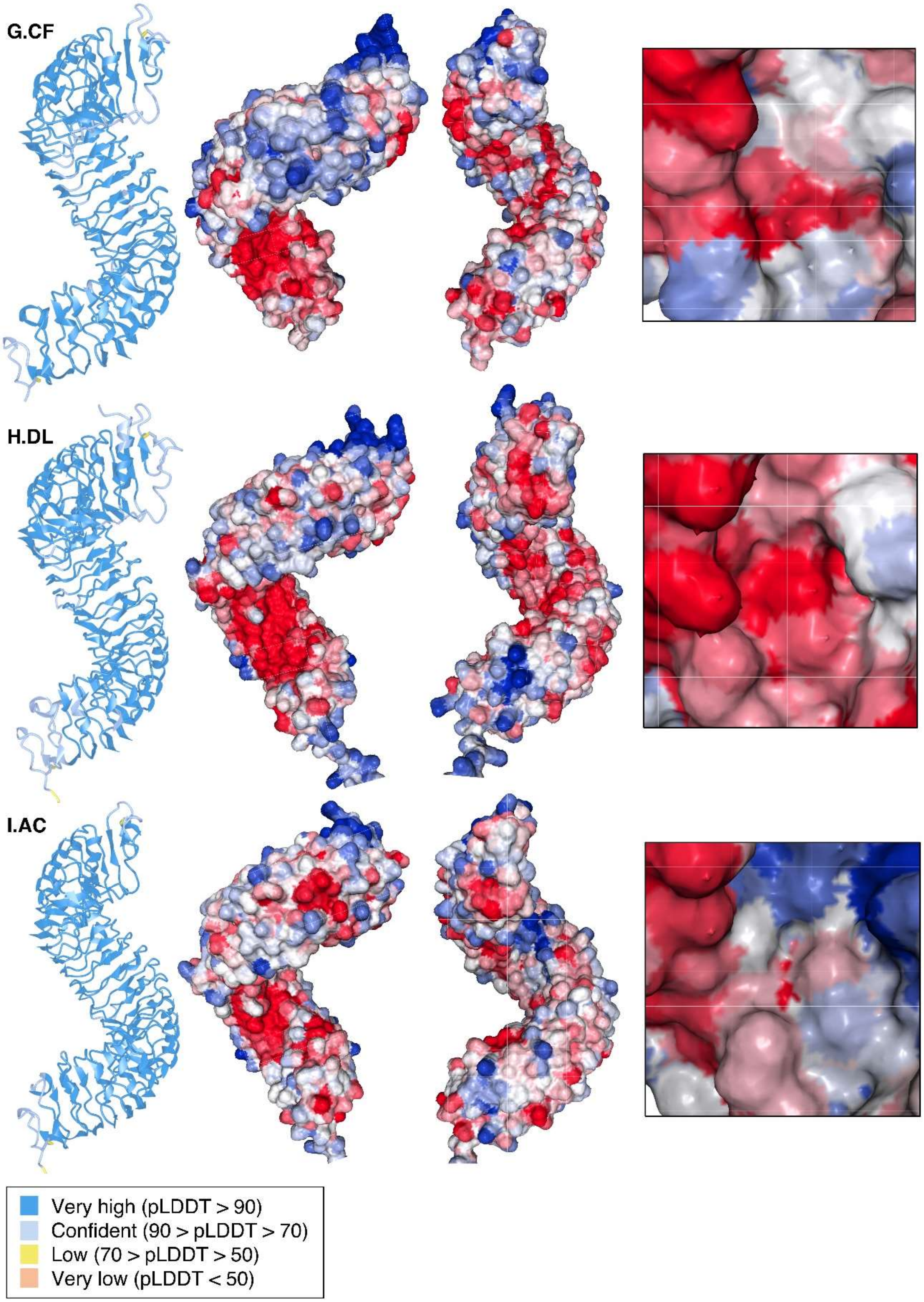
G.CF
H.DL
I.AC
Very high (pLDDT > 90)
Confident (90 > pLDDT > 70)
Low (70 > pLDDT > 50)
Very low (pLDDT < 50)

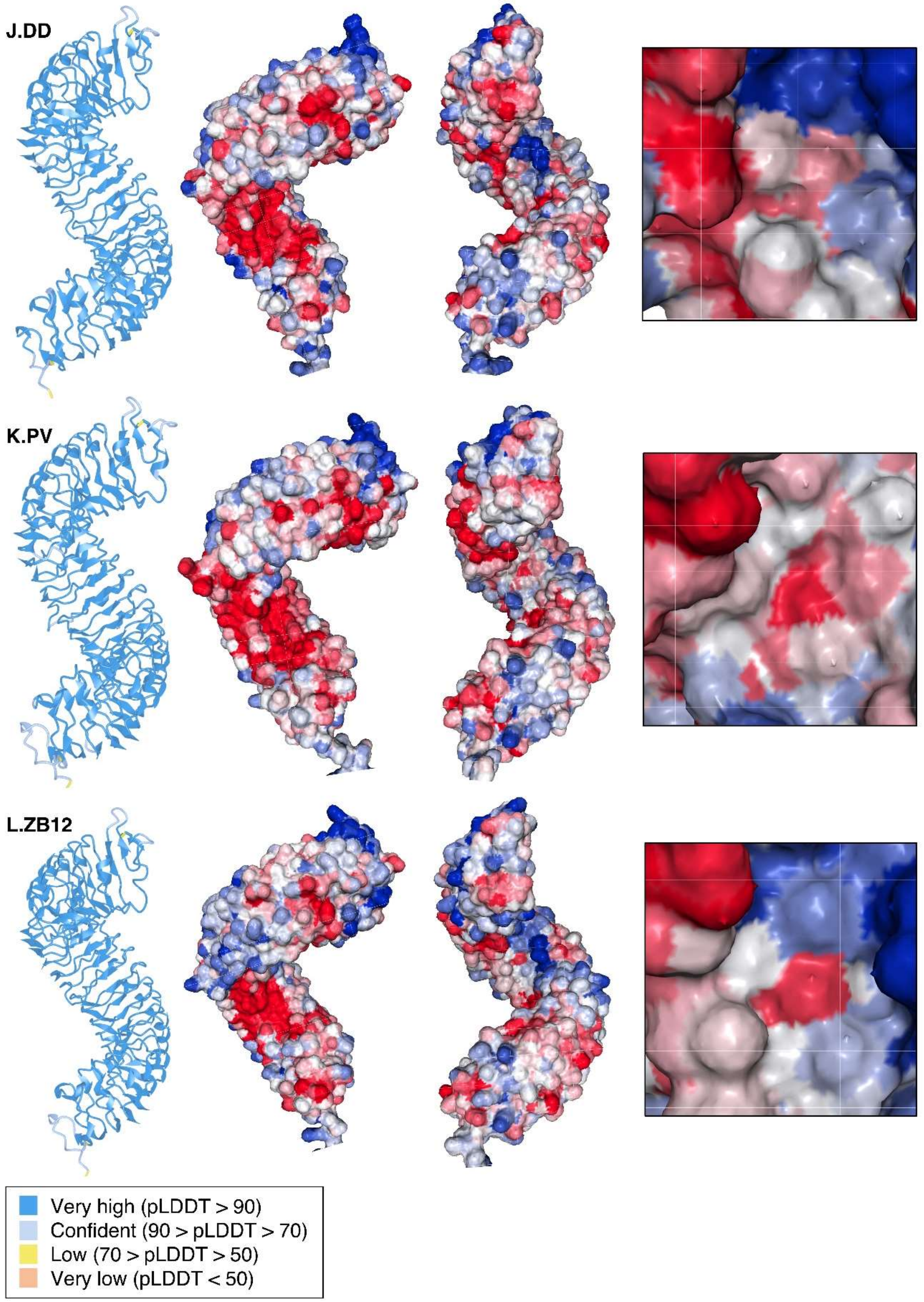
J.DD
K.PV
L.ZB12
Very high (pLDDT > 90)
Confident (90 > pLDDT > 70)
Low (70 > pLDDT > 50)
Very low (pLDDT < 50)

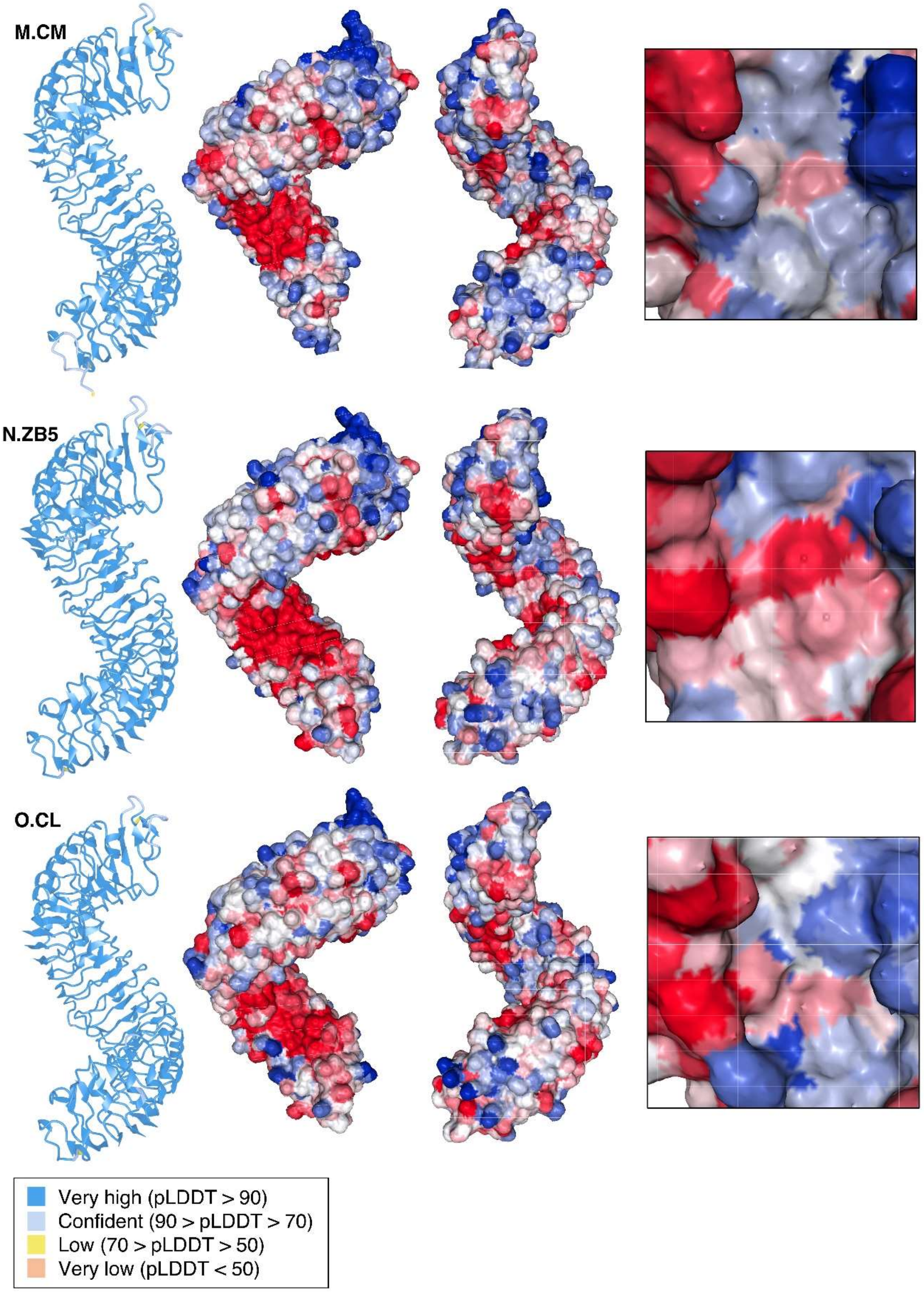
M.CM
N.ZB5
O.CL
Very high (pLDDT > 90)
Confident (90 > pLDDT > 70)
Low (70 > pLDDT > 50)
Very low (pLDDT < 50)

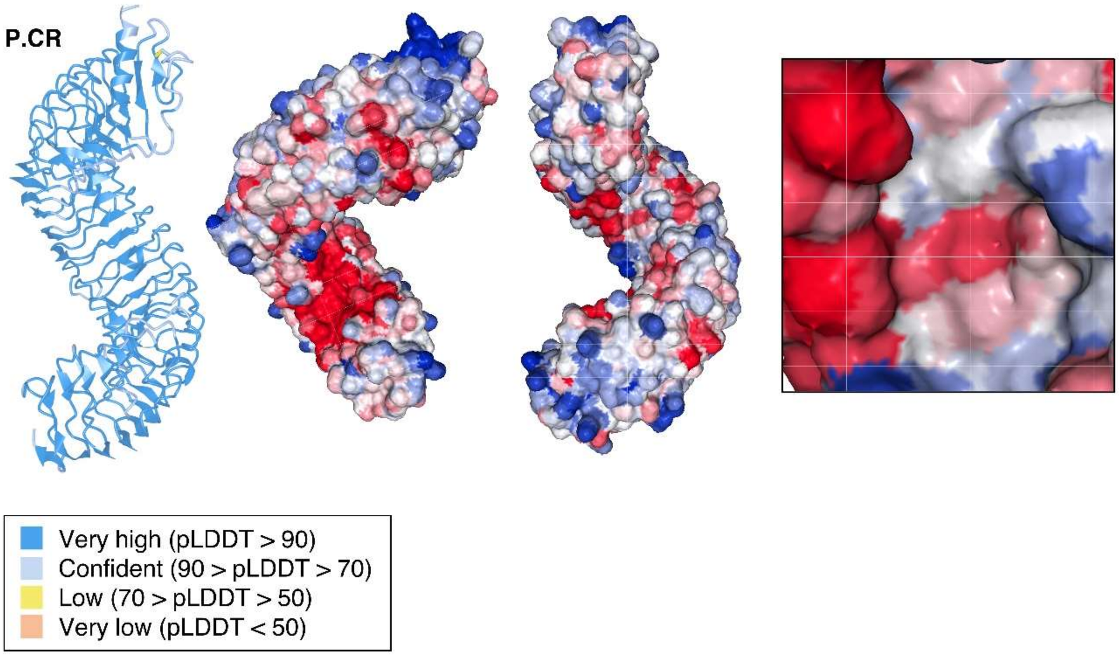


**Fig. S20. Alphafold3 structure prediction of SCORE orthologs and surface charge distribution.**

Alphafold3 structure prediction of SCORE$^{Ecto}$ from tested SCORE orthologs. Left structure: The predicted protein structure is color-coded based on pLDDT scores (see legend below). Middle structures: SCORE$^{Ecto}$ with surface charges displayed. Red indicates negative charge, blue indicates positive charge, and white indicates neutral charge. Right box: Surface charge distribution on the inner LRR surface between the 8$^{th}$-11$^{th}$ LRR motifs. (**A-P**) DW-CR refers to the original SCORE orthologs tested in Fig. 4D. Protein structures were visualized using iCn3D, and surface charges were calculated using protein-sol (24).

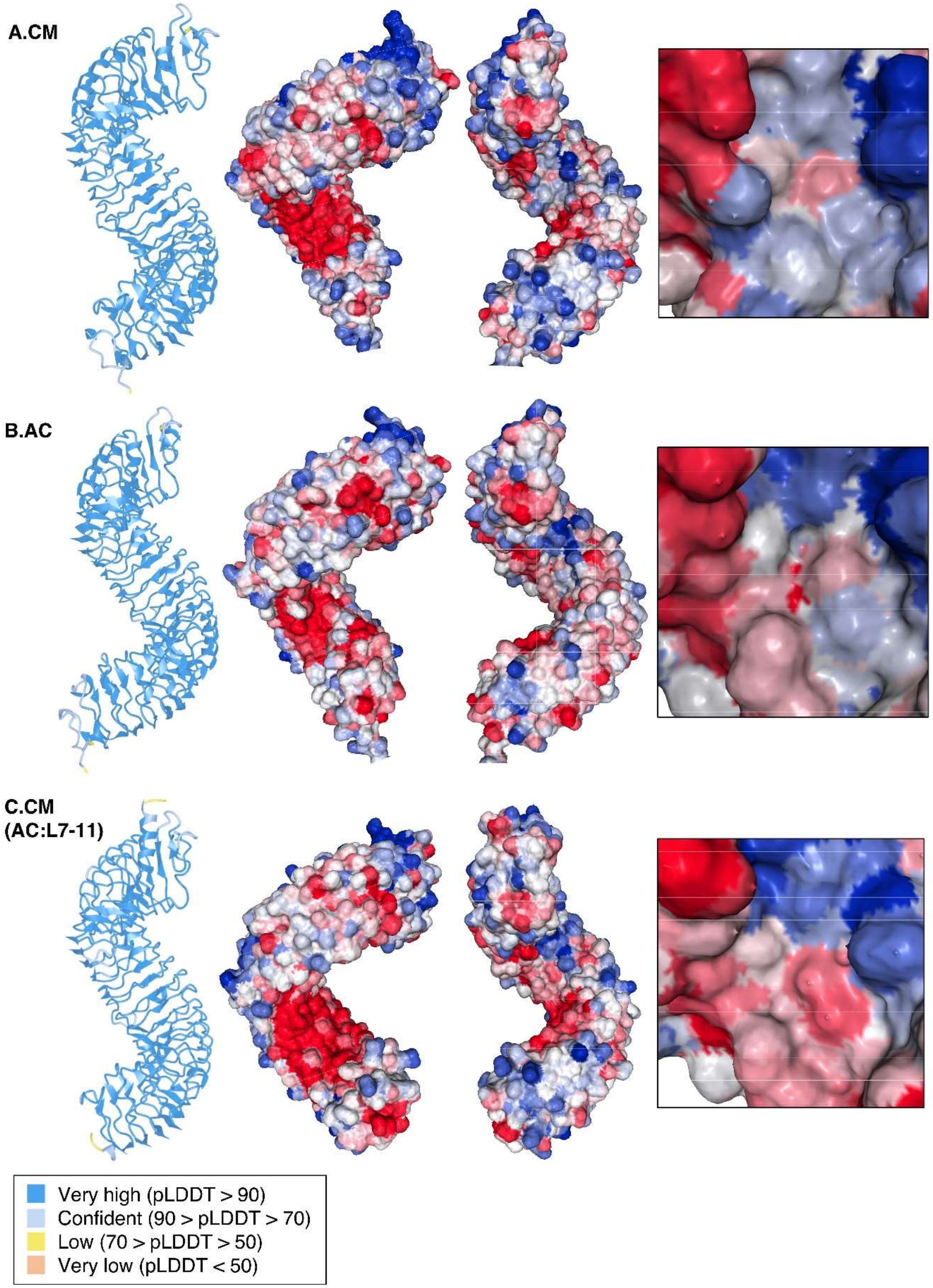
A.CM
B.AC
C.CM (AC:L7-11)
Very high (pLDDT > 90)
Confident (90 > pLDDT > 70)
Low (70 > pLDDT > 50)
Very low (pLDDT < 50)

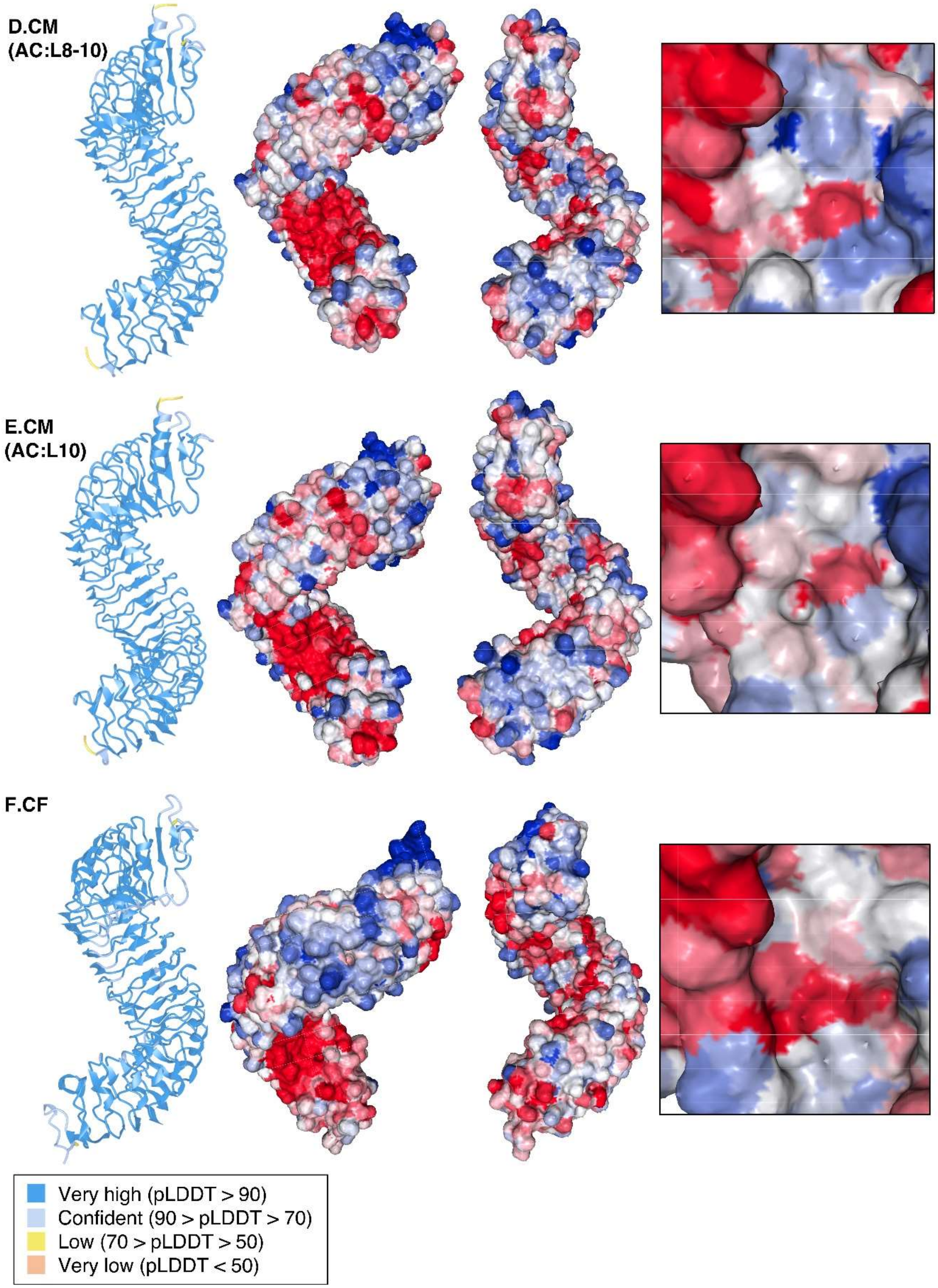
D.CM
(AC:L8-10)
E.CM
(AC:L10)
F.CF
Very high (pLDDT > 90)
Confident (90 > pLDDT > 70)
Low (70 > pLDDT > 50)
Very low (pLDDT < 50)

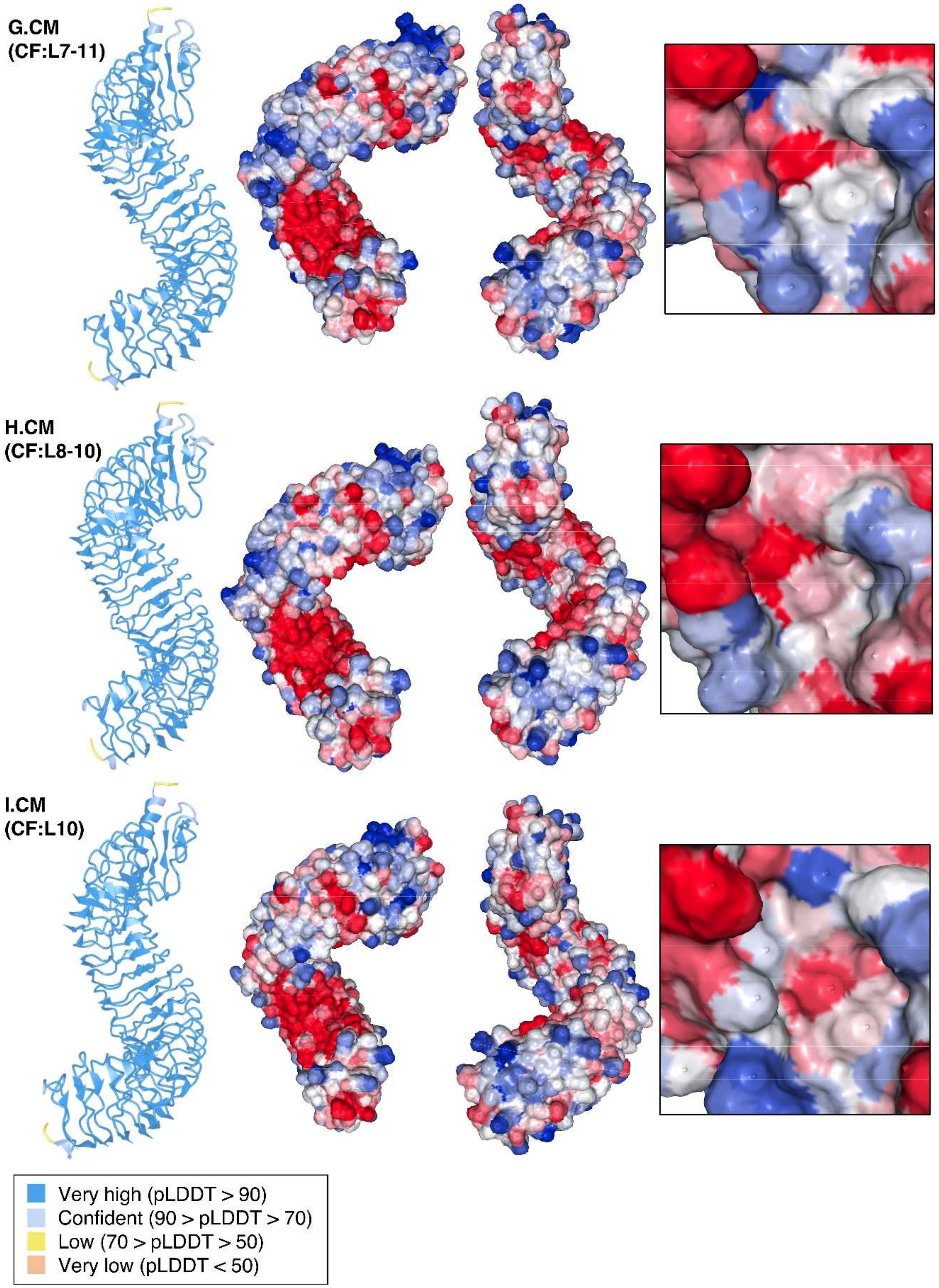
G.CM
(CF:L7-11)
H.CM
(CF:L8-10)
I.CM
(CF:L10)
Very high (pLDDT > 90)
Confident (90 > pLDDT > 70)
Low (70 > pLDDT > 50)
Very low (pLDDT < 50)

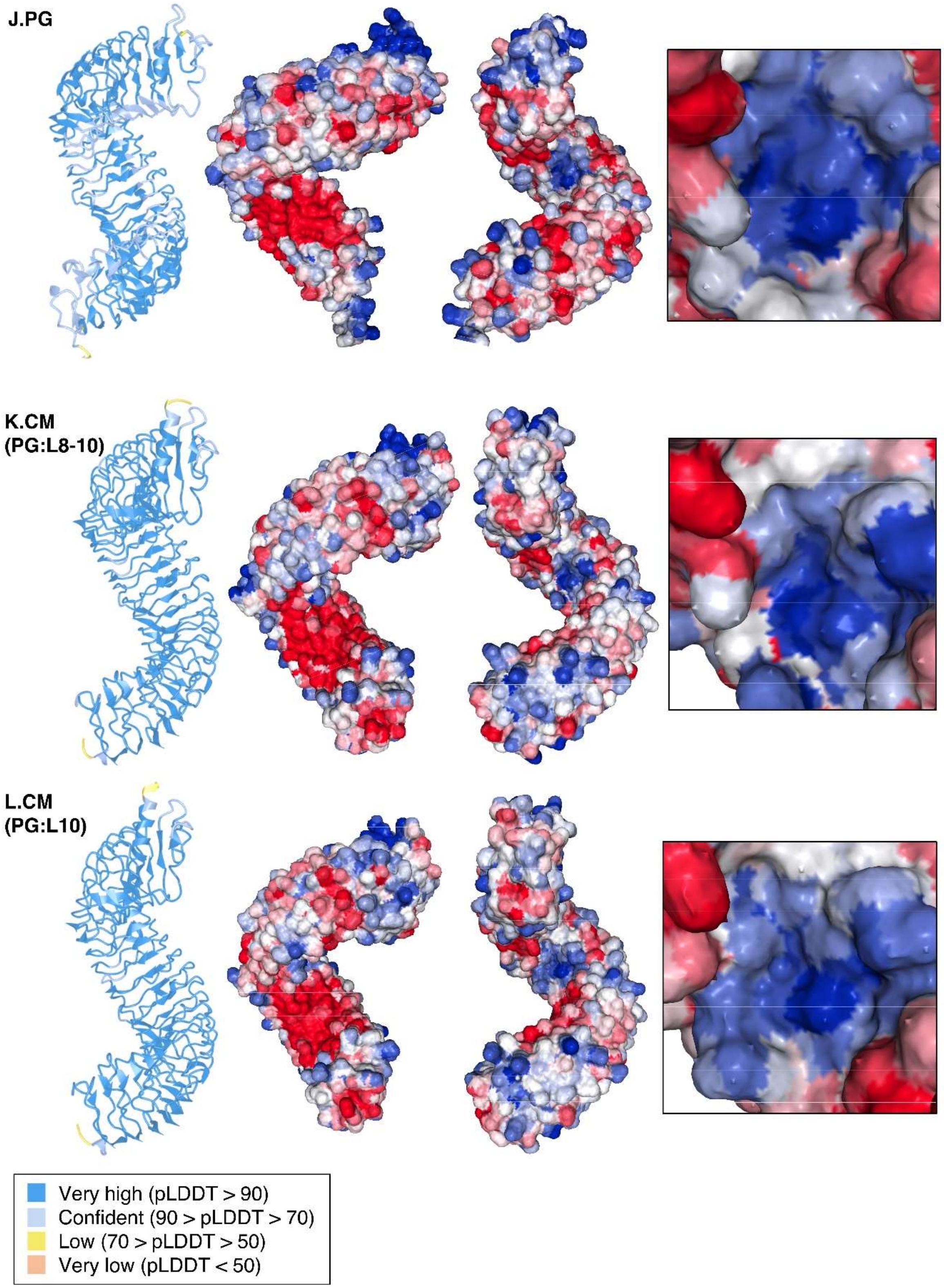
J.PG
K.CM
(PG:L8-10)
L.CM
(PG:L10)
Very high (pLDDT > 90)
Confident (90 > pLDDT > 70)
Low (70 > pLDDT > 50)
Very low (pLDDT < 50)

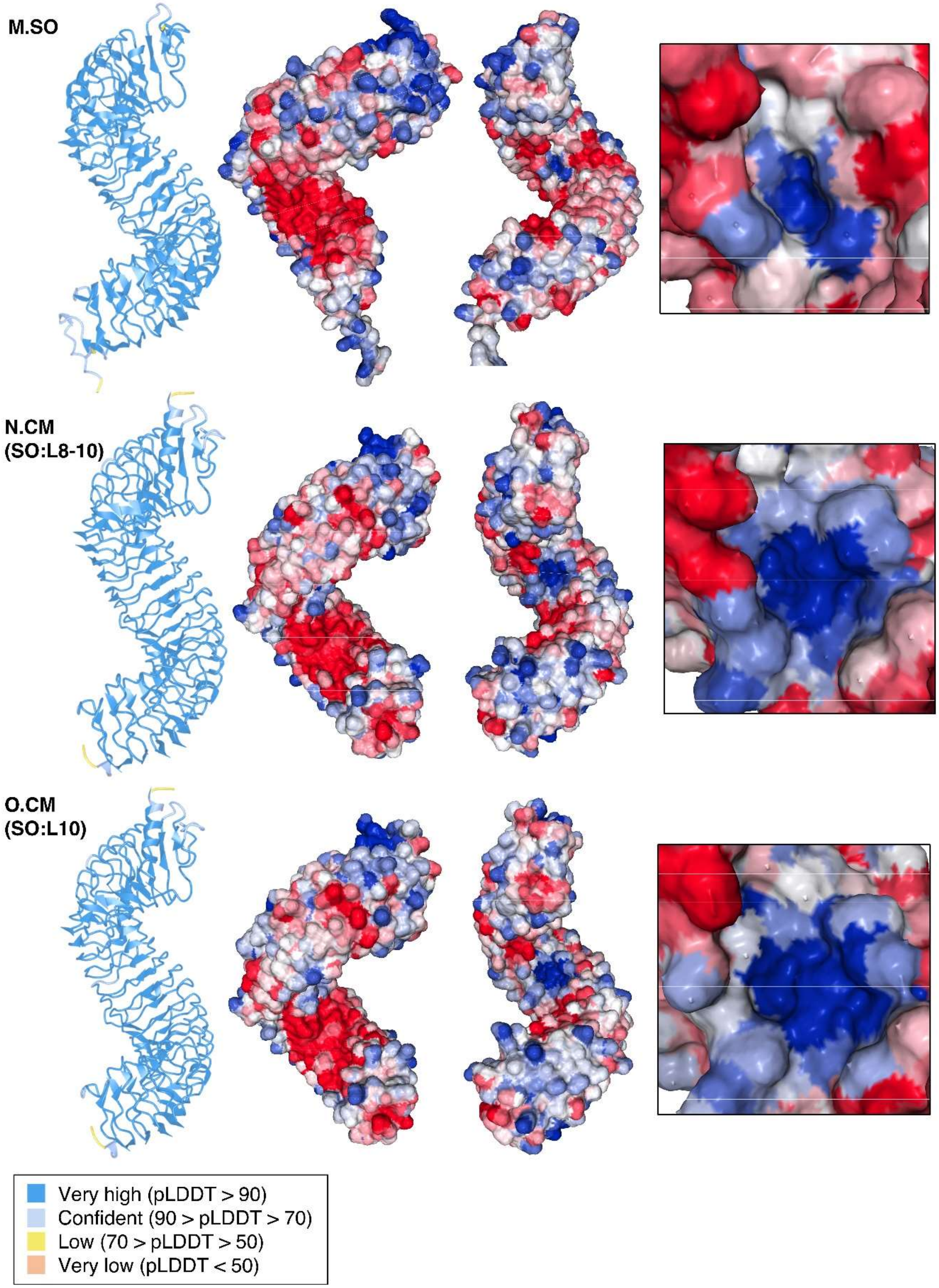
M.SO
N.CM
(SO:L8-10)
O.CM
(SO:L10)
Very high (pLDDT > 90)
Confident (90 > pLDDT > 70)
Low (70 > pLDDT > 50)
Very low (pLDDT < 50)

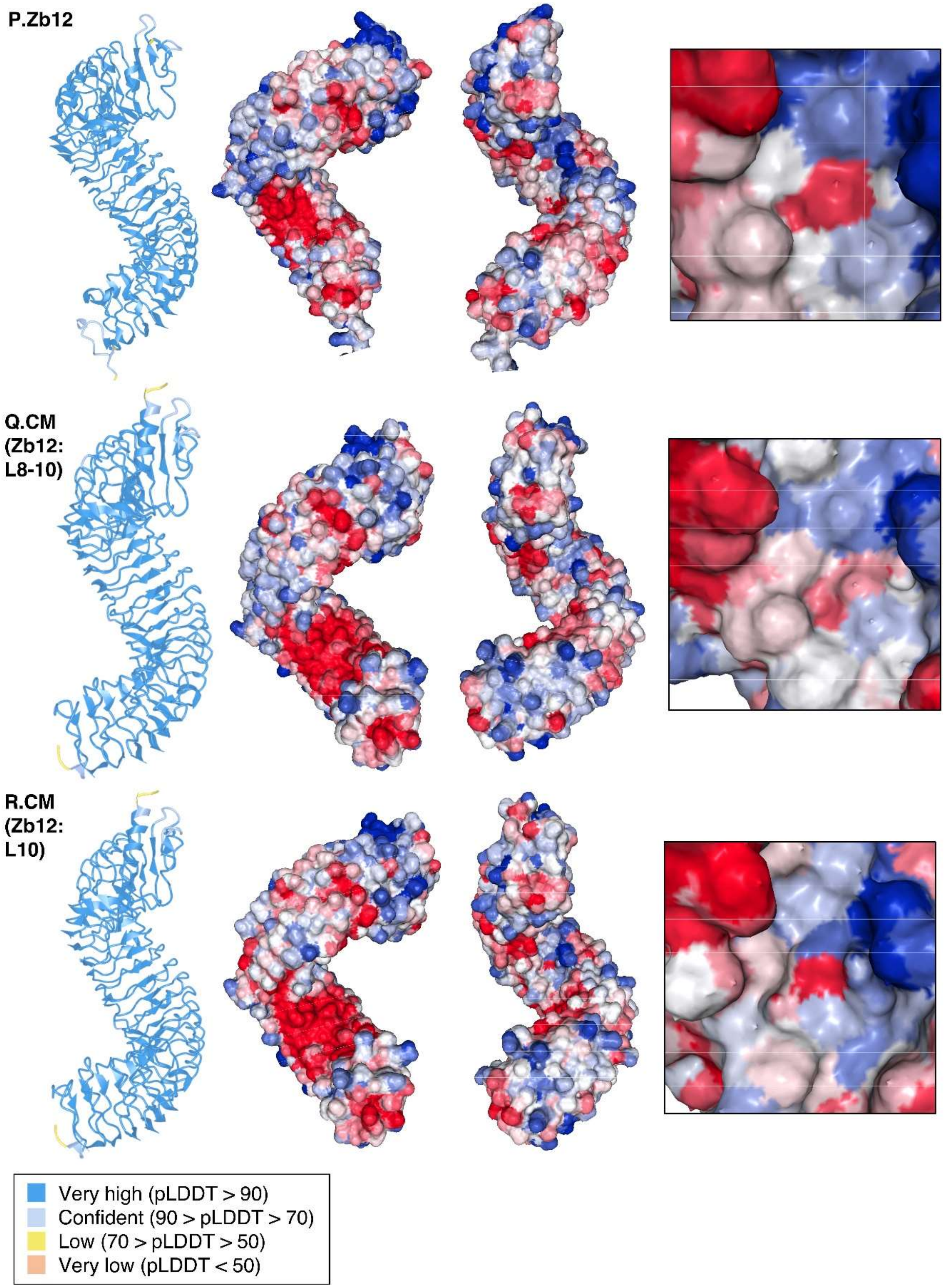

P.Zb12
Q.CM (Zb12: L8-10)
R.CM (Zb12: L10)
Very high (pLDDT > 90)
Confident (90 > pLDDT > 70)
Low (70 > pLDDT > 50)
Very low (pLDDT < 50)

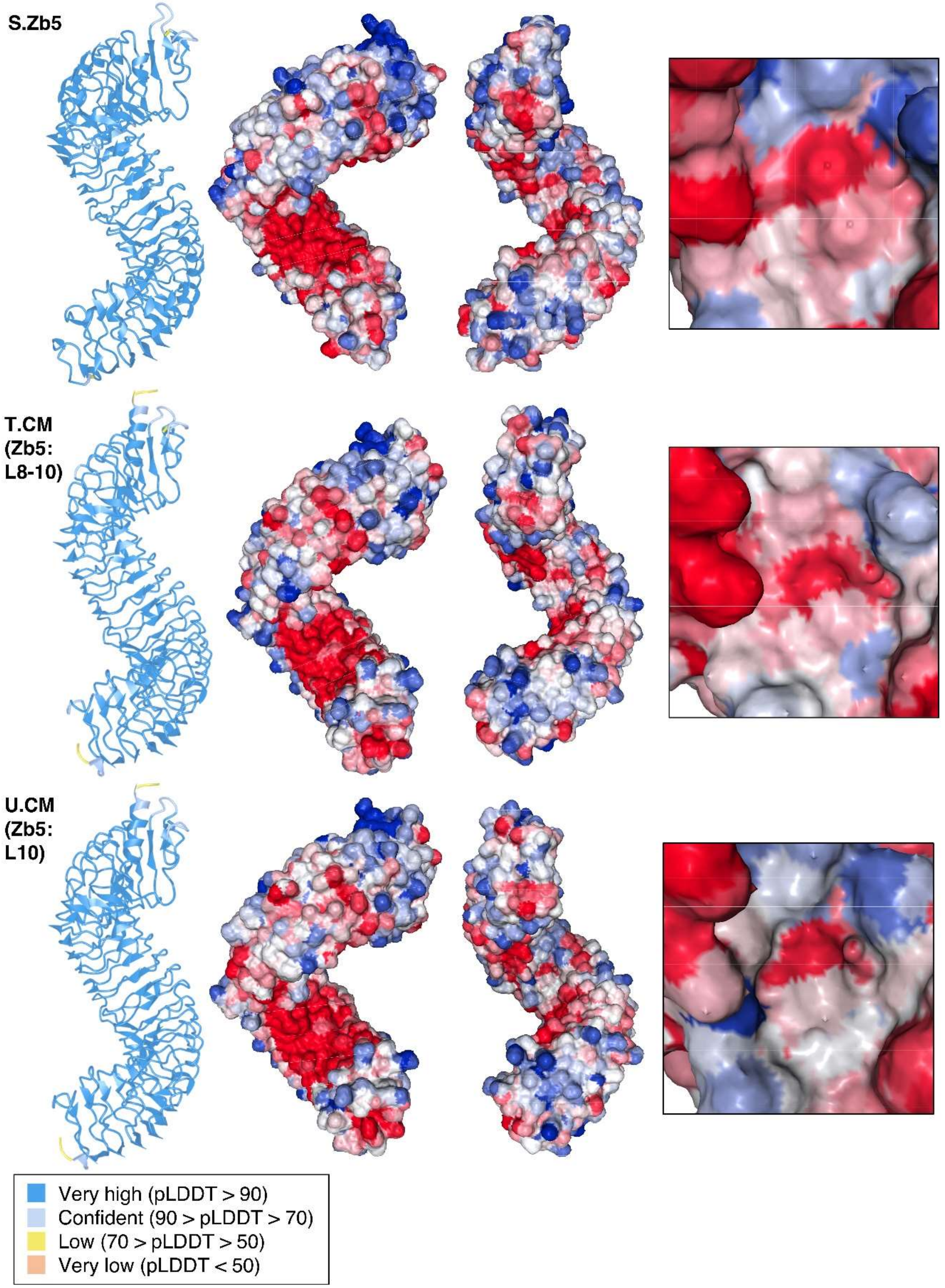
S.Zb5
T.CM (Zb5: L8-10)
U.CM (Zb5: L10)
Very high (pLDDT > 90)
Confident (90 > pLDDT > 70)
Low (70 > pLDDT > 50)
Very low (pLDDT < 50)

**Fig. S21. Alphafold3 structure prediction and surface charge distribution of SCORE orthologs and variants (CM) generated from domain swaps.**

Left structure: The predicted protein structure is color-coded based on pLDDT scores (see legend below). Middle structures: SCORE$^{Ecto}$ with surface charges displayed. Red indicates negative charge, blue indicates positive charge, and white indicates neutral charge. Right box: Surface charge distribution on the inner LRR surface between the 8$^{th}$-11$^{th}$ LRR motifs. For label of receptors, CM, AC, CF, SO, PG, ZB5 and ZB12 refers to the original SCORE orthologs tested in Fig. 4D. CM with different LRR motifs from other orthologs are indicated from (**A-U**). For example, CM(AC:L7-11) refers to CM swapped with the 7$^{th}$-11$^{th}$ LRR motifs from AC (also see Figure 5D). Protein structures were visualized using iCn3D, and surface charges were calculated using protein-sol (24).

**A**

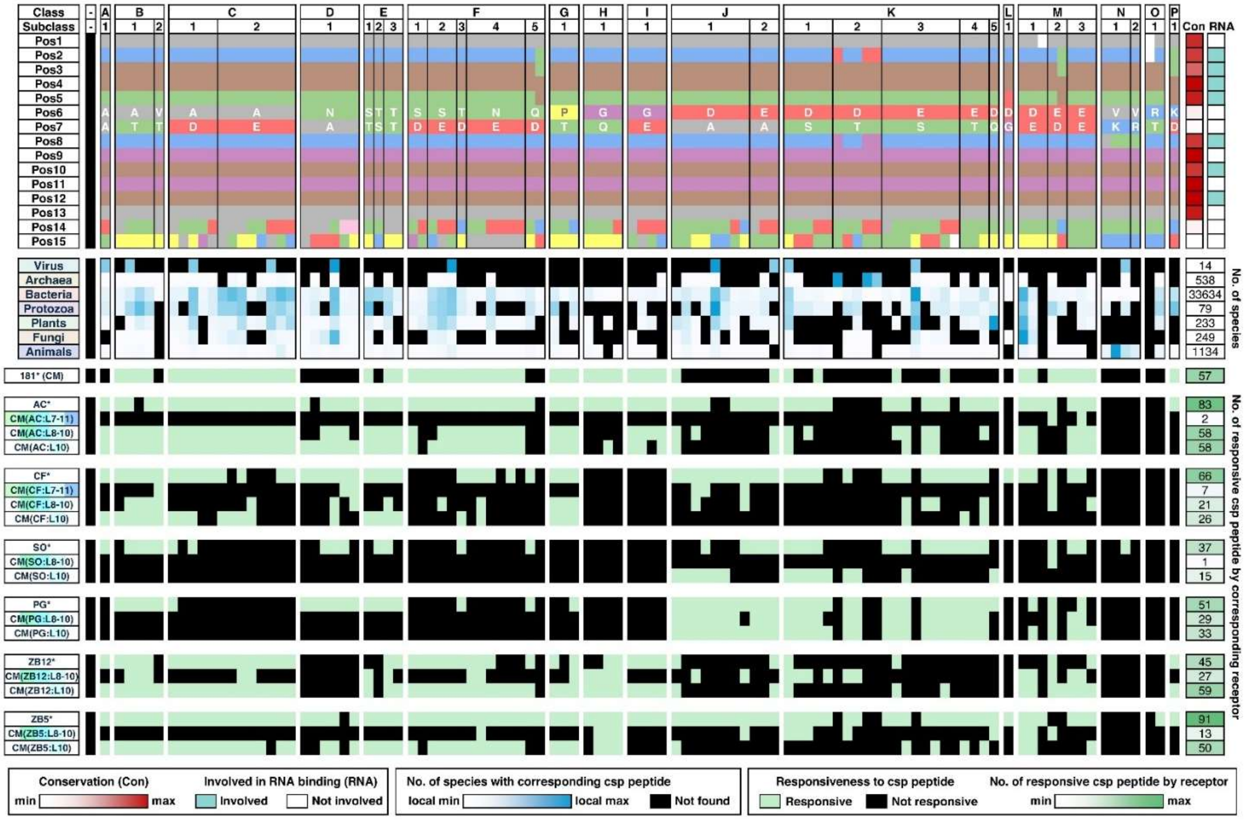
Class
Subclass
Con
RNA
Virus
Archaea
Bacteria
Protozoa
Plants
Fungi
Animals
No. of species
181* (CM)
AC*
CM(AC:L7-11)
CM(AC:L8-10)
CM(AC:L10)
CF*
CM(CF:L7-11)
CM(CF:L8-10)
CM(CF:L10)
SO*
CM(SO:L8-10)
CM(SO:L10)
PG*
CM(PG:L8-10)
CM(PG:L10)
ZB12*
CM(ZB12:L8-10)
CM(ZB12:L10)
ZB5*
CM(ZB5:L8-10)
CM(ZB5:L10)
No. of responsive csp peptide by corresponding receptor
Conservation (Con)
min
max
Involved in RNA binding (RNA)
Involved
Not involved
No. of species with corresponding csp peptide
local min
local max
Not found
Responsiveness to csp peptide
Responsive
Not responsive
No. of responsive csp peptide by receptor
min
max

B

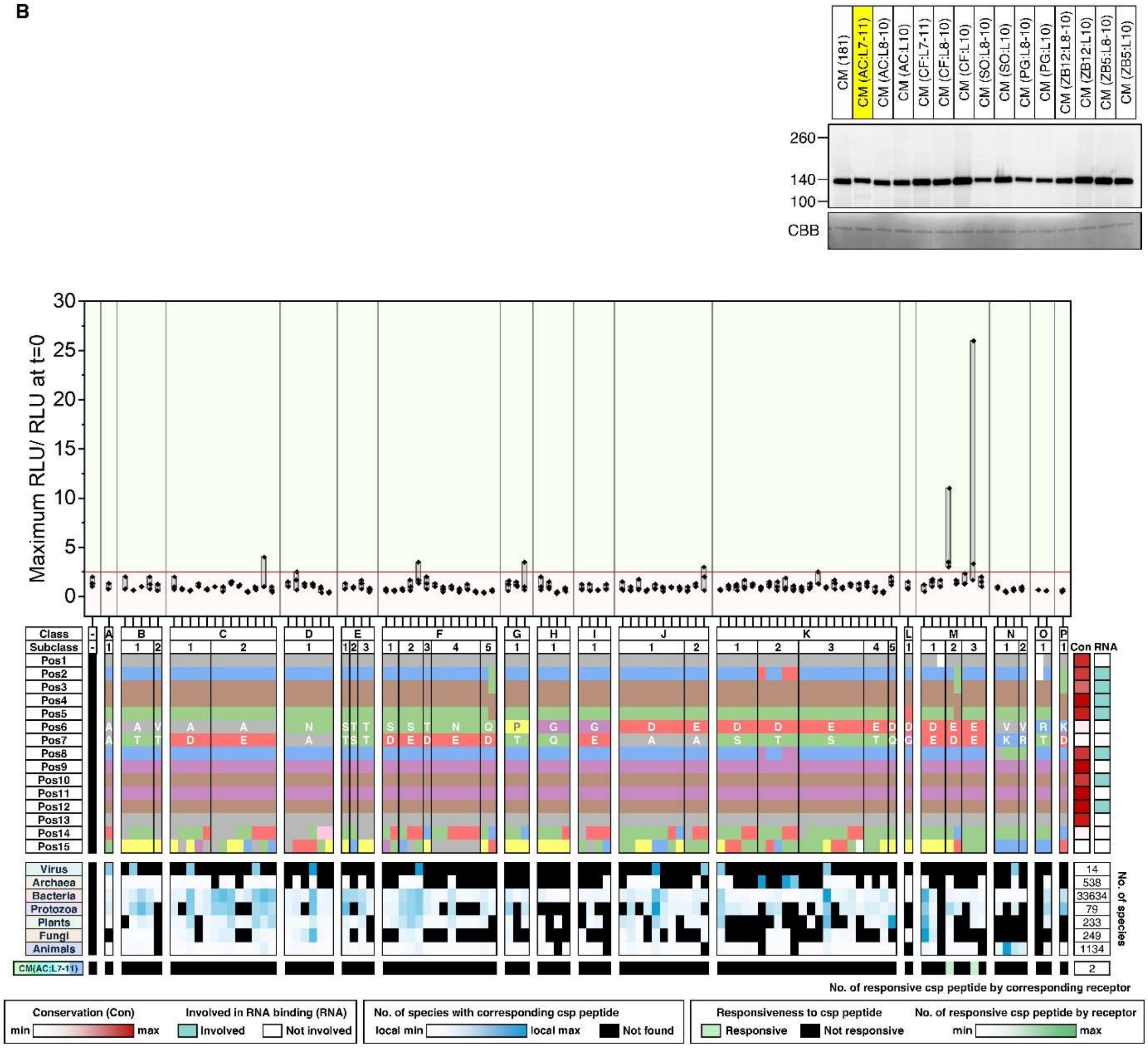
CM (181)
CM (AC:L7-11)
CM (AC:L8-10)
CM (AC:L10)
CM (CF:L7-11)
CM (CF:L8-10)
CM (CF:L10)
CM (SO:L8-10)
CM (SO:L10)
CM (PG:L8-10)
CM (PG:L10)
CM (ZB12:L8-10)
CM (ZB12:L10)
CM (ZB5:L8-10)
CM (ZB5:L10)
260
140
100
CBB
Maximum RLU/ RLU at t=0
Class
Subclass
Virus
Archaea
Bacteria
Protozoa
Plants
Fungi
Animals
CM(AC:L7-11)
Con RNA
No. of species
14
538
33634
79
233
249
1134
2
No. of responsive csp peptide by corresponding receptor
Conservation (Con)
Involved in RNA binding (RNA)
min
max
Involved
Not involved
No. of species with corresponding csp peptide
local min
local max
Not found
Responsiveness to csp peptide
No. of responsive csp peptide by receptor
Responsive
Not responsive

C

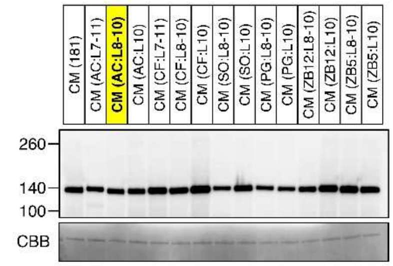

CM (181)
CM (AC:L7-11)
CM (AC:L8-10)
CM (AC:L10)
CM (CF:L7-11)
CM (CF:L8-10)
CM (CF:L10)
CM (SO:L8-10)
CM (SO:L10)
CM (PG:L8-10)
CM (PG:L10)
CM (ZB12:L8-10)
CM (ZB12:L10)
CM (ZB5:L8-10)
CM (ZB5:L10)
260
140
100
CBB


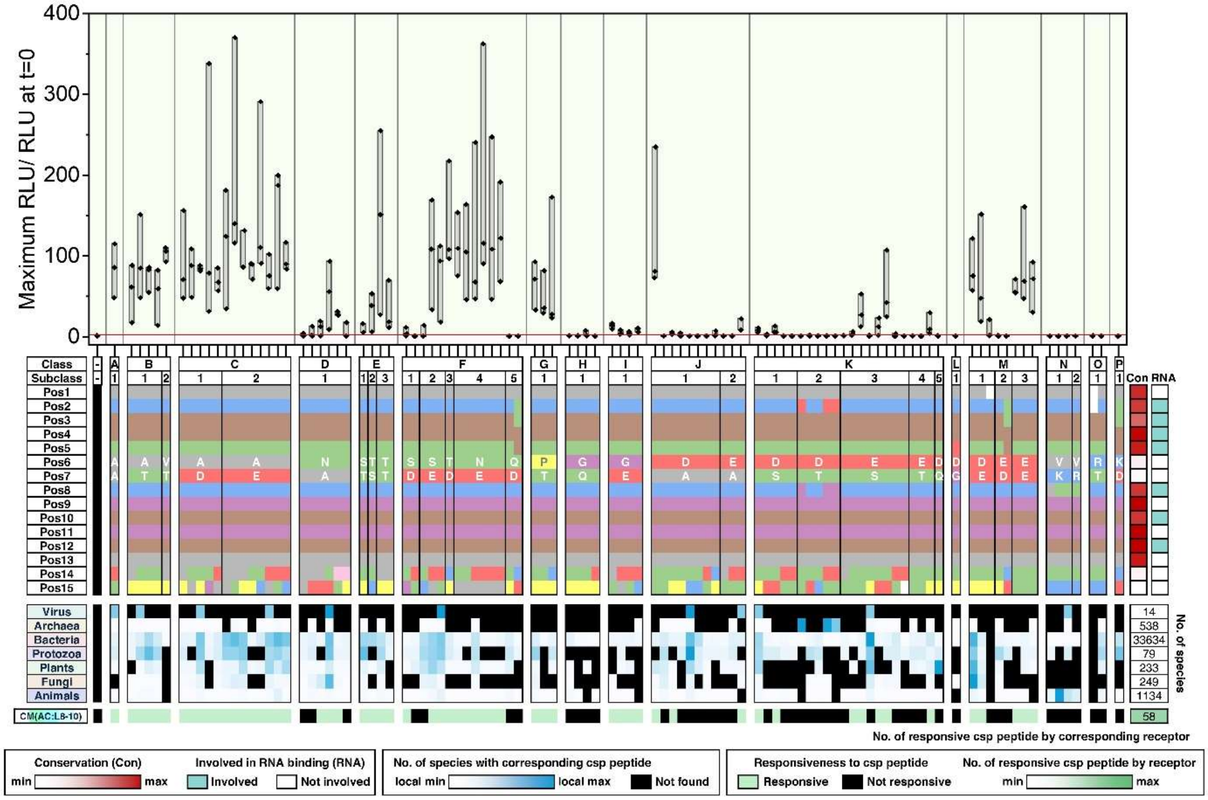

Maximum RLU/ RLU at t=0
400
300
200
100
0
Class
Subclass
Pos1
Pos2
Pos3
Pos4
Pos5
Pos6
Pos7
Pos8
Pos9
Pos10
Pos11
Pos12
Pos13
Pos14
Pos15
Con
RNA
Virus
Archaea
Bacteria
Protozoa
Plants
Fungi
Animals
No. of species
14
538
33634
79
233
249
1134
CM(AC:L8-10)
58
No. of responsive csp peptide by corresponding receptor
Conservation (Con)
min
max
Involved in RNA binding (RNA)
Involved
Not involved
No. of species with corresponding csp peptide
local min
local max
Not found
Responsiveness to csp peptide
Responsive
Not responsive
No. of responsive csp peptide by receptor
min
max

**D**

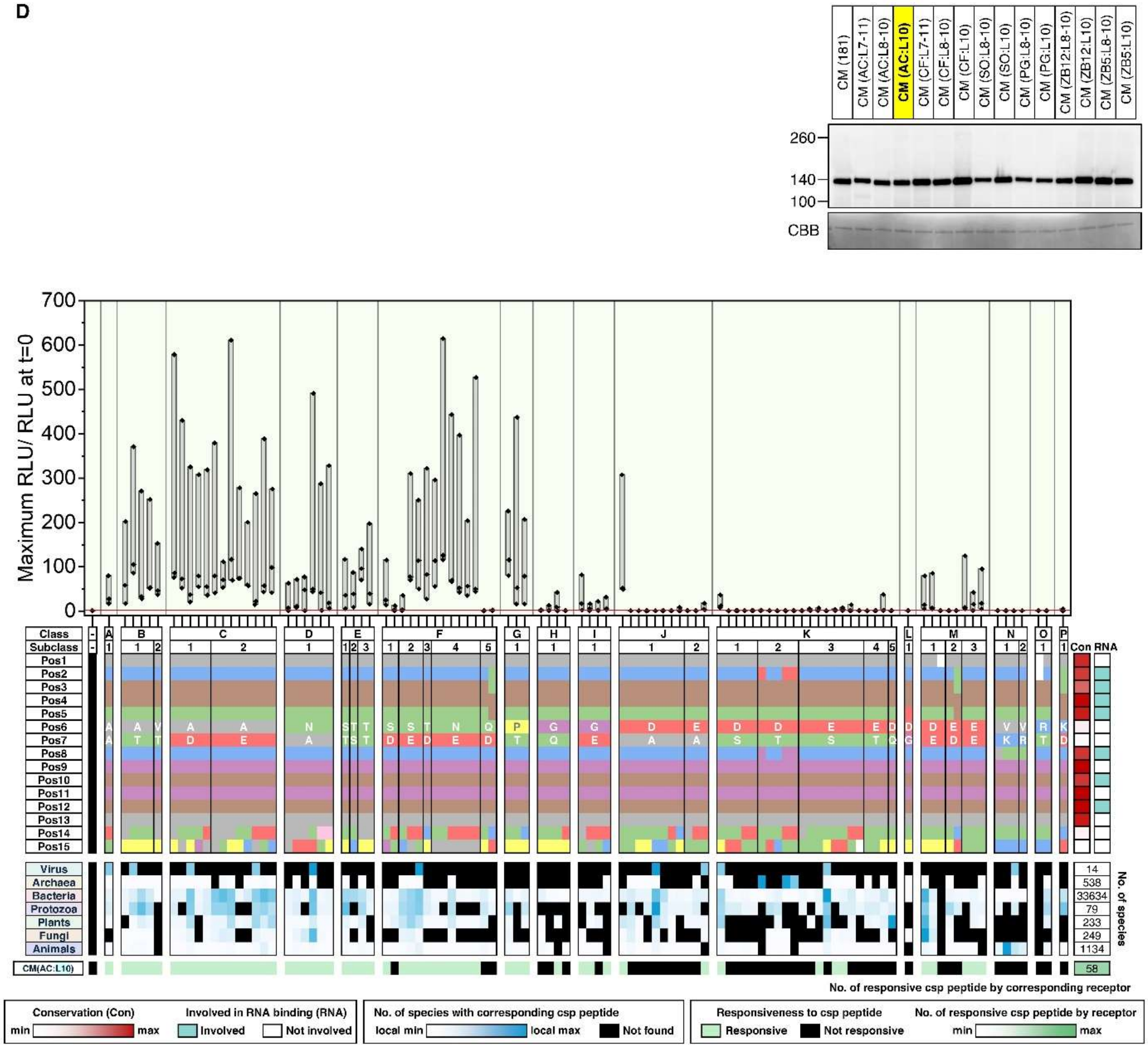
CM (181)
CM (AC:L7-11)
CM (AC:L8-10)
CM (AC:L10)
CM (CF:L7-11)
CM (CF:L8-10)
CM (CF:L10)
CM (SO:L8-10)
CM (SO:L10)
CM (PG:L8-10)
CM (PG:L10)
CM (ZB12:L8-10)
CM (ZB12:L10)
CM (ZB5:L8-10)
CM (ZB5:L10)
260
140
100
CBB
Maximum RLU/ RLU at t=0
700
600
500
400
300
200
100
0
Class
Subclass
Pos1
Pos2
Pos3
Pos4
Pos5
Pos6
Pos7
Pos8
Pos9
Pos10
Pos11
Pos12
Pos13
Pos14
Pos15
Con
RNA
Virus
Archaea
Bacteria
Protozoa
Plants
Fungi
Animals
14
538
33634
79
233
249
1134
No. of species
CM(AC:L10)
58
No. of responsive csp peptide by corresponding receptor
Conservation (Con)
min
max
Involved in RNA binding (RNA)
Involved
Not involved
No. of species with corresponding csp peptide
local min
local max
Not found
Responsiveness to csp peptide
Responsive
Not responsive
No. of responsive csp peptide by receptor
min
max

E

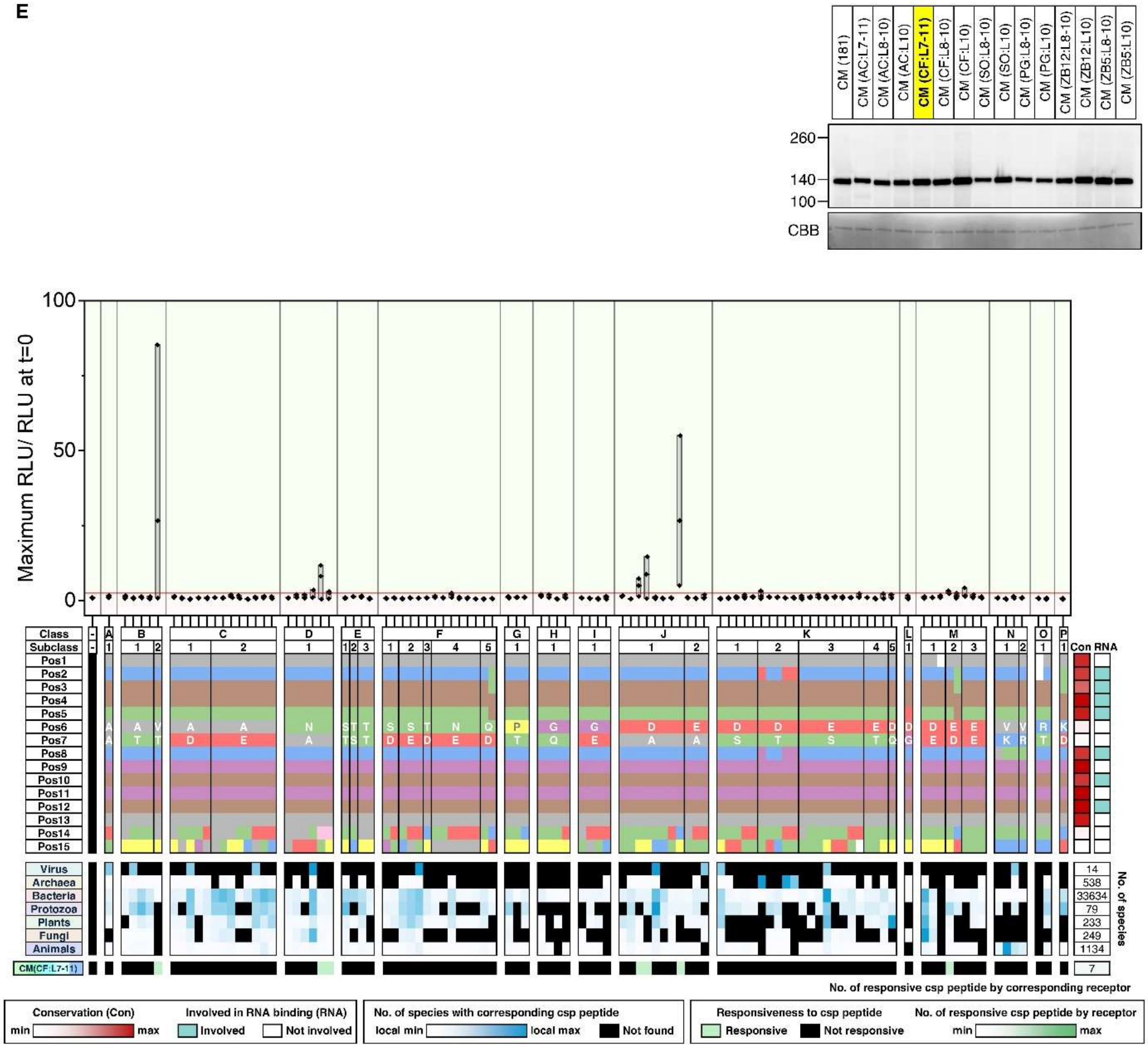

CM (181)
CM (AC:L7-11)
CM (AC:L8-10)
CM (AC:L10)
CM (CF:L7-11)
CM (CF:L8-10)
CM (CF:L10)
CM (SO:L8-10)
CM (SO:L10)
CM (PG:L8-10)
CM (PG:L10)
CM (ZB12:L8-10)
CM (ZB12:L10)
CM (ZB5:L8-10)
CM (ZB5:L10)
260
140
100
CBB
Maximum RLU/ RLU at t=0
100
50
0
Class
Subclass
Pos1
Pos2
Pos3
Pos4
Pos5
Pos6
Pos7
Pos8
Pos9
Pos10
Pos11
Pos12
Pos13
Pos14
Pos15
Con
RNA
Virus
Archaea
Bacteria
Protozoa
Plants
Fungi
Animals
No. of species
14
538
33634
79
233
249
1134
CM(CF:L7-11)
7
No. of responsive csp peptide by corresponding receptor
Conservation (Con)
min
max
Involved in RNA binding (RNA)
Involved
Not involved
No. of species with corresponding csp peptide
local min
local max
Not found
Responsiveness to csp peptide
Responsive
Not responsive
No. of responsive csp peptide by receptor
min
max

F

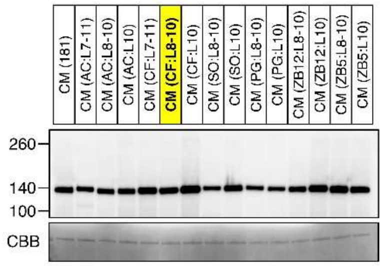
CM (181)
CM (AC:L7-11)
CM (AC:L8-10)
CM (AC:L10)
CM (CF:L7-11)
CM (CF:L8-10)
CM (CF:L10)
CM (SO:L8-10)
CM (SO:L10)
CM (PG:L8-10)
CM (PG:L10)
CM (ZB12:L8-10)
CM (ZB12:L10)
CM (ZB5:L8-10)
CM (ZB5:L10)
260
140
100
CBB

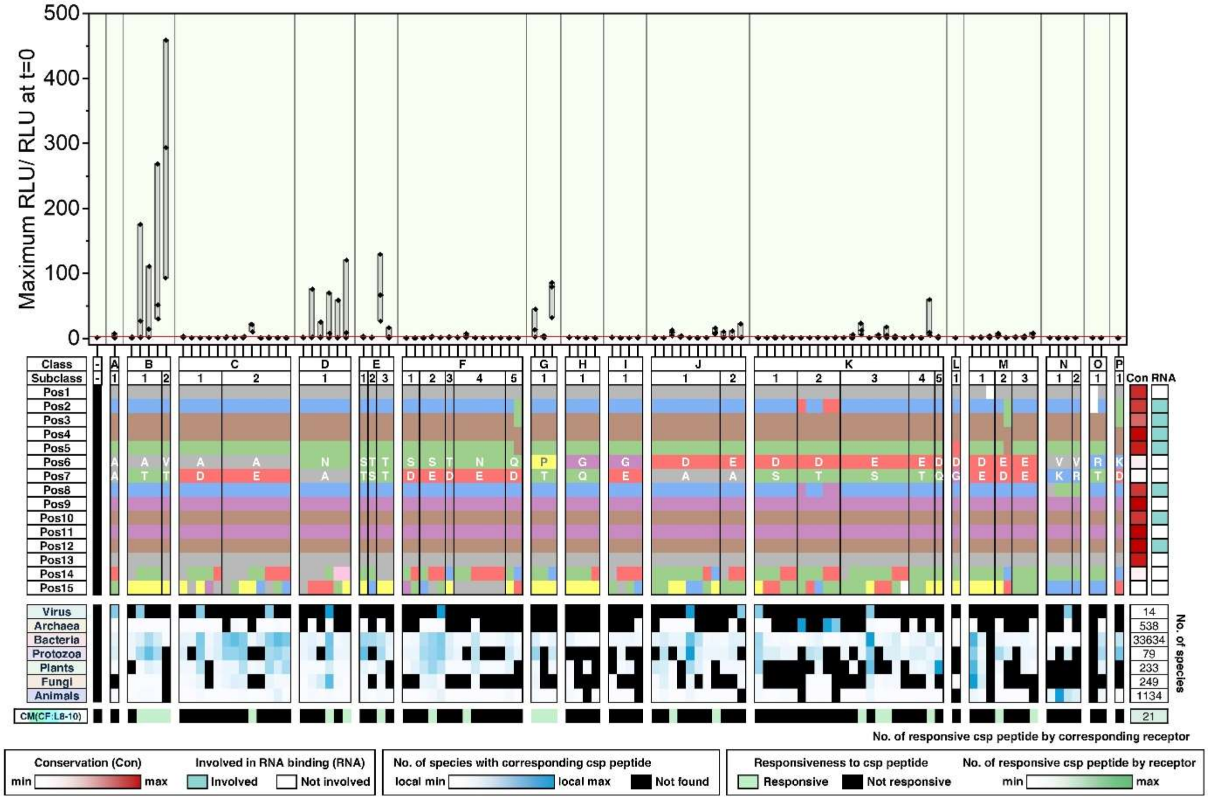
Maximum RLU/ RLU at t=0
500
400
300
200
100
0
Class
Subclass
Pos1
Pos2
Pos3
Pos4
Pos5
Pos6
Pos7
Pos8
Pos9
Pos10
Pos11
Pos12
Pos13
Pos14
Pos15
Con
RNA
Virus
Archaea
Bacteria
Protozoa
Plants
Fungi
Animals
No. of species
14
538
33634
79
233
249
1134
CM(CF:L8-10)
21
No. of responsive csp peptide by corresponding receptor
Conservation (Con)
min
max
Involved in RNA binding (RNA)
Involved
Not involved
No. of species with corresponding csp peptide
local min
local max
Not found
Responsiveness to csp peptide
Responsive
Not responsive
No. of responsive csp peptide by receptor
min
max

G

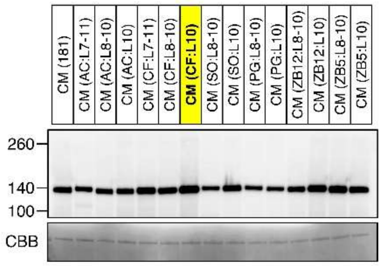
CM (181)
CM (AC:L7-11)
CM (AC:L8-10)
CM (AC:L10)
CM (CF:L7-11)
CM (CF:L8-10)
CM (CF:L10)
CM (SO:L8-10)
CM (SO:L10)
CM (PG:L8-10)
CM (PG:L10)
CM (ZB12:L8-10)
CM (ZB12:L10)
CM (ZB5:L8-10)
CM (ZB5:L10)
260
140
100
CBB

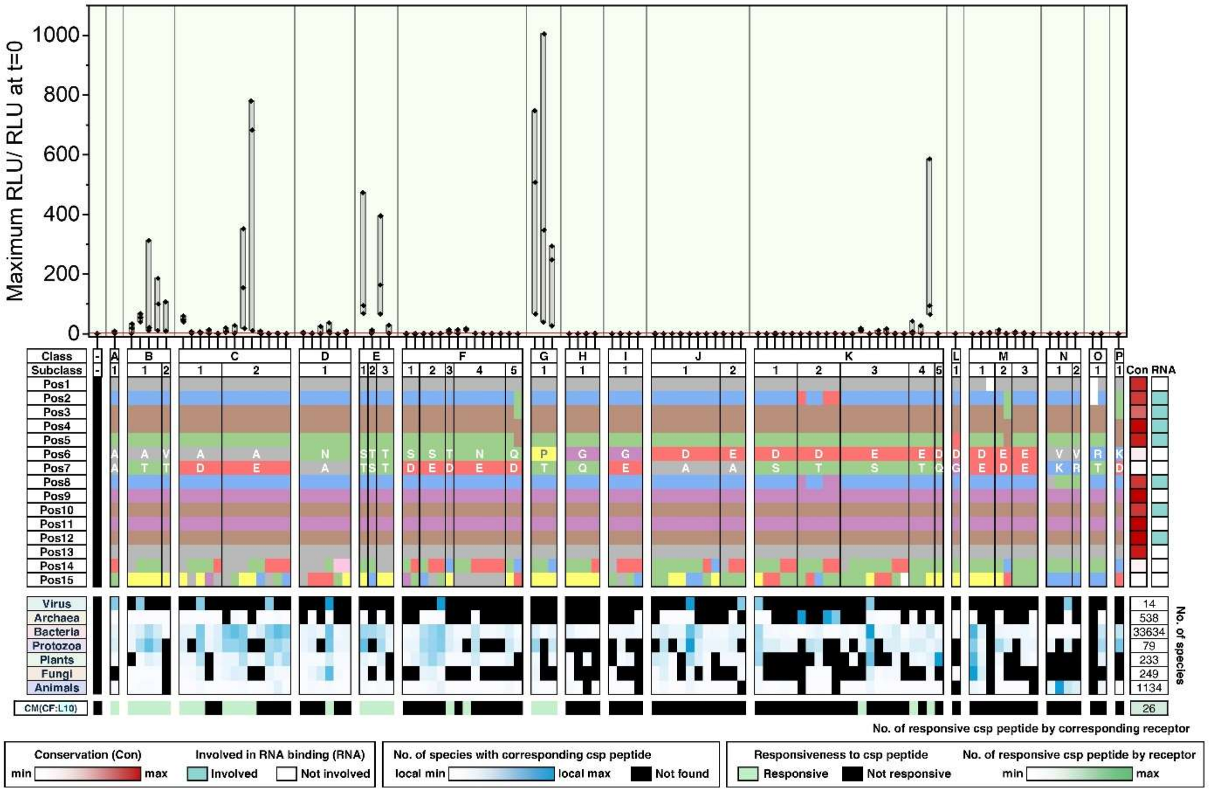
Maximum RLU/ RLU at t=0
1000
800
600
400
200
0
Class
Subclass
Pos1
Pos2
Pos3
Pos4
Pos5
Pos6
Pos7
Pos8
Pos9
Pos10
Pos11
Pos12
Pos13
Pos14
Pos15
Con
RNA
Virus
Archaea
Bacteria
Protozoa
Plants
Fungi
Animals
14
538
33634
79
233
249
1134
No. of species
CM(CF:L10)
26
No. of responsive csp peptide by corresponding receptor
Conservation (Con)
min
max
Involved in RNA binding (RNA)
Involved
Not involved
No. of species with corresponding csp peptide
local min
local max
Not found
Responsiveness to csp peptide
Responsive
Not responsive
No. of responsive csp peptide by receptor
min
max

**H**

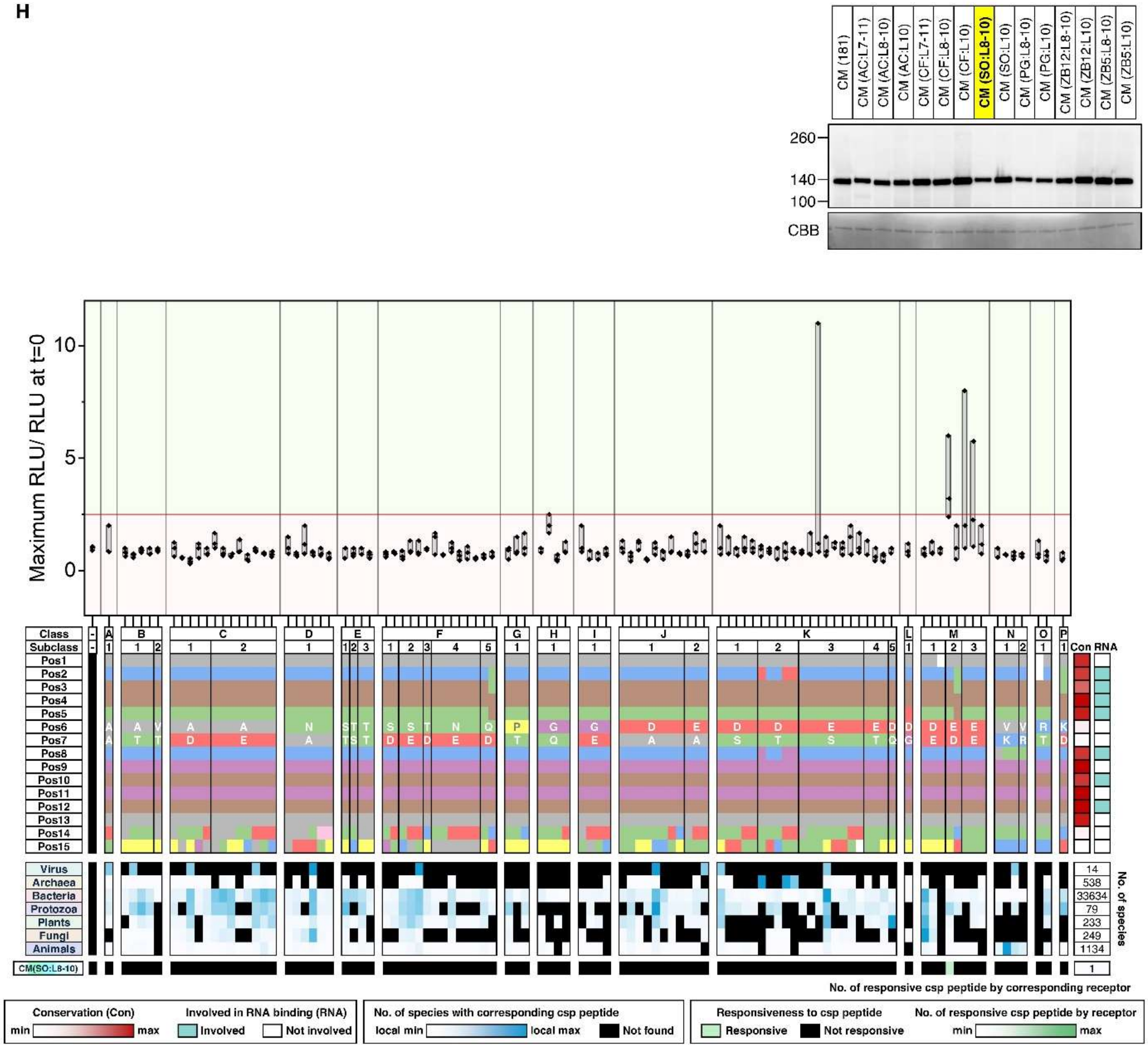

CM (181)
CM (AC:L7-11)
CM (AC:L8-10)
CM (AC:L10)
CM (CF:L7-11)
CM (CF:L8-10)
CM (CF:L10)
CM (SO:L8-10)
CM (SO:L10)
CM (PG:L8-10)
CM (PG:L10)
CM (ZB12:L8-10)
CM (ZB12:L10)
CM (ZB5:L8-10)
CM (ZB5:L10)
260
140
100
CBB
Maximum RLU/ RLU at t=0
10
5
0
Class
Subclass
Pos1
Pos2
Pos3
Pos4
Pos5
Pos6
Pos7
Pos8
Pos9
Pos10
Pos11
Pos12
Pos13
Pos14
Pos15
Con
RNA
Virus
Archaea
Bacteria
Protozoa
Plants
Fungi
Animals
14
538
33634
79
233
249
1134
No. of species
CM(SO:L8-10)
1
No. of responsive csp peptide by corresponding receptor
Conservation (Con)
min
max
Involved in RNA binding (RNA)
Involved
Not involved
No. of species with corresponding csp peptide
local min
local max
Not found
Responsiveness to csp peptide
Responsive
Not responsive
No. of responsive csp peptide by receptor
min
max

I

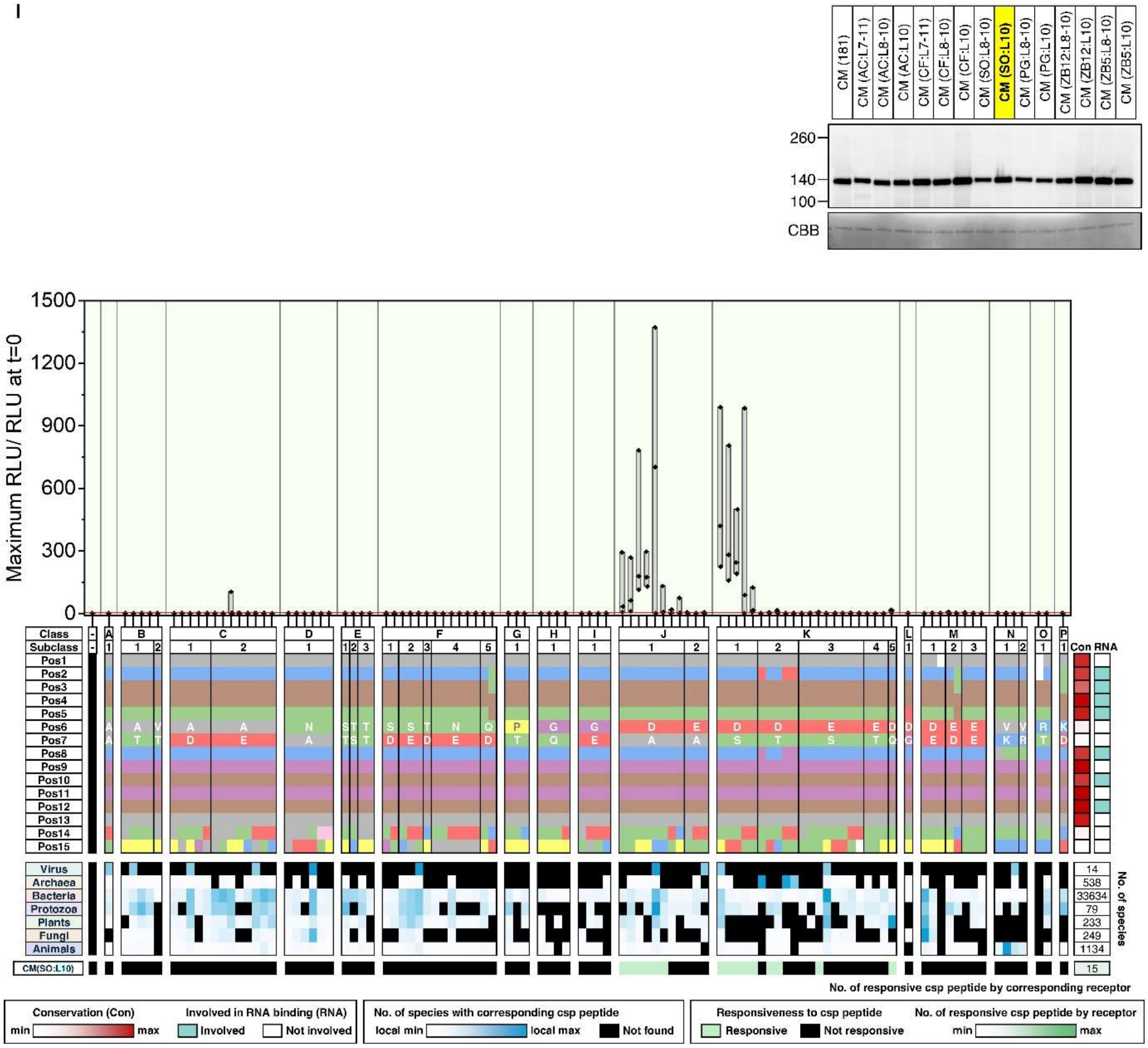

CM (181)
CM (AC:L7-11)
CM (AC:L8-10)
CM (AC:L10)
CM (CF:L7-11)
CM (CF:L8-10)
CM (CF:L10)
CM (SO:L8-10)
CM (SO:L10)
CM (PG:L8-10)
CM (PG:L10)
CM (ZB12:L8-10)
CM (ZB12:L10)
CM (ZB5:L8-10)
CM (ZB5:L10)
260
140
100
CBB
Maximum RLU/ RLU at t=0
1500
1200
900
600
300
0
Class
Subclass
Con
RNA
Virus
Archaea
Bacteria
Protozoa
Plants
Fungi
Animals
14
538
33634
79
233
249
1134
No. of species
CM(SO:L10)
15
No. of responsive csp peptide by corresponding receptor
Conservation (Con)
min
max
Involved in RNA binding (RNA)
Involved
Not involved
No. of species with corresponding csp peptide
local min
local max
Not found
Responsiveness to csp peptide
Responsive
Not responsive
No. of responsive csp peptide by receptor
min
max

**J**

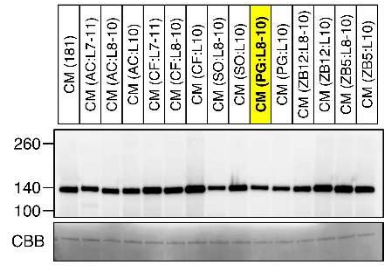

CM (181)
CM (AC:L7-11)
CM (AC:L8-10)
CM (AC:L10)
CM (CF:L7-11)
CM (CF:L8-10)
CM (CF:L10)
CM (SO:L8-10)
CM (SO:L10)
CM (PG:L8-10)
CM (PG:L10)
CM (ZB12:L8-10)
CM (ZB12:L10)
CM (ZB5:L8-10)
CM (ZB5:L10)
260
140
100
CBB


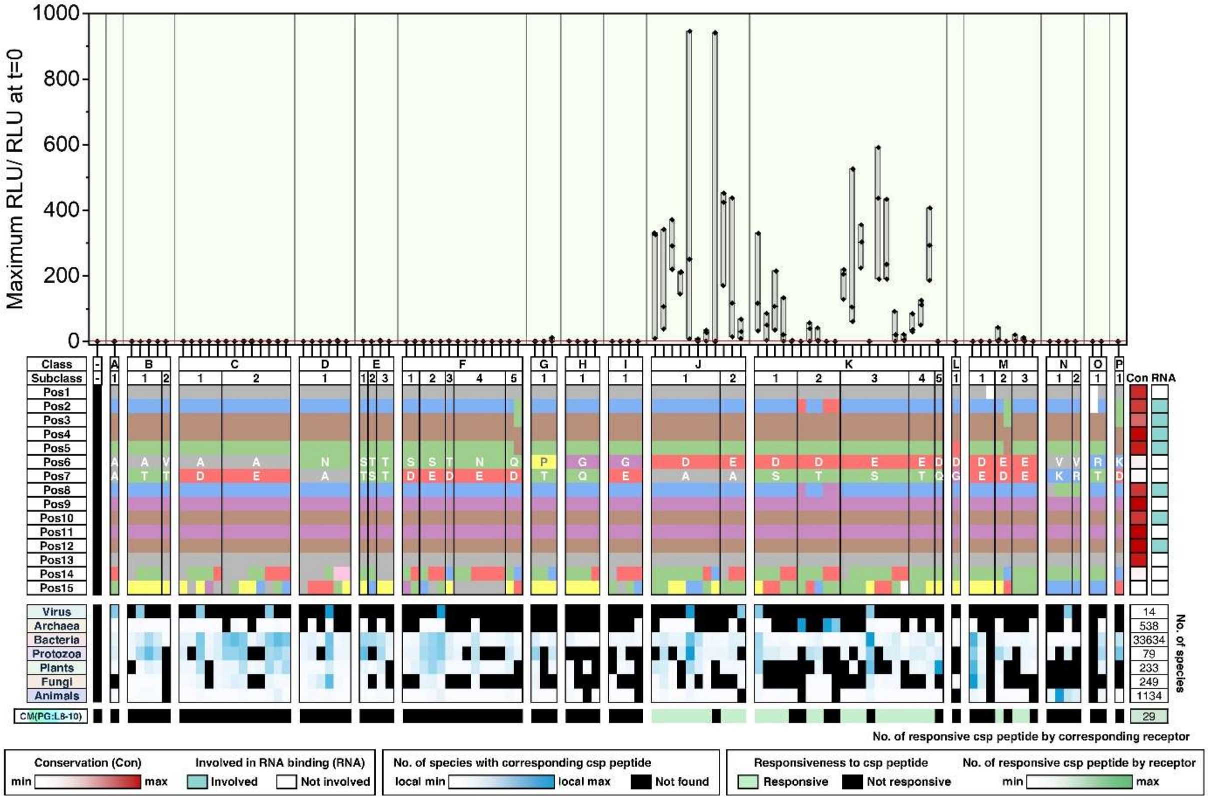

Maximum RLU/ RLU at t=0
1000
800
600
400
200
0
Class
Subclass
Pos1
Pos2
Pos3
Pos4
Pos5
Pos6
Pos7
Pos8
Pos9
Pos10
Pos11
Pos12
Pos13
Pos14
Pos15
Con
RNA
Virus
Archaea
Bacteria
Protozoa
Plants
Fungi
Animals
14
538
33634
79
233
249
1134
No. of species
CM(PG:L8-10)
29
No. of responsive csp peptide by corresponding receptor
Conservation (Con)
min
max
Involved in RNA binding (RNA)
Involved
Not involved
No. of species with corresponding csp peptide
local min
local max
Not found
Responsiveness to csp peptide
Responsive
Not responsive
No. of responsive csp peptide by receptor
min
max

**K**

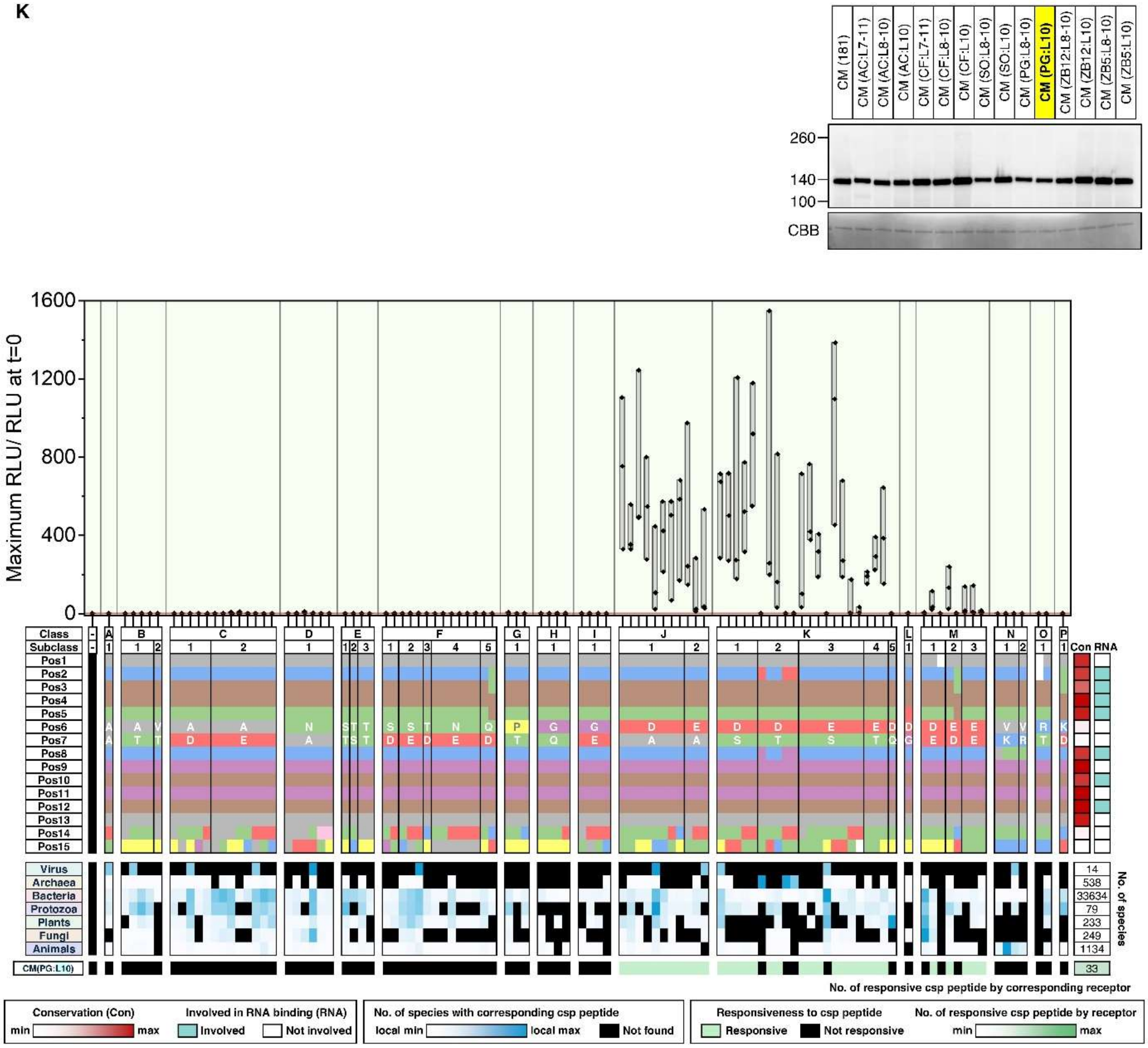
CM (181)
CM (AC:L7-11)
CM (AC:L8-10)
CM (AC:L10)
CM (CF:L7-11)
CM (CF:L8-10)
CM (CF:L10)
CM (SO:L8-10)
CM (SO:L10)
CM (PG:L8-10)
CM (PG:L10)
CM (ZB12:L8-10)
CM (ZB12:L10)
CM (ZB5:L8-10)
CM (ZB5:L10)
260
140
100
CBB
Maximum RLU/ RLU at t=0
1600
1200
800
400
0
Class
Subclass
Pos1
Pos2
Pos3
Pos4
Pos5
Pos6
Pos7
Pos8
Pos9
Pos10
Pos11
Pos12
Pos13
Pos14
Pos15
Con
RNA
Virus
Archaea
Bacteria
Protozoa
Plants
Fungi
Animals
14
538
33634
79
233
249
1134
No. of species
CM(PG:L10)
33
No. of responsive csp peptide by corresponding receptor
Conservation (Con)
min
max
Involved in RNA binding (RNA)
Involved
Not involved
No. of species with corresponding csp peptide
local min
local max
Not found
Responsiveness to csp peptide
Responsive
Not responsive
No. of responsive csp peptide by receptor
min
max

L

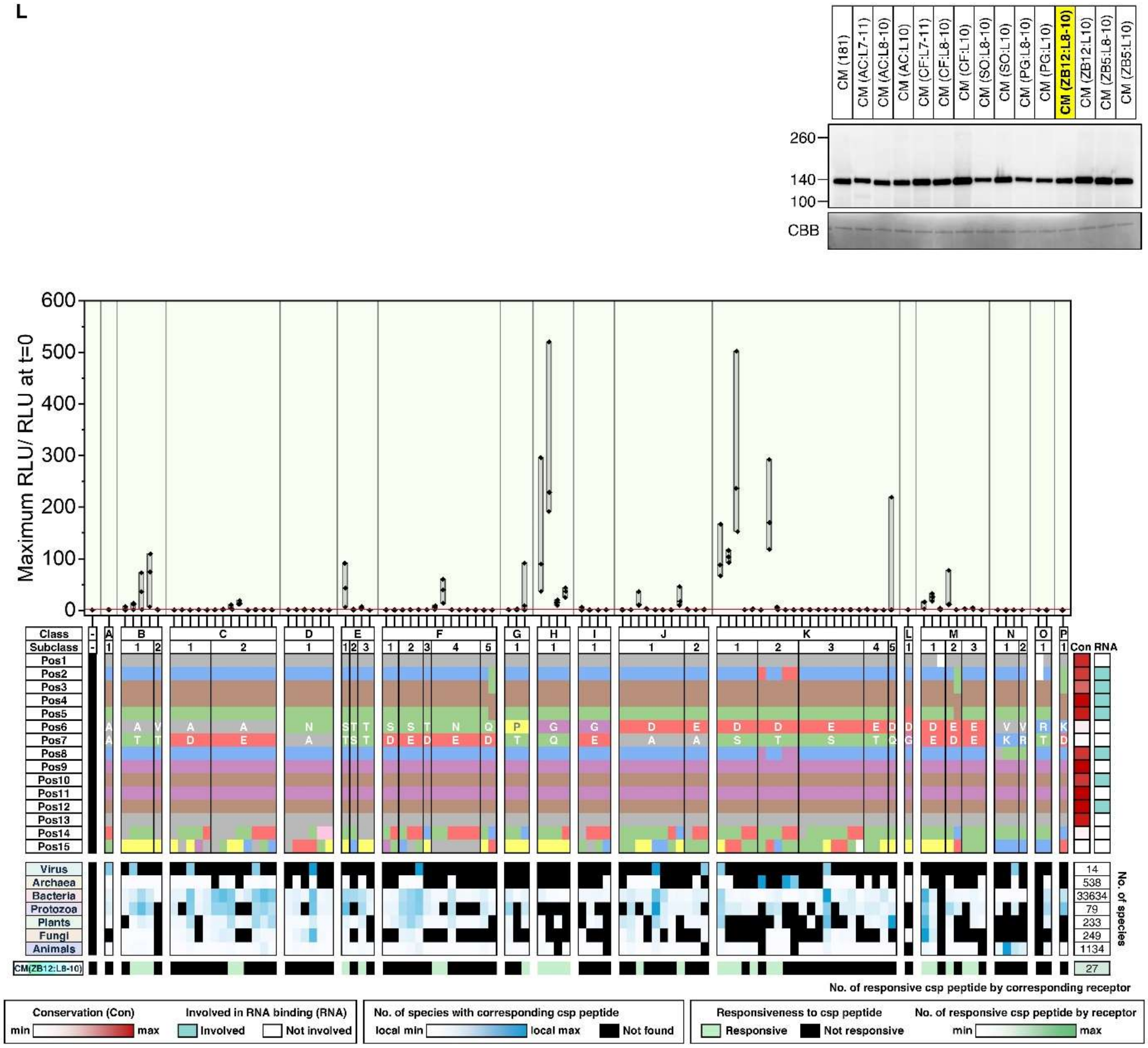
CM (181)
CM (AC:L7-11)
CM (AC:L8-10)
CM (AC:L10)
CM (CF:L7-11)
CM (CF:L8-10)
CM (CF:L10)
CM (SO:L8-10)
CM (SO:L10)
CM (PG:L8-10)
CM (PG:L10)
CM (ZB12:L8-10)
CM (ZB12:L10)
CM (ZB5:L8-10)
CM (ZB5:L10)
260
140
100
CBB
Maximum RLU/ RLU at t=0
600
500
400
300
200
100
0
Class
Subclass
Pos1
Pos2
Pos3
Pos4
Pos5
Pos6
Pos7
Pos8
Pos9
Pos10
Pos11
Pos12
Pos13
Pos14
Pos15
Con
RNA
Virus
Archaea
Bacteria
Protozoa
Plants
Fungi
Animals
14
538
33634
79
233
249
1134
No. of species
CM(ZB12:L8-10)
27
No. of responsive csp peptide by corresponding receptor
Conservation (Con)
min
max
Involved in RNA binding (RNA)
Involved
Not involved
No. of species with corresponding csp peptide
local min
local max
Not found
Responsiveness to csp peptide
Responsive
Not responsive
No. of responsive csp peptide by receptor
min
max

M

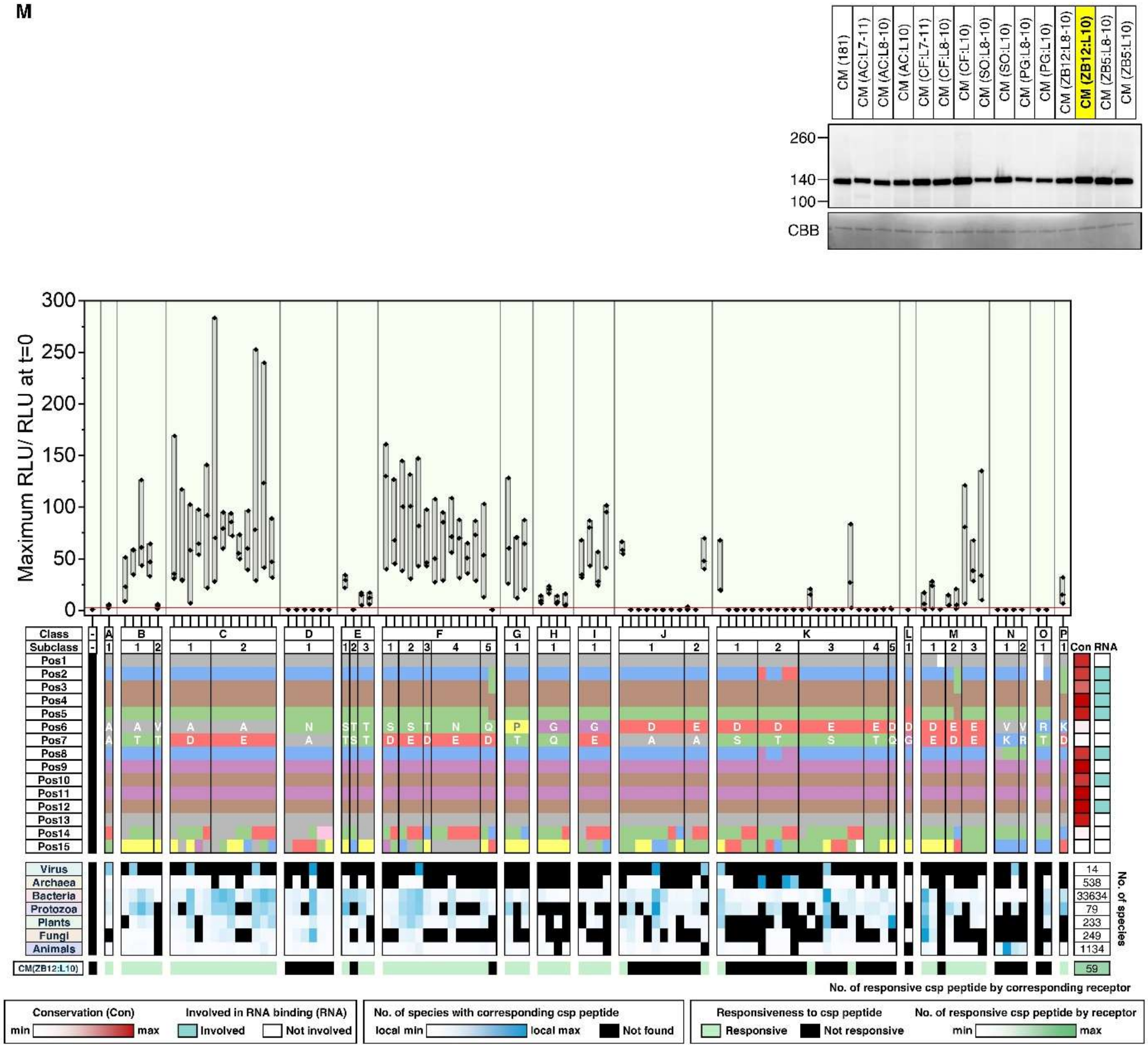
CM (181)
CM (AC:L7-11)
CM (AC:L8-10)
CM (AC:L10)
CM (CF:L7-11)
CM (CF:L8-10)
CM (CF:L10)
CM (SO:L8-10)
CM (SO:L10)
CM (PG:L8-10)
CM (PG:L10)
CM (ZB12:L8-10)
CM (ZB12:L10)
CM (ZB5:L8-10)
CM (ZB5:L10)
260
140
100
CBB
Maximum RLU/ RLU at t=0
Class
Subclass
Con RNA
Virus
Archaea
Bacteria
Protozoa
Plants
Fungi
Animals
No. of species
14
538
33634
79
233
249
1134
CM(ZB12:L10)
59
No. of responsive csp peptide by corresponding receptor
Conservation (Con)
Involved in RNA binding (RNA)
Involved
Not involved
No. of species with corresponding csp peptide
local min
local max
Not found
Responsiveness to csp peptide
Responsive
Not responsive
No. of responsive csp peptide by receptor
min
max

N

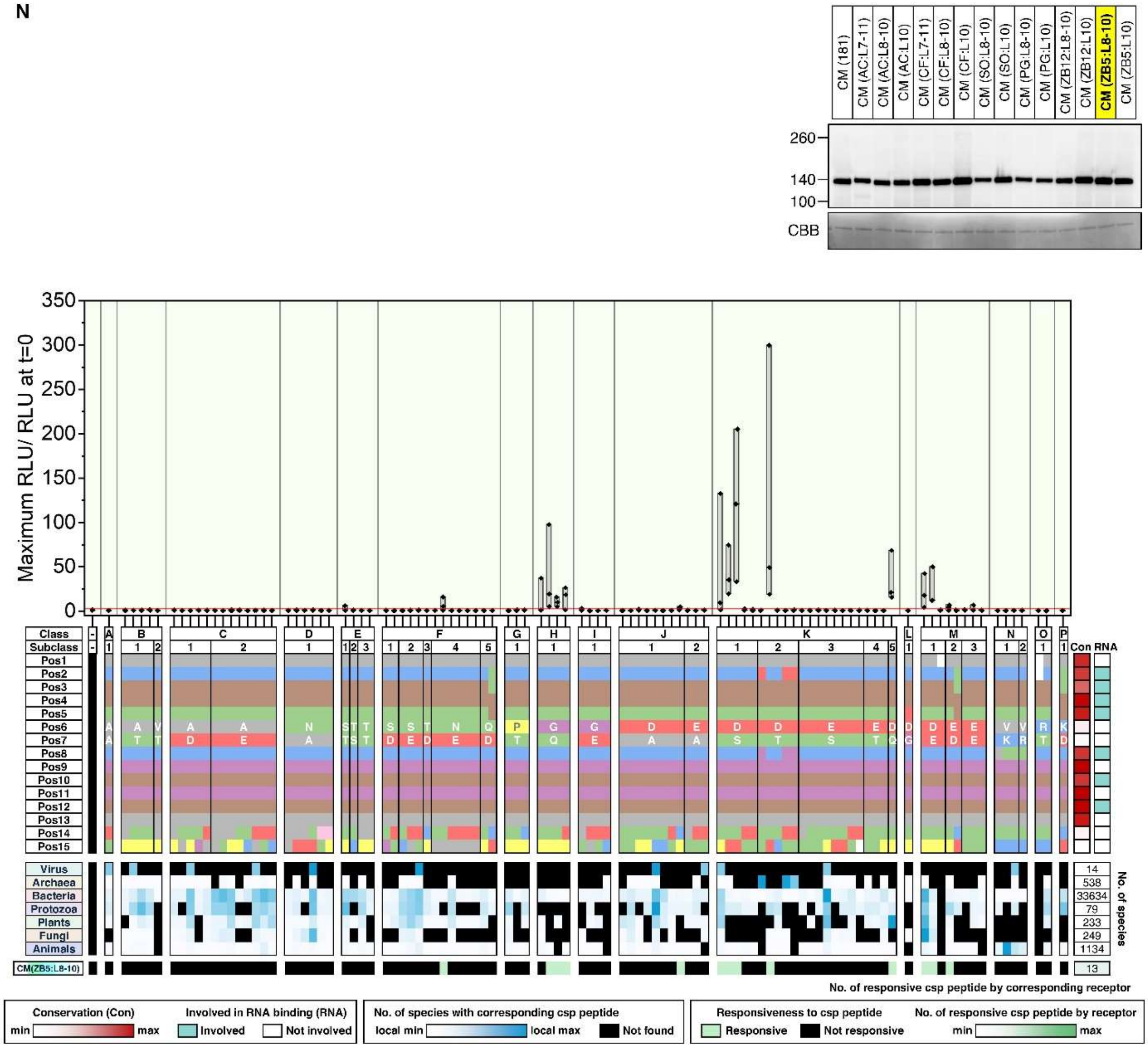
CM (181)
CM (AC:L7-11)
CM (AC:L8-10)
CM (AC:L10)
CM (CF:L7-11)
CM (CF:L8-10)
CM (CF:L10)
CM (SO:L8-10)
CM (SO:L10)
CM (PG:L8-10)
CM (PG:L10)
CM (ZB12:L8-10)
CM (ZB12:L10)
CM (ZB5:L8-10)
CM (ZB5:L10)
260
140
100
CBB
Maximum RLU/ RLU at t=0
350
300
250
200
150
100
50
0
Class
Subclass
Pos1
Pos2
Pos3
Pos4
Pos5
Pos6
Pos7
Pos8
Pos9
Pos10
Pos11
Pos12
Pos13
Pos14
Pos15
Con
RNA
Virus
Archaea
Bacteria
Protozoa
Plants
Fungi
Animals
14
538
33634
79
233
249
1134
No. of species
CM(ZB5:L8-10)
13
No. of responsive csp peptide by corresponding receptor
Conservation (Con)
min
max
Involved in RNA binding (RNA)
Involved
Not involved
No. of species with corresponding csp peptide
local min
local max
Not found
Responsiveness to csp peptide
Responsive
Not responsive
No. of responsive csp peptide by receptor
min
max

o

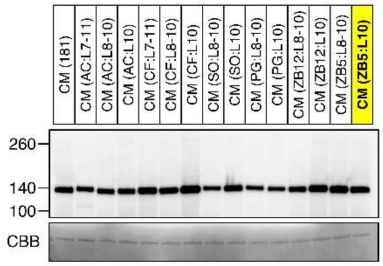

CM (181)
CM (AC:L7-11)
CM (AC:L8-10)
CM (AC:L10)
CM (CF:L7-11)
CM (CF:L8-10)
CM (CF:L10)
CM (SO:L8-10)
CM (SO:L10)
CM (PG:L8-10)
CM (PG:L10)
CM (ZB12:L8-10)
CM (ZB12:L10)
CM (ZB5:L8-10)
CM (ZB5:L10)
260
140
100
CBB


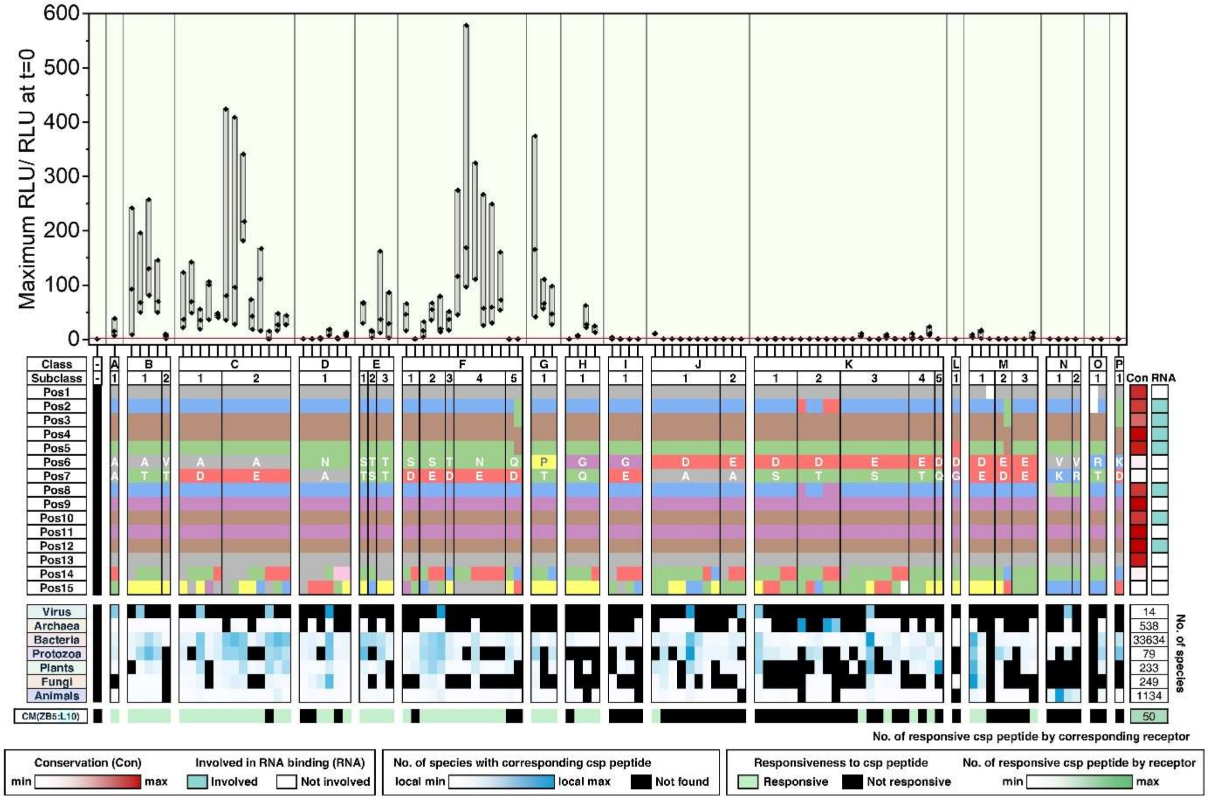

Maximum RLU/ RLU at t=0
600
500
400
300
200
100
0
Class
Subclass
Pos1
Pos2
Pos3
Pos4
Pos5
Pos6
Pos7
Pos8
Pos9
Pos10
Pos11
Pos12
Pos13
Pos14
Pos15
Con
RNA
Virus
Archaea
Bacteria
Protozoa
Plants
Fungi
Animals
14
538
33634
79
233
249
1134
No. of species
CM(ZB5:L10)
50
No. of responsive csp peptide by corresponding receptor
Conservation (Con)
min
max
Involved in RNA binding (RNA)
Involved
Not involved
No. of species with corresponding csp peptide
local min
local max
Not found
Responsiveness to csp peptide
Responsive
Not responsive
No. of responsive csp peptide by receptor
min
max

**Fig. S22. ROS data related to Fig 5D (domain swapping).**

**(A)** The top chart represents the 103 csp15 peptides that are synthesized in this study. The amino acid properties in each position (Pos) are indicated with corresponding colors used in Fig. 4A. Peptides are grouped into classes and subclasses based on residues at the positions 6 and 7. Red heatmap on the right represents residue conservation scores, and cyan blocks mark RNA binding residues in CSPs. The middle blue heatmap represents the prevalence of the 103 csp15 peptide in each kingdom. Black blocks indicate peptides absent from all species in a kingdom. The number of species searched in each kingdom is shown on the right. Bottom chart represents the CSP recognition capacity of SCORE orthologs and variants (CM) generated from domain swaps. A peptide was considered active if ROS readout exceeds the baseline by 2.5 times (green; see below). Non-responsive peptides are labelled in black. Green heatmap on the right indicates the number of responsive peptides (out of 103) for each receptor. **(B-O)** Top immunoblot: Samples used in the following ROS assays were analysed with immunoblotting to confirm receptor expression. CBB staining is included as loading controls. Middle bar chart represents the ROS responses to mock treatment or each of the 103 csp15 peptide by the corresponding receptors. Leaves were then treated with 100 nM of indicated peptides, and ROS production was recorded for 15 minutes. For the box plots, the center line represents the median; the box bounds represent the 25th and 75th percentiles; and the whiskers extend to 1.5 × the interquartile range (IQR) from the 25th and 75th percentiles. Data points from 3 technical replicates were analyzed. A peptide was considered active if at least 2 out of 3 technical replicates showed ROS levels exceeding the baseline by 2.5 times (marked by the red line). Bottom charts: The top chart displays the 103 csp15 peptides synthesized in this study. Amino acid properties at each position (Pos) are color-coded, as in in Fig. 4A. Peptides are grouped into classes and subclasses based on residues at the positions 6 and 7. A red heatmap on the right indicates residue conservation scores, while cyan blocks mark RNA-binding residues in CSPs. The middle blue heatmap shows the prevalence of each csp15 peptide across kingdom. Black blocks represent peptides absent from all species within a kingdom. The number of species analyzed in each kingdom is shown on the right. Bottom chart represents the CSP recognition capacity of the corresponding receptors in plants. As noted above, peptides were considered active if at least 2 out of 3 technical replicates showed ROS levels exceeding the baseline by 2.5 times (see above). Non-responsive peptides are labeled in black. The green heatmap on the right indicates the number of responsive peptides (out of 103) for each receptor.

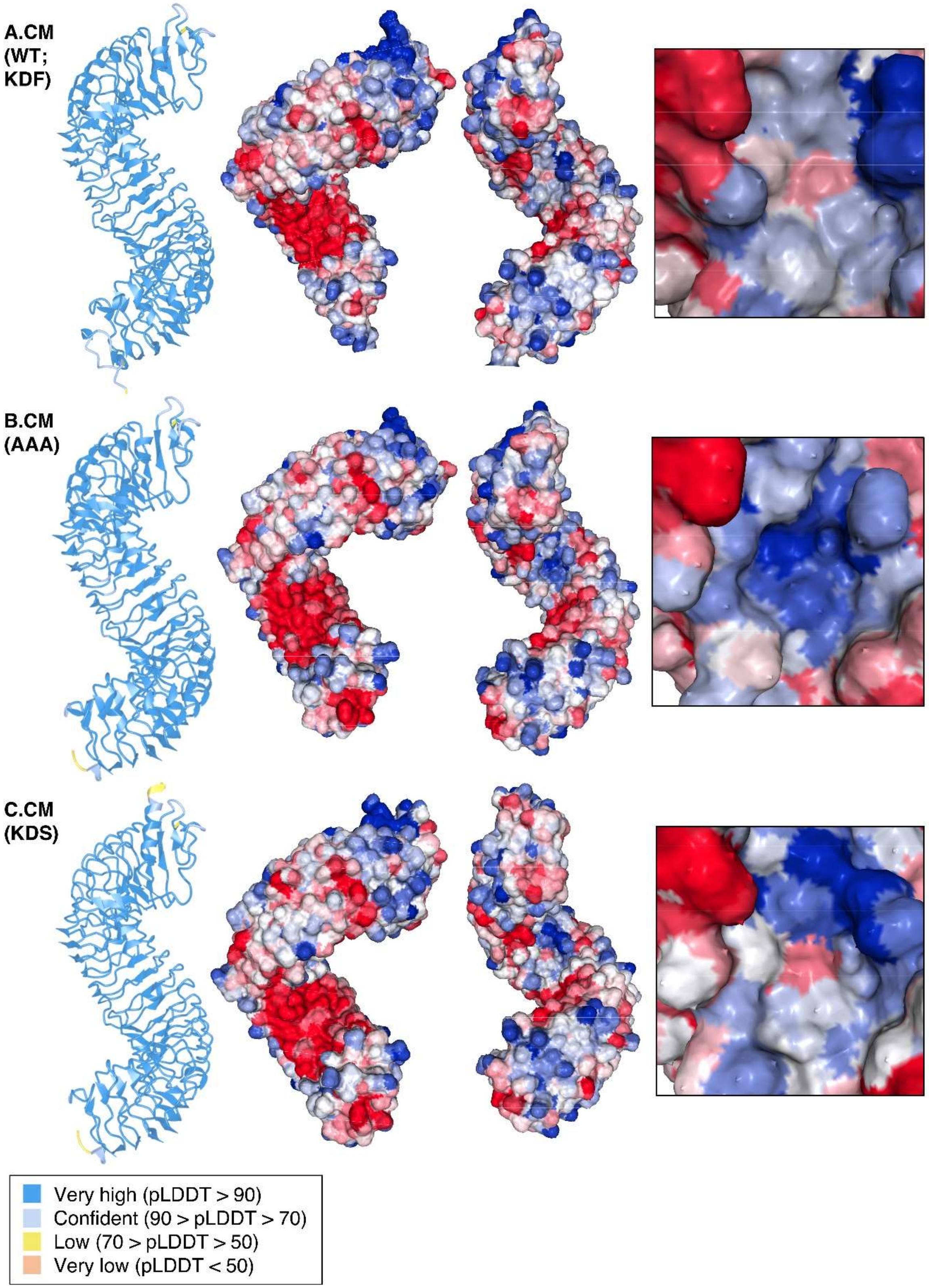
A.CM
(WT;
KDF)
B.CM
(AAA)
C.CM
(KDS)
Very high (pLDDT > 90)
Confident (90 > pLDDT > 70)
Low (70 > pLDDT > 50)
Very low (pLDDT < 50)

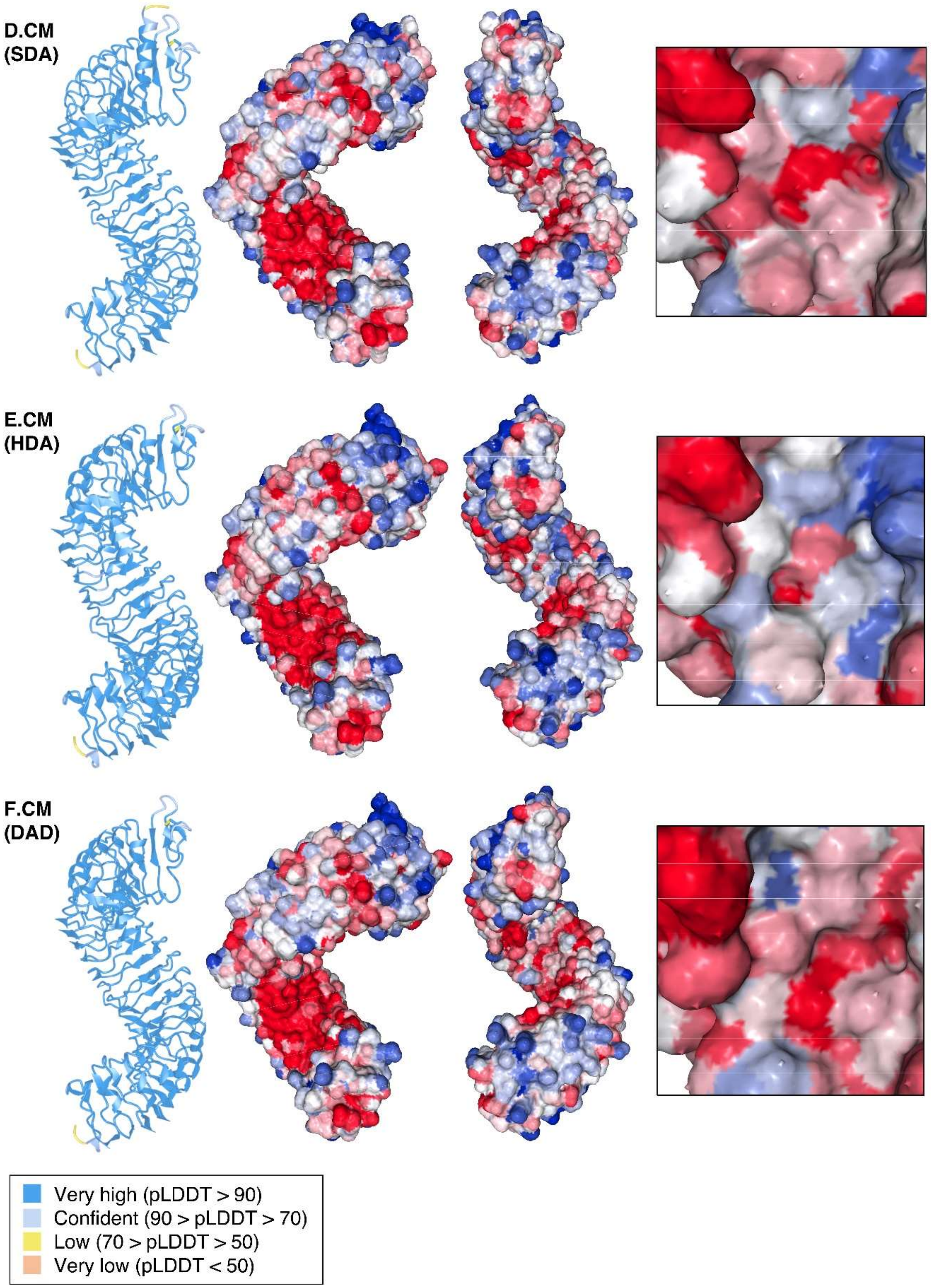
D.CM
(SDA)
E.CM
(HDA)
F.CM
(DAD)
Very high (pLDDT > 90)
Confident (90 > pLDDT > 70)
Low (70 > pLDDT > 50)
Very low (pLDDT < 50)

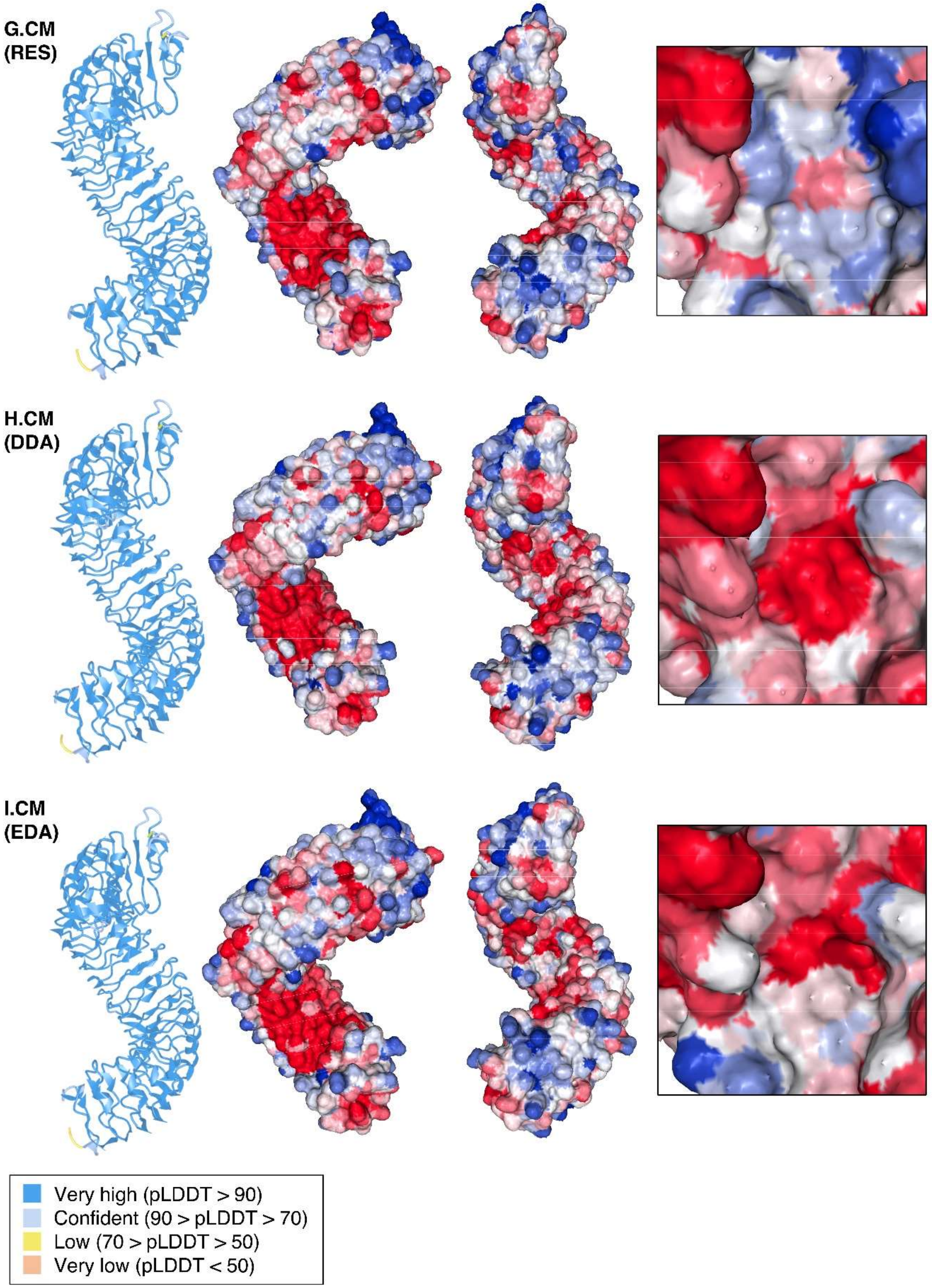
G.CM
(RES)
H.CM
(DDA)
I.CM
(EDA)
Very high (pLDDT > 90)
Confident (90 > pLDDT > 70)
Low (70 > pLDDT > 50)
Very low (pLDDT < 50)

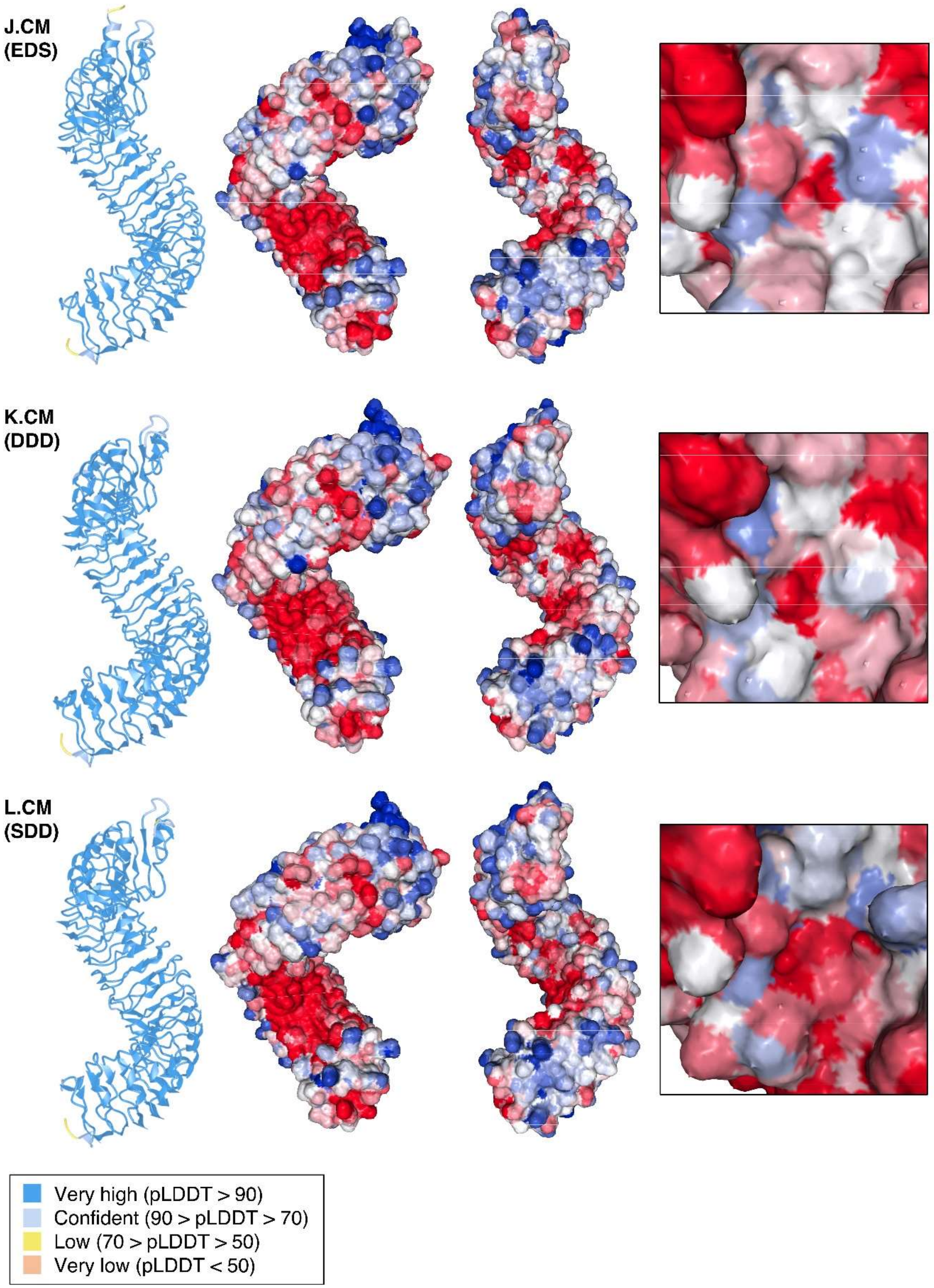
J.CM
(EDS)
K.CM
(DDD)
L.CM
(SDD)
Very high (pLDDT > 90)
Confident (90 > pLDDT > 70)
Low (70 > pLDDT > 50)
Very low (pLDDT < 50)

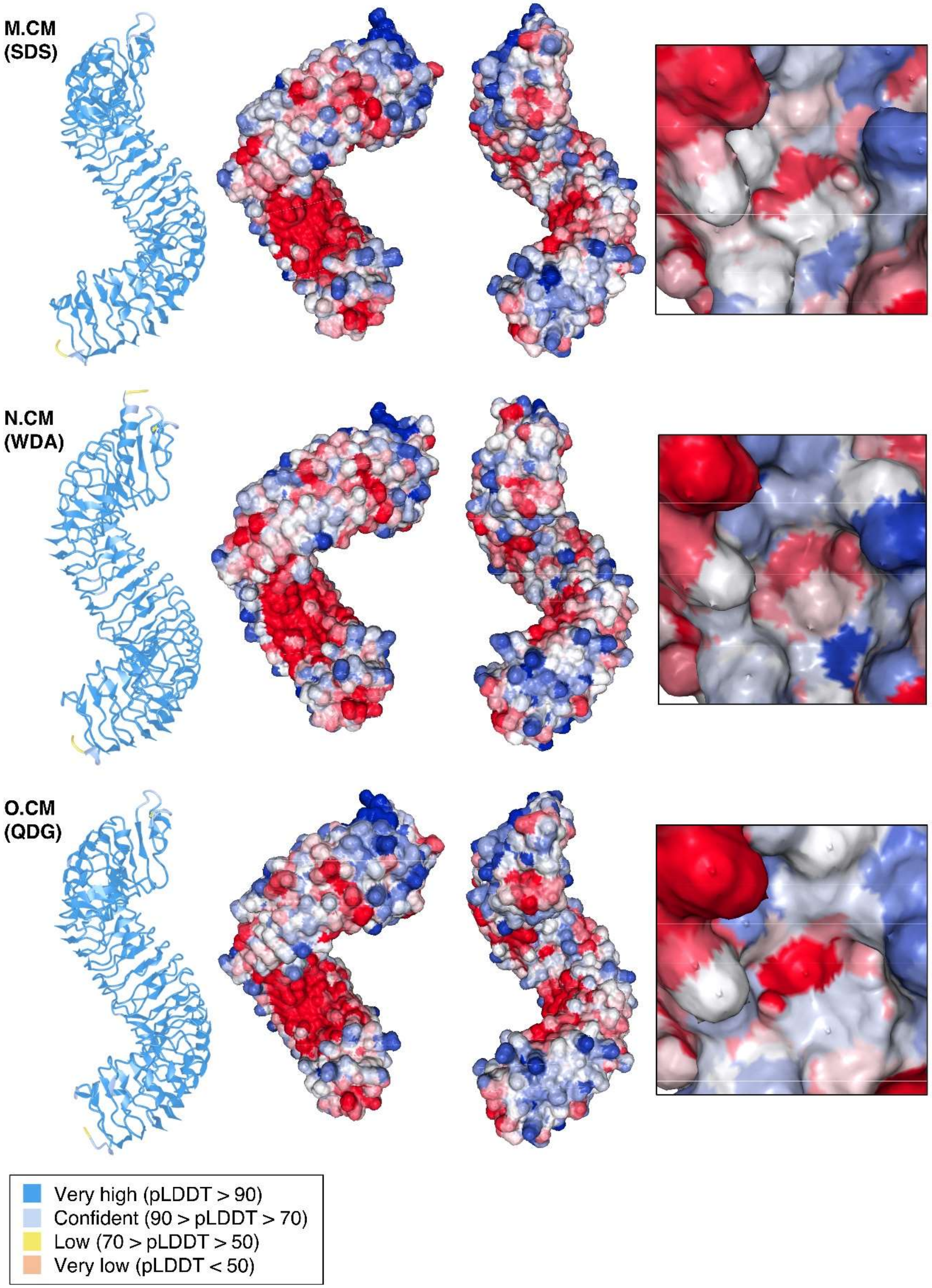
M.CM
(SDS)
N.CM
(WDA)
O.CM
(QDG)
Very high (pLDDT > 90)
Confident (90 > pLDDT > 70)
Low (70 > pLDDT > 50)
Very low (pLDDT < 50)

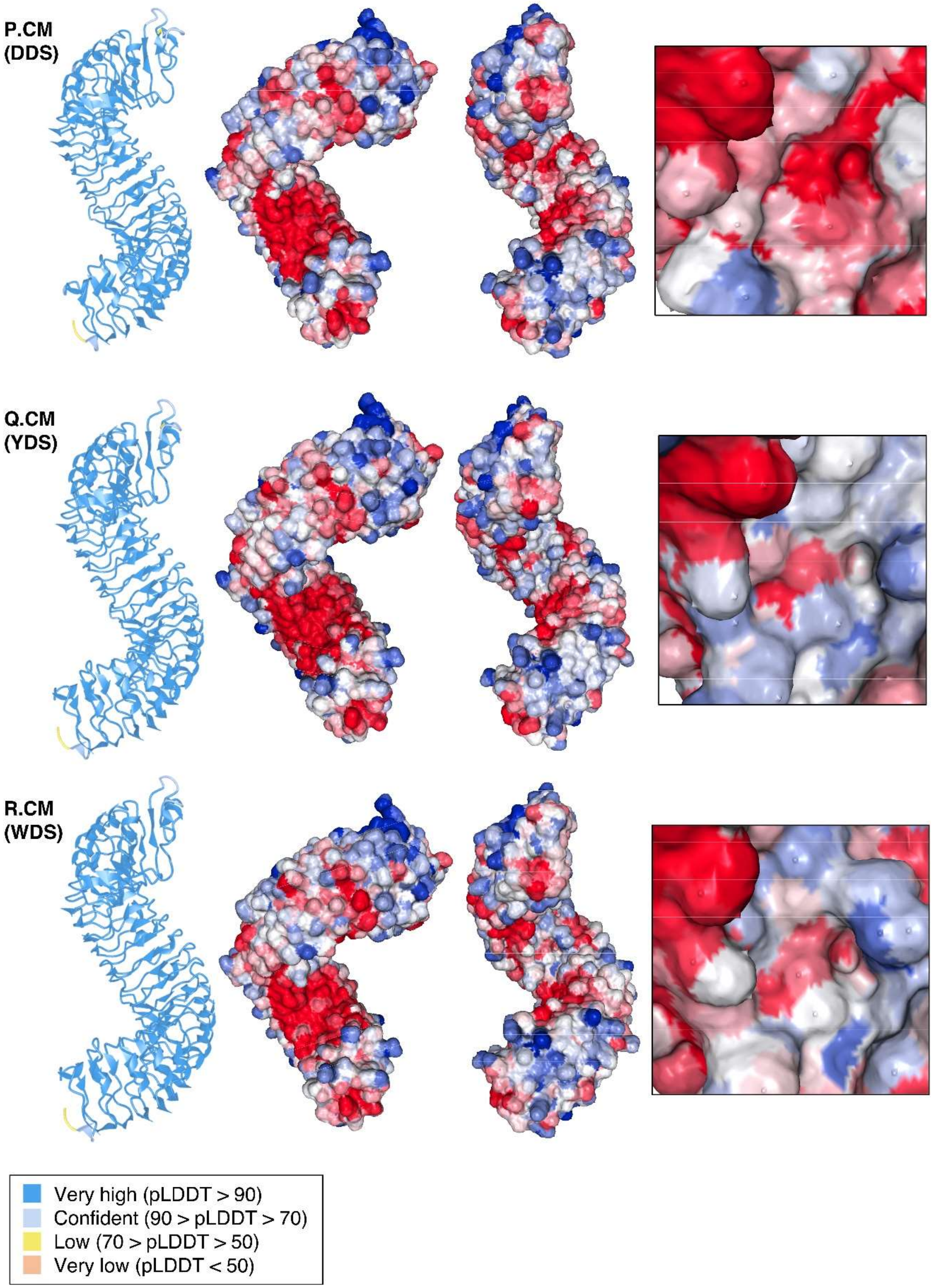
P.CM
(DDS)
Q.CM
(YDS)
R.CM
(WDS)
Very high (pLDDT > 90)
Confident (90 > pLDDT > 70)
Low (70 > pLDDT > 50)
Very low (pLDDT < 50)

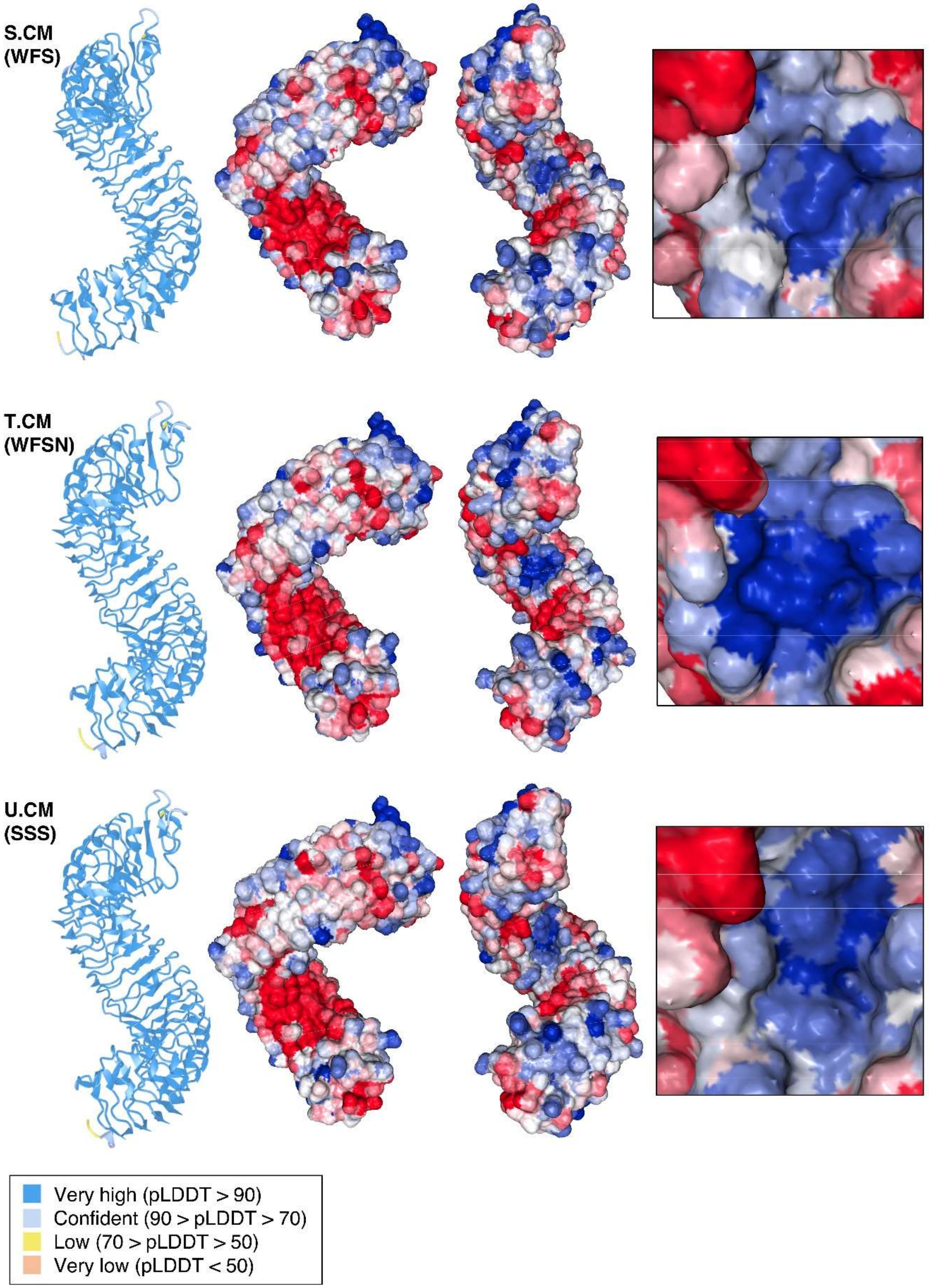

S.CM
(WFS)
T.CM
(WFSN)
U.CM
(SSS)
Very high (pLDDT > 90)
Confident (90 > pLDDT > 70)
Low (70 > pLDDT > 50)
Very low (pLDDT < 50)

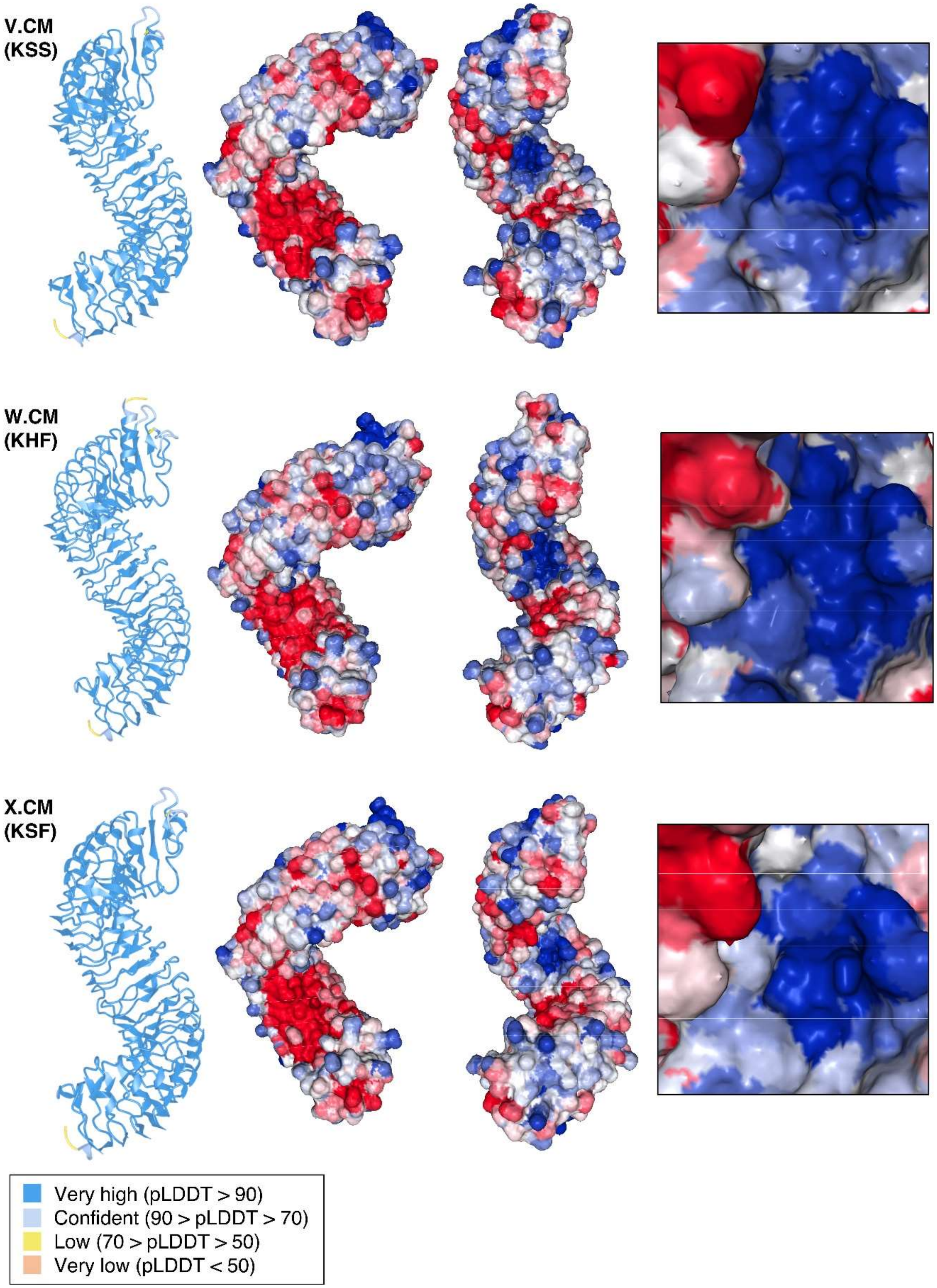
V.CM
(KSS)
W.CM
(KHF)
X.CM
(KSF)
Very high (pLDDT > 90)
Confident (90 > pLDDT > 70)
Low (70 > pLDDT > 50)
Very low (pLDDT < 50)

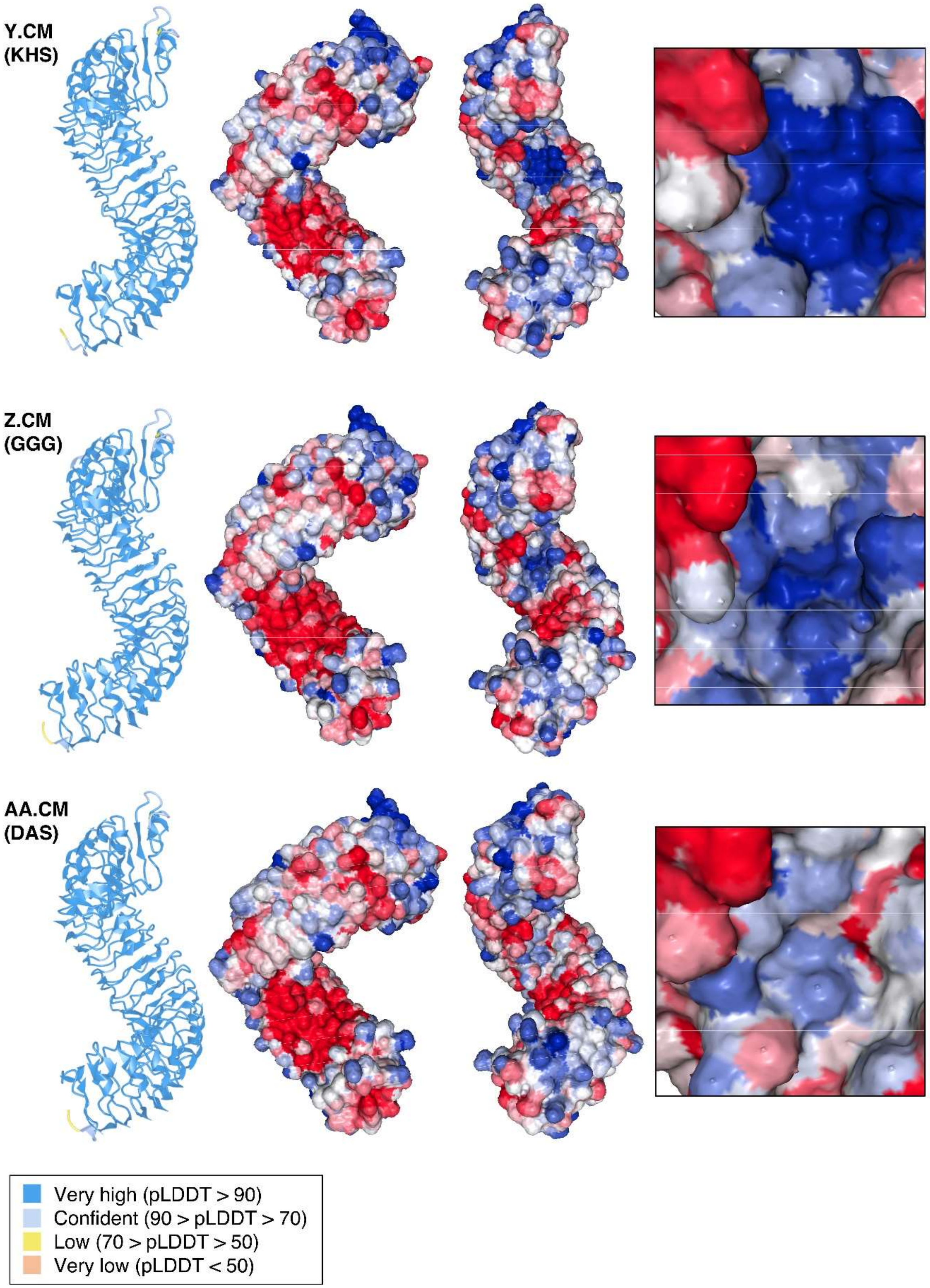
Y.CM
(KHS)
Z.CM
(GGG)
AA.CM
(DAS)
Very high (pLDDT > 90)
Confident (90 > pLDDT > 70)
Low (70 > pLDDT > 50)
Very low (pLDDT < 50)

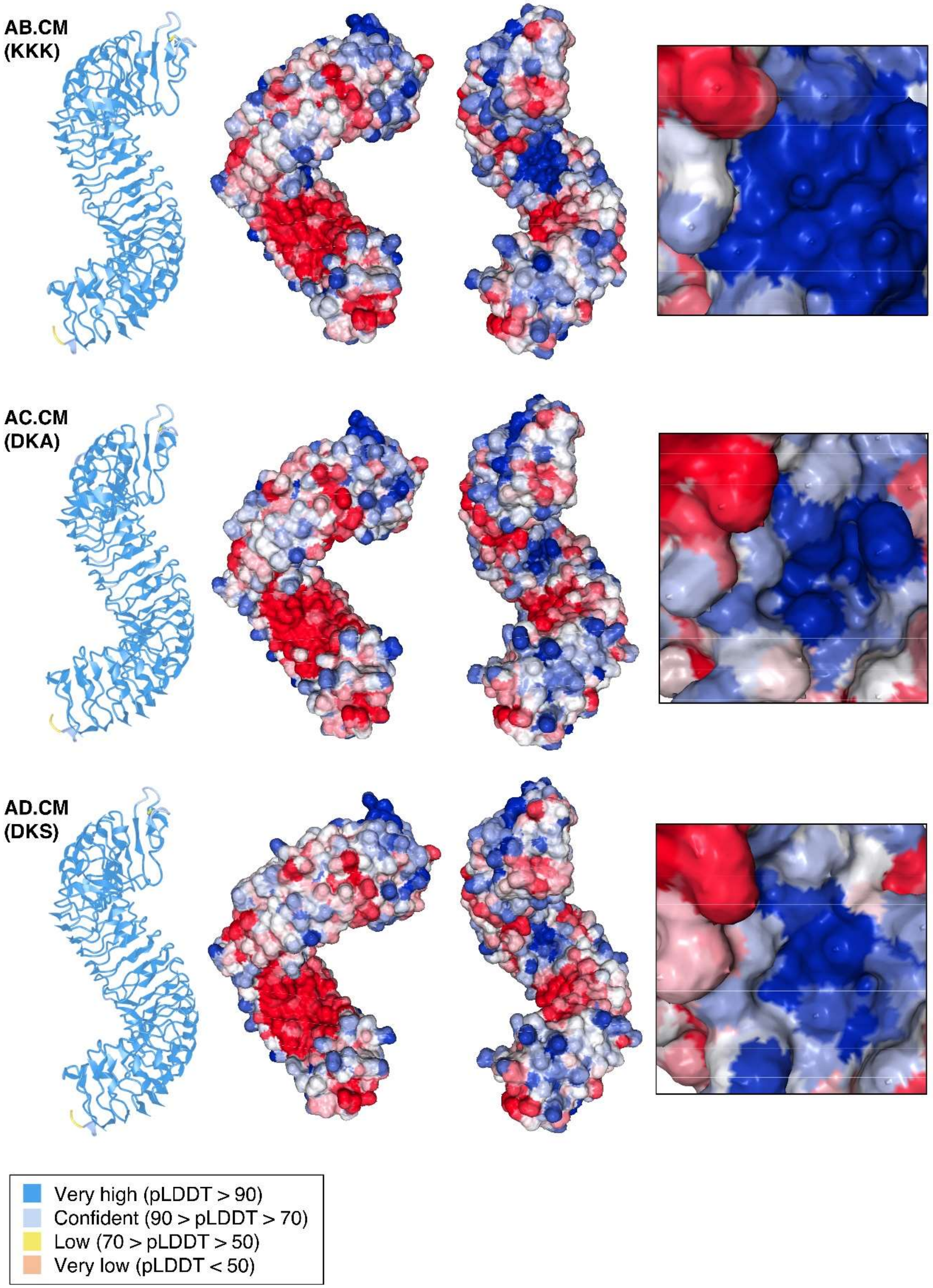
AB.CM
(KKK)
AC.CM
(DKA)
AD.CM
(DKS)
Very high (pLDDT > 90)
Confident (90 > pLDDT > 70)
Low (70 > pLDDT > 50)
Very low (pLDDT < 50)

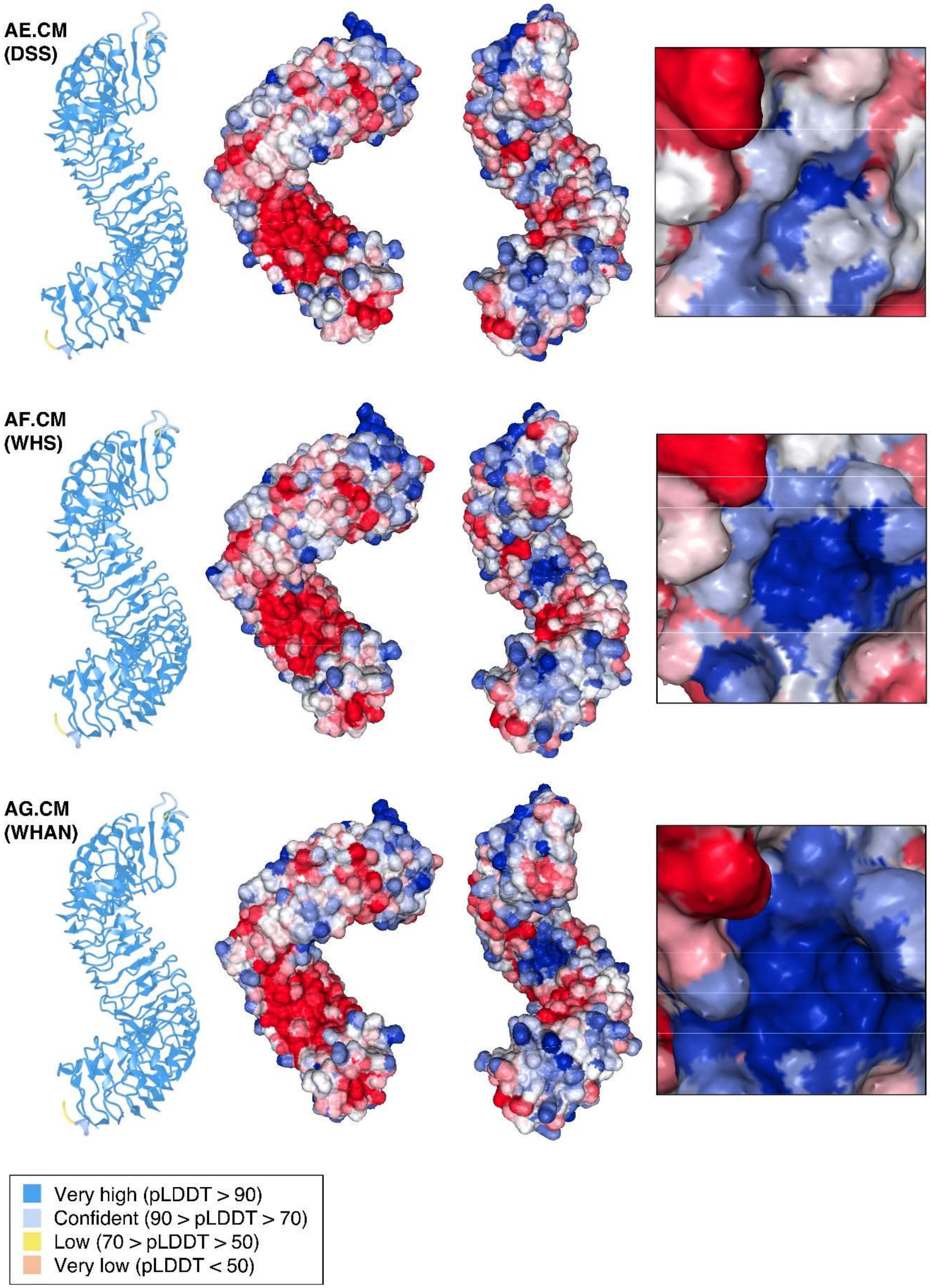
AE.CM
(DSS)
AF.CM
(WHS)
AG.CM
(WHAN)
Very high (pLDDT > 90)
Confident (90 > pLDDT > 70)
Low (70 > pLDDT > 50)
Very low (pLDDT < 50)

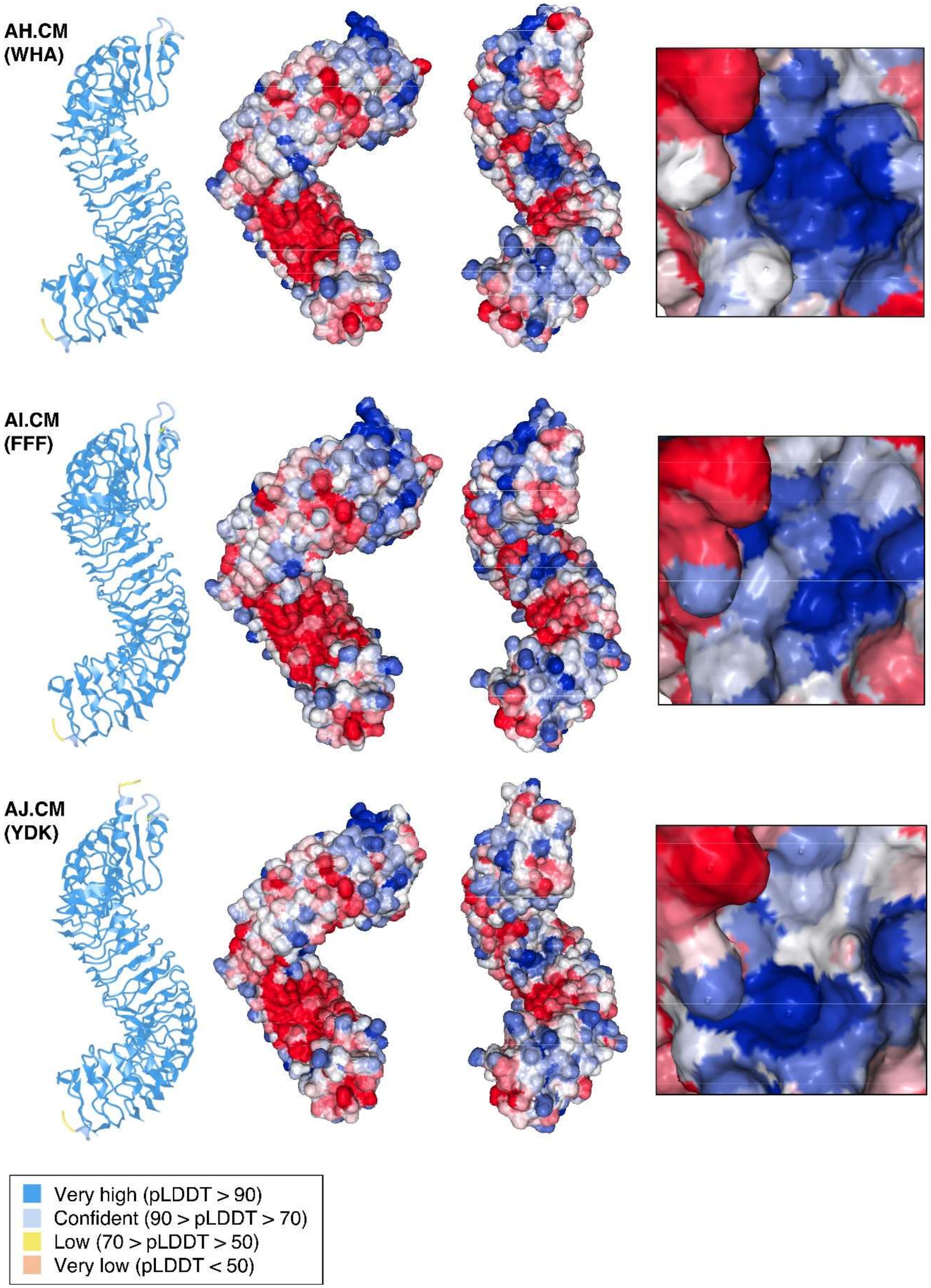

AH.CM
(WHA)
AI.CM
(FFF)
AJ.CM
(YDK)
Very high (pLDDT > 90)
Confident (90 > pLDDT > 70)
Low (70 > pLDDT > 50)
Very low (pLDDT < 50)

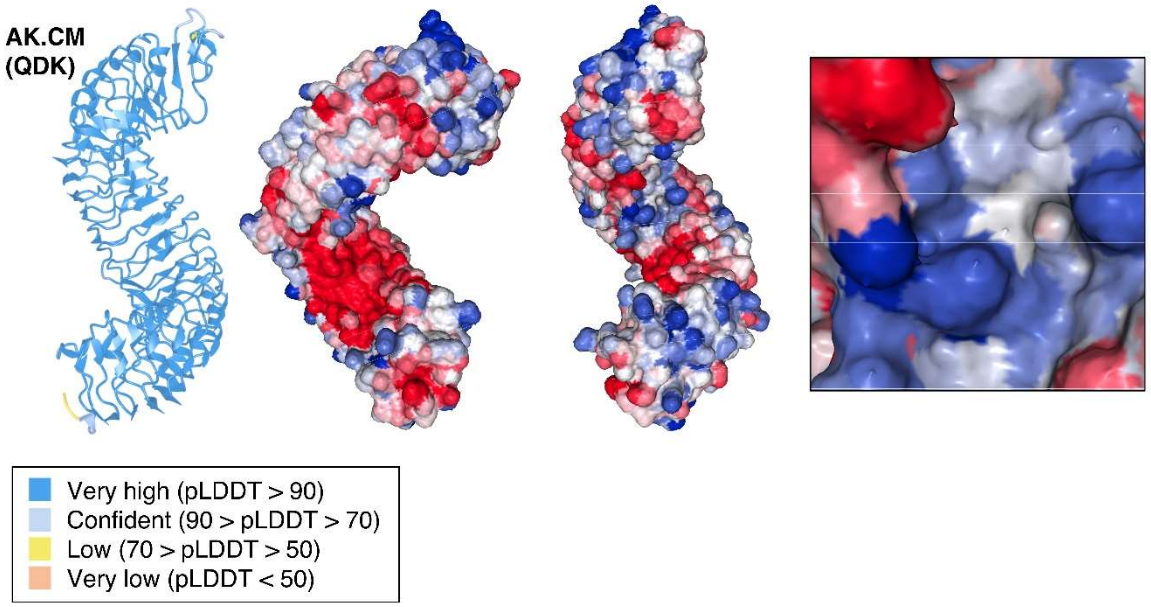


**Fig. S23. Alphafold3 structure prediction and surface charge distribution of SCORE (CM) and variants generated from amino acid substitutions.**

Left structure: The predicted protein structure is color-coded based on pLDDT scores (see legend below). Middle structures: SCORE$^{Ecto}$ with surface charges displayed. Red indicates negative charge, blue indicates positive charge, and white indicates neutral charge. Right box: Surface charge distribution on the inner LRR surface between the 8$^{th}$-11$^{th}$ LRR motifs. For labels of the SCORE (CM) variants generated from amino acid substitutions in **(A-AK)**, refer to Fig. 5D-E. Protein structures were visualized using iCn3D, and surface charges were calculated using protein-sol (24).

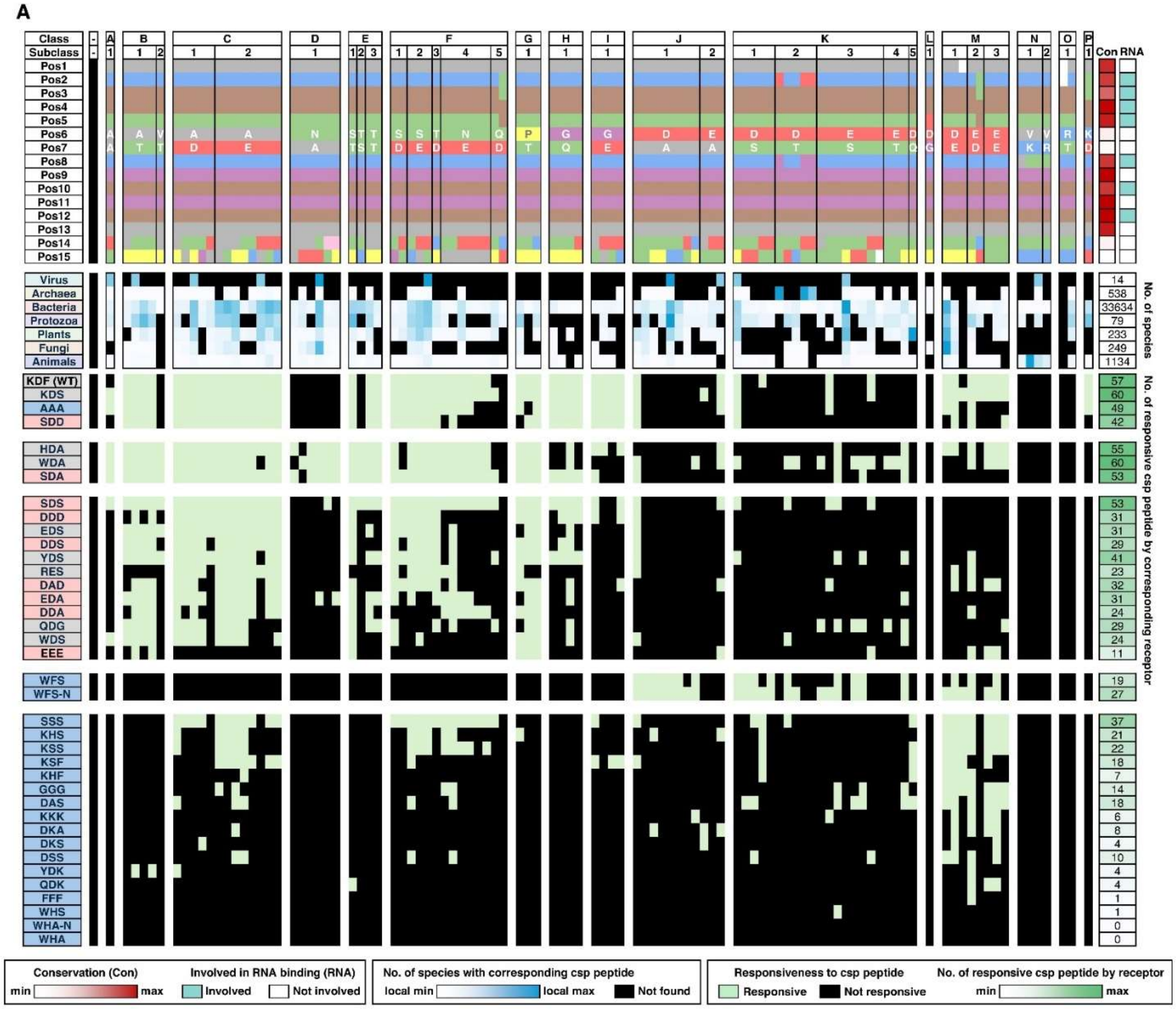
A
Class
Subclass
Pos1
Pos2
Pos3
Pos4
Pos5
Pos6
Pos7
Pos8
Pos9
Pos10
Pos11
Pos12
Pos13
Pos14
Pos15
Con
RNA
Virus
Archaea
Bacteria
Protozoa
Plants
Fungi
Animals
14
538
33634
79
233
249
1134
No. of species
KDF (WT)
KDS
AAA
SDD
HDA
WDA
SDA
SDS
DDD
EDS
DDS
YDS
RES
DAD
EDA
DDA
QDG
WDS
EEE
WFS
WFS-N
SSS
KHS
KSS
KSF
KHF
GGG
DAS
KKK
DKA
DKS
DSS
YDK
QDK
FFF
WHS
WHA-N
WHA
No. of responsive csp peptide by corresponding receptor
Conservation (Con)
min
max
Involved in RNA binding (RNA)
Involved
Not involved
No. of species with corresponding csp peptide
local min
local max
Not found
Responsiveness to csp peptide
Responsive
Not responsive
No. of responsive csp peptide by receptor
min
max

**B**

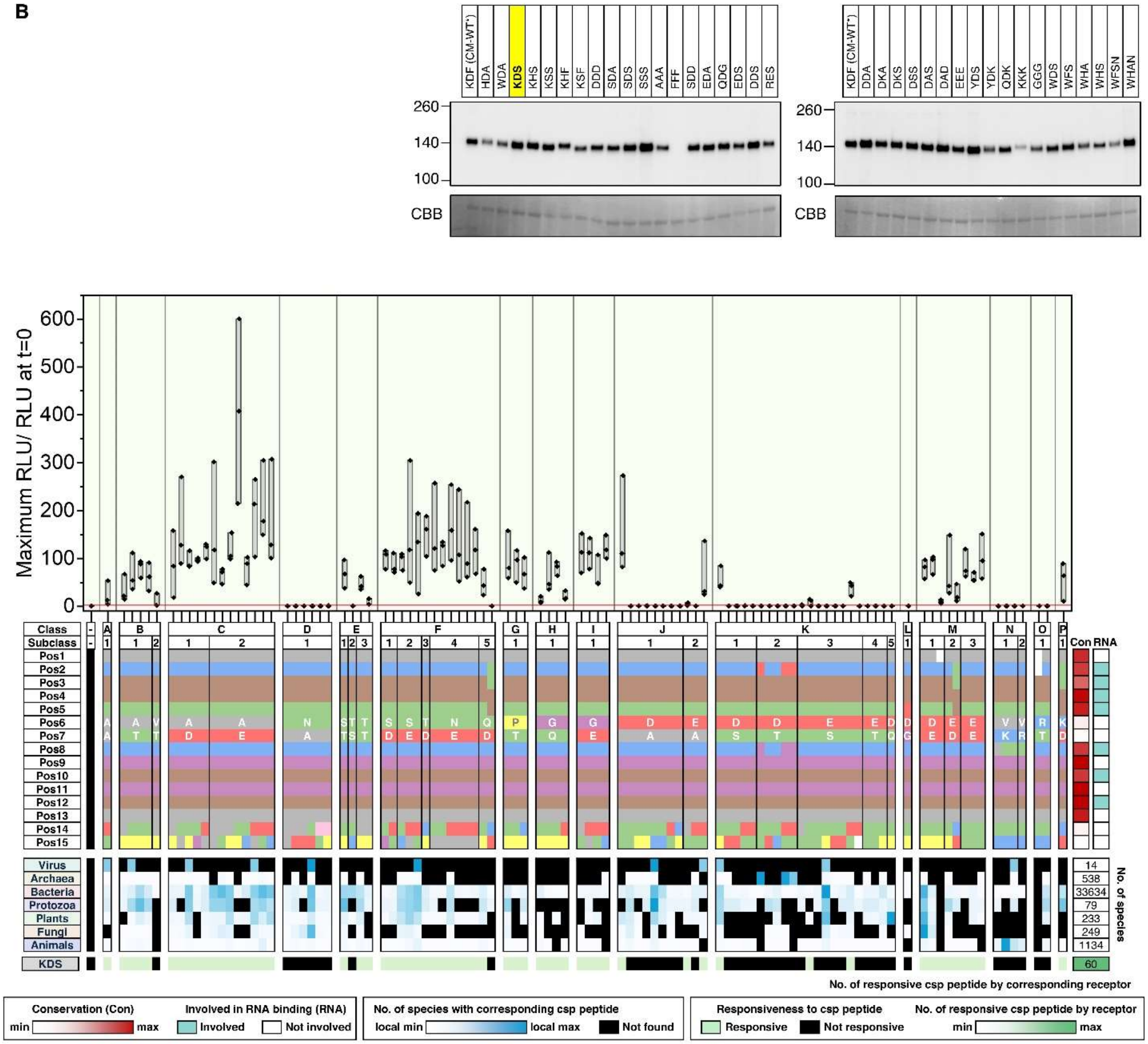

KDF (CM-WT*) HDA WDA KDS KHS KSS KHF KSF DDD SDA SDS SSS AAA FFF SDD EDA QDG EDS DDS RES
KDF (CM-WT*) DDA DKA DKS DSS DAS DAD EEE YDS YDK QDK KKK GGG WDS WFS WHA WHS WFSN WHAN
260
140
100
CBB
Maximum RLU/ RLU at t=0
Class
Subclass
Virus
Archaea
Bacteria
Protozoa
Plants
Fungi
Animals
KDS
No. of species
14
538
33634
79
233
249
1134
60
Con RNA
No. of responsive csp peptide by corresponding receptor
Conservation (Con)
min max
Involved in RNA binding (RNA)
Involved
Not involved
No. of species with corresponding csp peptide
local min
local max
Not found
Responsiveness to csp peptide
Responsive
Not responsive
No. of responsive csp peptide by receptor
min max

c

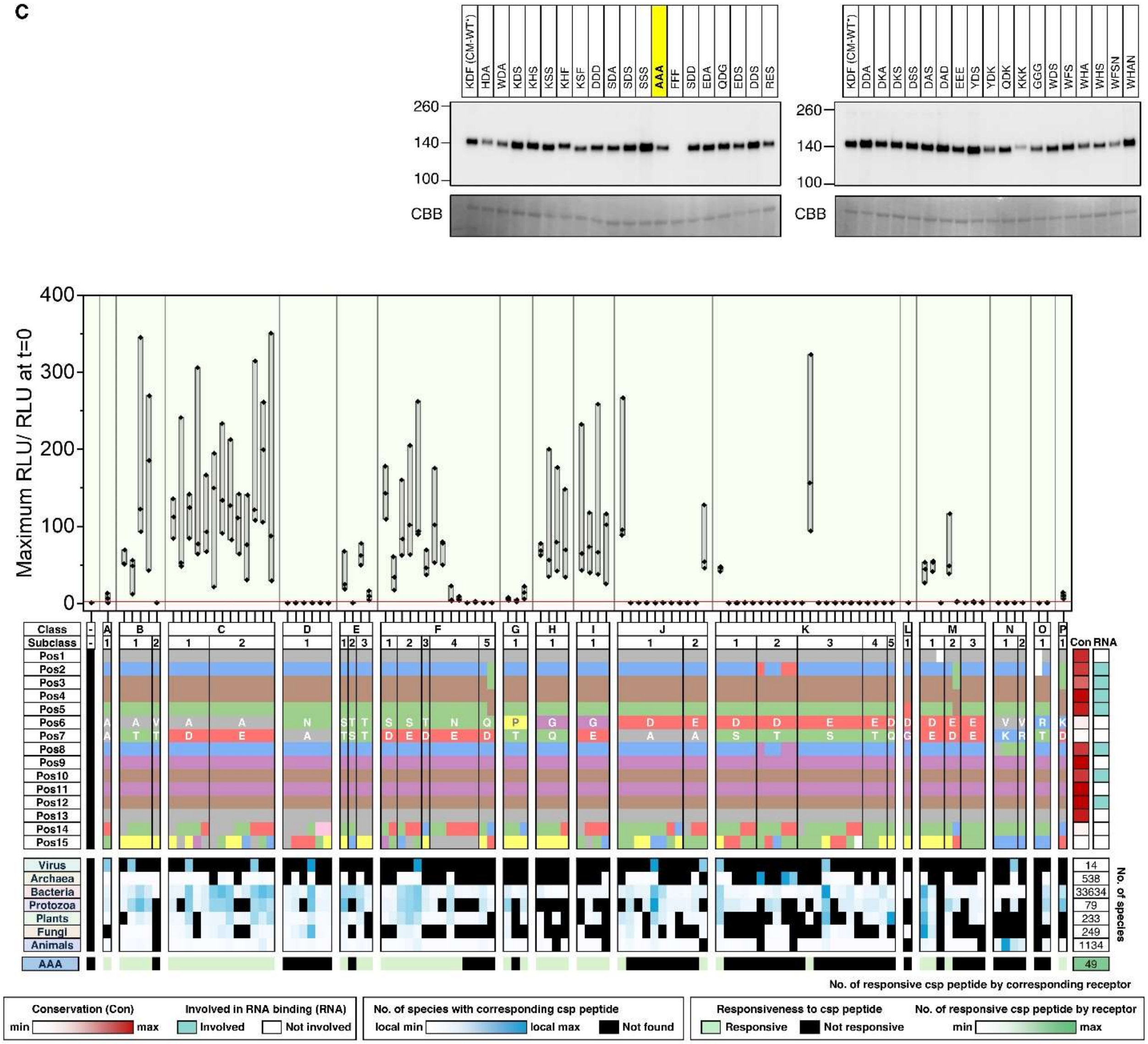
KDF (CM-WT*)
HDA
WDA
KDS
KHS
KSS
KHF
KSF
DDD
SDA
SDS
SSS
AAA
FFF
SDD
EDA
QDG
EDS
DDS
RES
DDA
DKA
DKS
DSS
DAS
DAD
EEE
YDS
YDK
QDK
KKK
GGG
WDS
WFS
WHA
WHS
WFSN
WHAN
260
140
100
CBB
Maximum RLU/ RLU at t=0
400
300
200
100
0
Class
Subclass
Pos1
Pos2
Pos3
Pos4
Pos5
Pos6
Pos7
Pos8
Pos9
Pos10
Pos11
Pos12
Pos13
Pos14
Pos15
Con
RNA
Virus
Archaea
Bacteria
Protozoa
Plants
Fungi
Animals
AAA
14
538
33634
79
233
249
1134
49
No. of species
No. of responsive csp peptide by corresponding receptor
Conservation (Con)
min
max
Involved in RNA binding (RNA)
Involved
Not involved
No. of species with corresponding csp peptide
local min
local max
Not found
Responsiveness to csp peptide
Responsive
Not responsive
No. of responsive csp peptide by receptor
min
max

D

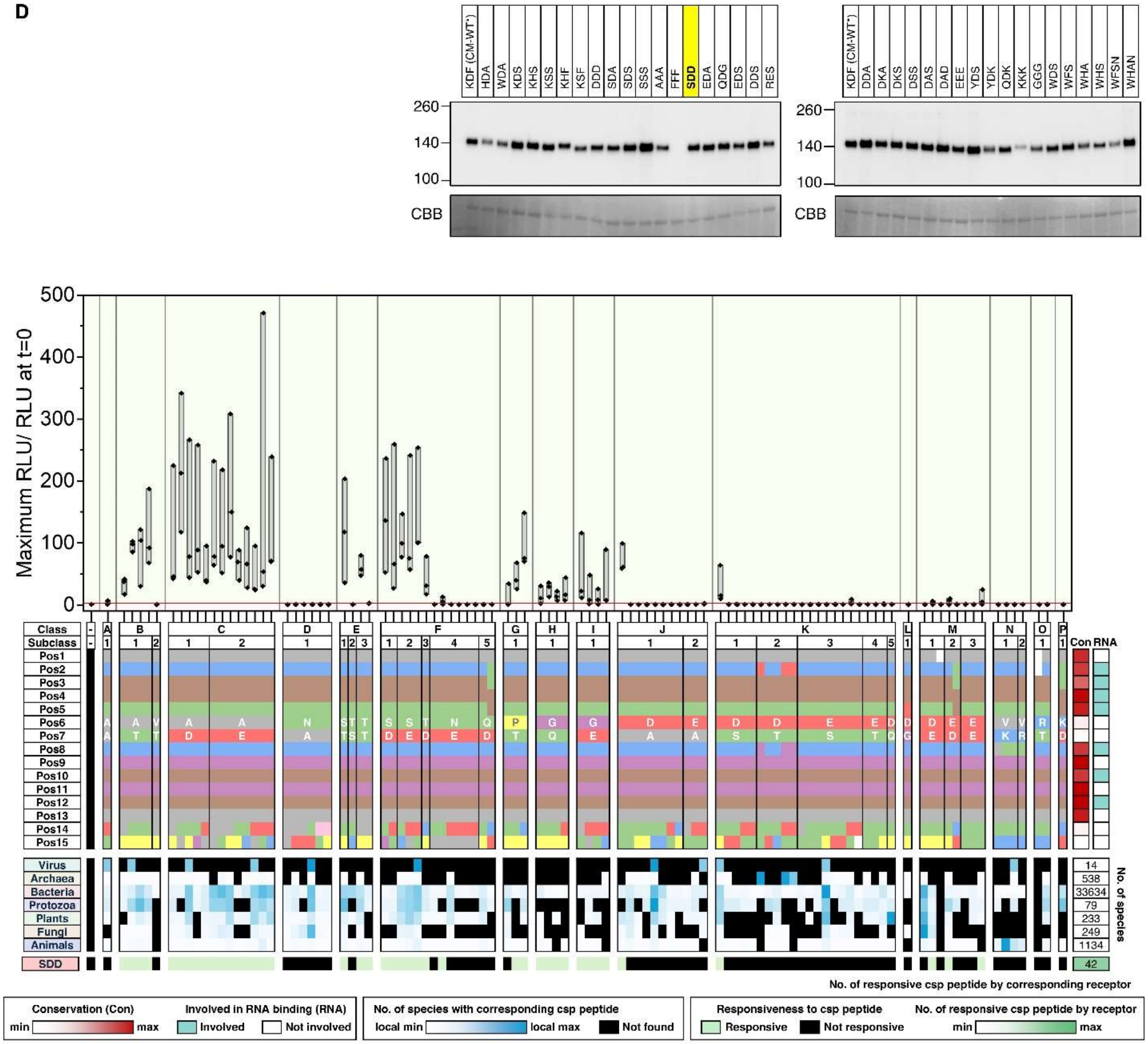

KDF (CM-WT*)
HDA
WDA
KDS
KHS
KSS
KHF
KSF
DDD
SDA
SDS
SSS
AAA
FFF
SDD
EDA
QDG
EDS
DDS
RES
DDA
DKA
DKS
DSS
DAS
DAD
EEE
YDS
YDK
QDK
KKK
GGG
WDS
WFS
WHA
WHS
WFSN
WHAN
260
140
100
CBB
Maximum RLU/ RLU at t=0
500
400
300
200
100
0
Class
Subclass
Pos1
Pos2
Pos3
Pos4
Pos5
Pos6
Pos7
Pos8
Pos9
Pos10
Pos11
Pos12
Pos13
Pos14
Pos15
Con
RNA
Virus
Archaea
Bacteria
Protozoa
Plants
Fungi
Animals
SDD
14
538
33634
79
233
249
1134
No. of species
42
No. of responsive csp peptide by corresponding receptor
Conservation (Con)
min
max
Involved in RNA binding (RNA)
Involved
Not involved
No. of species with corresponding csp peptide
local min
local max
Not found
Responsiveness to csp peptide
Responsive
Not responsive
No. of responsive csp peptide by receptor
min
max

E

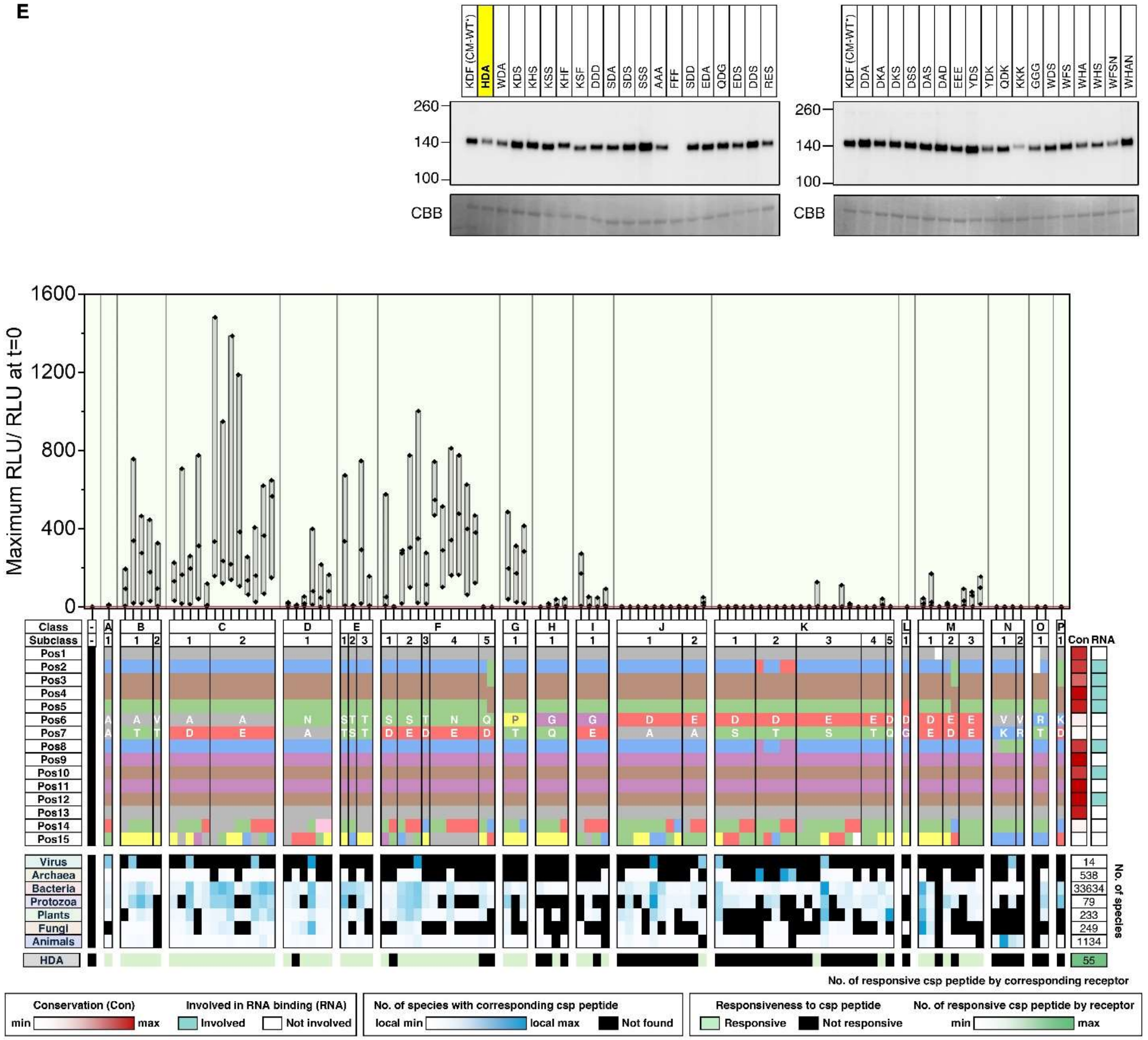
KDF (CM-WT*)
HDA
WDA
KDS
KHS
KSS
KHF
KSF
DDD
SDA
SDS
SSS
AAA
FFF
SDD
EDA
QDG
EDS
DDS
RES
DDA
DKA
DKS
DSS
DAS
DAD
EEE
YDS
YDK
QDK
KKK
GGG
WDS
WFS
WHA
WHS
WFSN
WHAN
260
140
100
CBB
1600
1200
800
400
0
Maximum RLU/ RLU at t=0
Class
Subclass
Pos1
Pos2
Pos3
Pos4
Pos5
Pos6
Pos7
Pos8
Pos9
Pos10
Pos11
Pos12
Pos13
Pos14
Pos15
Con
RNA
Virus
Archaea
Bacteria
Protozoa
Plants
Fungi
Animals
HDA
14
538
33634
79
233
249
1134
55
No. of species
No. of responsive csp peptide by corresponding receptor
Conservation (Con)
min
max
Involved in RNA binding (RNA)
Involved
Not involved
No. of species with corresponding csp peptide
local min
local max
Not found
Responsiveness to csp peptide
Responsive
Not responsive
No. of responsive csp peptide by receptor
min
max

F

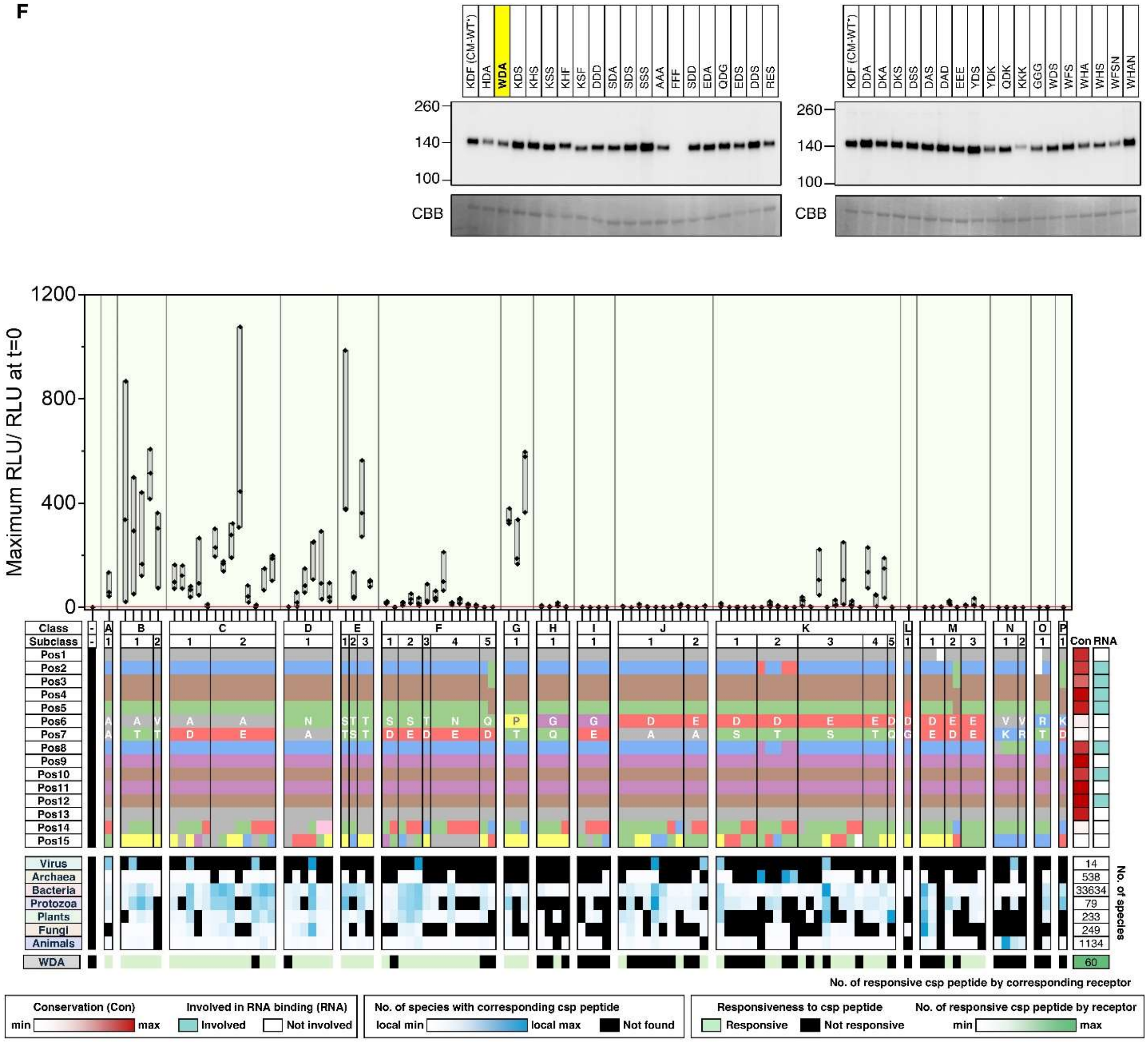
KDF (CM-WT*) HDA WDA KDS KHS KSS KHF KSF DDD SDA SDS SSS AAA FFF SDD EDA QDG EDS DDS RES
KDF (CM-WT*) DDA DKA DKS DSS DAS DAD EEE YDS YDK QDK KKK GGG WDS WFS WHA WHS WFSN WHAN
260
140
100
CBB
Maximum RLU/ RLU at t=0
1200
800
400
0
Class
Subclass
Pos1
Pos2
Pos3
Pos4
Pos5
Pos6
Pos7
Pos8
Pos9
Pos10
Pos11
Pos12
Pos13
Pos14
Pos15
Con
RNA
Virus
Archaea
Bacteria
Protozoa
Plants
Fungi
Animals
WDA
14
538
33634
79
233
249
1134
60
No. of species
No. of responsive csp peptide by corresponding receptor
Conservation (Con)
min
max
Involved in RNA binding (RNA)
Involved
Not involved
No. of species with corresponding csp peptide
local min
local max
Not found
Responsiveness to csp peptide
Responsive
Not responsive
No. of responsive csp peptide by receptor
min
max

G

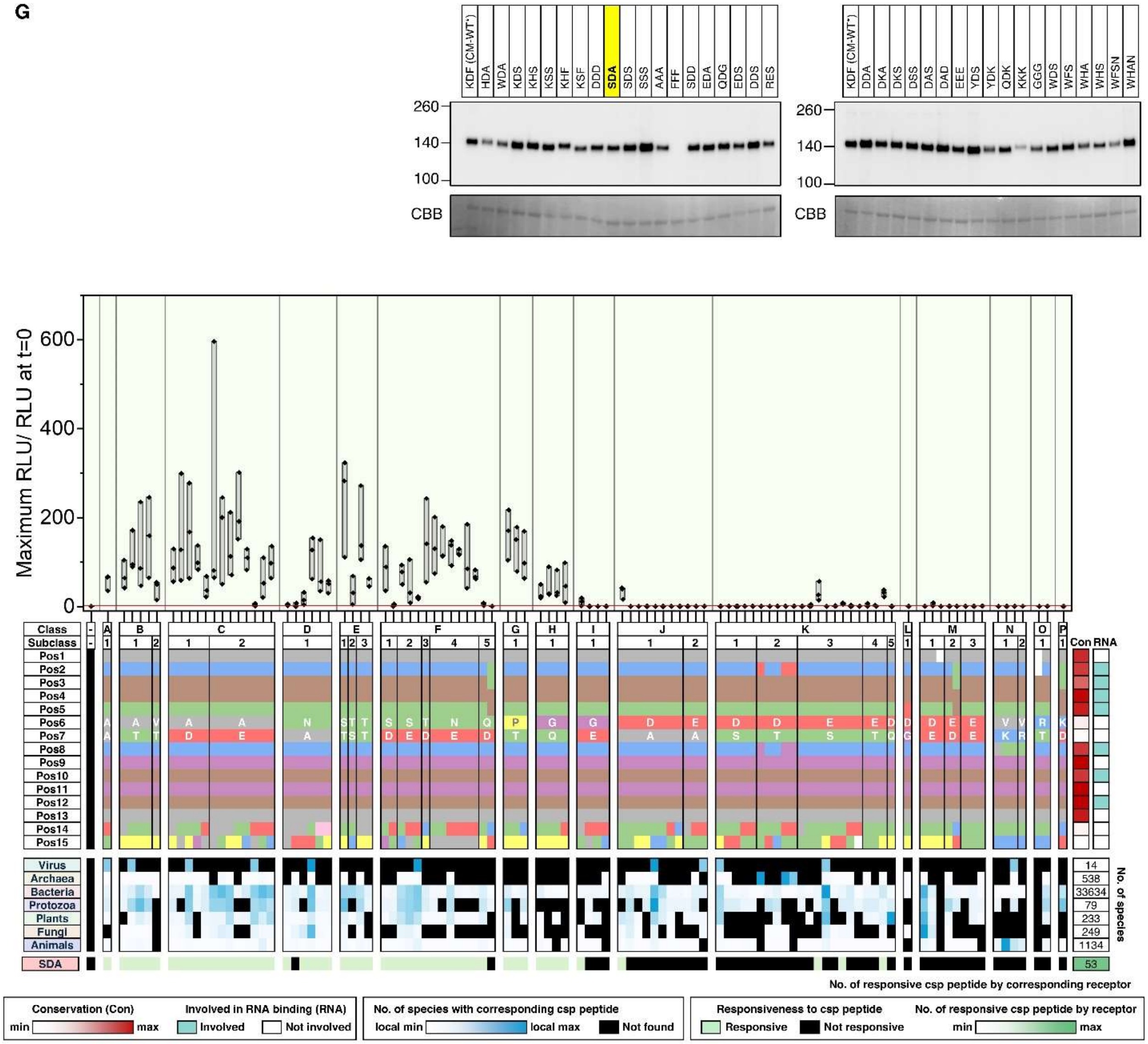
KDF (CM-WT*)
HDA
WDA
KDS
KHS
KSS
KHF
KSF
DDD
SDA
SDS
SSS
AAA
FFF
SDD
EDA
QDG
EDS
DDS
RES
DDA
DKA
DKS
DSS
DAS
DAD
EEE
YDS
YDK
QDK
KKK
GGG
WDS
WFS
WHA
WHS
WFSN
WHAN
260
140
100
CBB
Maximum RLU/ RLU at t=0
600
400
200
0
Class
Subclass
Pos1
Pos2
Pos3
Pos4
Pos5
Pos6
Pos7
Pos8
Pos9
Pos10
Pos11
Pos12
Pos13
Pos14
Pos15
Con
RNA
Virus
Archaea
Bacteria
Protozoa
Plants
Fungi
Animals
SDA
14
538
33634
79
233
249
1134
53
No. of species
No. of responsive csp peptide by corresponding receptor
Conservation (Con)
min
max
Involved in RNA binding (RNA)
Involved
Not involved
No. of species with corresponding csp peptide
local min
local max
Not found
Responsiveness to csp peptide
Responsive
Not responsive
No. of responsive csp peptide by receptor
min
max

H

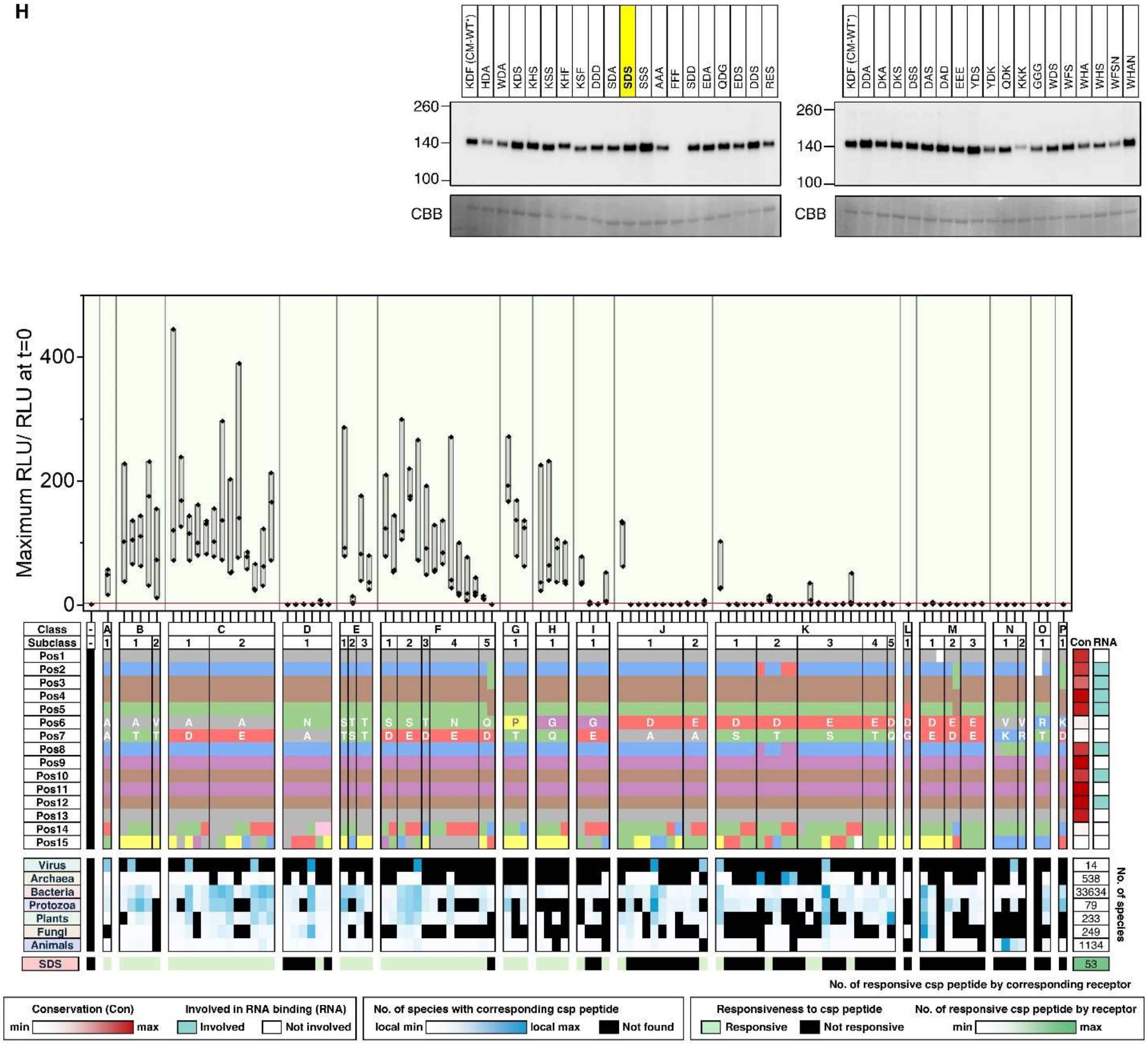
KDF (CM-WT*)
HDA
WDA
KDS
KHS
KSS
KHF
KSF
DDD
SDA
SDS
SSS
AAA
FFF
SDD
EDA
QDG
EDS
DDS
RES
DDA
DKA
DKS
DSS
DAS
DAD
EEE
YDS
YDK
QDK
KKK
GGG
WDS
WFS
WHA
WHS
WFSN
WHAN
260
140
100
CBB
Maximum RLU/ RLU at t=0
400
200
0
Class
Subclass
Pos1
Pos2
Pos3
Pos4
Pos5
Pos6
Pos7
Pos8
Pos9
Pos10
Pos11
Pos12
Pos13
Pos14
Pos15
Con
RNA
Virus
Archaea
Bacteria
Protozoa
Plants
Fungi
Animals
SDS
14
538
33634
79
233
249
1134
53
No. of species
No. of responsive csp peptide by corresponding receptor
Conservation (Con)
min
max
Involved in RNA binding (RNA)
Involved
Not involved
No. of species with corresponding csp peptide
local min
local max
Not found
Responsiveness to csp peptide
Responsive
Not responsive
No. of responsive csp peptide by receptor
min
max

I

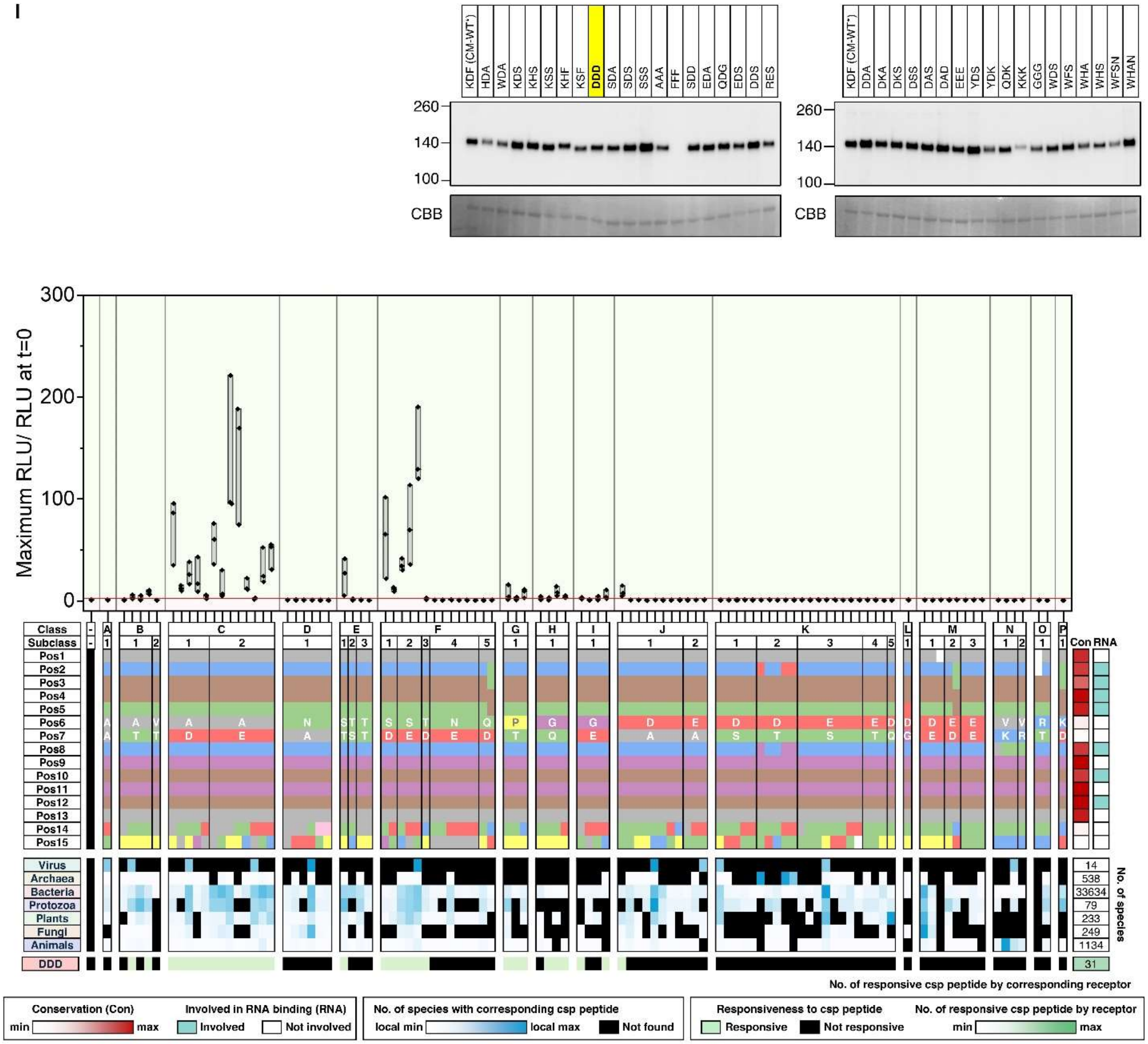

KDF (CM-WT*)
HDA
WDA
KDS
KHS
KSS
KHF
KSF
DDD
SDA
SDS
SSS
AAA
FFF
SDD
EDA
QDG
EDS
DDS
RES
DDA
DKA
DKS
DSS
DAS
DAD
EEE
YDS
YDK
QDK
KKK
GGG
WDS
WFS
WHA
WHS
WFSN
WHAN
260
140
100
CBB
Maximum RLU/ RLU at t=0
300
200
100
0
Class
Subclass
Pos1
Pos2
Pos3
Pos4
Pos5
Pos6
Pos7
Pos8
Pos9
Pos10
Pos11
Pos12
Pos13
Pos14
Pos15
Con
RNA
Virus
Archaea
Bacteria
Protozoa
Plants
Fungi
Animals
DDD
14
538
33634
79
233
249
1134
31
No. of species
No. of responsive csp peptide by corresponding receptor
Conservation (Con)
min
max
Involved in RNA binding (RNA)
Involved
Not involved
No. of species with corresponding csp peptide
local min
local max
Not found
Responsiveness to csp peptide
Responsive
Not responsive
No. of responsive csp peptide by receptor
min
max

**J**

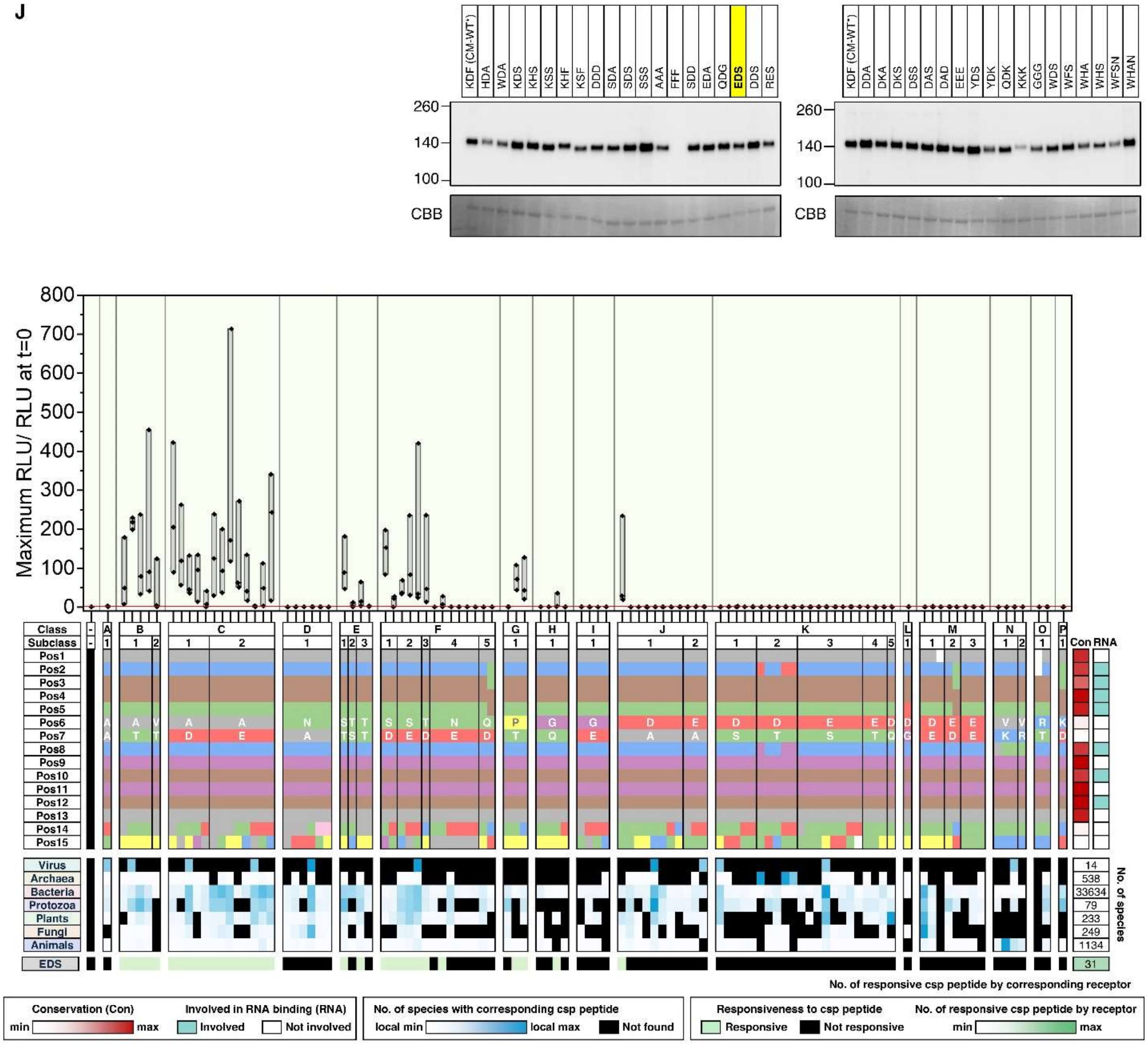
KDF (CM-WT*)
HDA
WDA
KDS
KHS
KSS
KHF
KSF
DDD
SDA
SDS
SSS
AAA
FFF
SDD
EDA
QDG
EDS
DDS
RES
DDA
DKA
DKS
DSS
DAS
DAD
EEE
YDS
YDK
QDK
KKK
GGG
WDS
WFS
WHA
WHS
WFSN
WHAN
260
140
100
CBB
Maximum RLU/ RLU at t=0
800
700
600
500
400
300
200
100
0
Class
Subclass
Virus
Archaea
Bacteria
Protozoa
Plants
Fungi
Animals
EDS
Con
RNA
No. of species
14
538
33634
79
233
249
1134
31
No. of responsive csp peptide by corresponding receptor
Conservation (Con)
min
max
Involved in RNA binding (RNA)
Involved
Not involved
No. of species with corresponding csp peptide
local min
local max
Not found
Responsiveness to csp peptide
Responsive
Not responsive
No. of responsive csp peptide by receptor
min
max

K

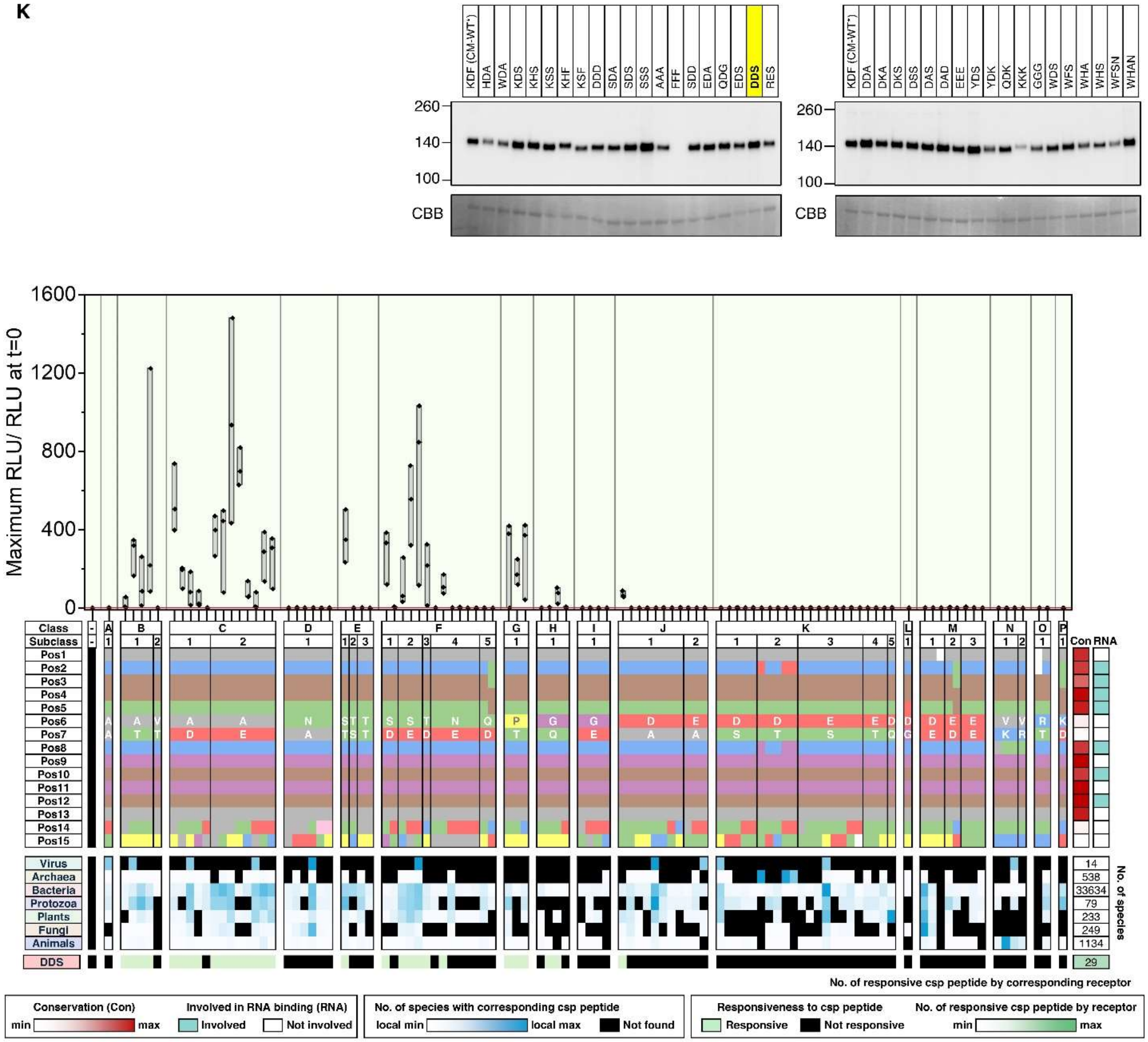
KDF (CM-WT*)
HDA
WDA
KDS
KHS
KSS
KHF
KSF
DDD
SDA
SDS
SSS
AAA
FFF
SDD
EDA
QDG
EDS
DDS
RES
DDA
DKA
DKS
DSS
DAS
DAD
EEE
YDS
YDK
QDK
KKK
GGG
WDS
WFS
WHA
WHS
WFSN
WHAN
260
140
100
CBB
Maximum RLU/ RLU at t=0
1600
1200
800
400
0
Class
Subclass
Virus
Archaea
Bacteria
Protozoa
Plants
Fungi
Animals
DDS
Con
RNA
No. of species
14
538
33634
79
233
249
1134
29
No. of responsive csp peptide by corresponding receptor
Conservation (Con)
min
max
Involved in RNA binding (RNA)
Involved
Not involved
No. of species with corresponding csp peptide
local min
local max
Not found
Responsiveness to csp peptide
Responsive
Not responsive
No. of responsive csp peptide by receptor
min
max

**L**

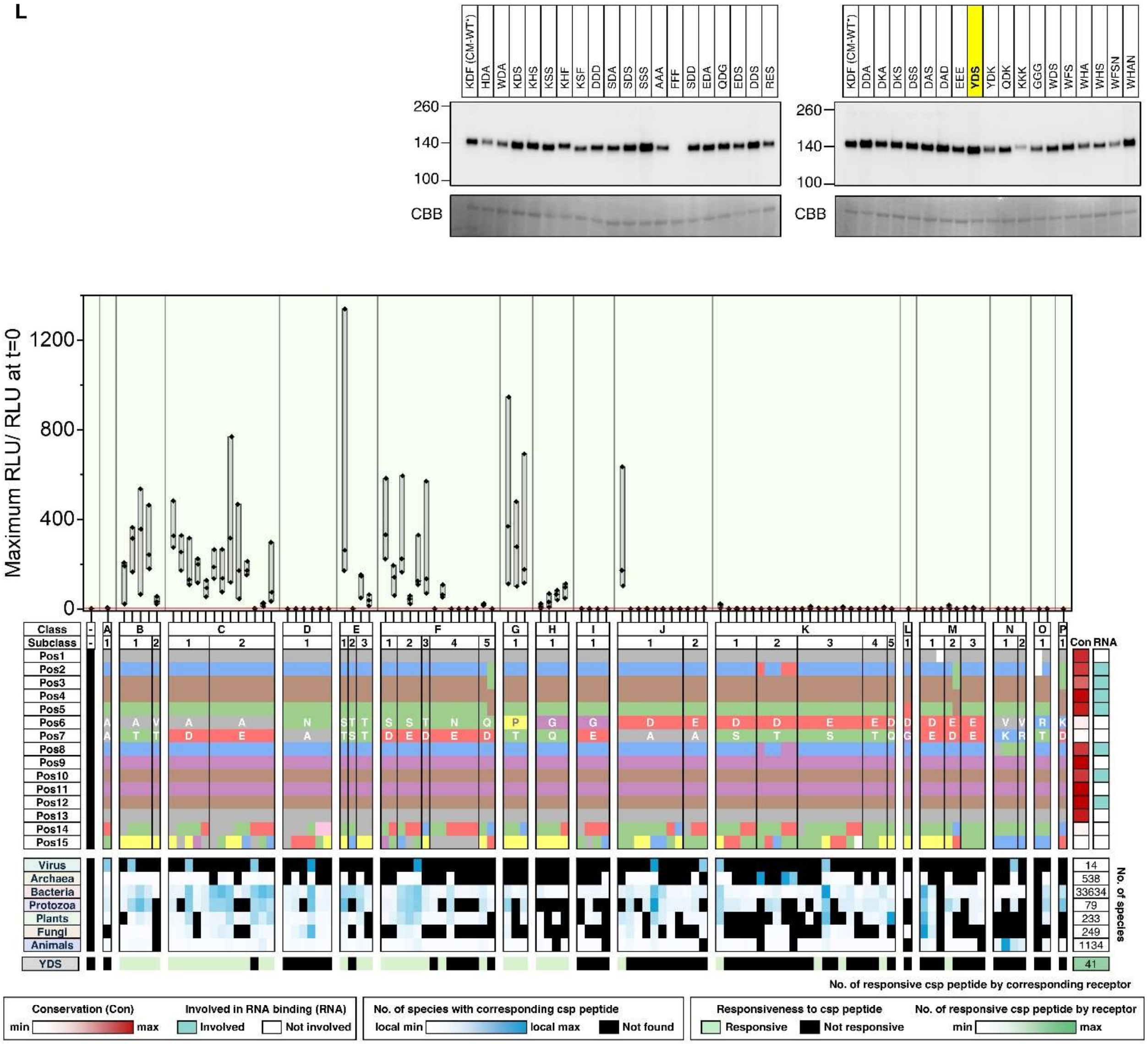
KDF (CM-WT*)
HDA
WDA
KDS
KHS
KSS
KHF
KSF
DDD
SDA
SDS
SSS
AAA
FFF
SDD
EDA
QDG
EDS
DDS
RES
DDA
DKA
DKS
DSS
DAS
DAD
EEE
YDS
YDK
QDK
KKK
GGG
WDS
WFS
WHA
WHS
WFSN
WHAN
260
140
100
CBB
Maximum RLU/ RLU at t=0
1200
800
400
0
Class
Subclass
Pos1
Pos2
Pos3
Pos4
Pos5
Pos6
Pos7
Pos8
Pos9
Pos10
Pos11
Pos12
Pos13
Pos14
Pos15
Con
RNA
Virus
Archaea
Bacteria
Protozoa
Plants
Fungi
Animals
YDS
No. of species
14
538
33634
79
233
249
1134
41
No. of responsive csp peptide by corresponding receptor
Conservation (Con)
min
max
Involved in RNA binding (RNA)
Involved
Not involved
No. of species with corresponding csp peptide
local min
local max
Not found
Responsiveness to csp peptide
Responsive
Not responsive
No. of responsive csp peptide by receptor
min
max

M

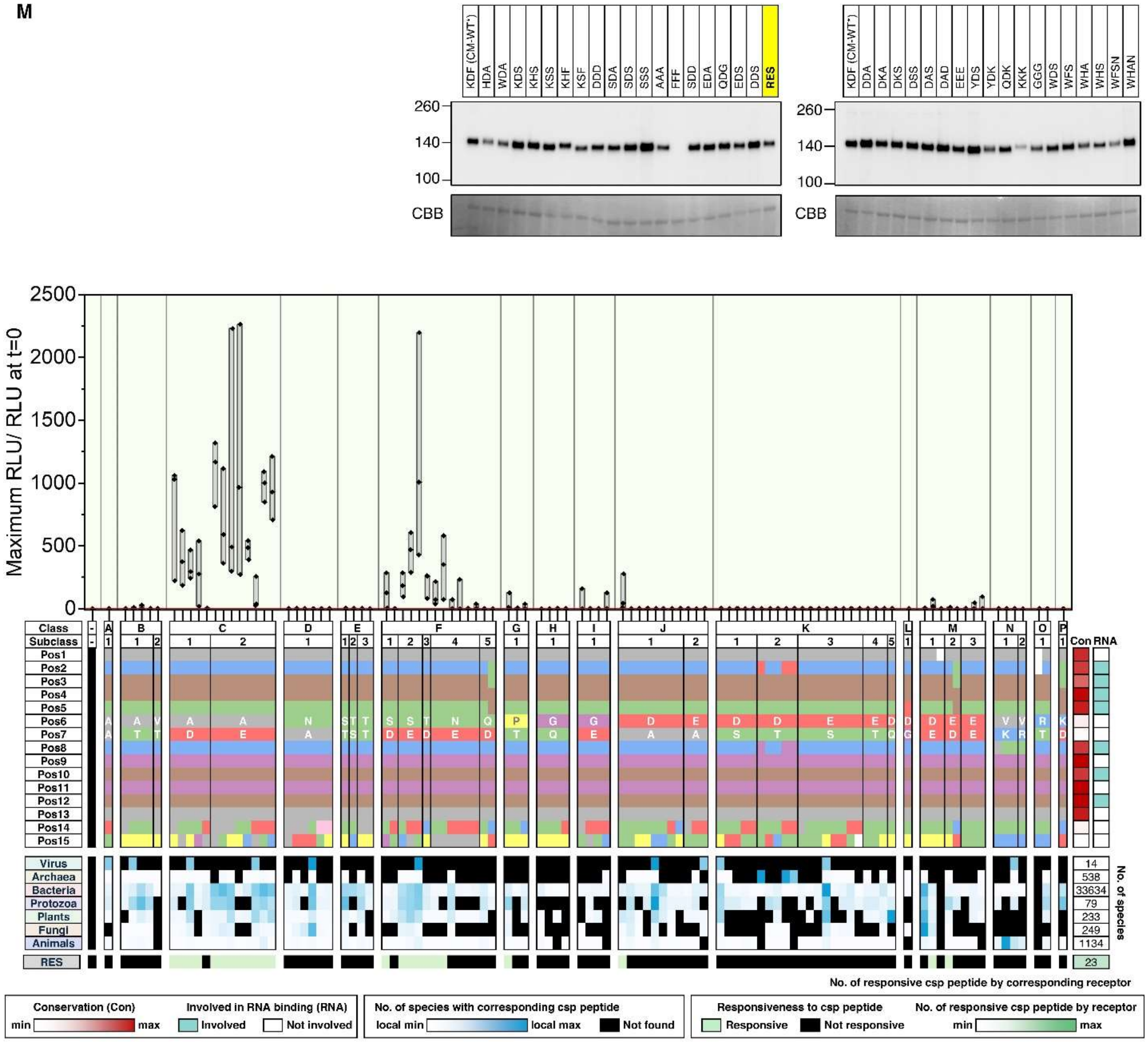
KDF (CM-WT')
RES
CBB
Maximum RLU/ RLU at t=0
Class
Subclass
Virus
Archaea
Bacteria
Protozoa
Plants
Fungi
Animals
No. of species
No. of responsive csp peptide by corresponding receptor
Conservation (Con)
Involved in RNA binding (RNA)
Involved
Not involved
No. of species with corresponding csp peptide
local min
local max
Not found
Responsiveness to csp peptide
Responsive
Not responsive
No. of responsive csp peptide by receptor

**N**

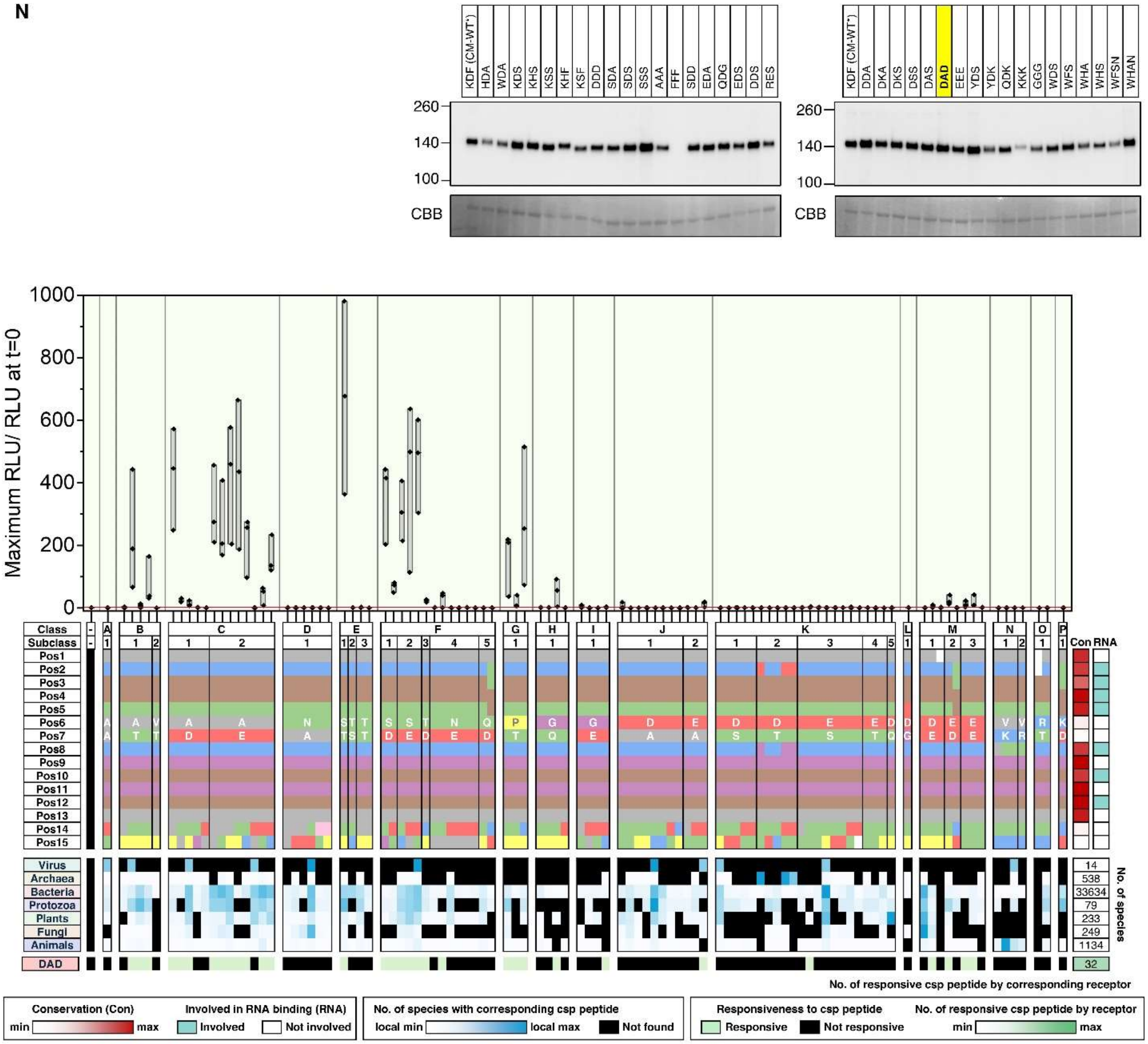

KDF (CM-WT') HDA WDA KDS KHS KSS KHF KSF DDD SDA SDS SSS AAA FFF SDD EDA QDG EDS DDS RES
KDF (CM-WT') DDA DKA DKS DSS DAS DAD EEE YDS YDK QDK KKK GGG WDS WFS WHA WHS WFSN WHAN
260
140
100
CBB
Maximum RLU/ RLU at t=0
Class
Subclass
Con
RNA
Virus
Archaea
Bacteria
Protozoa
Plants
Fungi
Animals
DAD
No. of species
No. of responsive csp peptide by corresponding receptor
Conservation (Con)
Involved in RNA binding (RNA)
min
max
Involved
Not involved
No. of species with corresponding csp peptide
local min
local max
Not found
Responsiveness to csp peptide
Responsive
Not responsive
No. of responsive csp peptide by receptor

o

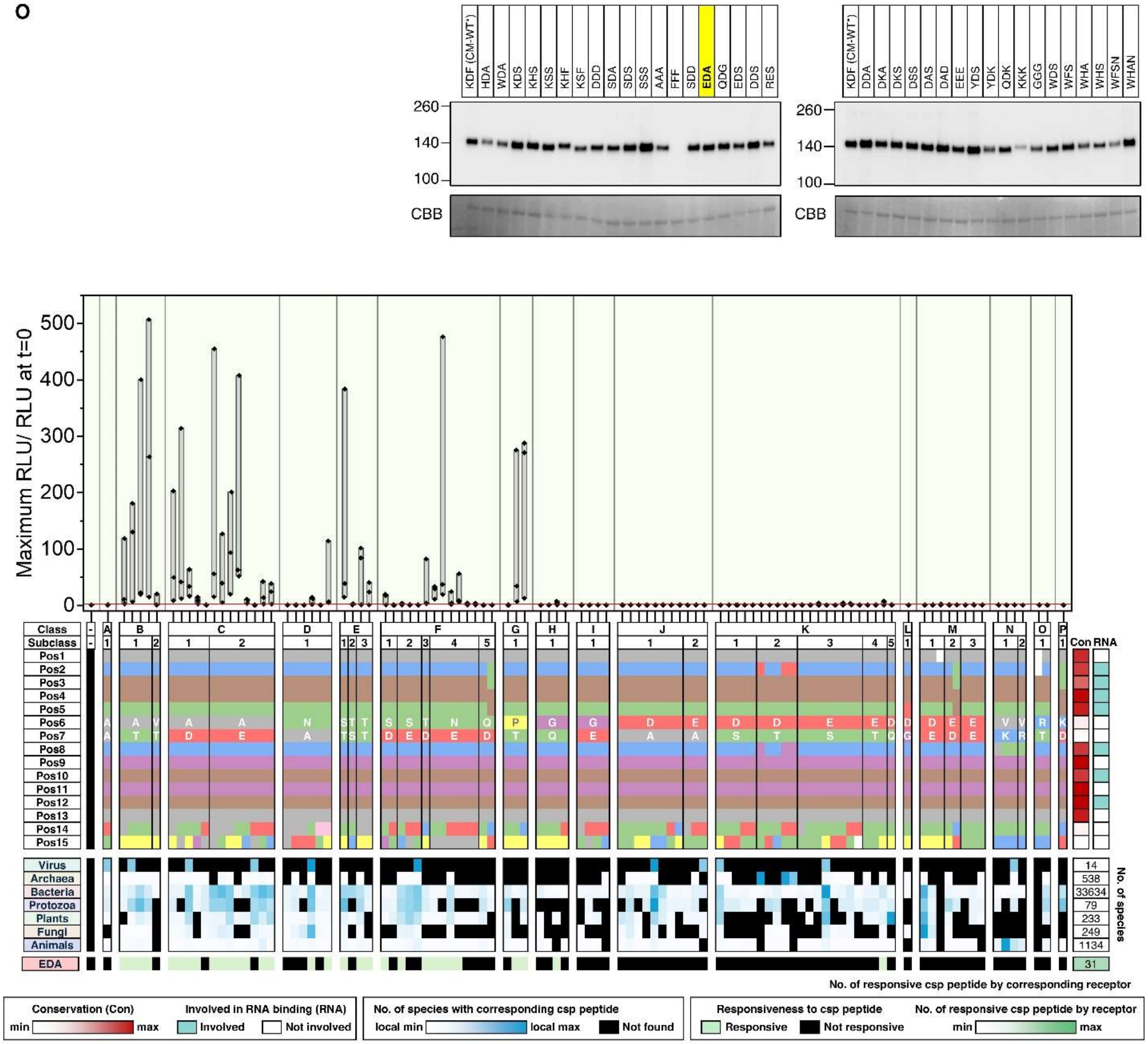
KDF (CM-WT')
HDA
WDA
KDS
KHS
KSS
KHF
KSF
DDD
SDA
SDS
SSS
AAA
FFF
SDD
EDA
QDG
EDS
DDS
RES
DDA
DKA
DKS
DSS
DAS
DAD
EEE
YDS
YDK
QDK
KKK
GGG
WDS
WFS
WHA
WHS
WFSN
WHAN
260
140
100
CBB
Maximum RLU/ RLU at t=0
Class
Subclass
Con
RNA
Virus
Archaea
Bacteria
Protozoa
Plants
Fungi
Animals
EDA
No. of species
14
538
33634
79
233
249
1134
31
No. of responsive csp peptide by corresponding receptor
Conservation (Con)
min
max
Involved in RNA binding (RNA)
Involved
Not involved
No. of species with corresponding csp peptide
local min
local max
Not found
Responsiveness to csp peptide
Responsive
Not responsive
No. of responsive csp peptide by receptor
min
max

**P**

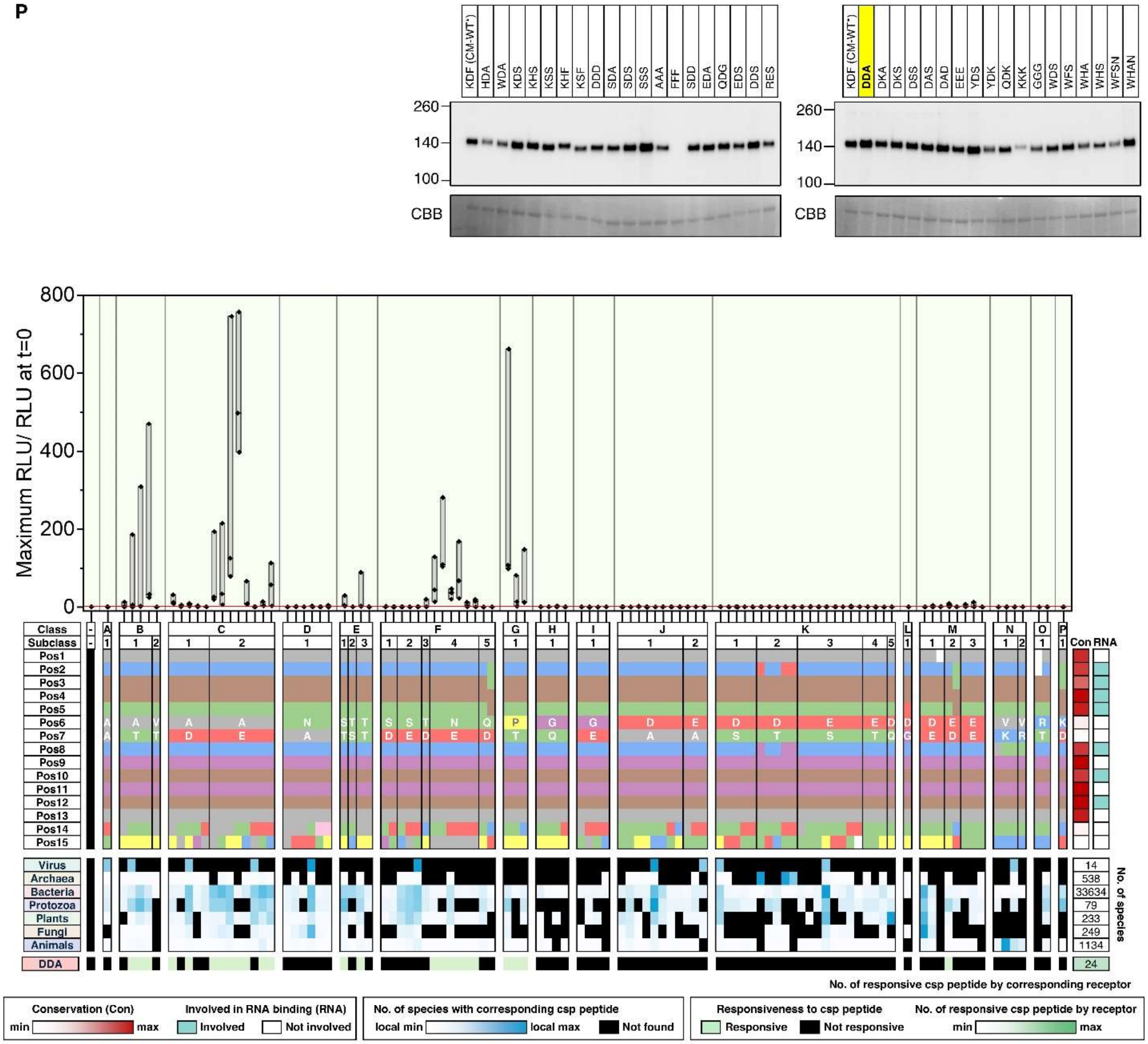

KDF (CM-WT*)
HDA
WDA
KDS
KHS
KSS
KHF
KSF
DDD
SDA
SDS
SSS
AAA
FFF
SDD
EDA
QDG
EDS
DDS
RES
DDA
DKA
DKS
DSS
DAS
DAD
EEE
YDS
YDK
QDK
KKK
GGG
WDS
WFS
WHA
WHS
WFSN
WHAN
260
140
100
CBB
Maximum RLU/ RLU at t=0
800
600
400
200
0
Class
Subclass
Pos1
Pos2
Pos3
Pos4
Pos5
Pos6
Pos7
Pos8
Pos9
Pos10
Pos11
Pos12
Pos13
Pos14
Pos15
Con
RNA
Virus
Archaea
Bacteria
Protozoa
Plants
Fungi
Animals
DDA
No. of species
14
538
33634
79
233
249
1134
24
No. of responsive csp peptide by corresponding receptor
Conservation (Con)
min
max
Involved in RNA binding (RNA)
Involved
Not involved
No. of species with corresponding csp peptide
local min
local max
Not found
Responsiveness to csp peptide
Responsive
Not responsive
No. of responsive csp peptide by receptor
min
max

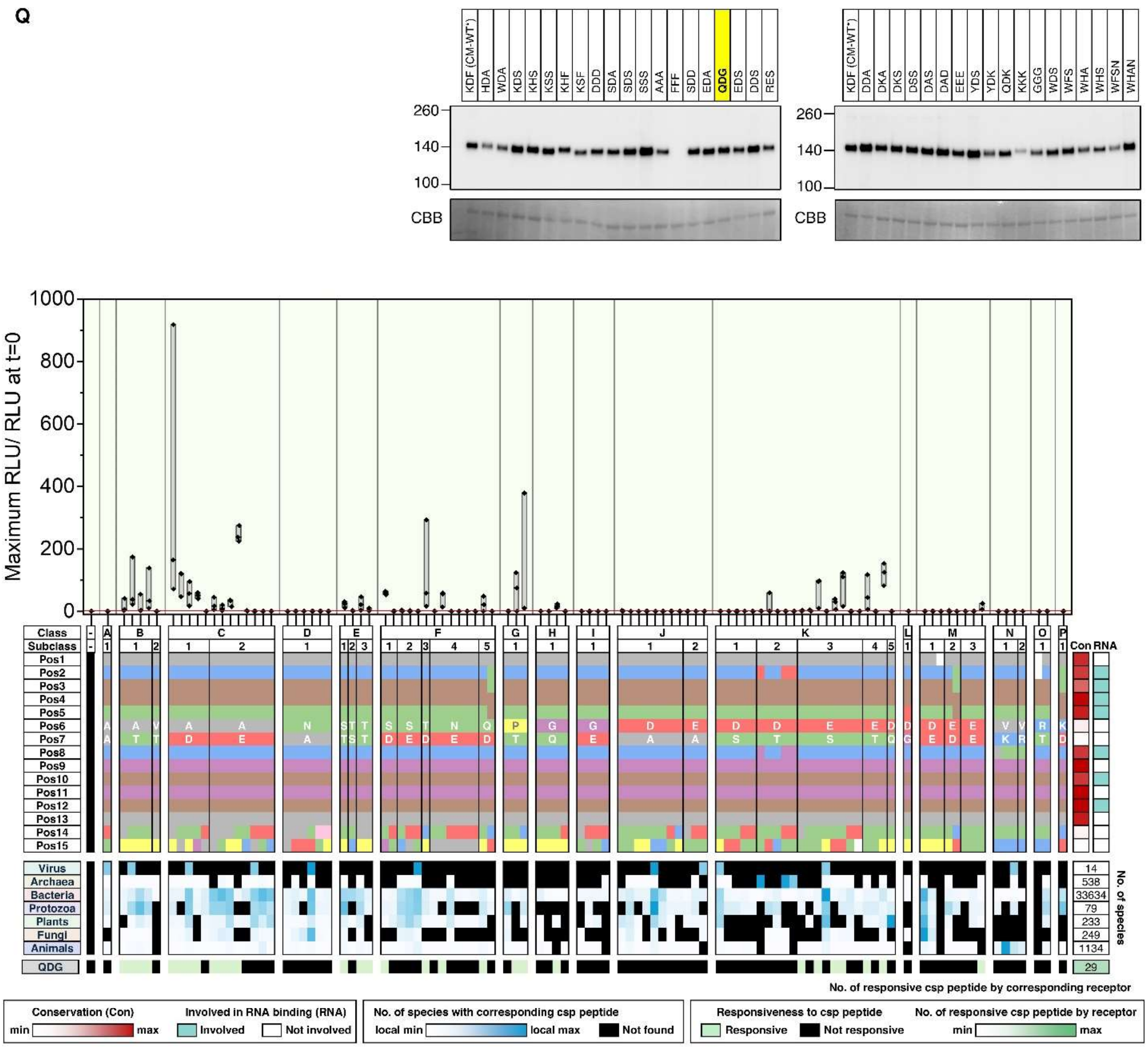
Q
KDF (CM-WT*)
HDA
WDA
KDS
KHS
KSS
KHF
KSF
DDD
SDA
SDS
SSS
AAA
FFF
SDD
EDA
QDG
EDS
DDS
RES
DDA
DKA
DKS
DSS
DAS
DAD
EEE
YDS
YDK
QDK
KKK
GGG
WDS
WFS
WHA
WHS
WFSN
WHAN
260
140
100
CBB
Maximum RLU/ RLU at t=0
Class
Subclass
Virus
Archaea
Bacteria
Protozoa
Plants
Fungi
Animals
QDG
No. of species
No. of responsive csp peptide by corresponding receptor
Conservation (Con)
Involved in RNA binding (RNA)
No. of species with corresponding csp peptide
Responsiveness to csp peptide
No. of responsive csp peptide by receptor

R

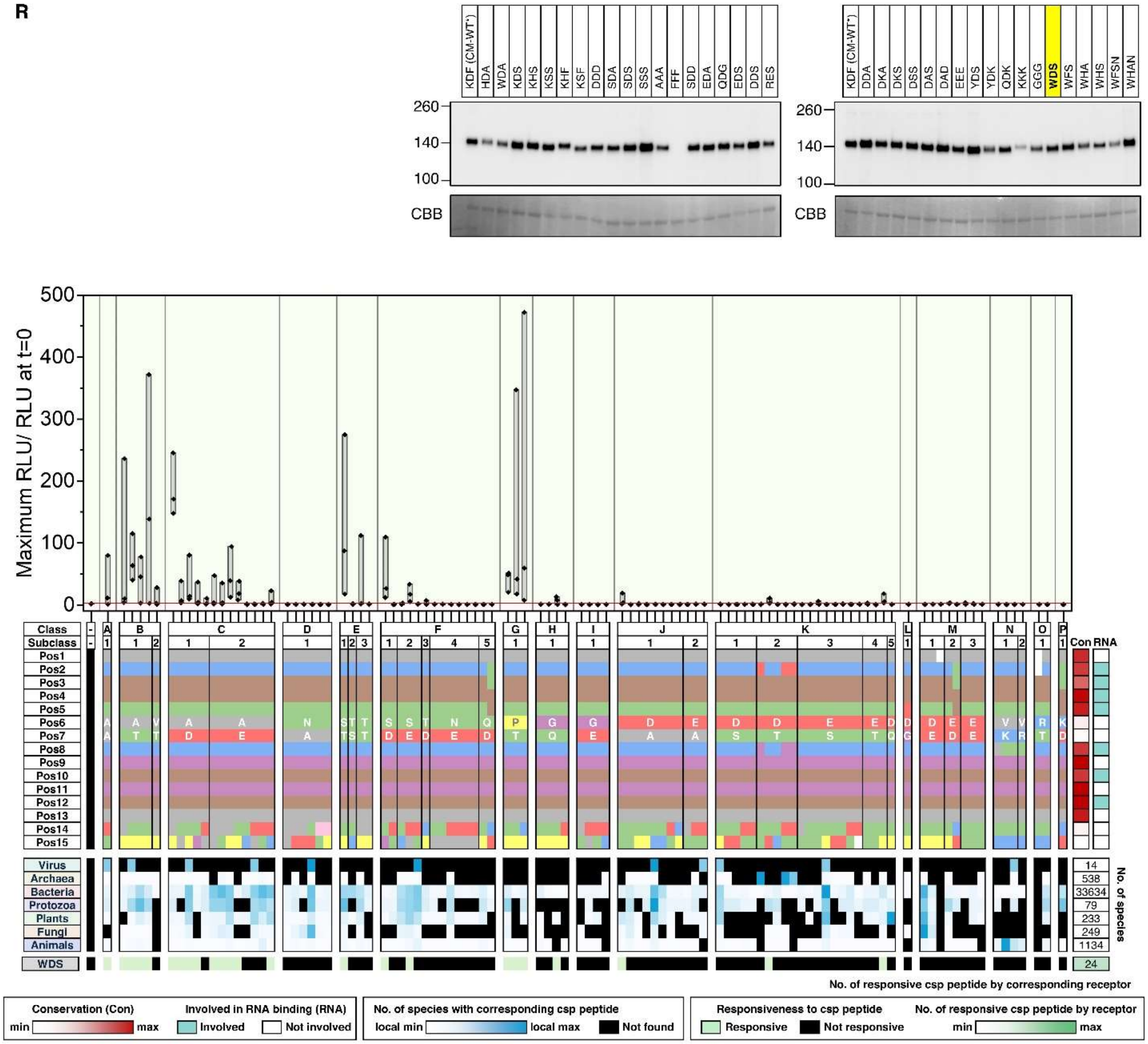

KDF (CM-WT*)
HDA
WDA
KDS
KHS
KSS
KHF
KSF
DDD
SDA
SDS
SSS
AAA
FFF
SDD
EDA
QDG
EDS
DDS
RES
DDA
DKA
DKS
DSS
DAS
DAD
EEE
YDS
YDK
QDK
KKK
GGG
WDS
WFS
WHA
WHS
WFSN
WHAN
260
140
100
CBB
Maximum RLU/ RLU at t=0
500
400
300
200
100
0
Class
Subclass
Pos1
Pos2
Pos3
Pos4
Pos5
Pos6
Pos7
Pos8
Pos9
Pos10
Pos11
Pos12
Pos13
Pos14
Pos15
Con
RNA
Virus
Archaea
Bacteria
Protozoa
Plants
Fungi
Animals
WDS
14
538
33634
79
233
249
1134
24
No. of species
No. of responsive csp peptide by corresponding receptor
Conservation (Con)
min
max
Involved in RNA binding (RNA)
Involved
Not involved
No. of species with corresponding csp peptide
local min
local max
Not found
Responsiveness to csp peptide
Responsive
Not responsive
No. of responsive csp peptide by receptor
min
max

S

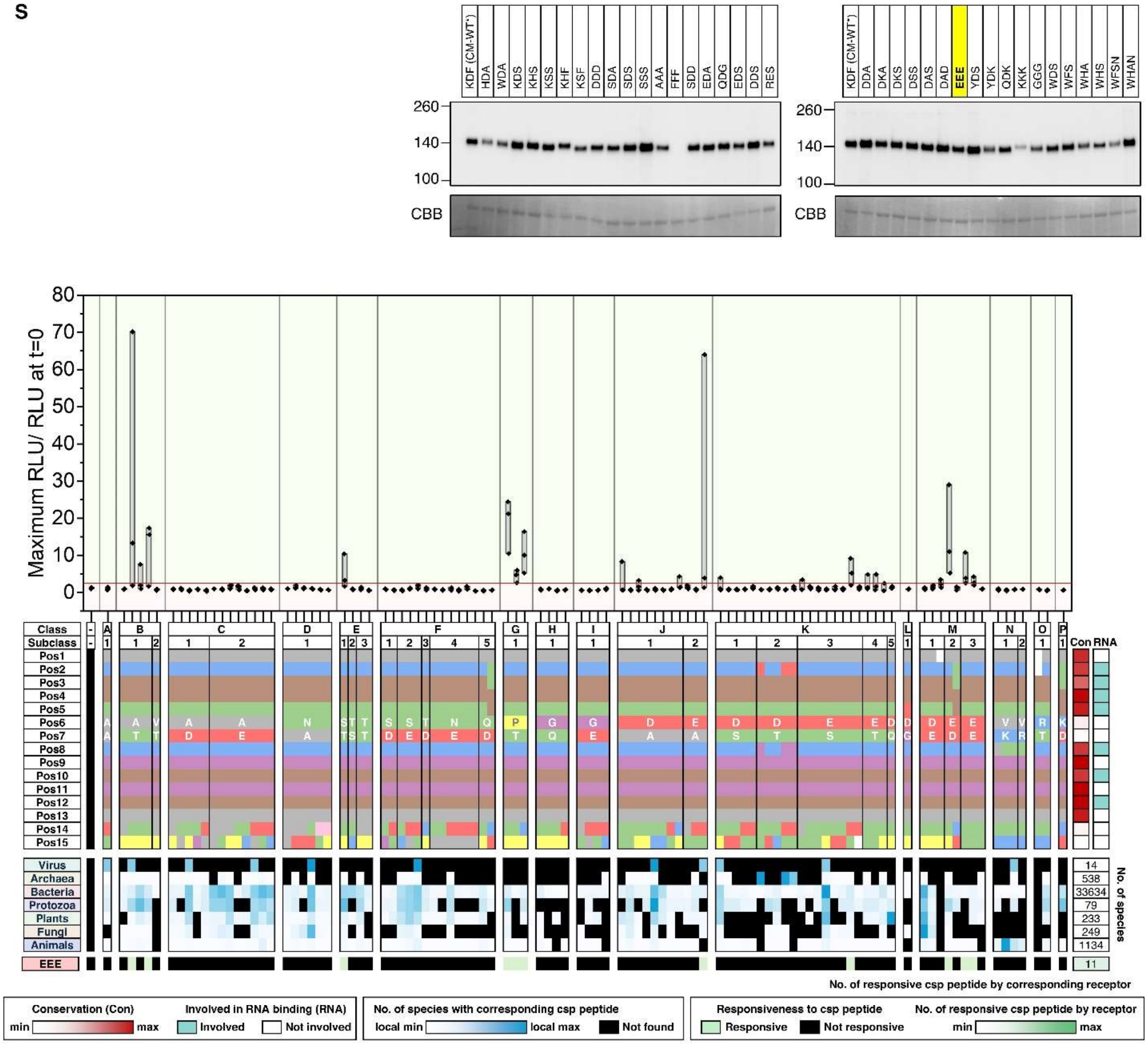
KDF (CM-WT*)
HDA
WDA
KDS
KHS
KSS
KHF
KSF
DDD
SDA
SDS
SSS
AAA
FFF
SDD
EDA
QDG
EDS
DDS
RES
260
140
100
CBB
KDF (CM-WT*)
DDA
DKA
DKS
DSS
DAS
DAD
EEE
YDS
YDK
QDK
KKK
GGG
WDS
WFS
WHA
WHS
WFSN
WHAN
Maximum RLU/ RLU at t=0
80
70
60
50
40
30
20
10
0
Class
Subclass
Pos1
Pos2
Pos3
Pos4
Pos5
Pos6
Pos7
Pos8
Pos9
Pos10
Pos11
Pos12
Pos13
Pos14
Pos15
Con
RNA
Virus
Archaea
Bacteria
Protozoa
Plants
Fungi
Animals
EEE
No. of species
14
538
33634
79
233
249
1134
11
No. of responsive csp peptide by corresponding receptor
Conservation (Con)
min
max
Involved in RNA binding (RNA)
Involved
Not involved
No. of species with corresponding csp peptide
local min
local max
Not found
Responsiveness to csp peptide
Responsive
Not responsive
No. of responsive csp peptide by receptor
min
max

**T**

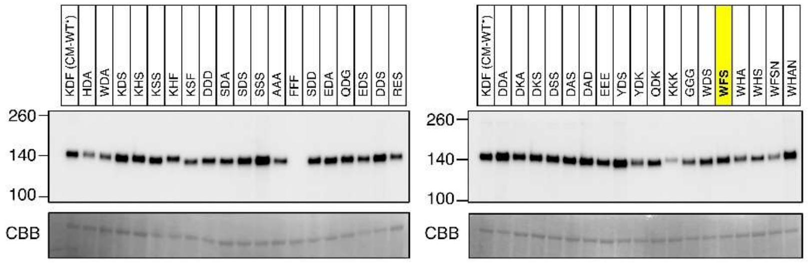

KDF (CM-WT*)
HDA
WDA
KDS
KHS
KSS
KHF
KSF
DDD
SDA
SDS
SSS
AAA
FFF
SDD
EDA
QDG
EDS
DDS
RES
260
140
100
CBB
KDF (CM-WT*)
DDA
DKA
DKS
DSS
DAS
DAD
EEE
YDS
YDK
QDK
KKK
GGG
WDS
WFS
WHA
WHS
WFSN
WHAN
260
140
100
CBB


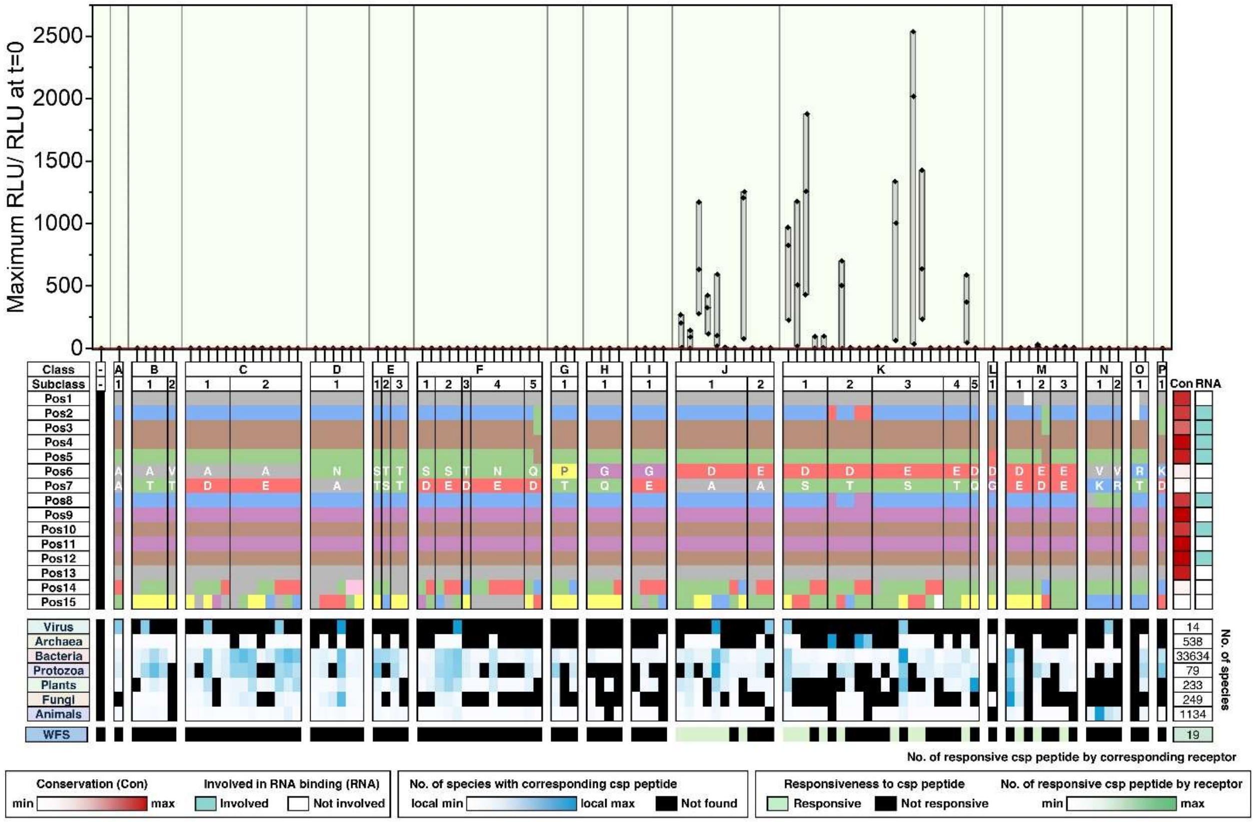

Maximum RLU/ RLU at t=0
2500
2000
1500
1000
500
0
Class
Subclass
Pos1
Pos2
Pos3
Pos4
Pos5
Pos6
Pos7
Pos8
Pos9
Pos10
Pos11
Pos12
Pos13
Pos14
Pos15
Con
RNA
Virus
Archaea
Bacteria
Protozoa
Plants
Fungi
Animals
WFS
14
538
33634
79
233
249
1134
19
No. of species
No. of responsive csp peptide by corresponding receptor
Conservation (Con)
min
max
Involved in RNA binding (RNA)
Involved
Not involved
No. of species with corresponding csp peptide
local min
local max
Not found
Responsiveness to csp peptide
Responsive
Not responsive
No. of responsive csp peptide by receptor
min
max

U

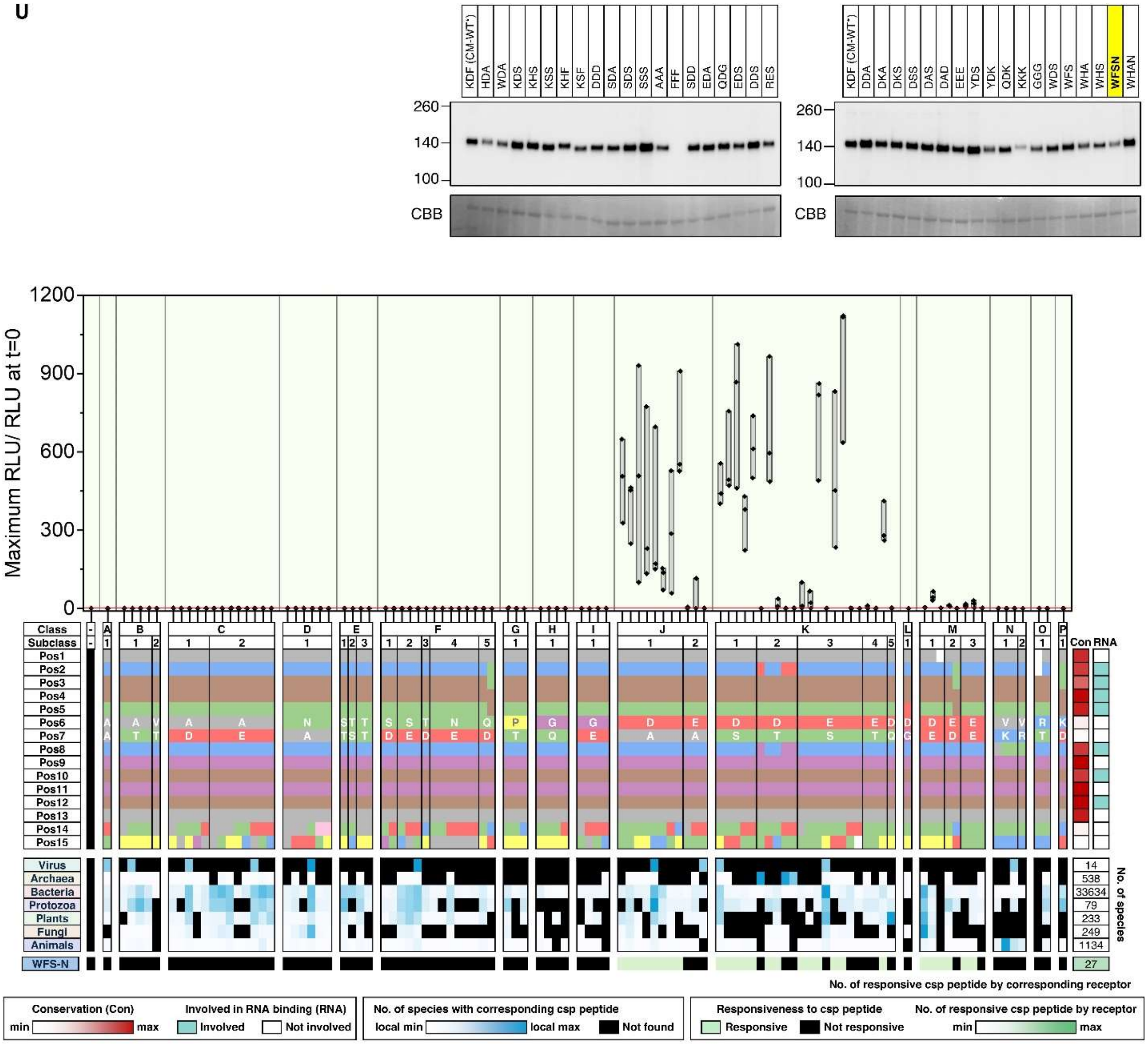
KDF (CM-WT*)
HDA
WDA
KDS
KHS
KSS
KHF
KSF
DDD
SDA
SDS
SSS
AAA
FFF
SDD
EDA
QDG
EDS
DDS
RES
DDA
DKA
DKS
DSS
DAS
DAD
EEE
YDS
YDK
QDK
KKK
GGG
WDS
WFS
WHA
WHS
WFSN
WHAN
260
140
100
CBB
Maximum RLU/ RLU at t=0
1200
900
600
300
0
Class
Subclass
Pos1
Pos2
Pos3
Pos4
Pos5
Pos6
Pos7
Pos8
Pos9
Pos10
Pos11
Pos12
Pos13
Pos14
Pos15
Con
RNA
Virus
Archaea
Bacteria
Protozoa
Plants
Fungi
Animals
WFS-N
14
538
33634
79
233
249
1134
27
No. of species
No. of responsive csp peptide by corresponding receptor
Conservation (Con)
min
max
Involved in RNA binding (RNA)
Involved
Not involved
No. of species with corresponding csp peptide
local min
local max
Not found
Responsiveness to csp peptide
Responsive
Not responsive
No. of responsive csp peptide by receptor
min
max

V

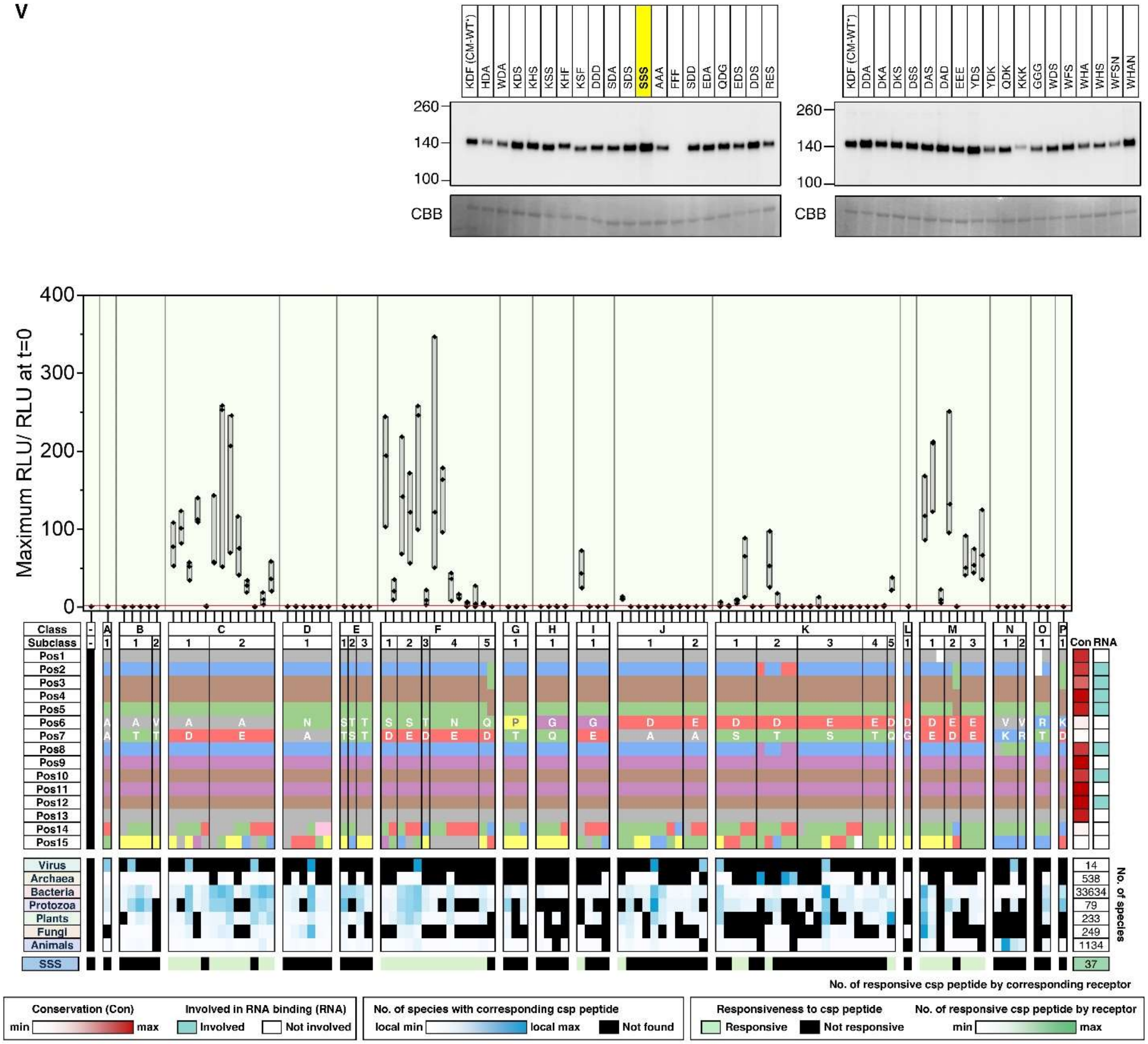
KDF (CM-WT*)
HDA
WDA
KDS
KHS
KSS
KHF
KSF
DDD
SDA
SDS
SSS
AAA
FFF
SDD
EDA
QDG
EDS
DDS
RES
DDA
DKA
DKS
DSS
DAS
DAD
EEE
YDS
YDK
QDK
KKK
GGG
WDS
WFS
WHA
WHS
WFSN
WHAN
260
140
100
CBB
Maximum RLU/ RLU at t=0
400
300
200
100
0
Class
Subclass
Pos1
Pos2
Pos3
Pos4
Pos5
Pos6
Pos7
Pos8
Pos9
Pos10
Pos11
Pos12
Pos13
Pos14
Pos15
Con
RNA
Virus
Archaea
Bacteria
Protozoa
Plants
Fungi
Animals
SSS
14
538
33634
79
233
249
1134
37
No. of species
No. of responsive csp peptide by corresponding receptor
Conservation (Con)
min
max
Involved in RNA binding (RNA)
Involved
Not involved
No. of species with corresponding csp peptide
local min
local max
Not found
Responsiveness to csp peptide
Responsive
Not responsive
No. of responsive csp peptide by receptor
min
max

W

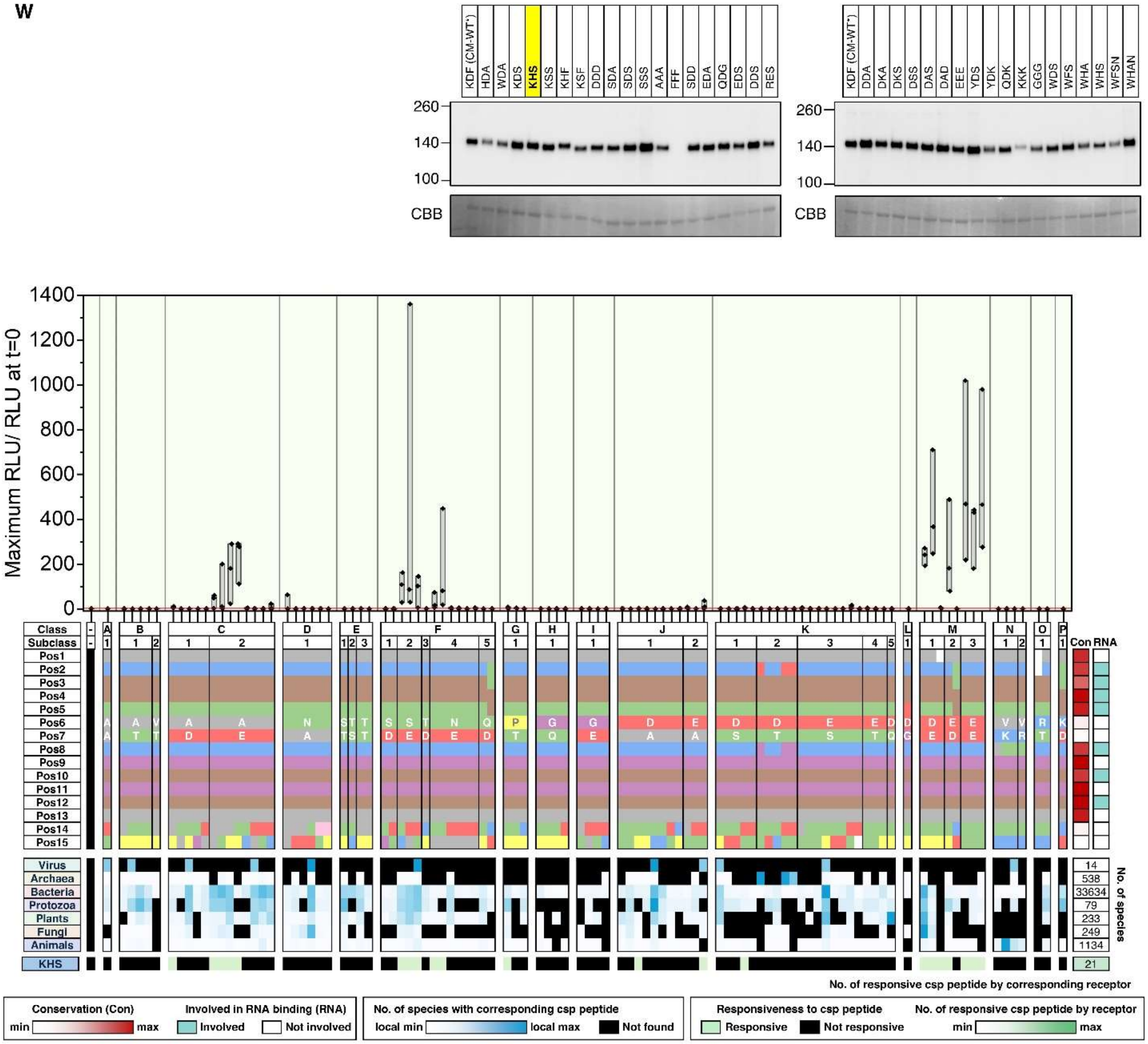
KDF (CM-WT*)
HDA
WDA
KDS
KHS
KSS
KHF
KSF
DDD
SDA
SDS
SSS
AAA
FFF
SDD
EDA
QDG
EDS
DDS
RES
DDA
DKA
DKS
DSS
DAS
DAD
EEE
YDS
YDK
QDK
KKK
GGG
WDS
WFS
WHA
WHS
WFSN
WHAN
260
140
100
CBB
Maximum RLU/ RLU at t=0
1400
1200
1000
800
600
400
200
0
Class
Subclass
Pos1
Pos2
Pos3
Pos4
Pos5
Pos6
Pos7
Pos8
Pos9
Pos10
Pos11
Pos12
Pos13
Pos14
Pos15
Con
RNA
Virus
Archaea
Bacteria
Protozoa
Plants
Fungi
Animals
KHS
14
538
33634
79
233
249
1134
21
No. of species
No. of responsive csp peptide by corresponding receptor
Conservation (Con)
min
max
Involved in RNA binding (RNA)
Involved
Not involved
No. of species with corresponding csp peptide
local min
local max
Not found
Responsiveness to csp peptide
Responsive
Not responsive
No. of responsive csp peptide by receptor
min
max

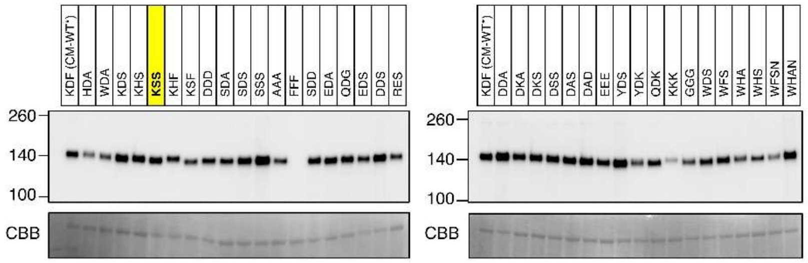
KDF (CM-WT')
HDA
WDA
KDS
KHS
KSS
KHF
KSF
DDD
SDA
SDS
SSS
AAA
FFF
SDD
EDA
QDG
EDS
DDS
RES
260
140
100
CBB
KDF (CM-WT')
DDA
DKA
DKS
DSS
DAS
DAD
EEE
YDS
YDK
QDK
KKK
GGG
WDS
WFS
WHA
WHS
WFSN
WHAN
260
140
100
CBB

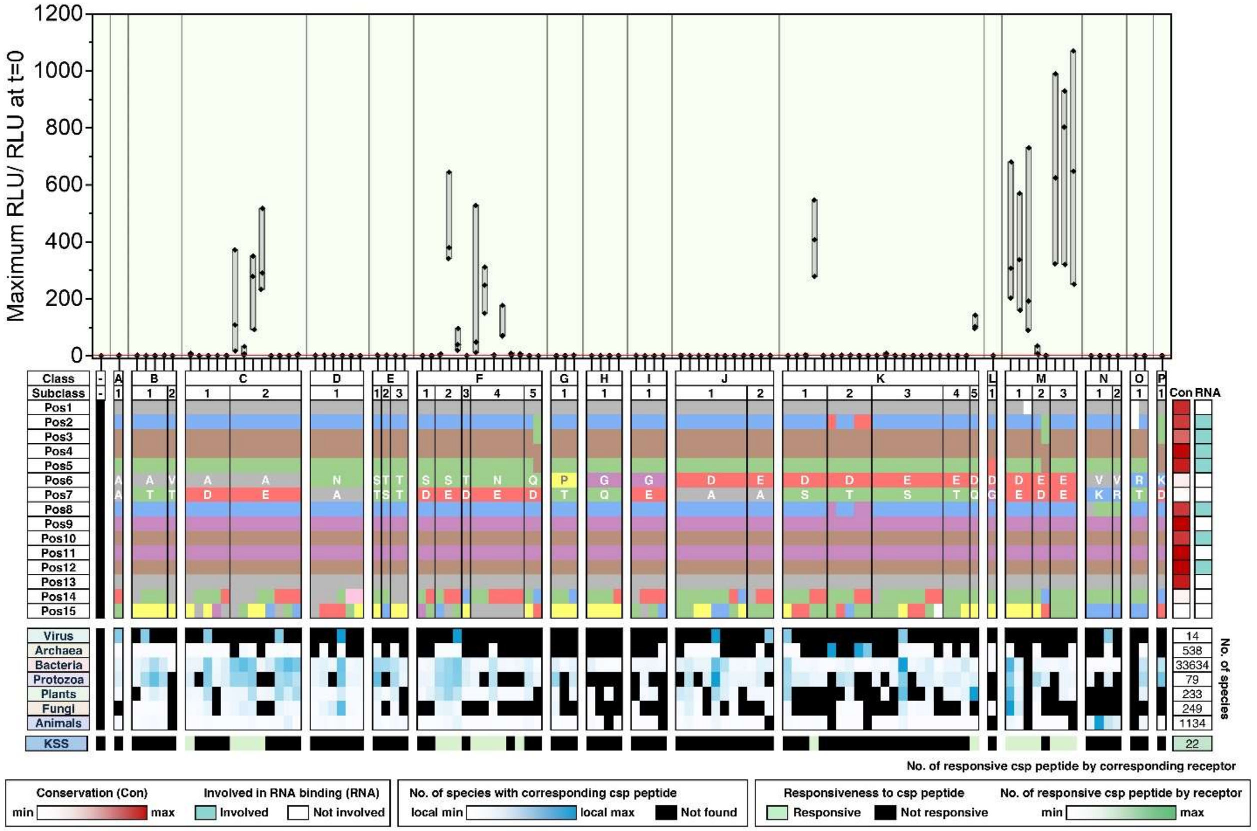
Maximum RLU/ RLU at t=0
1200
1000
800
600
400
200
0
Class
Subclass
Pos1
Pos2
Pos3
Pos4
Pos5
Pos6
Pos7
Pos8
Pos9
Pos10
Pos11
Pos12
Pos13
Pos14
Pos15
Con
RNA
Virus
Archaea
Bacteria
Protozoa
Plants
Fungi
Animals
KSS
No. of species
14
538
33634
79
233
249
1134
22
No. of responsive csp peptide by corresponding receptor
Conservation (Con)
min
max
Involved in RNA binding (RNA)
Involved
Not involved
No. of species with corresponding csp peptide
local min
local max
Not found
Responsiveness to csp peptide
Responsive
Not responsive
No. of responsive csp peptide by receptor
min
max

X

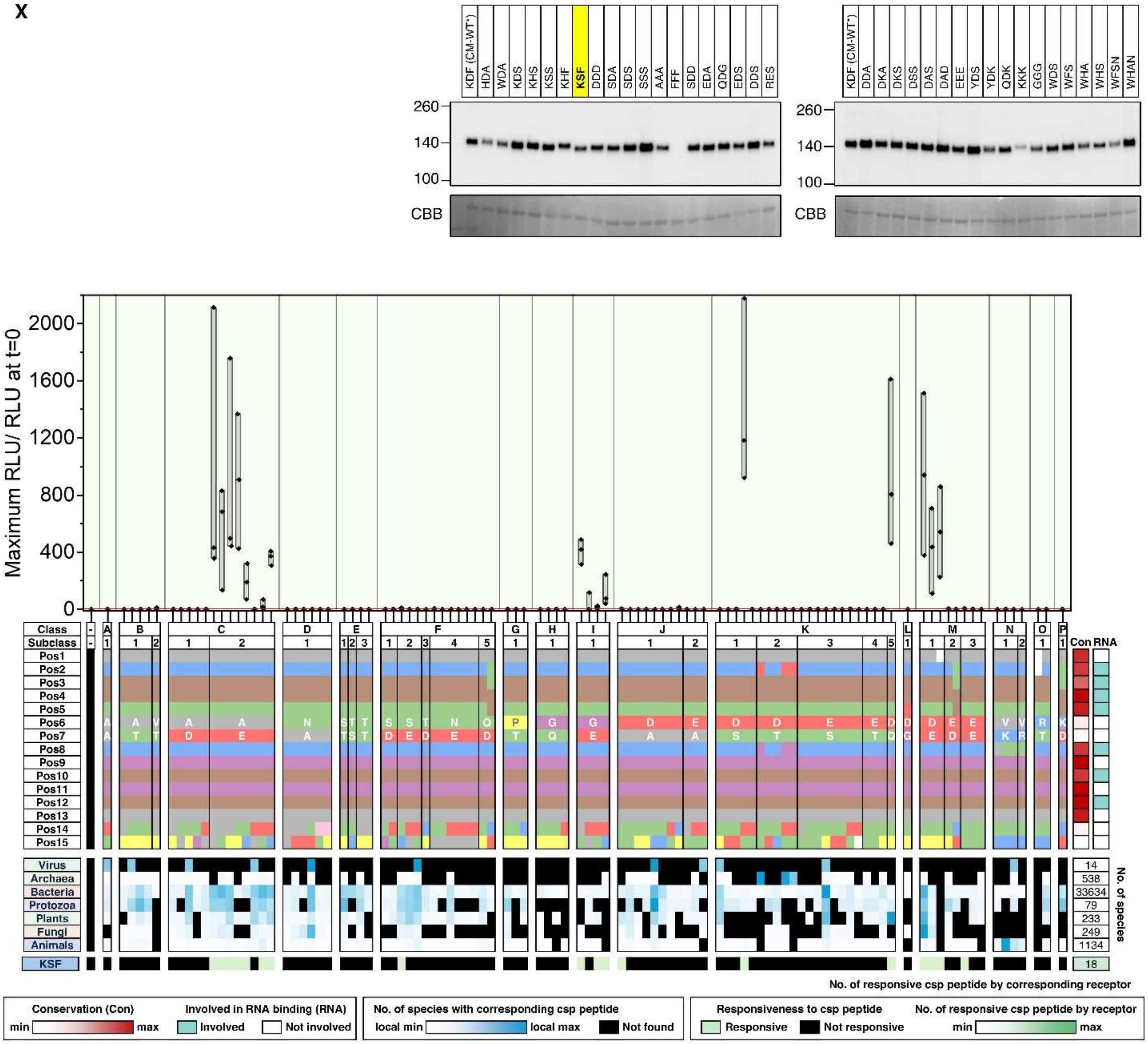

KDF (CM-WT*)
HDA
WDA
KDS
KHS
KSS
KHF
KSF
DDD
SDA
SDS
SSS
AAA
FFF
SDD
EDA
QDG
EDS
DDS
RES
DDA
DKA
DKS
DSS
DAS
DAD
EEE
YDS
YDK
QDK
KKK
GGG
WDS
WFS
WHA
WHS
WFSN
WHAN
260
140
100
CBB
Maximum RLU/ RLU at t=0
2000
1600
1200
800
400
0
Class
Subclass
Pos1
Pos2
Pos3
Pos4
Pos5
Pos6
Pos7
Pos8
Pos9
Pos10
Pos11
Pos12
Pos13
Pos14
Pos15
Con
RNA
Virus
Archaea
Bacteria
Protozoa
Plants
Fungi
Animals
KSF
No. of species
14
538
33634
79
233
249
1134
18
No. of responsive csp peptide by corresponding receptor
Conservation (Con)
min
max
Involved in RNA binding (RNA)
Involved
Not involved
No. of species with corresponding csp peptide
local min
local max
Not found
Responsiveness to csp peptide
Responsive
Not responsive
No. of responsive csp peptide by receptor
min
max

Y

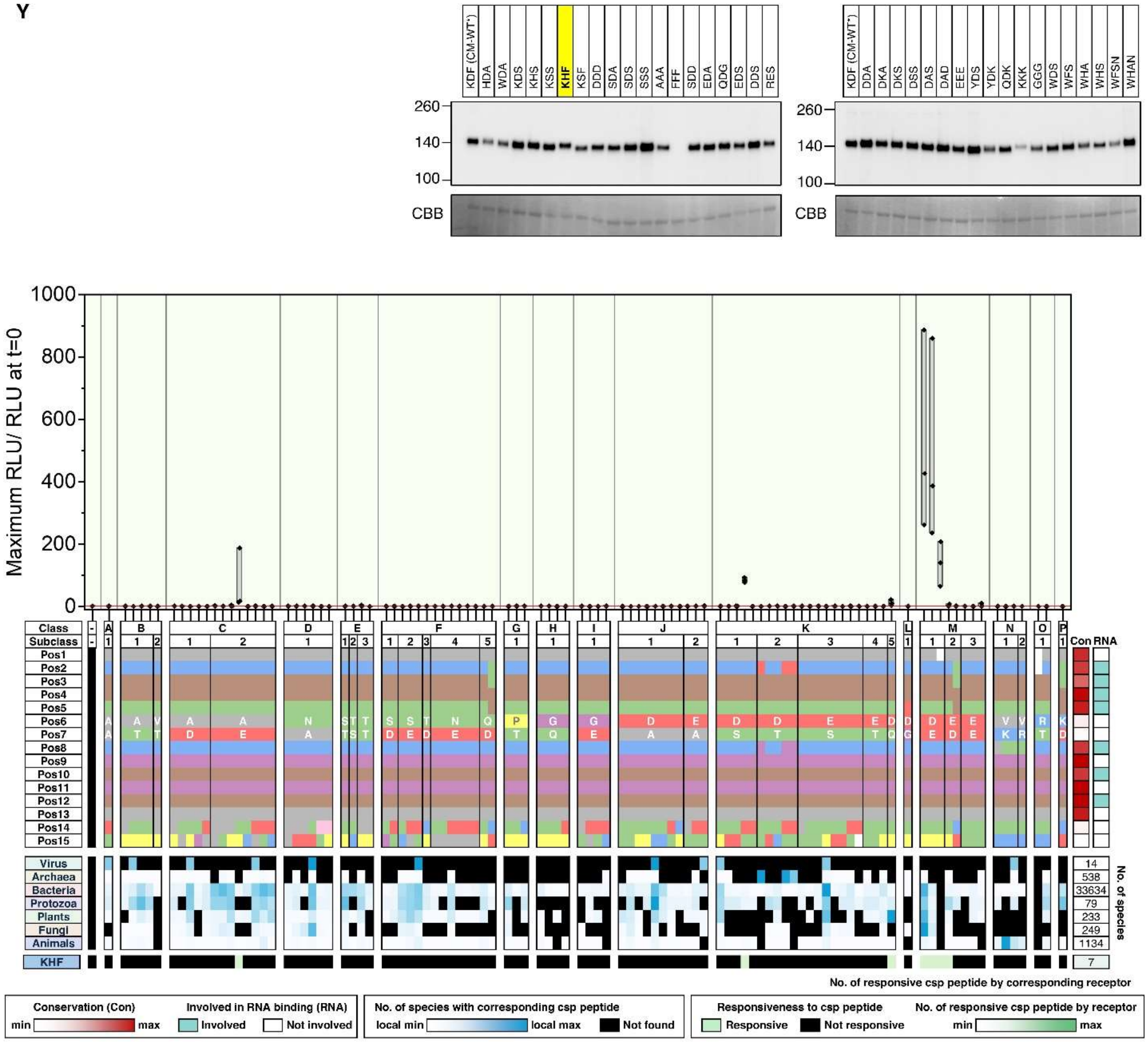
KDF (CM-WT*)
HDA
WDA
KDS
KHS
KSS
KHF
KSF
DDD
SDA
SDS
SSS
AAA
FFF
SDD
EDA
QDG
EDS
DDS
RES
DDA
DKA
DKS
DSS
DAS
DAD
EEE
YDS
YDK
QDK
KKK
GGG
WDS
WFS
WHA
WHS
WFSN
WHAN
260
140
100
CBB
Maximum RLU/ RLU at t=0
1000
800
600
400
200
0
Class
Subclass
Pos1
Pos2
Pos3
Pos4
Pos5
Pos6
Pos7
Pos8
Pos9
Pos10
Pos11
Pos12
Pos13
Pos14
Pos15
Con
RNA
Virus
Archaea
Bacteria
Protozoa
Plants
Fungi
Animals
KHF
No. of species
14
538
33634
79
233
249
1134
7
No. of responsive csp peptide by corresponding receptor
Conservation (Con)
min
max
Involved in RNA binding (RNA)
Involved
Not involved
No. of species with corresponding csp peptide
local min
local max
Not found
Responsiveness to csp peptide
Responsive
Not responsive
No. of responsive csp peptide by receptor
min
max

**Z**

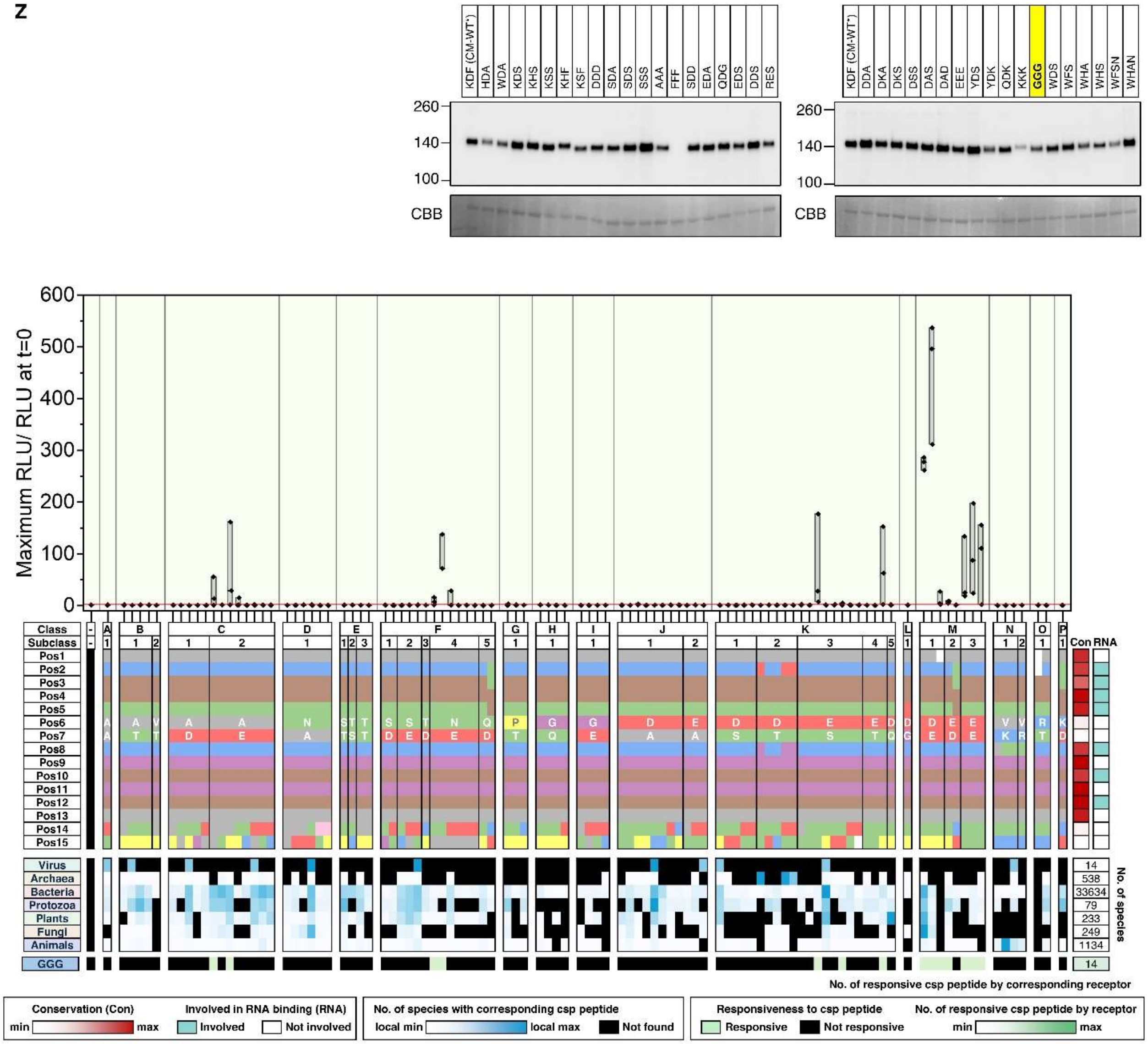

KDF (CM-WT*)
HDA
WDA
KDS
KHS
KSS
KHF
KSF
DDD
SDA
SDS
SSS
AAA
FFF
SDD
EDA
QDG
EDS
DDS
RES
DDA
DKA
DKS
DSS
DAS
DAD
EEE
YDS
YDK
QDK
KKK
GGG
WDS
WFS
WHA
WHS
WFSN
WHAN
260
140
100
CBB
Maximum RLU/ RLU at t=0
Class
Subclass
Con
RNA
Virus
Archaea
Bacteria
Protozoa
Plants
Fungi
Animals
No. of species
14
538
33634
79
233
249
1134
GGG
No. of responsive csp peptide by corresponding receptor
Conservation (Con)
min
max
Involved in RNA binding (RNA)
Involved
Not involved
No. of species with corresponding csp peptide
local min
local max
Not found
Responsiveness to csp peptide
Responsive
Not responsive
No. of responsive csp peptide by receptor

AA

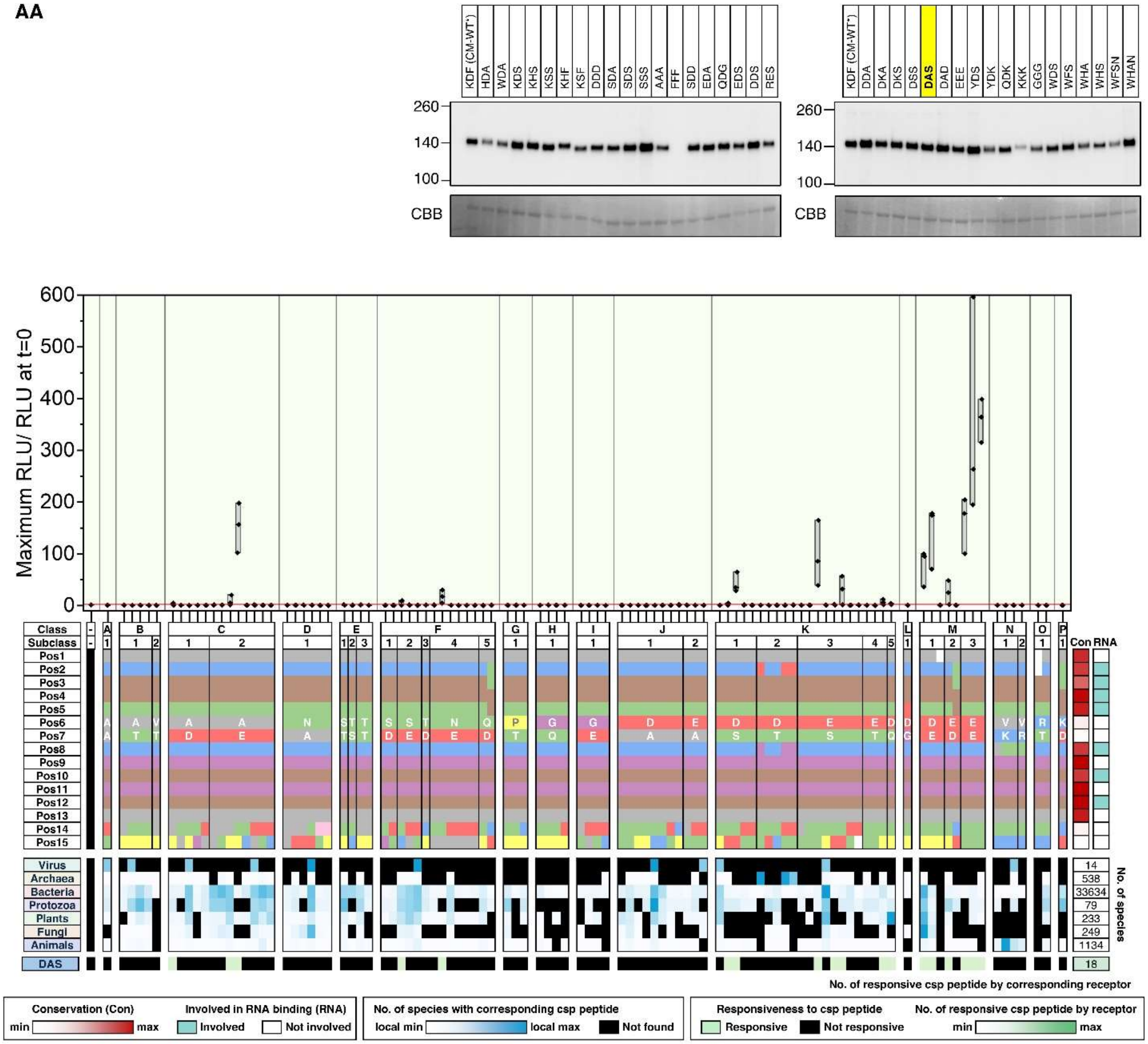
KDF (CM-WT*)
HDA
WDA
KDS
KHS
KSS
KHF
KSF
DDD
SDA
SDS
SSS
AAA
FFF
SDD
EDA
QDG
EDS
DDS
RES
260
140
100
CBB
KDF (CM-WT*)
DDA
DKA
DKS
DSS
DAS
DAD
EEE
YDS
YDK
QDK
KKK
GGG
WDS
WFS
WHA
WHS
WFSN
WHAN
Maximum RLU/ RLU at t=0
600
500
400
300
200
100
0
Class
Subclass
Pos1
Pos2
Pos3
Pos4
Pos5
Pos6
Pos7
Pos8
Pos9
Pos10
Pos11
Pos12
Pos13
Pos14
Pos15
Con
RNA
Virus
Archaea
Bacteria
Protozoa
Plants
Fungi
Animals
DAS
14
538
33634
79
233
249
1134
18
No. of species
No. of responsive csp peptide by corresponding receptor
Conservation (Con)
min
max
Involved in RNA binding (RNA)
Involved
Not involved
No. of species with corresponding csp peptide
local min
local max
Not found
Responsiveness to csp peptide
Responsive
Not responsive
No. of responsive csp peptide by receptor
min
max

**AB**

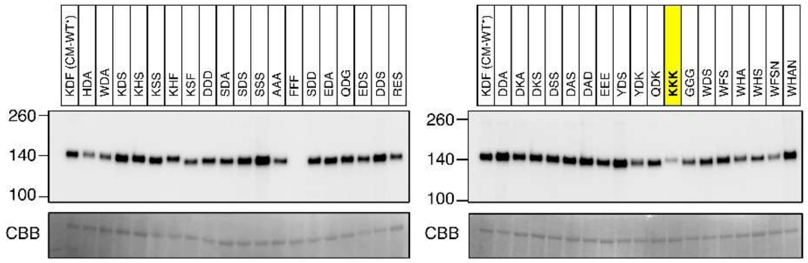
KDF (CM-WT*)
HDA
WDA
KDS
KHS
KSS
KHF
KSF
DDD
SDA
SDS
SSS
AAA
FFF
SDD
EDA
QDG
EDS
DDS
RES
DDA
DKA
DKS
DSS
DAS
DAD
EEE
YDS
YDK
QDK
KKK
GGG
WDS
WFS
WHA
WHS
WFSN
WHAN
260
140
100
CBB

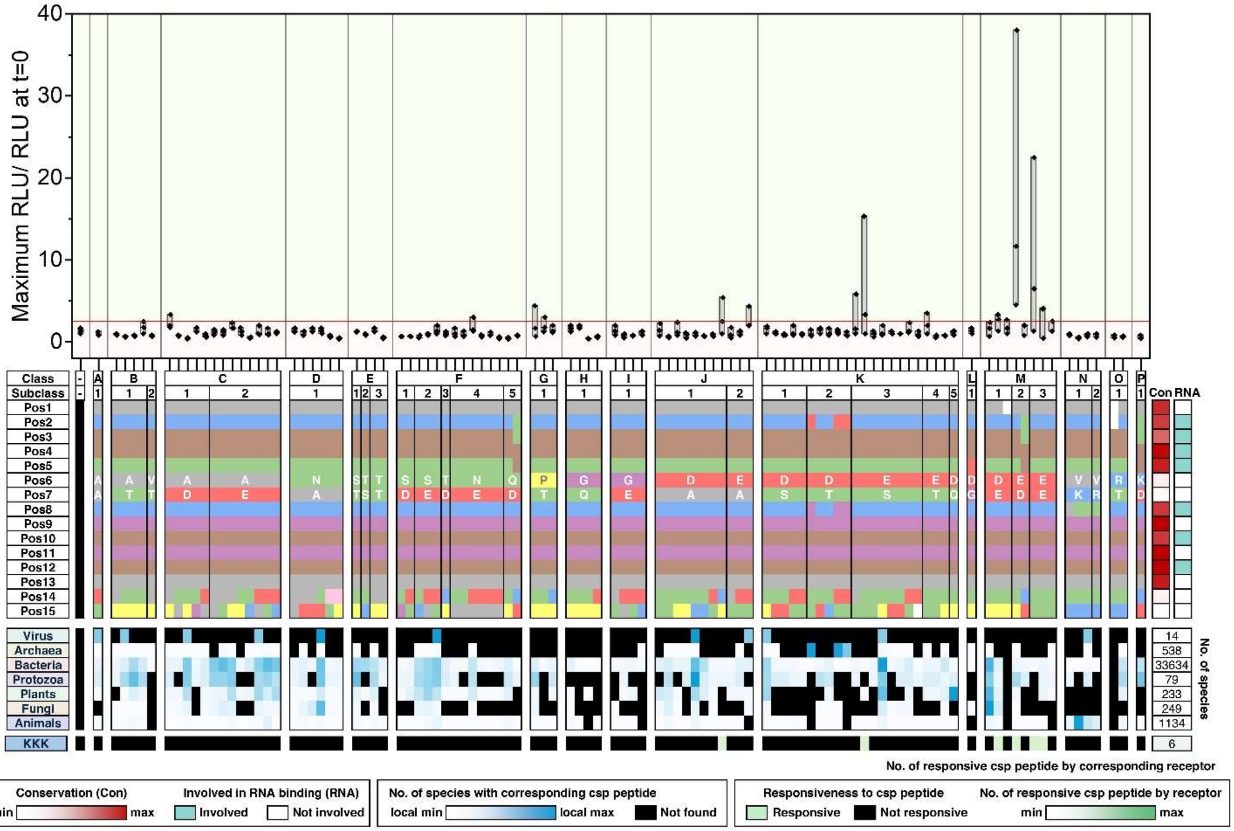
Maximum RLU/ RLU at t=0
40
30
20
10
0
Class
Subclass
Pos1
Pos2
Pos3
Pos4
Pos5
Pos6
Pos7
Pos8
Pos9
Pos10
Pos11
Pos12
Pos13
Pos14
Pos15
Con
RNA
Virus
Archaea
Bacteria
Protozoa
Plants
Fungi
Animals
KKK
14
538
33634
79
233
249
1134
6
No. of species
No. of responsive csp peptide by corresponding receptor
Conservation (Con)
min
max
Involved in RNA binding (RNA)
Involved
Not involved
No. of species with corresponding csp peptide
local min
local max
Not found
Responsiveness to csp peptide
Responsive
Not responsive
No. of responsive csp peptide by receptor
min
max

**AC**

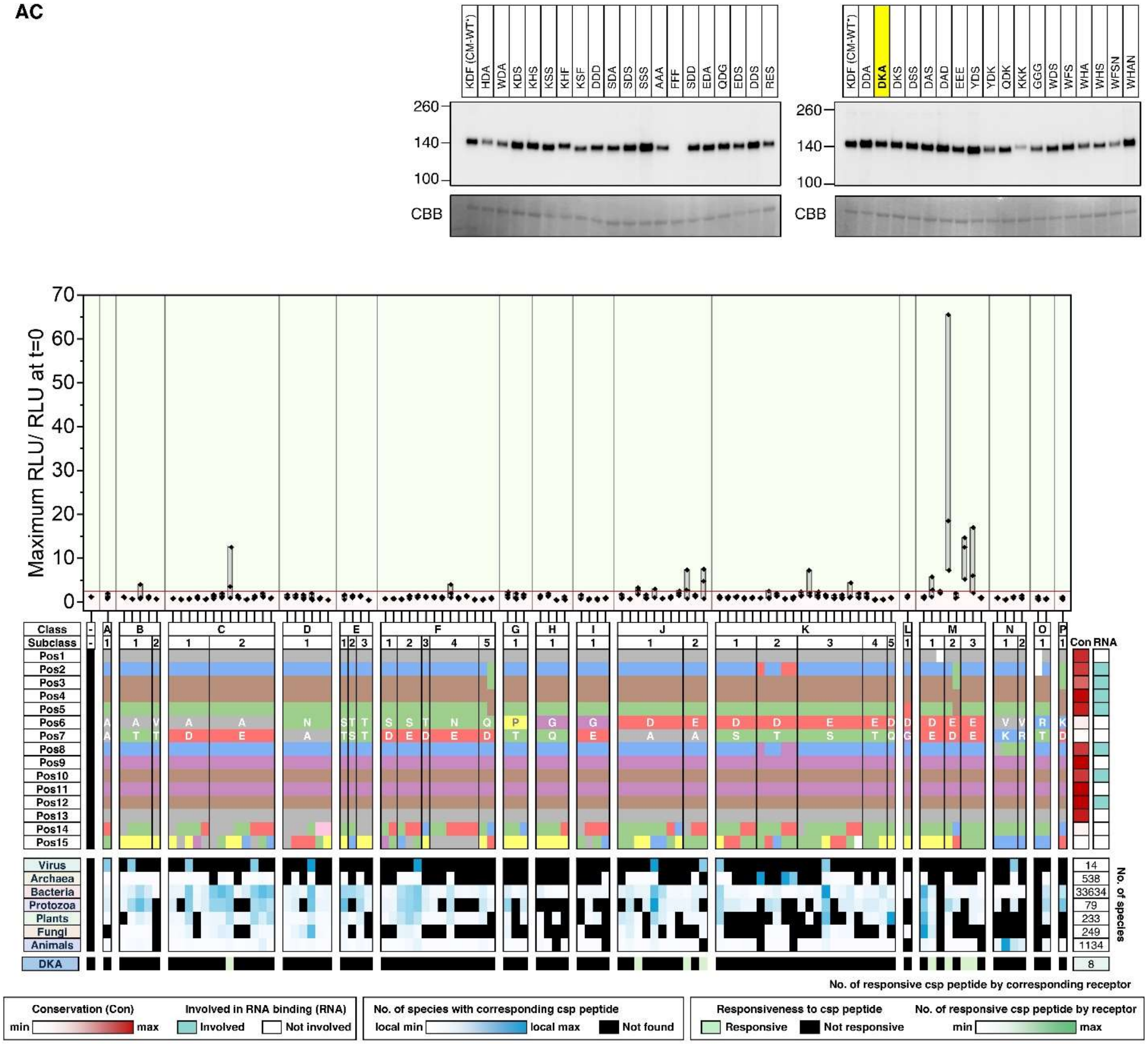
KDF (CM-WT*)
HDA
WDA
KDS
KHS
KSS
KHF
KSF
DDD
SDA
SDS
SSS
AAA
FFF
SDD
EDA
QDG
EDS
DDS
RES
KDF (CM-WT*)
DDA
DKA
DKS
DSS
DAS
DAD
EEE
YDS
YDK
QDK
KKK
GGG
WDS
WFS
WHA
WHS
WFSN
WHAN
260
140
100
CBB
Maximum RLU/ RLU at t=0
Class
Subclass
Pos1
Pos2
Pos3
Pos4
Pos5
Pos6
Pos7
Pos8
Pos9
Pos10
Pos11
Pos12
Pos13
Pos14
Pos15
Con
RNA
Virus
Archaea
Bacteria
Protozoa
Plants
Fungi
Animals
DKA
No. of species
14
538
33634
79
233
249
1134
8
No. of responsive csp peptide by corresponding receptor
Conservation (Con)
min
max
Involved in RNA binding (RNA)
Involved
Not involved
No. of species with corresponding csp peptide
local min
local max
Not found
Responsiveness to csp peptide
Responsive
Not responsive
No. of responsive csp peptide by receptor
min
max

AD

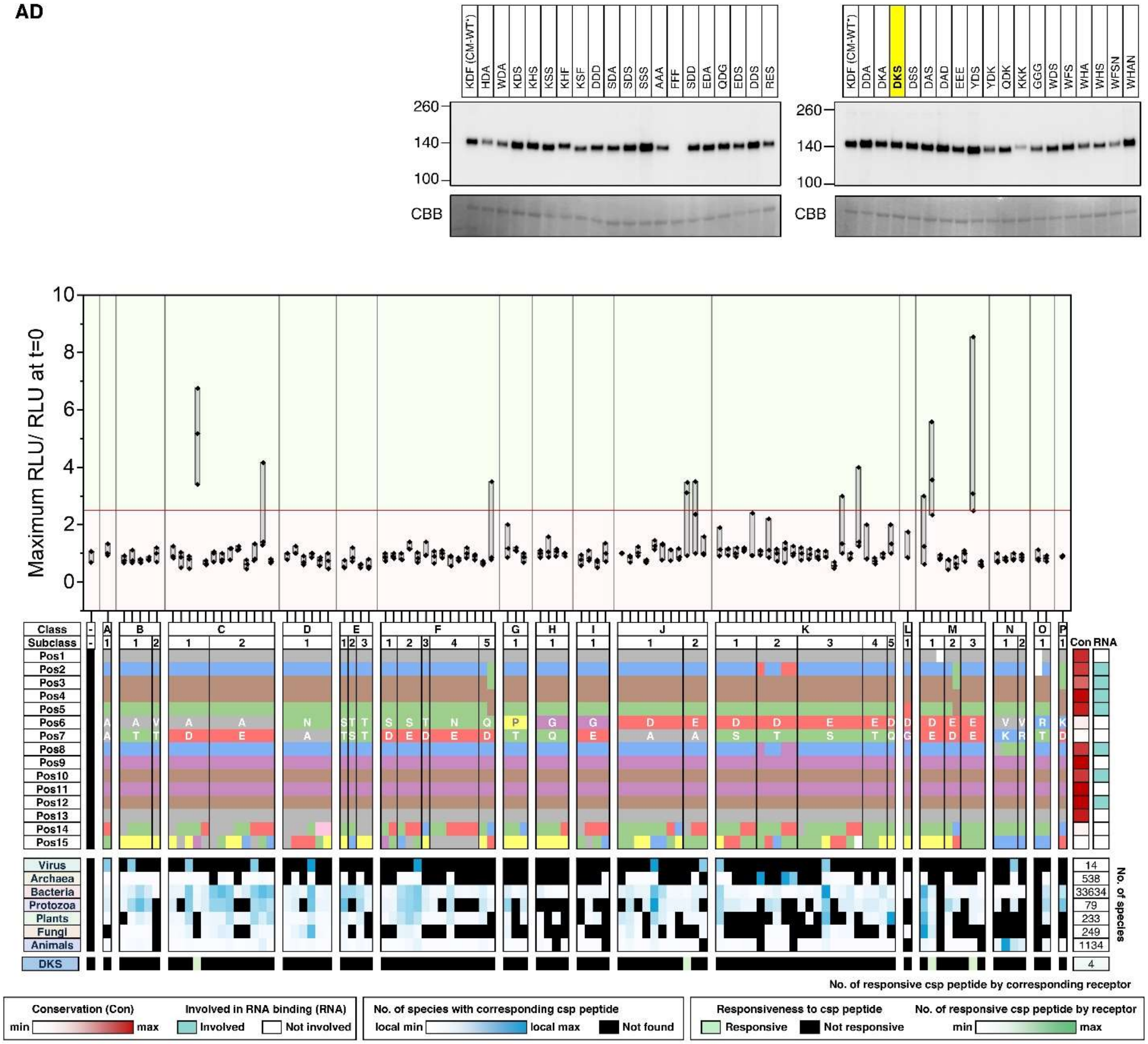
KDF (CM-WT*)
HDA
WDA
KDS
KHS
KSS
KHF
KSF
DDD
SDA
SDS
SSS
AAA
FFF
SDD
EDA
QDG
EDS
DDS
RES
DDA
DKA
DKS
DSS
DAS
DAD
EEE
YDS
YDK
QDK
KKK
GGG
WDS
WFS
WHA
WHS
WFSN
WHAN
260
140
100
CBB
Maximum RLU/ RLU at t=0
Class
Subclass
Virus
Archaea
Bacteria
Protozoa
Plants
Fungi
Animals
DKS
Con
RNA
No. of species
14
538
33634
79
233
249
1134
4
No. of responsive csp peptide by corresponding receptor
Conservation (Con)
min
max
Involved in RNA binding (RNA)
Involved
Not involved
No. of species with corresponding csp peptide
local min
local max
Not found
Responsiveness to csp peptide
Responsive
Not responsive
No. of responsive csp peptide by receptor

**AE**

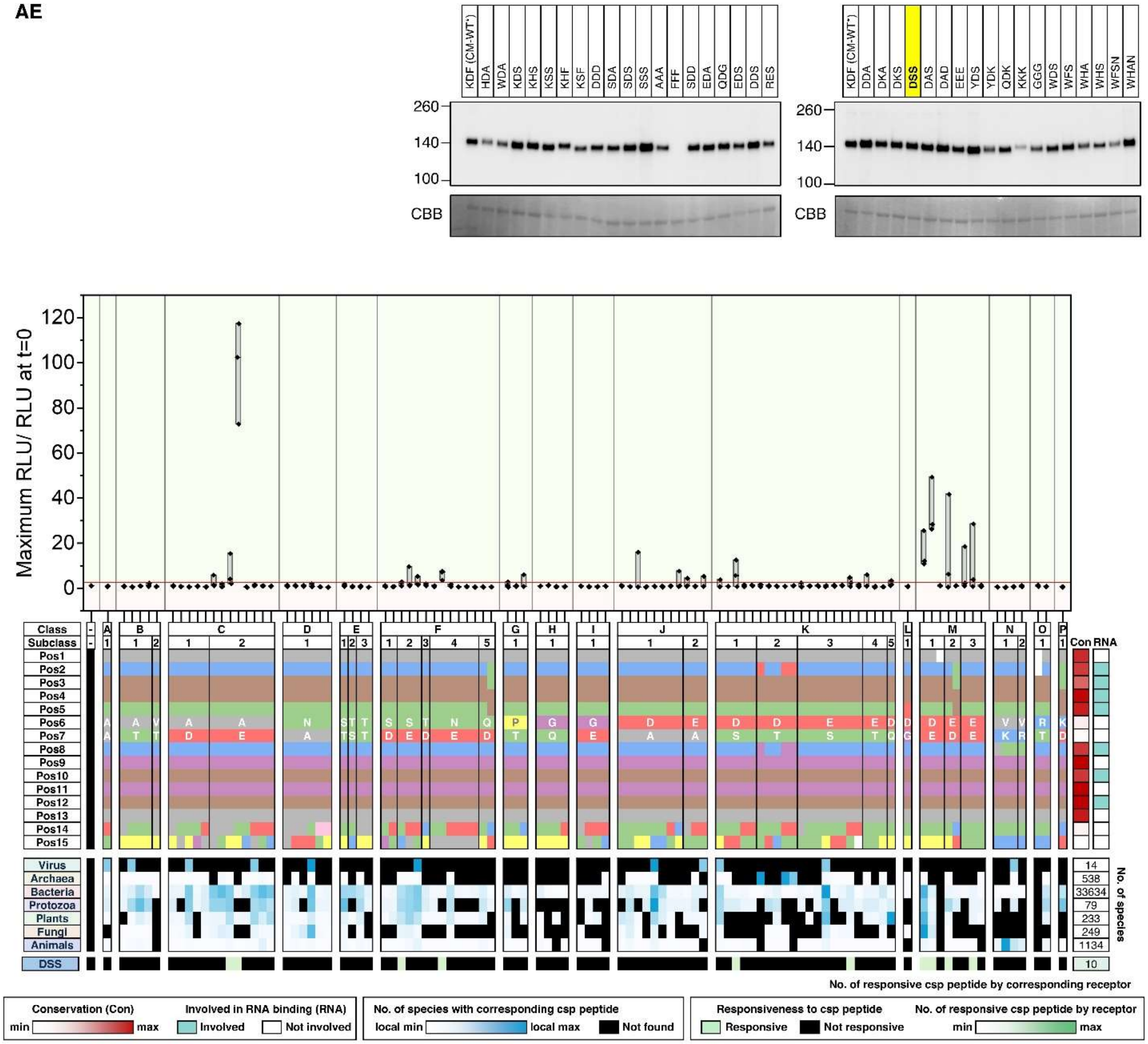

AF

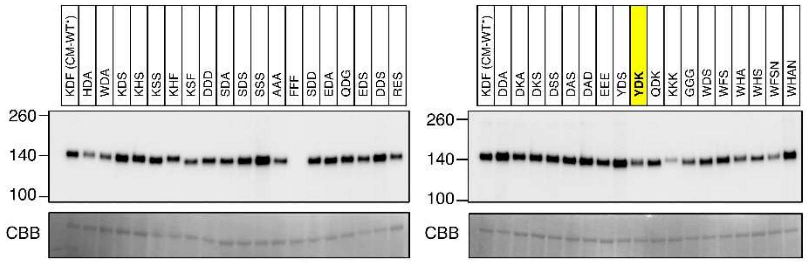
KDF (CM-WT')
HDA
WDA
KDS
KHS
KSS
KHF
KSF
DDD
SDA
SDS
SSS
AAA
FFF
SDD
EDA
QDG
EDS
DDS
RES
260
140
100
CBB
KDF (CM-WT')
DDA
DKA
DKS
DSS
DAS
DAD
EEE
YDS
YDK
QDK
KKK
GGG
WDS
WFS
WHA
WHS
WFSN
WHAN
260
140
100
CBB

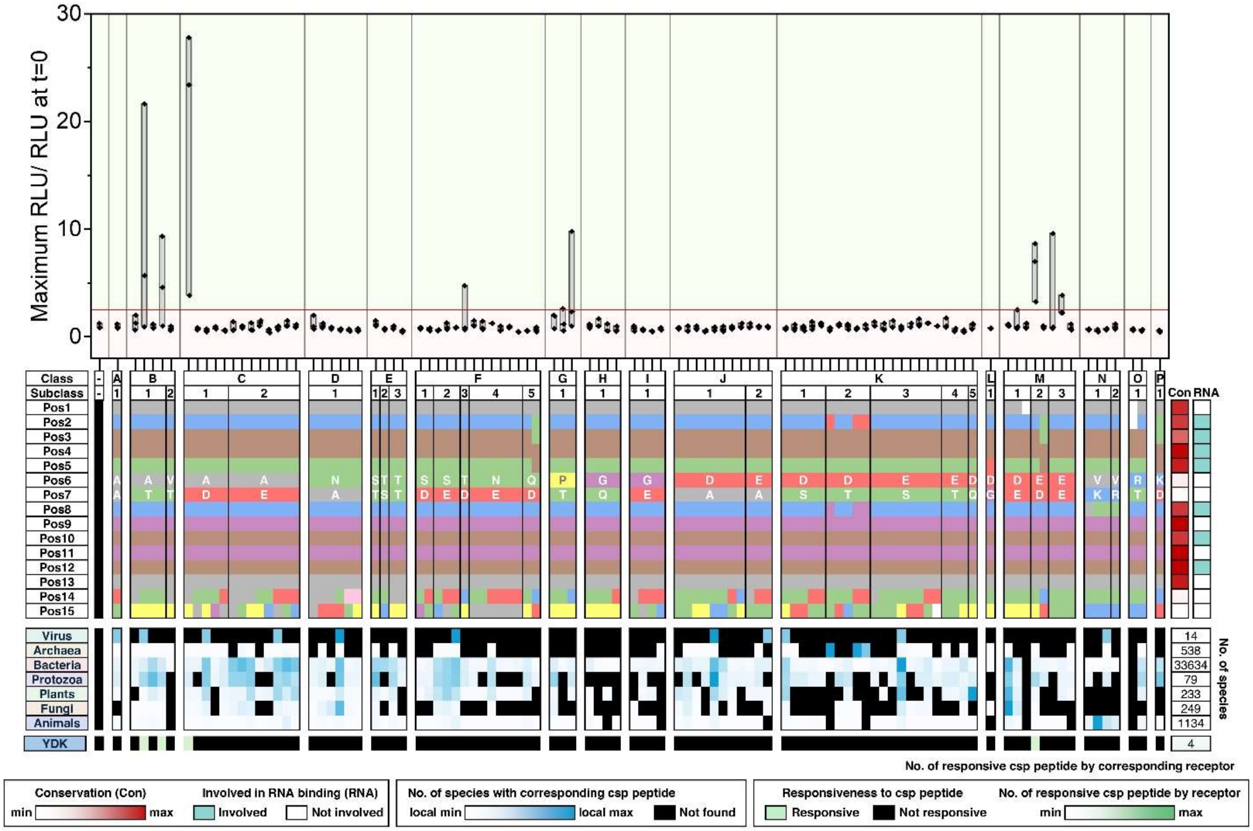
Maximum RLU/ RLU at t=0
30
20
10
0
Class
Subclass
Pos1
Pos2
Pos3
Pos4
Pos5
Pos6
Pos7
Pos8
Pos9
Pos10
Pos11
Pos12
Pos13
Pos14
Pos15
Con
RNA
Virus
Archaea
Bacteria
Protozoa
Plants
Fungi
Animals
YDK
14
538
33634
79
233
249
1134
4
No. of species
No. of responsive csp peptide by corresponding receptor
Conservation (Con)
min
max
Involved in RNA binding (RNA)
Involved
Not involved
No. of species with corresponding csp peptide
local min
local max
Not found
Responsiveness to csp peptide
Responsive
Not responsive
No. of responsive csp peptide by receptor
min
max

**AG**

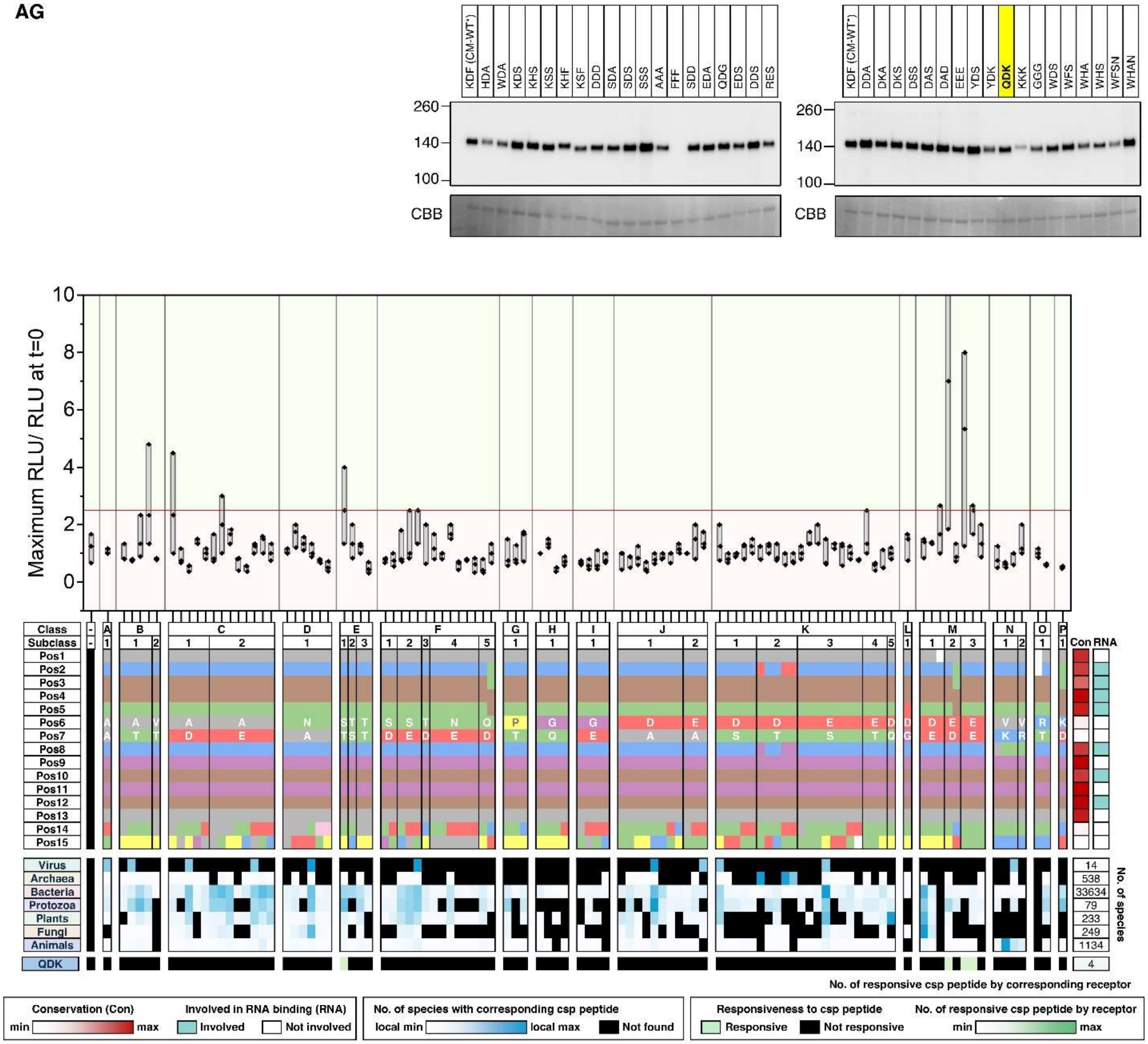
KDF (CM-WT*)
HDA
WDA
KDS
KHS
KSS
KHF
KSF
DDD
SDA
SDS
SSS
AAA
FFF
SDD
EDA
QDG
EDS
DDS
RES
DDA
DKA
DKS
DSS
DAS
DAD
EEE
YDS
YDK
QDK
KKK
GGG
WDS
WFS
WHA
WHS
WFSN
WHAN
260
140
100
CBB
Maximum RLU/ RLU at t=0
Class
Subclass
Pos1
Pos2
Pos3
Pos4
Pos5
Pos6
Pos7
Pos8
Pos9
Pos10
Pos11
Pos12
Pos13
Pos14
Pos15
Con
RNA
Virus
Archaea
Bacteria
Protozoa
Plants
Fungi
Animals
QDK
14
538
33634
79
233
249
1134
4
No. of species
No. of responsive csp peptide by corresponding receptor
Conservation (Con)
min
max
Involved in RNA binding (RNA)
Involved
Not involved
No. of species with corresponding csp peptide
local min
local max
Not found
Responsiveness to csp peptide
Responsive
Not responsive
No. of responsive csp peptide by receptor
min
max

**AH**

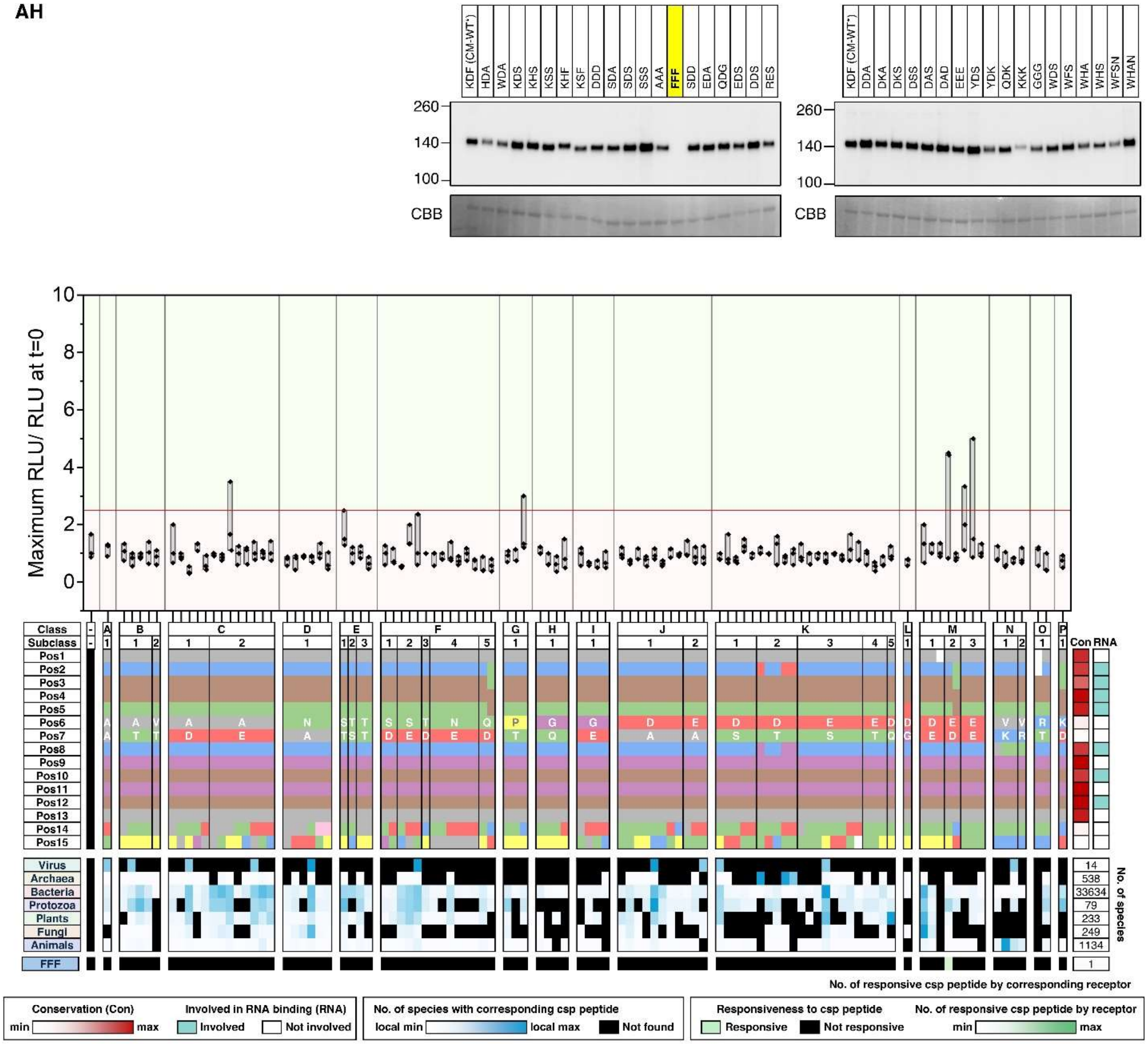
KDF (CM-WT*)
HDA
WDA
KDS
KHS
KSS
KHF
KSF
DDD
SDA
SDS
SSS
AAA
FFF
SDD
EDA
QDG
EDS
DDS
RES
DDA
DKA
DKS
DSS
DAS
DAD
EEE
YDS
YDK
QDK
KKK
GGG
WDS
WFS
WHA
WHS
WFSN
WHAN
260
140
100
CBB
Maximum RLU/ RLU at t=0
10
8
6
4
2
0
Class
Subclass
Pos1
Pos2
Pos3
Pos4
Pos5
Pos6
Pos7
Pos8
Pos9
Pos10
Pos11
Pos12
Pos13
Pos14
Pos15
Con
RNA
Virus
Archaea
Bacteria
Protozoa
Plants
Fungi
Animals
FFF
14
538
33634
79
233
249
1134
1
No. of species
No. of responsive csp peptide by corresponding receptor
Conservation (Con)
min
max
Involved in RNA binding (RNA)
Involved
Not involved
No. of species with corresponding csp peptide
local min
local max
Not found
Responsiveness to csp peptide
Responsive
Not responsive
No. of responsive csp peptide by receptor
min
max

AI

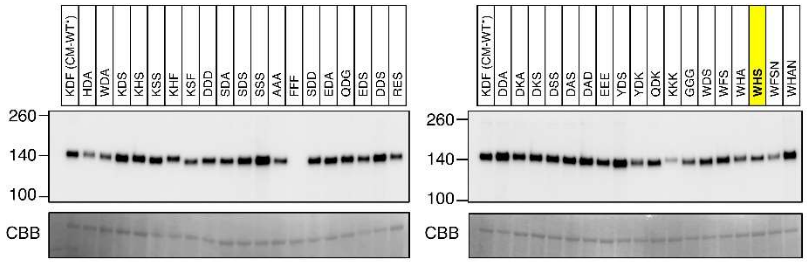

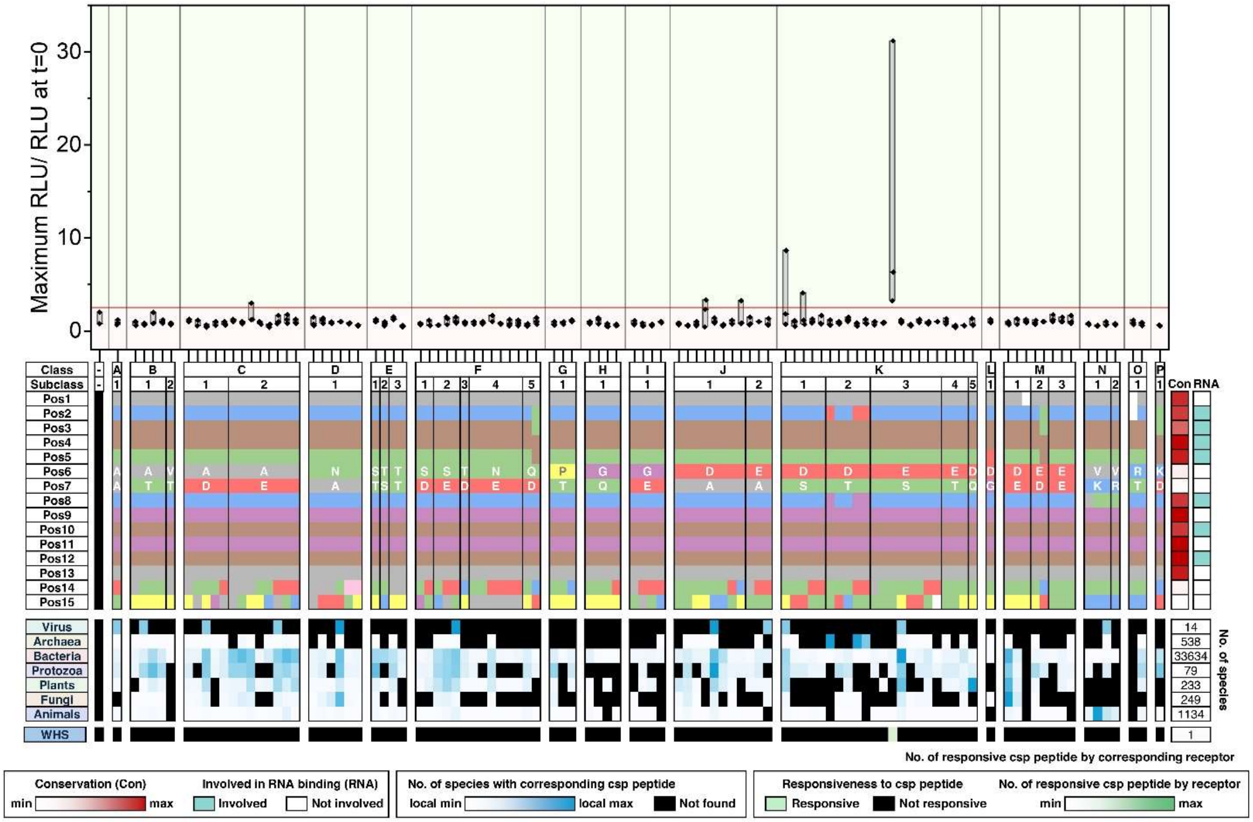

Maximum RLU/ RLU at t=0
Class
Subclass
Virus
Archaea
Bacteria
Protozoa
Plants
Fungi
Animals
WHS
No. of species
No. of responsive csp peptide by corresponding receptor
Conservation (Con)
Involved in RNA binding (RNA)
No. of species with corresponding csp peptide
Responsiveness to csp peptide
No. of responsive csp peptide by receptor

AJ

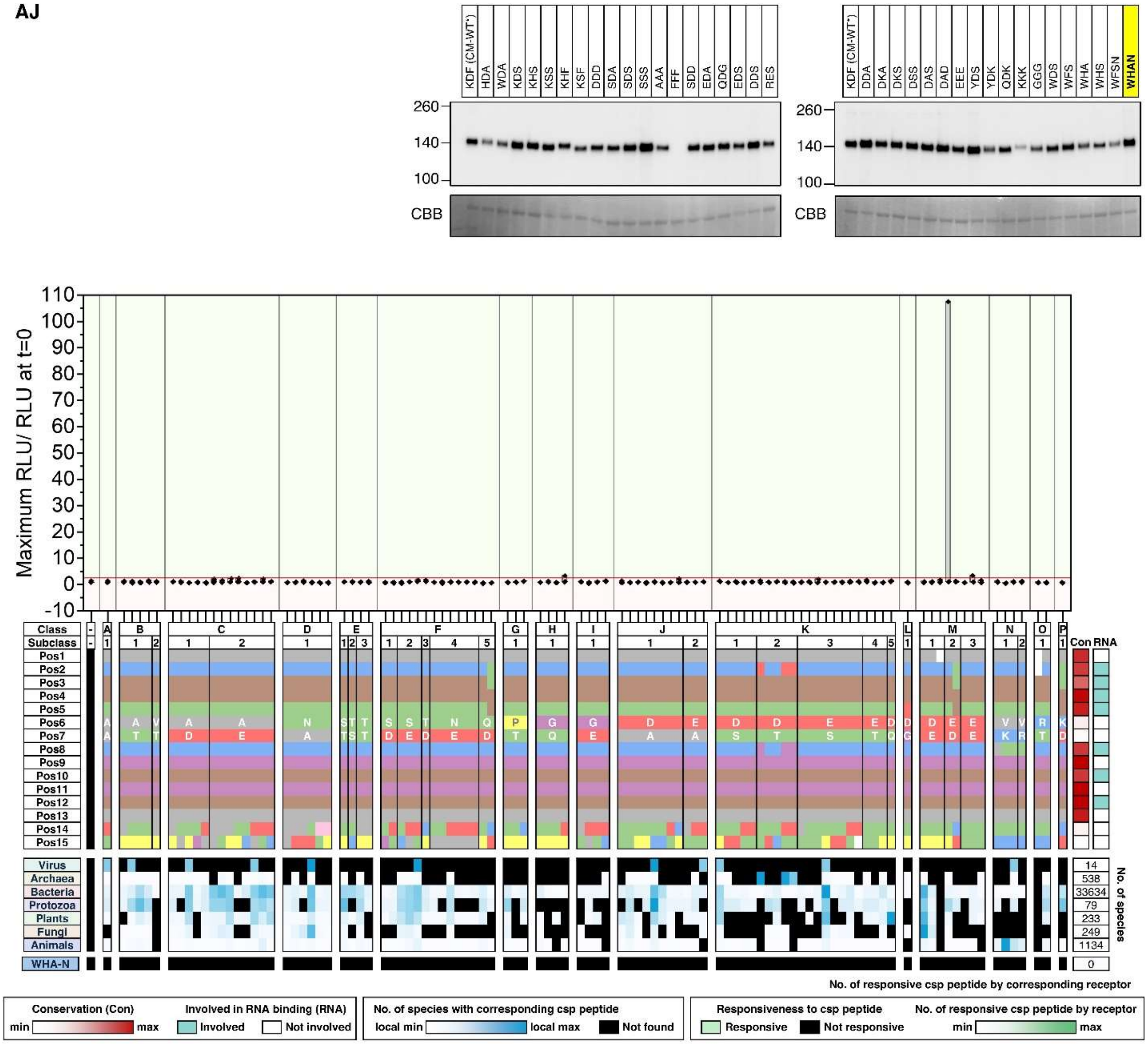
KDF (CM-WT')
HDA
WDA
KDS
KHS
KSS
KHF
KSF
DDD
SDA
SDS
SSS
AAA
FFF
SDD
EDA
QDG
EDS
DDS
RES
260
140
100
CBB
KDF (CM-WT')
DDA
DKA
DKS
DSS
DAS
DAD
EEE
YDS
YDK
QDK
KKK
GGG
WDS
WFS
WHA
WHS
WFSN
WHAN
260
140
100
CBB
Maximum RLU/ RLU at t=0
110
100
90
80
70
60
50
40
30
20
10
0
-10
Class
Subclass
Pos1
Pos2
Pos3
Pos4
Pos5
Pos6
Pos7
Pos8
Pos9
Pos10
Pos11
Pos12
Pos13
Pos14
Pos15
Con
RNA
Virus
Archaea
Bacteria
Protozoa
Plants
Fungi
Animals
14
538
33634
79
233
249
1134
No. of species
WHA-N
0
No. of responsive csp peptide by corresponding receptor
Conservation (Con)
min
max
Involved in RNA binding (RNA)
Involved
Not involved
No. of species with corresponding csp peptide
local min
local max
Not found
Responsiveness to csp peptide
Responsive
Not responsive
No. of responsive csp peptide by receptor
min
max

**AK**

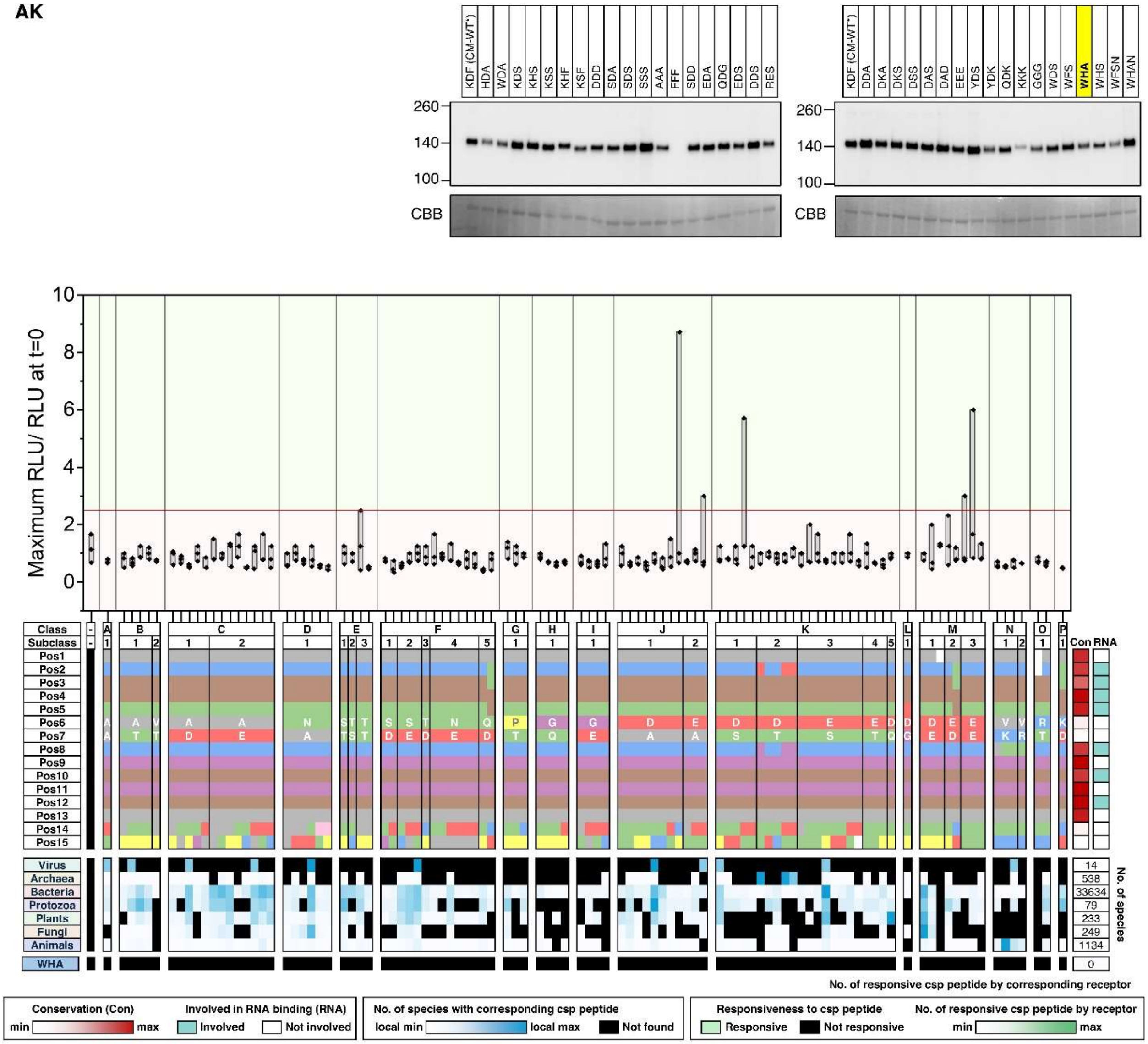

**Fig. S24. ROS data related to Fig 5D (amino acid substitutions).**

**(A)** The top chart represents the 103 csp15 peptides that are synthesized in this study. The amino acid properties in each position (Pos) are indicated with corresponding colors used in Fig. 4A. Peptides are grouped into classes and subclasses based on residues at the positions 6 and 7. Red heatmap on the right represents residue conservation scores, and cyan blocks mark RNA binding residues in CSPs. The middle blue heatmap represents the prevalence of the 103 csp15 peptide in each kingdom. Black blocks indicate peptides absent from all species in a kingdom. The number of species searched in each kingdom is shown on the right. Bottom chart represents the CSP recognition capacity of SCORE (CM) and variants generated from amino acid substitutions. A peptide was considered active if ROS readout exceeds the baseline by 2.5 times (green; see below). Non-responsive peptides are labelled in black. Green heatmap on the right indicates the number of responsive peptides (out of 103) for each receptor. **(B-AK)** Top immunoblot: Samples used in the following ROS assays were analysed with immunoblotting to confirm receptor expression. CBB staining is included as loading controls. Middle bar chart represents the ROS responses to mock treatment or each of the 103 csp15 peptide by the corresponding receptors. Leaves were then treated with 100 nM of indicated peptides, and ROS production was recorded for 15 minutes. For the box plots, the center line represents the median; the box bounds represent the 25th and 75th percentiles; and the whiskers extend to 1.5 × the interquartile range (IQR) from the 25th and 75th percentiles. Data points from 3 technical replicates were analyzed. A peptide was considered active if at least 2 out of 3 technical replicates showed ROS levels exceeding the baseline by 2.5 times (marked by the red line). Bottom charts: The top chart displays the 103 csp15 peptides synthesized in this study. Amino acid properties at each position (Pos) are color-coded, as in in Fig. 4A. Peptides are grouped into classes and subclasses based on residues at the positions 6 and 7. A red heatmap on the right indicates residue conservation scores, while cyan blocks mark RNA-binding residues in CSPs. The middle blue heatmap shows the prevalence of each csp15 peptide across kingdom. Black blocks represent peptides absent from all species within a kingdom. The number of species analyzed in each kingdom is shown on the right. Bottom chart represents the CSP recognition capacity of the corresponding receptors in plants. As noted above, peptides were considered active if at least 2 out of 3 technical replicates showed ROS levels exceeding the baseline by 2.5 times (see above). Non-responsive peptides are labeled in black. The green heatmap on the right indicates the number of responsive peptides (out of 103) for each receptor.

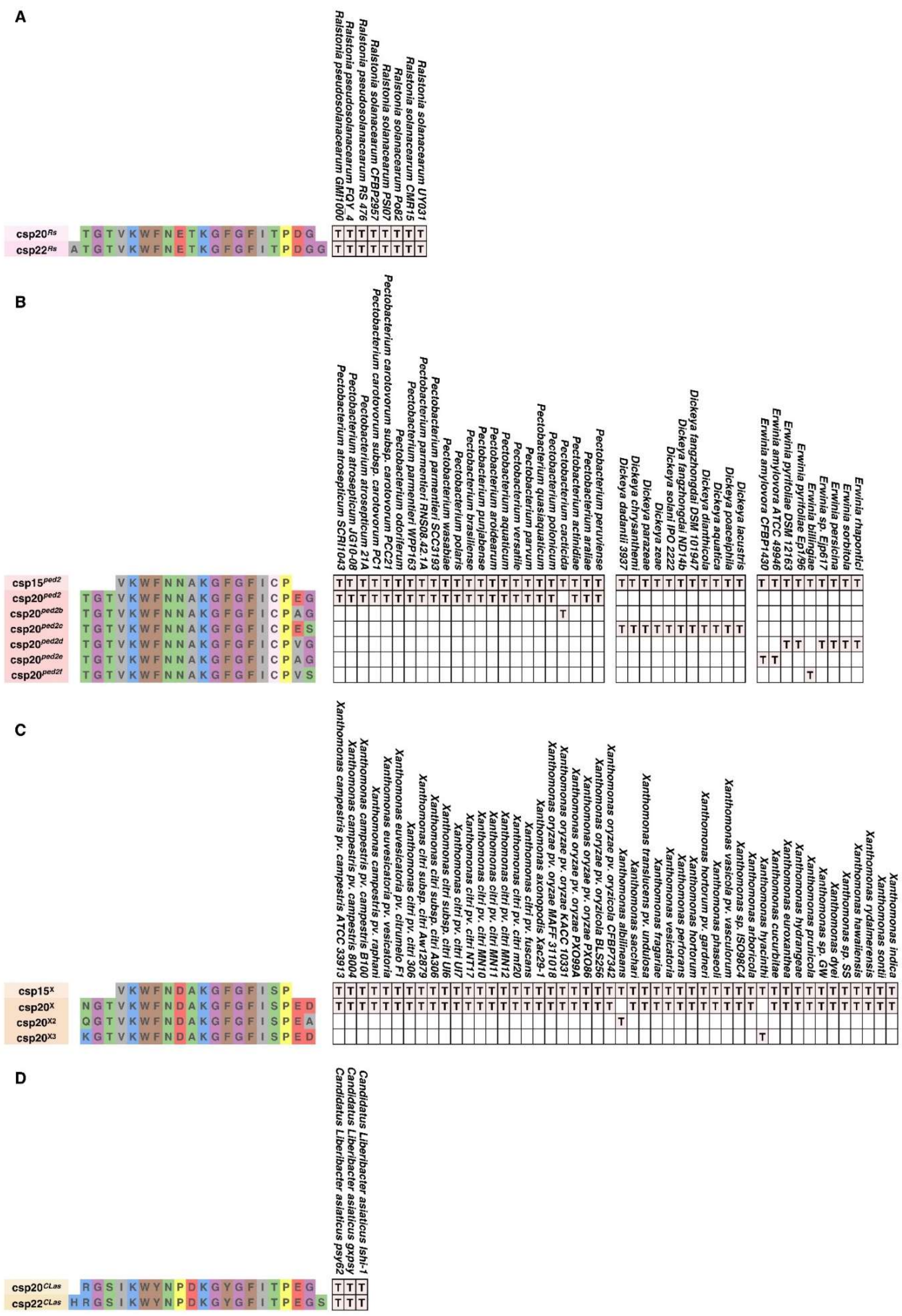
A
Ralstonia solanacearum UY031
Ralstonia solanacearum CMR15
Ralstonia solanacearum Po82
Ralstonia solanacearum PSI07
Ralstonia solanacearum CFBP2957
Ralstonia pseudosolanacearum RS 476
Ralstonia pseudosolanacearum FQY_4
Ralstonia pseudosolanacearum GMI1000
csp20Rs TGTVKWFNETKGFGFITPDG
csp22Rs ATGTVKWFNETKGFGFITPDGG
B
csp15ped2 VKWFNNAKGFGFICP
csp20ped2 TGTVKWFNNAKGFGFICPEG
csp20ped2b TGTVKWFNNAKGFGFICPAG
csp20ped2c TGTVKWFNNAKGFGFICPES
csp20ped2d TGTVKWFNNAKGFGFICPVG
csp20ped2e TGTVKWFNNAKGFGFICPAG
csp20ped2f TGTVKWFNNAKGFGFICPVS
C
csp15X VKWFNDAKGFGFISP
csp20X NGTVKWFNDAKGFGFISPED
csp20X2 QGTVKWFNDAKGFGFISPEA
csp20X3 KGTVKWFNDAKGFGFISPED
D
Candidatus Liberibacter asiaticus Ishi-1
Candidatus Liberibacter asiaticus gxpsy
Candidatus Liberibacter asiaticus psy62
csp20CLas RGSIKWYNPDKGYGFITPEG
csp22CLas HRGSIKWYNPDKGYGFITPEGS

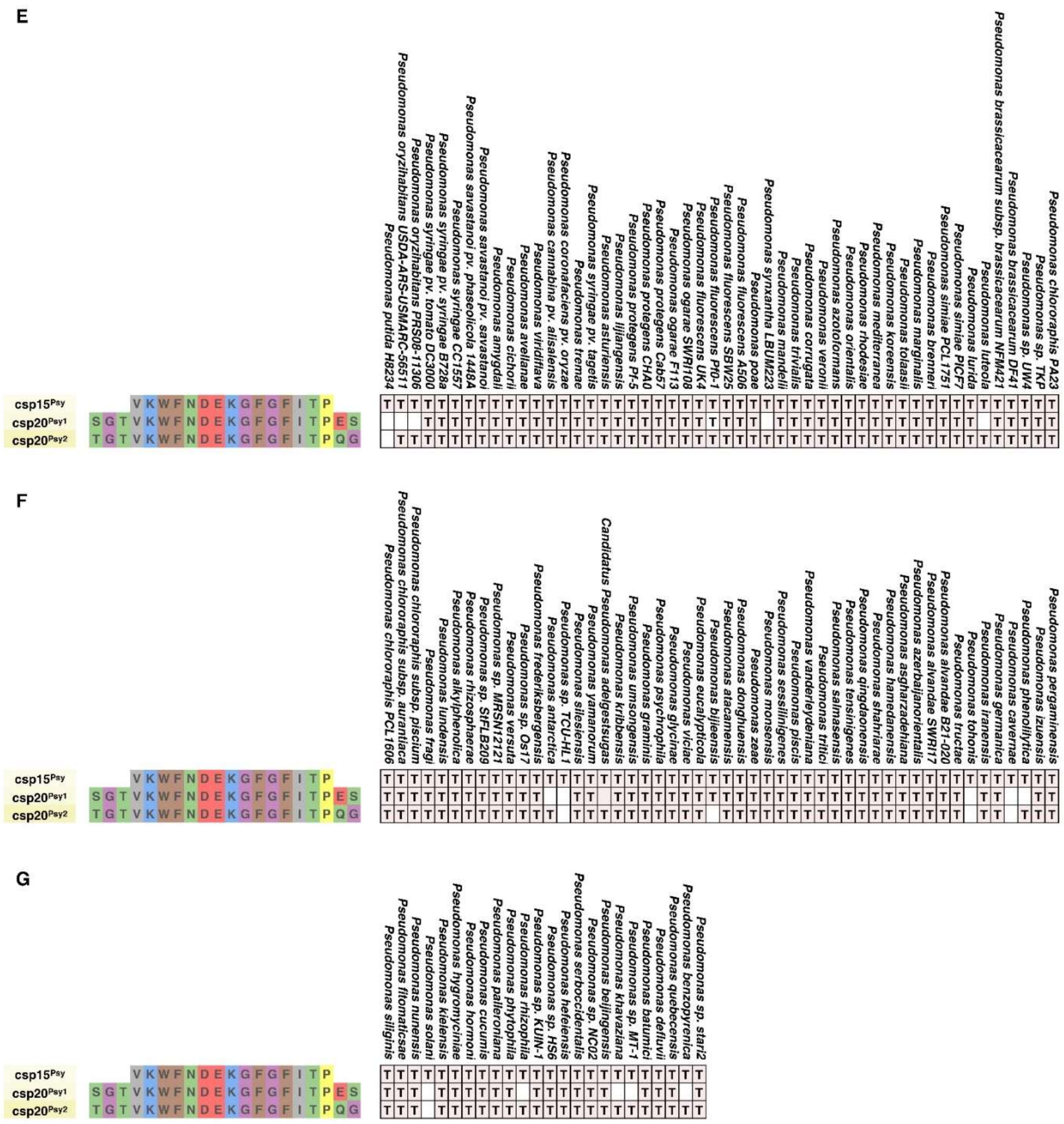
E
csp15Psy VKWFNDEKGFGFITP
csp20Psy1 SGTVKWFNDEKGFGFITPES
csp20Psy2 TGTVKWFNDEKGFGFITPQG
Pseudomonas chlororaphis PA23
Pseudomonas sp. TKP
Pseudomonas sp. UW4
Pseudomonas brassicacearum DF41
Pseudomonas brassicacearum subsp. brassicacearum NFM421
Pseudomonas luteola
Pseudomonas lurida
Pseudomonas simiae PICF7
Pseudomonas simiae PCL1751
Pseudomonas bremeri
Pseudomonas marginalis
Pseudomonas tolaasii
Pseudomonas koreensis
Pseudomonas mediterranea
Pseudomonas rhodesiae
Pseudomonas orientalis
Pseudomonas azotoformans
Pseudomonas veronii
Pseudomonas corrugata
Pseudomonas trivialis
Pseudomonas mandelii
Pseudomonas synxantha LBUM223
Pseudomonas poae
Pseudomonas fluorescens A506
Pseudomonas fluorescens SBW25
Pseudomonas fluorescens Pf0-1
Pseudomonas fluorescens UK4
Pseudomonas ogarae SWR108
Pseudomonas ogarae F113
Pseudomonas protegens Cab57
Pseudomonas protegens CHA0
Pseudomonas protegens Pf-5
Pseudomonas lijiangensis
Pseudomonas asturiensis
Pseudomonas tremae
Pseudomonas syringae pv. tagetis
Pseudomonas viridiflava
Pseudomonas cannabina pv. alisalensis
Pseudomonas coronafaciens pv. oryzae
Pseudomonas avellanae
Pseudomonas cichorii
Pseudomonas amygdali
Pseudomonas savastanoi pv. savastanoi
Pseudomonas savastanoi pv. phaseolicola 1448A
Pseudomonas syringae CC1557
Pseudomonas syringae pv. syringae B728a
Pseudomonas syringae pv. tomato DC3000
Pseudomonas oryzihabitans PRS08-11306
Pseudomonas oryzihabitans USDA-ARS-USMARC-56511
Pseudomonas putida H8234
F
csp15Psy VKWFNDEKGFGFITP
csp20Psy1 SGTVKWFNDEKGFGFITPES
csp20Psy2 TGTVKWFNDEKGFGFITPQG
Pseudomonas pergaminensis
Pseudomonas izuensis
Pseudomonas phenolilytica
Pseudomonas cavernae
Pseudomonas germanica
Pseudomonas iranensis
Pseudomonas tohonis
Pseudomonas fructae
Pseudomonas alvandae B21-020
Pseudomonas alvandae SWRI17
Pseudomonas azerbaijanorientalis
Pseudomonas asgharzadehiana
Pseudomonas hamedanensis
Pseudomonas shahriarae
Pseudomonas qingdaonensis
Pseudomonas tensinigenes
Pseudomonas salmasensis
Pseudomonas tritici
Pseudomonas vanderleydeniana
Pseudomonas piscis
Pseudomonas sessilinigenes
Pseudomonas monsensis
Pseudomonas zeae
Pseudomonas donghuensis
Pseudomonas atacamensis
Pseudomonas bijieensis
Pseudomonas eucalypticola
Pseudomonas viciae
Pseudomonas glycinae
Pseudomonas psychrophila
Pseudomonas graminis
Pseudomonas umsongensis
Pseudomonas kribbensis
Candidatus Pseudomonas adelgestsugas
Pseudomonas yamanorum
Pseudomonas silesiensis
Pseudomonas sp. TCU-HL1
Pseudomonas antarctica
Pseudomonas frederiksbergensis
Pseudomonas sp. OS17
Pseudomonas versuta
Pseudomonas sp. MRSN12121
Pseudomonas sp. SFLB209
Pseudomonas rhizosphaerae
Pseudomonas alkylphenolica
Pseudomonas lundensis
Pseudomonas fragi
Pseudomonas chlororaphis subsp. piscium
Pseudomonas chlororaphis subsp. aurantiaca
Pseudomonas chlororaphis PCL1606
G
csp15Psy VKWFNDEKGFGFITP
csp20Psy1 SGTVKWFNDEKGFGFITPES
csp20Psy2 TGTVKWFNDEKGFGFITPQG
Pseudomonas sp. stari2
Pseudomonas benzopyrenica
Pseudomonas quebecensis
Pseudomonas defluvii
Pseudomonas batumici
Pseudomonas sp. MT-1
Pseudomonas khavaziana
Pseudomonas beijingensis
Pseudomonas sp. NC02
Pseudomonas serboccidentalis
Pseudomonas helleriensis
Pseudomonas sp. HS6
Pseudomonas sp. KUIN-1
Pseudomonas rhizophila
Pseudomonas phytophila
Pseudomonas paleroniana
Pseudomonas cucumis
Pseudomonas hormoni
Pseudomonas hygromyciniae
Pseudomonas kielensis
Pseudomonas solani
Pseudomonas nunensis
Pseudomonas fitomaticsae
Pseudomonas siliginis

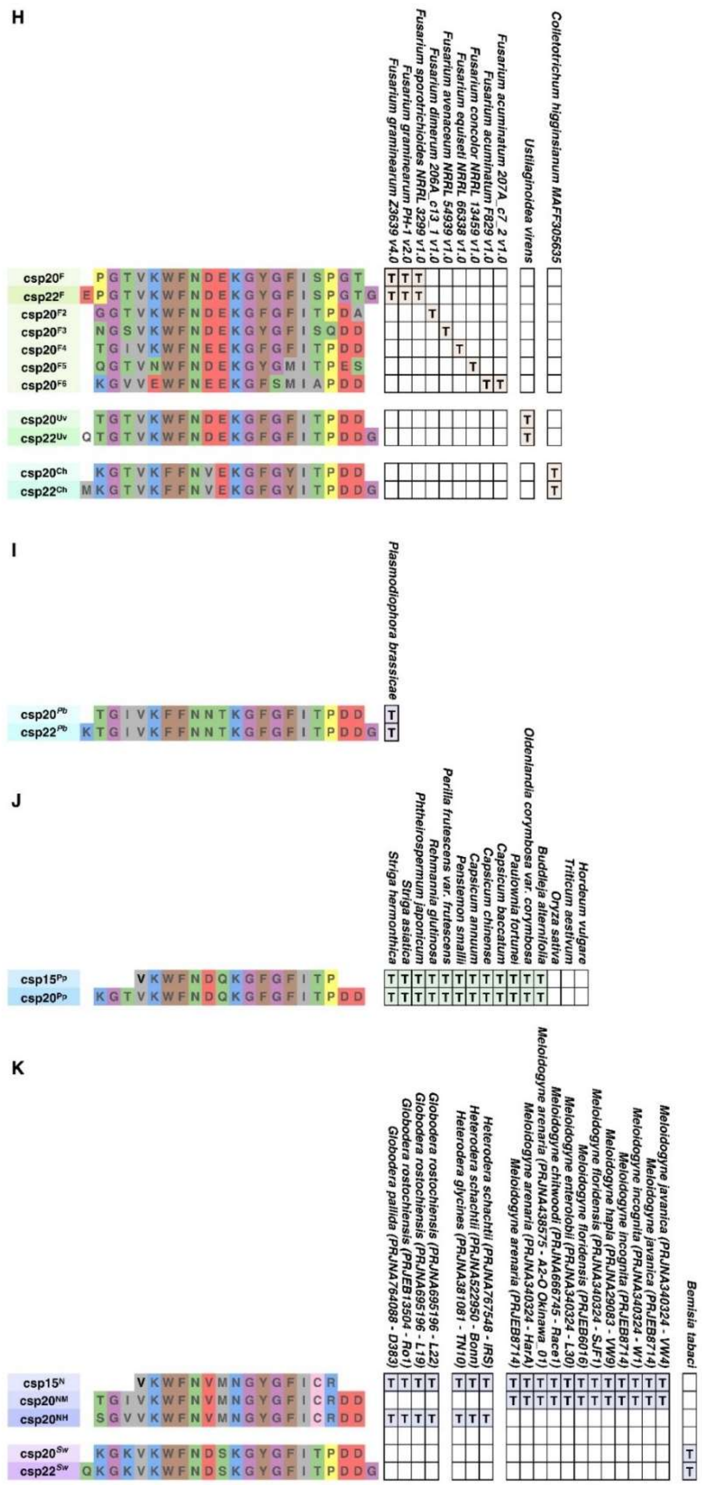

**Fig. S25. Conservation of CSP peptides in pathogens and pests.**

The names and sequences of CSP peptides from Fig 6 are listed on the left. The presence of each CSP peptide in the corresponding genome is indicated with a "T", while its absence is left blank.

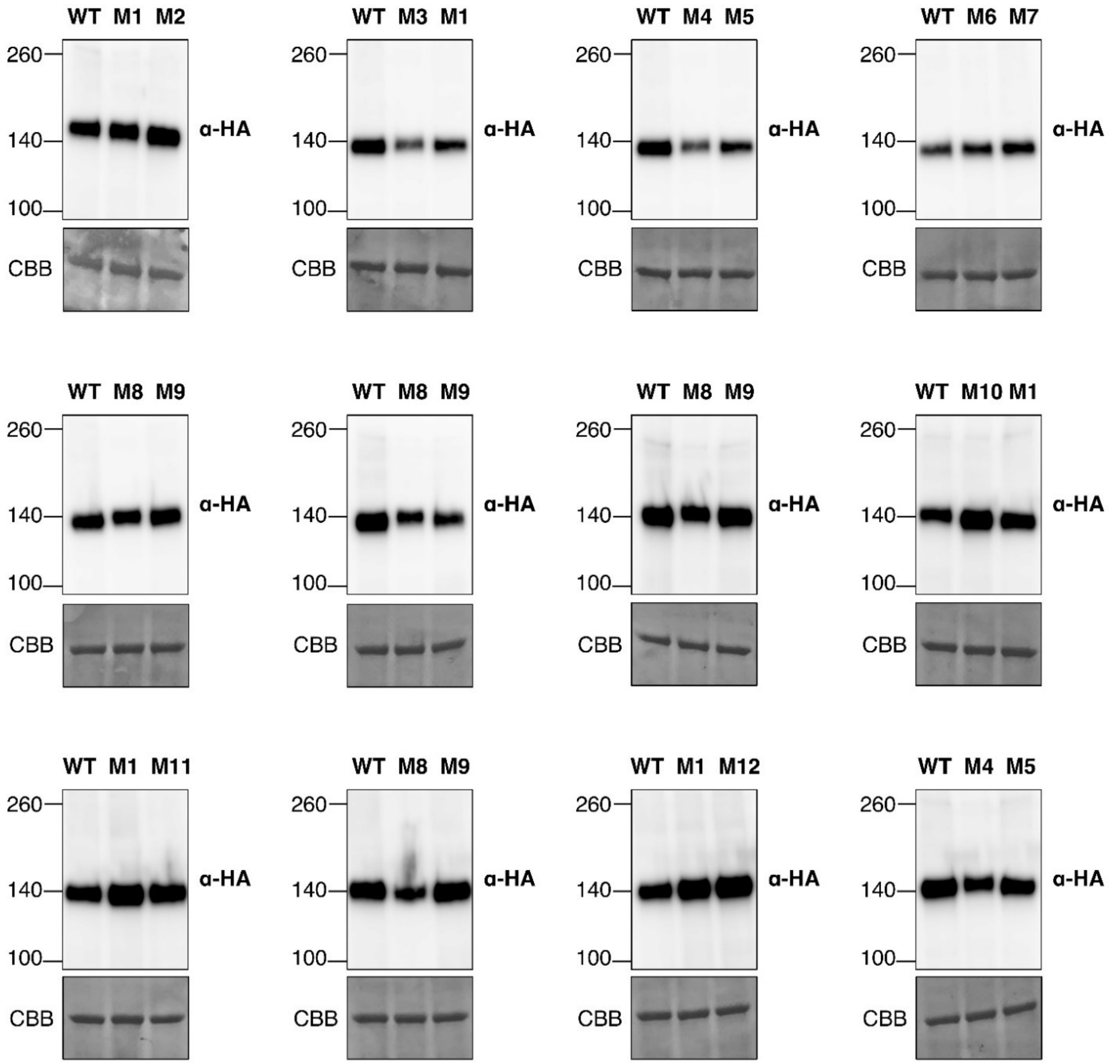


**Fig. S26. Expression of SCORE (CM) and variants in Fig 6.**

Samples used in the ROS assays in Fig 6 were analysed with immunoblotting to confirm receptor expression. CBB staining is included as loading controls.

**Data S1. (separate file)**

MS data from fractions A6-A9 in Fig 3C. The table provides detailed information on the proteins identified from fractions A6 to A9 in Fig. 3C. Protein abundance ratio for A8/A6, A9/A6, A8/A7, A9/A7 are highlighted. Green represents higher abundance and red represents lower abundance.

**Data S2. (separate file)**

Concentration of bacterial suspensions, full-length MAMPs, and peptides for each experiment.
